# Localized FLASH Radiotherapy Reduces Long-Term Skin and Muscle Damage While Preserving Systemic Homeostasis

**DOI:** 10.1101/2025.09.04.674020

**Authors:** Giulia Furini, Eduarda Mota Da Silva, Alice Usai, Gaia Scabia, Claudia Kusmic, Francesco Faita, Andrea Cavalieri, Mariagrazia Celentano, Mario Costa, Filippo Rossi, Giulia Asero, Roberta Di Pietro, Emanuela Guerra, Stefano Lattanzio, Tonia Luca, Sergio Castorina, Roberto Cusano, Riccardo Berutti, Jessica Milia, Simone Capaccioli, Alessandra Gonnelli, Noemi Giannini, Fabiola Paiar, Saverio Cinti, Fabio Di Martino, Margherita Maffei

**Author notes:** co-first author (Giulia Furini, Eduarda Mota Da Silva).

## Abstract

Radiotherapy (RT) is a cornerstone treatment for nearly 50% of cancer patients, but its curative potential and safe dosing are constrained by cumulative toxicity to surrounding healthy tissues. Delivering RT at ultra-high dose rates (FLASH-RT) represents a transformative strategy, as it appears to maintain tumor control, while sparing normal tissue. Melanoma is among the most radioresistant tumors, and skin and muscle are invariably exposed during RT, also in case of deep seated tumors. Here, we compared electron FLASH-RT and conventional RT (CONV-RT) in melanoma-bearing and naïve mice assessing tumor control, tissue integrity, and systemic homeostasis over the medium to long term. Both modalities achieved comparable tumor suppression. However, CONV-RT induced persistent skin damage, dermal fibrosis, muscle dysfunction and systemic inflammatory-metabolic alterations, while FLASH-RT largely spared normal tissue and systemic balance. Bulk RNA sequencing revealed striking differences: FLASH induced minimal transcriptional disruption in skin and muscle, whereas CONV-RT triggered thousands of differentially expressed genes, including massive activation of fibrosis, inflammation, cell death-related pathways in skin, and broad dysregulation of genes linked to muscle function, remodeling and the unfolded protein response. Histological and ultrastructural analyses corroborated the findings, showing reduced immune infiltration in the skin and preserved tissue architecture both in skin and muscle following FLASH. In conclusion our study not only confirms the protective nature of FLASH but also provides novel mechanistic insights into the cascade linking local injury to systemic dysfunction under CONV-RT, reinforcing the translational potential of FLASH to expand the therapeutic window of radiotherapy.

**One Sentence Summary:** Comparative analysis of FLASH and conventional radiotherapy in a murine model of melanoma and naïve mice - Therapeutical efficacy, local and systemic effects.

## INTRODUCTION

Radiotherapy (RT) represents one of the main pillars of modern oncology, with approximately 50–60% of patients receiving RT during the course of their disease and contributing to definitive cure in nearly 40% of cases. Despite remarkable technological advancements in dose delivery and image guidance, one of the most critical challenges remains its unavoidable toxicity to surrounding normal tissues. The skin in particular represents both a barrier and a vulnerable target: it is exposed during irradiation of superficial as well as deep-seated tumors, and radiation-induced skin damage is among the most frequent and clinically relevant adverse effects. Indeed, skin toxicity is reported in nearly 90% of patients undergoing RT, ranging from erythema and desquamation to late effects such as atrophy, fibrosis, and necrosis, with fibrosis-related dysfunction recognized as the most severe long-term complication (*1*).

Melanoma exemplifies the dual challenge of tumor radioresistance and treatment-associated toxicity. It ranks as the 17th most common cancer worldwide, with approximately 331,700 new cases in 2022, and its incidence continues to rise, particularly in Europe and Australia/New Zealand (*2*). Despite significant progress achieved through immune checkpoint inhibitors and targeted therapies, the prognosis for advanced melanoma remains poor, with a global 5-year survival rate of ∼50% (*3*). Radiotherapy, often used as part of multimodal regimens, provides local control in specific clinical scenarios; however, the intrinsic radioresistance of melanoma, coupled with frequent relapses, underscores the urgent need for novel and more effective RT approaches (*4*).

In recent years, ultra-high dose rate radiotherapy (FLASH-RT) has emerged as a promising strategy to mitigate toxicity without compromising tumor control. FLASH-RT delivers radiation at dose rates several orders of magnitude higher than conventional RT (CONV-RT), typically ≥40 Gy/s compared to ∼0.1–0.5 Gy/s. Strikingly, multiple preclinical studies have reported that FLASH-RT spares normal tissues across various organs—including brain, lung, intestine, and skin—while maintaining iso-effective tumor control. This so-called “FLASH effect” has been demonstrated in mice (*5–7*) as well as in larger animals such as minipigs and cats (*8*), suggesting broad translational potential. Importantly, the magnitude of this protective effect appears dose-dependent, with higher total doses often producing more pronounced tissue sparing (*5, 8*).

Despite these promising observations, the mechanisms underlying the FLASH effect remain poorly understood. Hypotheses include oxygen depletion during ultra-fast delivery, altered redox signaling, and differential activation of DNA damage responses (*9*). However, definitive mechanistic evidence is still lacking. Moreover, while tumor control by FLASH-RT has been shown to be comparable to CONV-RT in models of sarcoma and lung carcinoma (*10, 11*), data on melanoma remain scarce. Considering the notorious radioresistance of melanoma, this tumor represents a particularly stringent benchmark for assessing the therapeutic relevance of FLASH-RT.

Another underexplored dimension is the relationship between local irradiation and systemic responses. Radiation not only damages tissues directly within the beam path but also triggers complex systemic effects, including metabolic, endocrine, and immune alterations (*12*). These broader responses may influence both treatment tolerance and efficacy. Yet, few studies have examined how FLASH versus CONV irradiation differentially impact systemic physiology beyond the irradiated tissue. For example, changes in energy balance regulating hormones such as leptin and ghrelin, or transcriptional remodeling in non-tumor tissues, may reveal previously unrecognized dimensions of the FLASH effect.

To address these gaps, we investigated the comparative impact of electron FLASH versus CONV-RT in a syngeneic melanoma model. In addition to evaluating tumor growth control and skin toxicity—classical endpoints in FLASH research—we extended our analysis to systemic parameters, including food and water intake, body weight (BW) dynamics, endocrine profiles, and gene expression programs in healthy irradiated tissues.

For this purpose, we exploited a clinical linear accelerator (linac) modified to deliver both CONV and electron FLASH beams within the same platform (*13*), ensuring maximal comparability. This approach enabled us to characterize, in a translationally relevant setting, how different dose-rate modalities shape not only local toxicity but also systemic responses in irradiated mice.

## RESULTS

### Comparable Tumor Control of FLASH- and CONV-RT in a Murine Model of Melanoma

Mice were intradermally injected with melanoma cells into the proximal region of the left hind limb, at the level of the quadriceps muscle (day 0). Three days post-injection, tumor development was confirmed by ultrasound imaging in 100% of animals (Fig. 1A, D). Animals were then randomly assigned to one of three groups to be untreated - controls (CNT), treated with FLASH- or CONV-RT at day 5 post-injection. Between days 11 and 17 post-injection, CNT mice reached humane endpoints due to tumor volumes exceeding 1 cm³ and were euthanized. In contrast, irradiated mice exhibited delayed tumor progression and prolonged survival, with greater efficacy observed at the higher radiation dose (Fig. 1C-E) for both modalities. Given the superior tumor control achieved with 35 Gy, we focused subsequent analyses on this experimental condition.

**Fig. 1.**
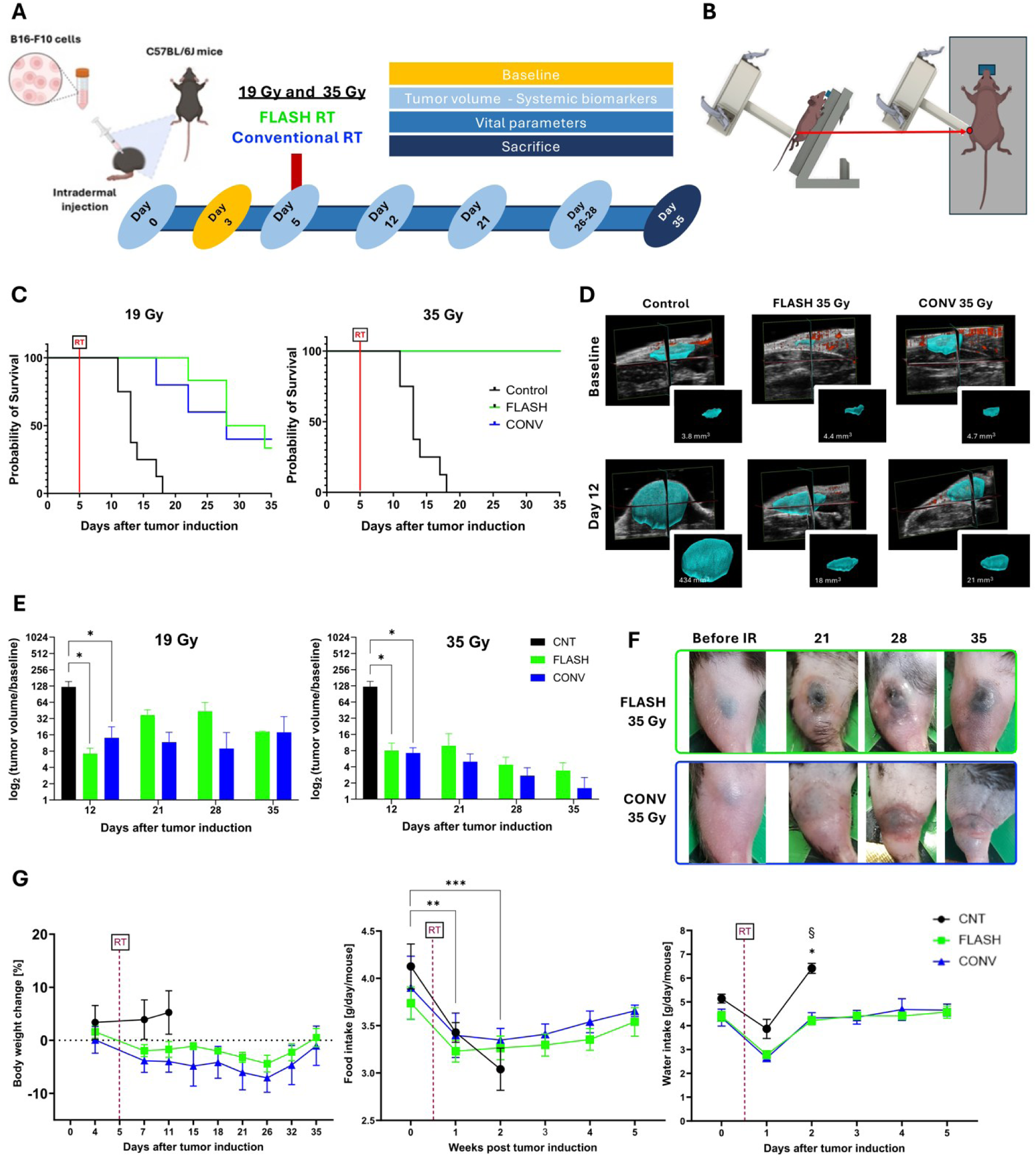
Comparative effects of FLASH- and CONV-RT on tumor progression, survival and vital parameters. **(A)** Experimental design/timeline describing the experiments performed in tumor bearing mice, see Material and Methods for further details. **(B)** Mice irradiation setup: on the left, mice positioned on a 3D-printed support. On the right, precise placement of the applicator onto the mouse thigh. **(C)** Kaplan–Meier survival curves of melanoma-bearing mice not irradiated (CNT, n = 8) or irradiated with FLASH- or CONV-RT (19 Gy, n = 5 per group; 35 Gy, n = 4 per group). All animals were sacrificed by day 35. **(D)** Representative ultrasound images of tumors treated with 35 Gy and used for volumetric analysis. **(E)** Mean tumor volume change over time for FLASH- or CONV-treated mice irradiated at 19 or 35 Gy, calculated as the ratio of each volume to its baseline measurement, taken immediately prior to RT (19 Gy, n = 5-7 per group; 35 Gy, n = 4-7 per group. 19 Gy RT fixed effect = 0.001; 35 Gy RT fixed effect = 0.0086). **(F)** Representative macroscopic images of tumor-bearing legs from two animals treated with either 35 Gy FLASH or CONV-RT as indicated. Images were taken at baseline and at days 21, 28, and 35 post-RT. **(G)** Vital parameters for CNT and mice irradiated with 35 Gy. Body weight change expressed as relative difference from baseline (time fixed effect = 0.0113), food intake (time fixed effect = 0.0009) and water intake (fixed effect: time <0.0001, RT = 0.0035). Data are presented as mean±SEM. Statistical analysis was performed using a matched mixed-effect model, followed by appropriate post-hoc tests. Symbols between groups: * = CNT vs CONV, § = CNT vs FLASH; comparisons between different timepoints in CNT as indicated by brackets; *p<0.05, **p<0.01, ***p<0.001, §p<0.05.

CNT, FLASH- and CONV-treated mice did not differ significantly in BW trajectories (Fig. 1G). In the CNT not irradiated mice, in which tumor size was rapidly reaching the humane endpoint, food intake decreased dramatically over time compared to the treated groups (Fig. 1G), while water intake increased. No notable differences were observed in circulating levels of glucose, insulin, ghrelin, or the inflammatory markers IL-6 and TNF-α (Fig. S2). Interestingly, macroscopic evaluation revealed that the healthy skin surrounding the tumor lesion appeared better preserved in FLASH-RT mice compared to those receiving CONV-RT (Fig. 1F).

Taken together, these findings support the iso-effective tumor control achieved by both RT modalities in a dose-dependent manner, while suggesting a potential protective effect of FLASH irradiation on adjacent healthy tissues.

### FLASH-RT Minimizes Damage and Transcriptional Perturbations in Skin

Given the reduced degree of skin damage observed in the peritumoral area of FLASH-treated mice, we sought to further investigate this sparing effect in a controlled setting using naïve, tumor-free animals (experimental design in Fig. 2A). 35 Gy dose was selected to assess the impact of RT on normal tissue. To more precisely evaluate radiation-induced skin toxicity, we developed a custom scoring system by integrating and refining existing methodologies (*5, 14*) to improve the resolution and consistency of damage assessment (Fig. 2B). Using this approach, we found that CONV-RT mice displayed significantly higher skin damage scores, particularly at early time points, compared to the FLASH-RT group (treatment effect p<0.01, Fig. 2C). These findings support a tissue-sparing effect of FLASH-RT on healthy skin.

**Fig. 2.**
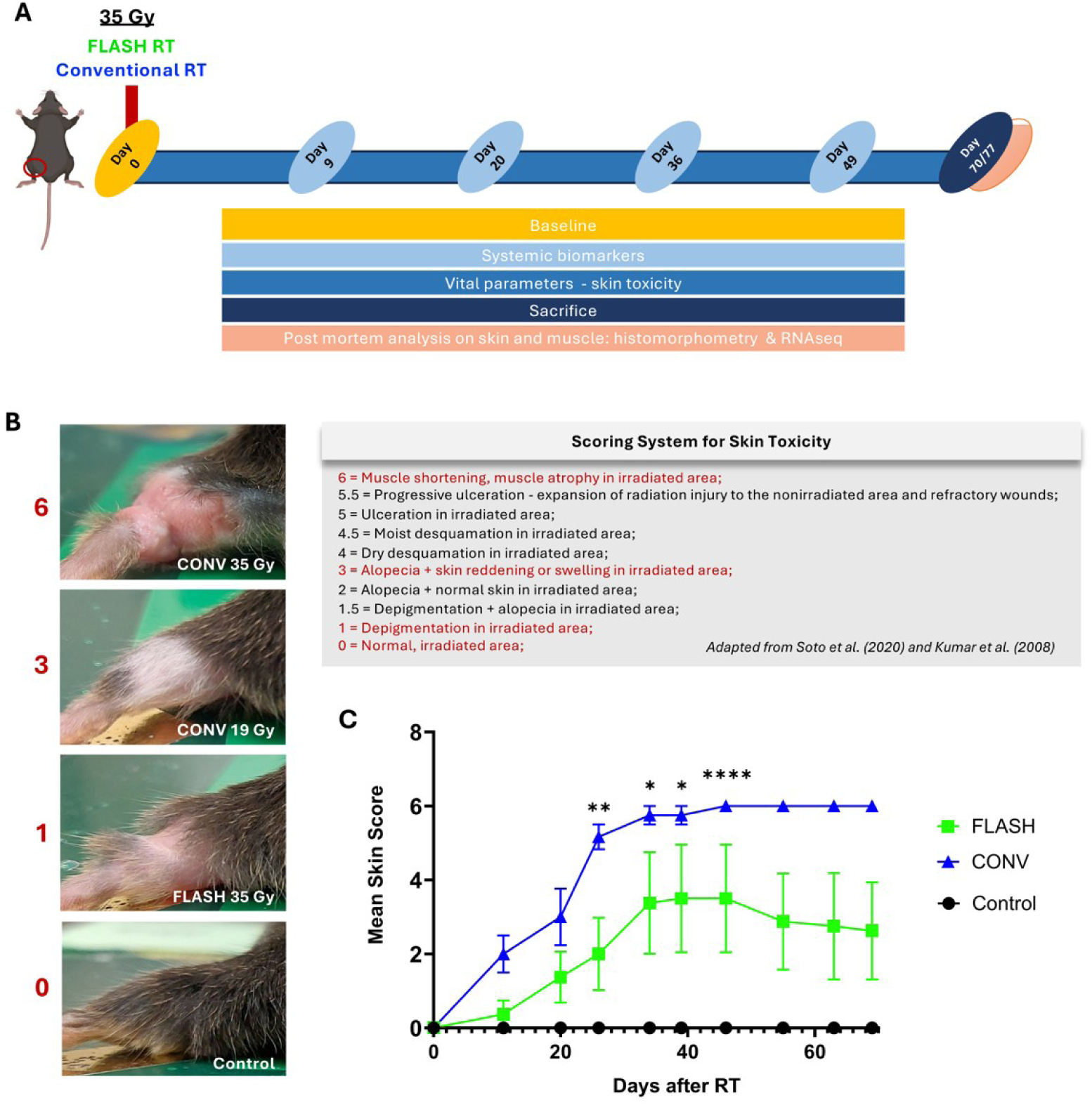
Skin toxicity in naïve mice. **(A)** Experimental design/timeline describing the experiments performed in naïve mice, see Material and Methods for further details. Mice were treated with 35 Gy CONV- or FLASH-RT to the left hindlimb; matched non-irradiated mice served as CNT. **(B)** Skin toxicity scores were assessed in naïve mice. Skin reactions were evaluated longitudinally using a semi-quantitative scoring system (0–6 scale) developed within the present project, with higher scores indicating increased severity of skin damage (see representative images in the left panel). Scores were independently and blindly assigned by three researchers using the defined 0–6 scale and averaged to obtain a final score for each mouse. **(C)** Longitudinal quantification of skin toxicity scores (fixed effect: time = 0.0181, RT = 0.0181). Data are presented as mean±SEM (n = 3-4 per group). Statistical analysis was performed using a matched mixed-effect model, followed by appropriate post-hoc tests. Symbols between groups: * = CNT vs CONV; *p<0.05, **p<0.01, ****p<0.0001.

To further investigate the molecular mechanisms underlying the FLASH effect, we performed bulk RNA sequencing (RNA-seq) on skin samples collected from mice in the CNT, FLASH-, and CONV-RT groups (Fig. 3, Figs. S3, S4, Tables S1-S4). Both FLASH- and CONV-RT samples segregated distinctly from controls (Fig. 3A, Fig. S3A), indicating that RT— regardless of dose rate—induces a measurable transcriptomic response. However, the extent of gene expression changes was markedly greater in CONV-treated mice compared to the FLASH-RT group (Fig. 3A). Using a cutoff of FDR-adjusted p<0.05 and |fold change|>1.5, we identified 2,461 differentially expressed genes (DEGs) in the CONV-RT group (1,513 upregulated and 948 downregulated) relative to CNT, whereas only 93 DEGs (42 upregulated and 51 downregulated) were detected in FLASH-treated samples (Fig. 3B). Among the common upregulated genes, to note is the presence of Keratin 6B, likely contributing to an over-keratinization of the skin and thickening of the epidermis in response to stress. Interestingly, this gene was the most upregulated, both in terms of fold change and adjusted p-value in CONV-RT (13.89; adj. p-value = 1.69×10^-69^) and in this group only was associated with other strongly upregulated keratins such as Keratin 6A, 16 and 17, known to be involved in injury-induced activation and hyperproliferation of keratinocytes (*15, 16*). Conversely, Leptin (Lep) and Perilipin 1 (Plin1) were consistently downregulated in both experimental conditions, indicating a potential reduction in the presence and function of the white adipose tissue (WAT) within the murine skin (Fig. 3B) (*17, 18*).

**Fig. 3.**
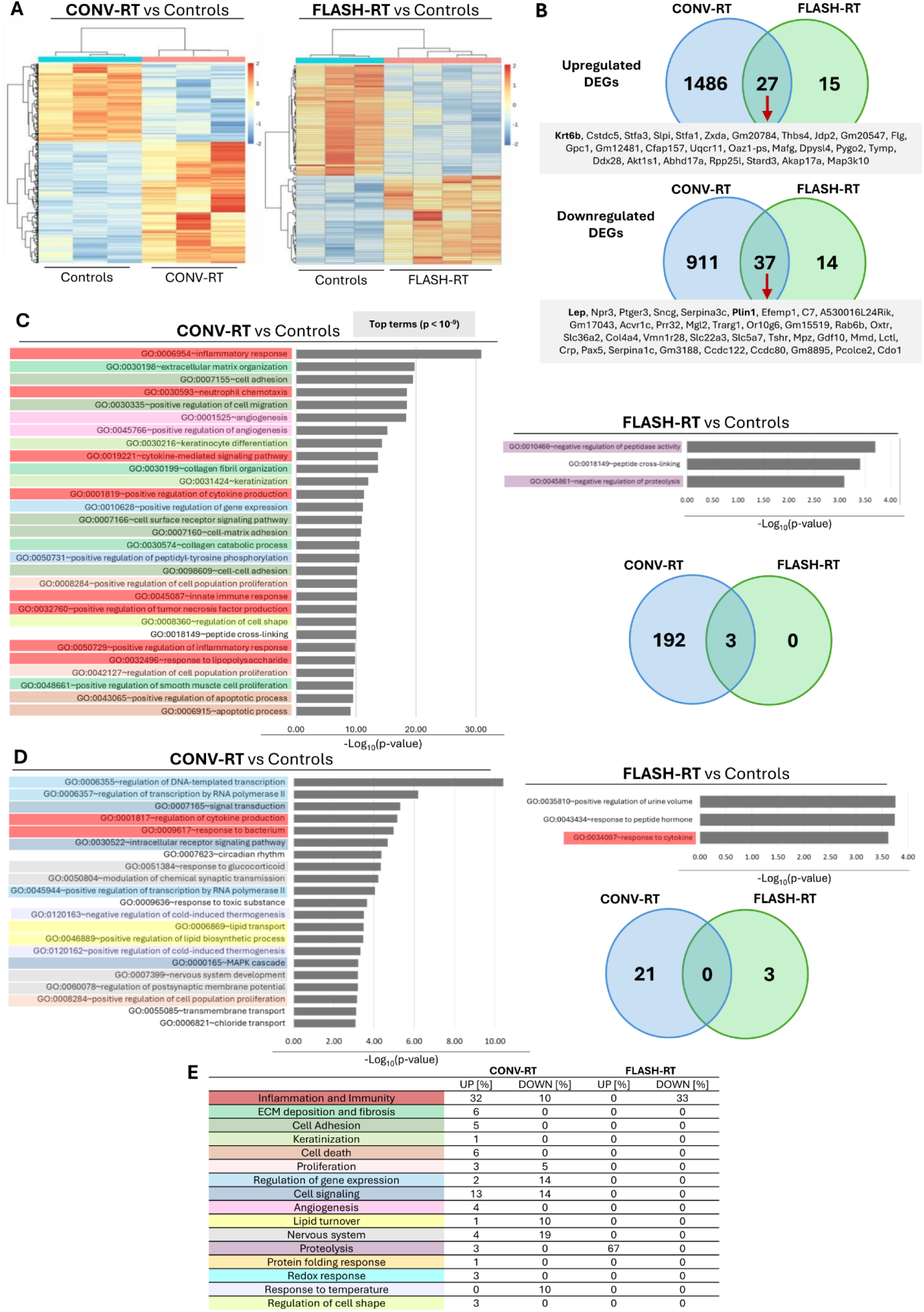
Transcriptomic analysis of left hindlimb skin from mice subjected to CONV- and FLASH-RT. RNA-seq was performed on skin samples collected from the irradiated region of the left hindlimb 70 days after mice received 35 Gy electron irradiation with either CONV-RT or FLASH-RT, as well as from matched non-irradiated CNT (n = 3-4 per group), as detailed in the Methods section. Differential gene expression analysis among irradiated and CNT groups was conducted using DESeq2. Genes with an absolute fold change>1.5 and a false discovery rate (FDR)-adjusted p-value<0.05 were considered significantly differentially expressed. **(A)** Heatmaps showing differentially expressed genes in CONV vs. CNT and FLASH vs. CNT comparisons. Color range reflects the log₂ fold change (FC) between the irradiated group and CNT. **(B)** Venn diagrams representing the number of significantly upregulated and downregulated genes, and the overlap between FLASH-RT and CONV-RT conditions. **(C–E)**. Functional enrichment analysis of Gene Ontology (GO) biological process (BP) terms among upregulated **(C)** and downregulated **(D)** genes in both CONV-RT and FLASH-RT groups (p<0.001). Bars represent enriched GO terms ranked by decreasing significance (-log10 p-value). Venn diagrams summarize the number of enriched terms identified in each analysis and their overlap. For CONV-RT upregulated genes, only the top enriched terms (p<10^-9^) are shown; see also Supplementary Figure 3 for additional significant terms. The various annotation terms collected in the figure are color-coded according to their broad categories, which are detailed in the table **(E)**; this table also includes the percentages of terms belonging to each category across the different comparisons shown.

When gene ontology (GO) analysis was applied to this dataset, 656 enriched biological process (BP) terms were identified among the genes specifically upregulated in the CONV-RT group (p<0.05), while only 6 significantly enriched terms were found in the FLASH-RT group compared to controls. A more stringent threshold for enrichment significance (p<0.001) resulted in the identification of 195 significantly enriched BP terms among the upregulated genes in the CONV-RT group, whereas only 3 BP terms remained significantly enriched in the FLASH-RT group (Fig. 3C, Fig. S3B), all of which overlapped with those enriched in the CONV-RT group. As expected, pathway analysis performed using the KEGG database reflected a similar scenario, with 81 enriched terms identified in the CONV-RT group and 2 specifics of the FLASH-RT group within the upregulated genes (p<0.05) (Fig S4A). In CONV-RT, the most prominently enriched processes and pathways were related to inflammation, fibrosis, keratinization, and cell–matrix interactions, as well as cell death and cell proliferation. Importantly, none of these themes were observed in the FLASH samples (Fig. 3C, E; Figs. S3B, S4A). Of note, the analysis of CONV-RT upregulated genes highlighted a strong enrichment in both innate and adaptive immunity and inflammatory processes and pathways, including pro-inflammatory cytokine signalling (IL-1, IL-6, TNFalpha, etc.) and leukocyte activity (Fig. 4C, E; Figs. S3B, S4A).

**Fig. 4.**
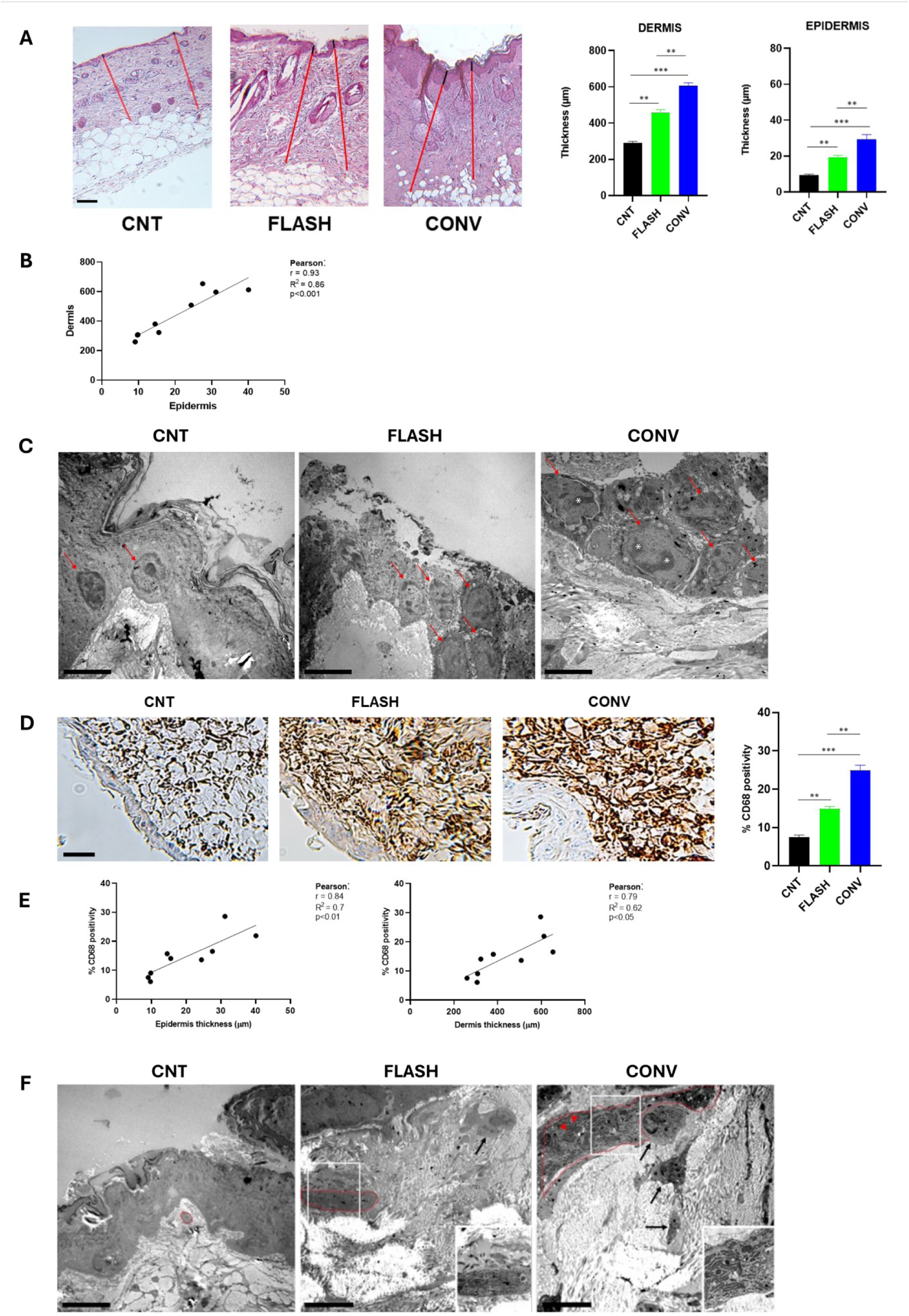
Effect of CONV- and FLASH-RT on histomorphology of healthy skin. Mice were subjected to 35 Gy CONV or FLASH-RT to the left hindlimb; matched non-irradiated mice served as CNT. Skin samples were collected 70 days post-RT upon sacrifice. **(A)** Light microscopy of hematoxylin and eosin-stained skin sections from CNT, FLASH, and CONV-treated mice with representative images of epidermal (black line) and dermal (red line) thickness. Quantification of the thickness of both skin layers is shown in the bar graphs. Scale bar: 100 µm. **(B)** Correlation analysis between epidermal and dermal thickness across all samples. **(C)** Transmission electron microscopy (TEM) images of the epidermis. Red arrows highlight nuclei of epithelial cells, which are arranged in a single layer in CNT and in multiple layers following irradiation (FLASH and CONV). Epithelial cells from irradiated skin display euchromatic nuclei. Asterisks denote prominent nucleoli, observed exclusively in the CONV-RT group, suggesting higher transcriptional activity. **(D)** Light microscopy of immunohistochemical staining for CD68 in skin sections from CNT, FLASH, and CONV-treated mice. Quantification of CD68 positivity is shown in the bar graph. Scale bar: 20 µm. **(E)** Correlation analysis between epidermal or dermal thickness and CD68 positivity. **(F)** TEM analysis of dermal tissue highlighting the inflammatory infiltrate and fibrotic remodeling. Black arrows indicate infiltrating macrophages within the dermal matrix. Red outlines mark fibroblasts of varying size and morphology, reflecting differing degrees of fibrotic response across samples. In CONV-treated mice only, fibroblasts exhibit a highly developed and dilated rough endoplasmic reticulum (RER) containing proteinaceous material (asterisks), as indicated by red arrowheads. Insets show magnified views of the boxed regions. Scale bar: 5 µm. In bar graphs, data are presented as mean ± SEM (n = 3-4 per group). Statistical analysis was performed using either one-way ANOVA or the Kruskal-Wallis test, followed by appropriate post-hoc tests; *p<0.01, **p<0.001.

Among the downregulated genes, applying the same stringent threshold described above (p<0.001), the analysis revealed 21 significantly enriched GO BP terms in CONV-RT mice, compared to only 3 in the FLASH-RT group (Fig. 3D). Similarly, 19 KEGG pathways were enriched in CONV-treated mice and 4 in FLASH-treated mice (p<0.05, Fig. S3B). Two KEGG pathways, “Regulation of Lipolysis in Adipocytes” and “Hormone Signalling” were enriched in both the CONV-RT and FLASH-RT groups (*19, 20*). As mentioned above, the downregulation of these genes is likely due to the intradermal WAT. In the case of downregulated genes, recurrent themes represented specifically in the CONV-RT enriched BP GO terms were lipid metabolism, gene expression regulation and neuron related function. Real-time quantitative PCR was used to validate the differential expression of selected genes identified by RNA-seq (Fig. S5).

Collectively, the data indicate that CONV-RT induces a more extensive transcriptional response in skin tissue compared to FLASH-RT, further supporting the notion of a tissue-sparing effect of FLASH.

### Sparing Effect of FLASH-RT on Skin Architecture and Histomorphology

To seek independent confirmation of molecular findings, we performed histological analyses. As shown in Fig. 4A, optical microscopy revealed thickening of the skin superficial layer under both irradiation conditions, but significantly more pronounced in the CONV-RT group, reflecting epidermal proliferation and increased collagen deposition. These changes were supported by morphometric analysis (Fig. 4A) and, consistently with a similar response to RT of the different skin compartments, we found a positive correlation between dermal and epidermal thickness (Fig. 4B). TEM data further corroborated these results: in unirradiated controls, TEM showed keratinocyte nuclei arranged in a single layer, consistent with normal squamous epithelium (Fig. 4C). In contrast, irradiated samples exhibited a multilayered epidermis composed of cells with enlarged euchromatic nuclei and hypertrophic nucleoli, indicative of enhanced transcriptional activity particularly pronounced in CONV-treated mice (Fig. 4C). This altered architecture suggests radiation-induced disruption of epidermal homeostasis, likely due to increased turnover and impaired differentiation, especially following CONV-RT exposure. TEM analysis of irradiated samples also revealed enlarged activated dermal fibroblasts with a well-developed endoplasmic reticulum, more prominent in CONV-compared to FLASH-RT. This aspect is associated with the elevated levels of collagen synthesis and secretion observed in irradiated skin (Fig. 4F).

We investigated the inflammatory infiltrate in skin samples and found that, while FLASH-irradiated skin was relatively spared, CONV-irradiated skin showed enhanced staining for the macrophage-specific marker CD68. Morphometry showed that the number of macrophages were significantly higher in dermis of mice irradiated by CONV-RT (Fig. 4D). The number of CD68 positive cells correlated positively with epidermis and dermis thickness (Fig. 4E). In line with these data, TEM analysis revealed the presence of numerous active macrophages in the extracellular matrix of irradiated dermal layer, indicating the presence of an inflammatory response (Fig. 4F). This immune cell infiltration suggests activation of the innate immune response following radiation exposure, contributing to tissue remodeling and repair.

All together, these findings point to a more robust and potentially damaging tissue response in CONV-irradiated skin compared to the relatively spared profile observed with FLASH-RT: major signatures of this damage include fibrosis and inflammation.

### CONV-RT, but not FLASH-RT, Impacts Vital Parameters and Circulating Metabolic and Inflammatory Markers

We next extended our analysis to evaluate the general health status of the animals. FLASH-RT and CNT groups exhibited overlapping survival curves with undefined median survival times, whereas three mice in the CONV-RT group were euthanized due to extensive lesions in the irradiated left leg, with one case showing lesion extension to off-target areas such as the posterior back and tail region (Fig. 5A). BW change was not different among the three groups, while both food and water intake were influenced by RT with CONV-treated mice displaying the highest values during the medium-term phase after irradiation (Fig. 5B). Glucose, on the other hand, tended to be lower in CONV-compared to FLASH-treated mice (Fig. 5C).

**Fig. 5.**
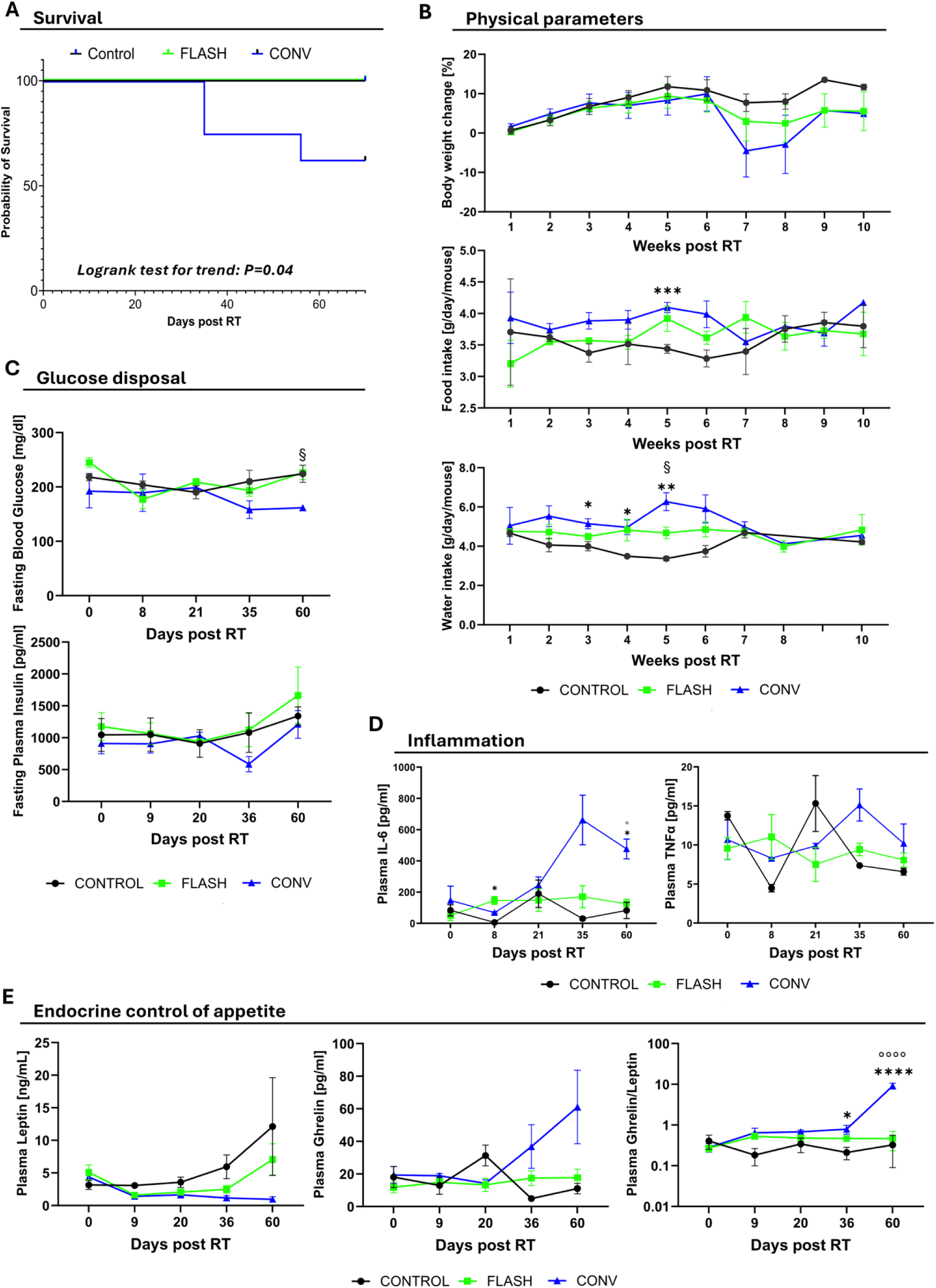
Overview of systemic effects in irradiated mice. Mice received 35 Gy of electron RT to the left hindlimb using either CONV or FLASH delivery. Non-irradiated animals served CNT. **(A)** Kaplan– Meier survival curves. Statistical analysis was performed using the Log-rank test for trend. **(B)** Longitudinal assessment of body weight expressed as percent change from pre-irradiation baseline (time fixed effect = 0.0044), food intake (RT fixed effect = 0.0181), and water intake (RT fixed effect = 0.0003). **(C)** Fasting blood glucose and insulin levels measured over time via tail vein sampling. **(D)** Longitudinal analysis of circulating pro-inflammatory cytokines TNF-α (time x RT fixed effect = 0.0158) and IL-6 (fixed effect: RT = 0.0077, time x RT = 0.008). **(E)** Circulating metabolic hormones involved in appetite regulation, ghrelin (fixed effect: RT = 0.0045; time x RT = 0.0042) and leptin (fixed effect: RT = 0.0116; time x RT = 0.0015). A starvation index was calculated as the plasma ghrelin-to-leptin ratio (source of variation: time <0.0001, RT <0.0001, time x RT <0.0001). Data are presented as mean±SEM (n = 3–8 mice / group). Statistical analysis was performed using two-way repeated measures ANOVA or a matched mixed-effects model, as appropriate, followed by post-hoc tests (comparisons between groups: * = CNT vs CONV; § = CNT vs FLASH; ° = FLASH vs CONV); *p<0.05, ** p<0.01, *** p< 0.001, ****p<0.0001, § p<0.05, ° p<0.05.

We also measured circulating markers related to inflammation and the endocrine regulation of energy balance. In CONV-RT mice there was a treatment dependent increase over time for circulating IL-6 (p<0.01, Fig. 5D). We then evaluated appetite-related hormones: leptin was reduced, most markedly in CONV-treated mice (RT fixed effect, p<0.05), whereas ghrelin showed an opposite trend, progressively increasing over time in the CONV-RT group only (RT fixed effect, p<0.01) (Fig. 5E). This endocrine profile is consistent with a systemic signal promoting enhanced appetite. To capture this imbalance, we calculated an “appetite index” (ghrelin/leptin ratio) as a proxy of perceived starvation. Notably, CONV-RT mice displayed persistently higher values compared to CNT and FLASH-RT, with differences becoming highly significant over time (Fig. 5E).

Overall, these data suggest that CONV-RT administered to the hindlimb not only affects the irradiated target area but also induces systemic alterations. This effect appears to be markedly reduced—or even absent—when FLASH-RT is used.

### Muscle Architecture and Signaling of the Irradiated Area are Impacted by CONV-, but not FLASH-RT

In the presence of a starvation condition—whether actual or merely perceived—the organism responds by preserving energy and sparing functions or tissues not deemed essential for survival. Among the known responses to starvation, muscle alterations have been described (*21, 22*). Analysis of semithin sections from irradiated left quadriceps revealed that muscle fibers in CONV-treated mice exhibited markedly irregular shapes compared to those from CNT or FLASH-treated mice, suggesting more pronounced structural damage upon CONV-RT (Fig. 6A). A progressively fainter staining from CNT to CONV-RT can also be noticed, this being usually associated with tissue damage. TEM analyses revealed the presence of tubular aggregates only in the FLASH-treated muscle fibers. These aggregates are typically associated with stress-related protein misfolding events and are a marker of reversible damage (*23*) (Fig. 6B). On the other hand, ultrastructural alterations that were either more severe or exclusively present following CONV-RT included nucleolar enlargement within myonuclei, mitochondrial hyperplasia and hypertrophy, widespread sarcomere disorganization, myofibrillar fragmentation, and disruption of the Z-line (Fig. 6B, panels 3, 6). Morphometric analysis of TEM images further reinforced these observations. A significant reduction in myofibrillar thickness together with the increase in inter-myofibrils spaces was documented across irradiated groups, with the most evident effects in the CONV-irradiated samples (Fig. 6B, panels 5, 6 and Fig. 6C). Given that myofibrillar density and thickness are critical determinants of contractile function, these data point to a greater degree of functional impairment following CONV irradiation.

**Figure 6.**
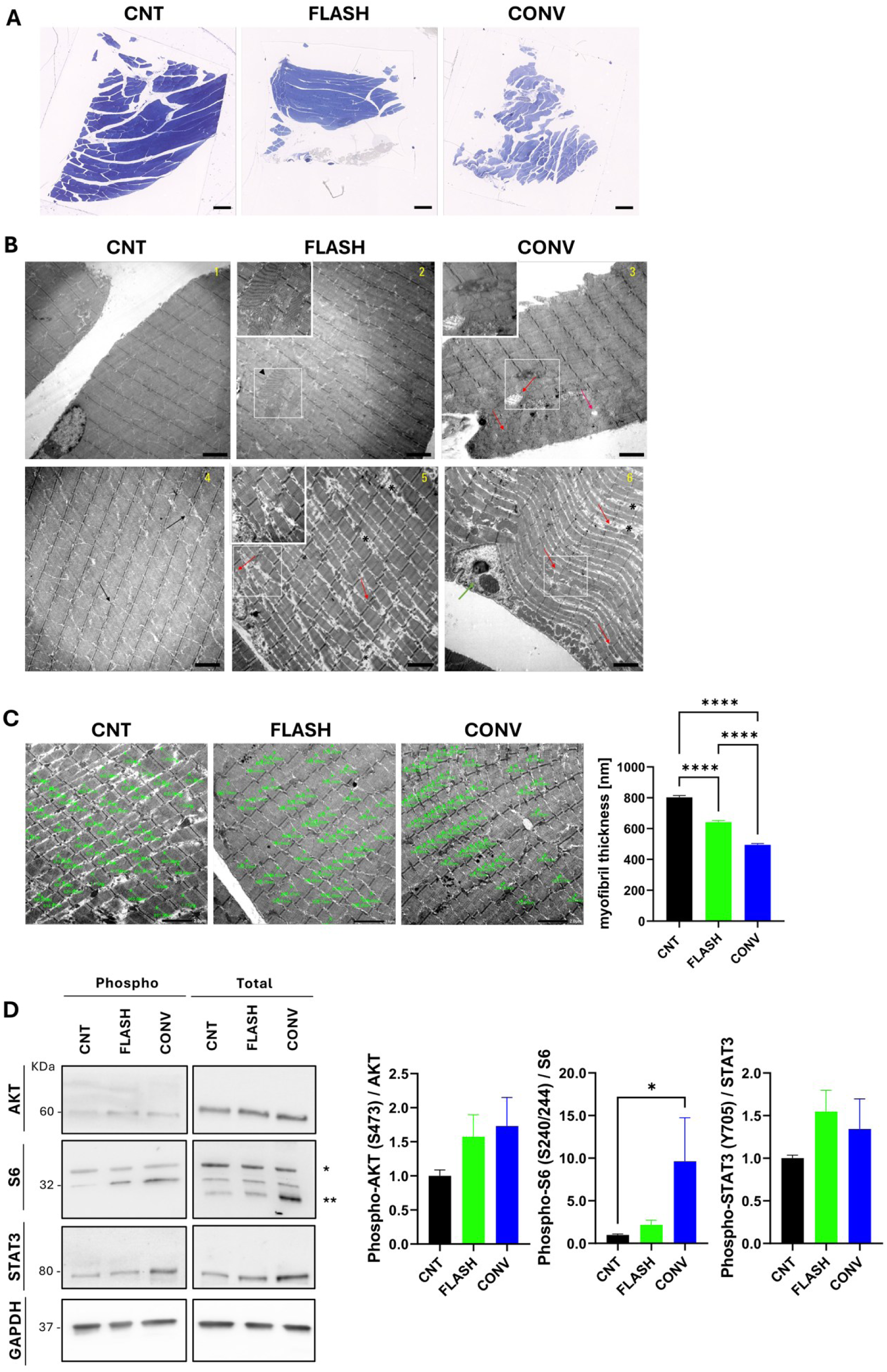
– Effects of FLASH- and CONV-RT on healthy skeletal muscle ultrastructure and signaling pathways. Mice were subjected to 35 Gy CONV or FLASH-RT to the left hindlimb; matched non-irradiated mice served as CNT. Skeletal muscle (*vastus lateralis*) samples were collected 70 days post-RT **(A)** Semithin sections of vastus lateralis muscle stained with toluidine blue. Representative images show irregular fiber morphology and regions of fainter staining, indicative of tissue damage, most pronounced in CONV-treated mice. Scale bar: 200 µm. **(B)** TEM of ultrathin sections stained with Uranyless and Lead Citrate. Black arrows: intact mitochondria (CNT, panel 4); Black arrowhead: tubular aggregates (FLASH-RT, panel 2); Red arrows: areas of mitochondrial hyperplasia and structural damage (predominantly in CONV-RT, panels 3, 5, 6); Green arrows: multiple nucleoli suggesting transcriptional activation (CONV-RT, panel 6); Black asterisks: enlarged intermyofibrillar sarcoplasmic spaces (more evident in CONV-RT, panels 5, 6). Inset boxes show magnified views. Scale bar: 2 µm. **(C)** Morphometric analysis of myofibril thickness at the M line on longitudinal skeletal muscle sections. Right panel shows the results of the morphometric analysis. Each field contained 30–50 myofibrils; three fields per sample were analyzed. Scale bar: 2 µm. **(D)** Western blot analysis of phosphorylated AKT, S6 ribosomal protein, and STAT3 in muscle lysates. The ratio of phosphorylated to total protein was calculated after normalization to GAPDH. In bar graphs, data are presented as mean±SEM (n = 3-4 per group). One-way ANOVA or Kruskal-Wallis test was performed as appropriate, followed by post-hoc tests; * p<0.05; **** p<0.001.

Collectively, these findings underscore the differential biological impact of CONV-RT and FLASH-RT on skeletal muscle integrity. While FLASH-RT appears to largely preserve muscle architecture, including the maintenance of myofibrils organization and subcellular structure, CONV-RT is associated with extensive architectural disruption, increased inter-myofibrillar spaces, severe damage at both the structural and ultrastructural levels and stress-induced increase in transcriptional activity and metabolic demand.

Muscle homeostasis is maintained by a finely tuned balance between protein synthesis and protein degradation, primarily regulated by signaling pathways such as mTOR/AKT for anabolic responses and STAT3 for catabolic control. RT can disrupt this balance, triggering compensatory or pathological adaptations. In irradiated mice, we observed a slight not significant increase in AKT activation, as measured by the phospho/total AKT ratio (Fig. 6D). Interestingly, total AKT protein levels were significantly upregulated in CONV-treated mice, but not in the FLASH-RT group (Fig. S6A), suggesting that the induction occurs at the transcriptional and/or translational level. We also detected increased activation of S6, a key downstream effector of the mTOR pathway and central regulator of protein synthesis (Fig. 6D). These findings may reflect a compensatory anabolic response aimed at counteracting prior or ongoing muscle fiber loss, as supported by the structural alterations observed in both histological and ultrastructural analyses. This observation is also consistent with the increased transcriptional activity mentioned above (Figure 6B). STAT3, a known upstream regulator of the ubiquitin–proteasome system, showed mild activation in irradiated muscles, regardless of the RT modality (Fig. 6D), indicating that a catabolic response might also be involved in the muscle loss observed in our model.

### Effects of FLASH- and CONV-RT on Skeletal Muscle Molecular Signature

Bulk RNA-seq performed on left quadriceps skeletal muscle revealed a transcriptional landscape closely resembling that observed in skin samples, with irradiated muscle transcriptomes clearly segregating from those of controls (Fig. 7A, Tables S5-S8). As in the skin, the comparison between CONV-RT and CNT mice yielded a markedly higher number of DEGs than the FLASH-RT versus CNT comparison (Fig. 7B). This divergence was clearly reflected in GO and KEGG enrichment analysis (Fig. 7C-E, Fig. S7). In the CONV group, 93 BPs were significantly enriched among the upregulated DEGs. The most prominently represented broad categories include lipid metabolism and turnover, muscle tissue remodeling, neuronal function and unfolded protein response, one of the upstream inducers of protein degradation (Fig. 7C-E, Fig S7A). Of note, response to starvation was also represented among significantly enriched process in CONV-RT mice lending support to the above-mentioned hypothesis of a perceived status of negative energy balance. Conversely, as for downregulated transcripts, the most significantly enriched BPs were associated with skeletal muscle structure and function (Fig. 7D). In the FLASH-RT group upregulated BPs were mainly related to altered inflammatory/immune profile, while no BP process was enriched among the downregulated genes (Fig. 7C-D).

**Figure 7.**
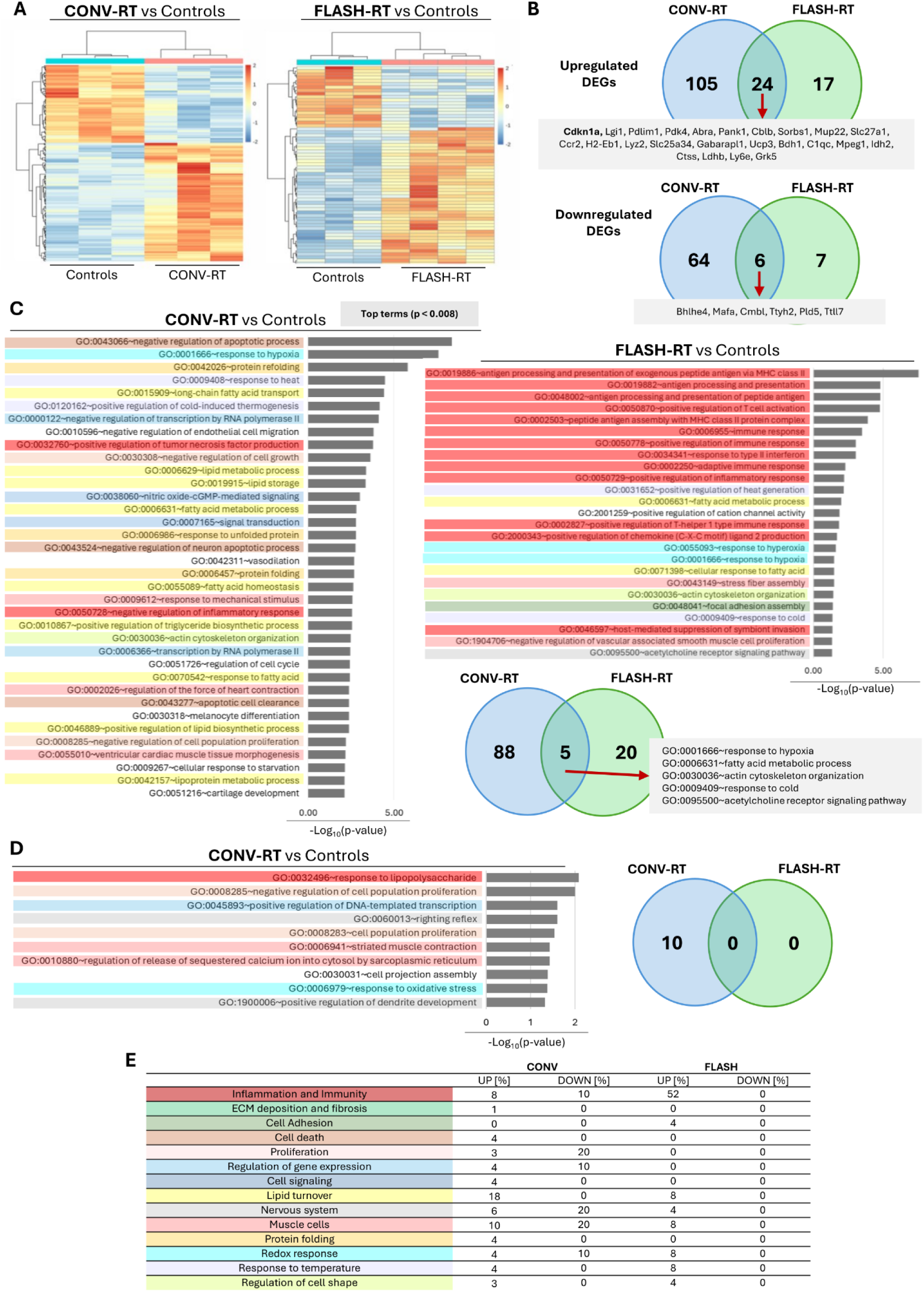
-Transcriptomic analysis of left hindlimb muscle from mice subjected to conventional and FLASH RT. RNA sequencing (RNA-seq) was performed on muscle samples (*vastus lateralis)* collected from the irradiated region of the left hindlimb 70 days after mice received 35 Gy electron irradiation with either CONV-RT or FLASH-RT as well as from matched non-irradiated CNT (n = 3-4 per group), as detailed in the Methods section. Differential gene expression analysis among irradiated and control groups was conducted using DESeq2. Genes with an absolute fold change > 1.5 and a false discovery rate (FDR)-adjusted p-value < 0.05 were considered significantly differentially expressed. **(A)** Heatmaps showing differentially expressed genes in CONV-RT vs. control and FLASH-RT vs. control comparisons. Color range reflects the log₂ fold change (FC) between the irradiated group and controls. **(B)** Venn diagrams representing the number of significantly upregulated and downregulated genes, and the overlap between FLASH-RT and CONV-RT conditions. **(C–E)**. Functional enrichment analysis of Gene Ontology (GO) biological process (BP) terms among upregulated **(C)** and downregulated **(D)** genes in both CONV-RT and FLASH-RT groups (p<0.05). Bars represent enriched GO terms ranked by decreasing significance (-log10 p-value). Venn diagrams summarize the number of enriched terms identified in each analysis and their overlap. The various annotation terms collected in the figure are color-coded according to their broad categories, which are detailed in the table **(E)**; this table also includes the percentages of terms belonging to each category across the different comparisons shown.

## DISCUSSION

In this study, we adopted a comprehensive preclinical approach to evaluate the dual objectives of tumor control in a murine melanoma model and healthy tissue preservation following electron FLASH-RT. Clinically speaking, FLASH-RT demonstrated iso-efficacy with CONV-RT in reducing melanoma tumor burden, while markedly limiting damage to surrounding healthy tissues—including skin, underlying skeletal muscle, and systemic metabolic and inflammatory parameters.

In line with prior reports, we found that FLASH-RT and CONV-RT achieved iso-effective tumor control in B16-F10 melanoma-bearing mice, with dose-dependent improvements in survival and tumor growth delay. A comparable tumor control between FLASH- and CONV-RT has been reported both with electron beams, as in the murine model of lung cancer (*10*), and with protons, as in two leg-injected sarcoma models (*11*). For the first time, our study demonstrates that the iso-efficacy of CONV and FLASH electron RT also applies to a syngeneic melanoma model. The higher dose (35 Gy) resulted in more prolonged tumor growth inhibition, yet complete remission was not observed in any of the experimental groups. This is consistent with other reports, where tumor complete eradication was absent (*11*). The known radioresistance of melanoma (*24*) therefore offered a stringent model to demonstrate the iso-efficacy of FLASH-relative to CONV-RT. Of note, proton FLASH in combination with immunotherapy has proven effective in increasing survival in mice bearing melanoma (*25*).

Interestingly, control mice bearing untreated tumors exhibited a progressive decline in food intake accompanied by an increase in water consumption, particularly near endpoint, when euthanasia was required. BW changes did not parallel food intake as tumor burden (0.7–1 g at sacrifice) contributed substantially (≈4–5%) to total mass. The increased water intake observed has been occasionally reported in cancer-bearing subjects (*26, 27*) although its etiology remains unclear.

One of the most striking differences between modalities was observed in healthy skin, where FLASH-RT resulted in markedly reduced acute and chronic damage. Using skin toxicity as primary endpoint, previous studies, employing both electron- and proton-source, report a FLASH effect with substantial sparing of healthy skin, in murine (*5–7*) as well as in larger animal models such as cats and minipigs (*8*), with higher doses better highlighting the protective effect. The 35 Gy dose used here lies within the effective range reported by Soto (*5*) (30–40 Gy) and Vozenin (*8*) (27–40 Gy). In our study, light microscopy revealed marked skin thickening after irradiation, particularly in the CONV group, closely associated with inflammatory infiltration, supporting the established link between epidermal proliferation, fibrosis, and inflammation (*28*). Ultrastructural analyses consistently indicated pronounced fibrosis, inflammatory infiltration, and keratinocyte hyperproliferation more pronounced upon CONV-RT. Transcriptomic analyses reinforced these findings: CONV-RT induced thousands of DEGs, particularly in pathways linked to inflammation, fibrosis, keratinization, apoptosis and extracellular matrix remodeling, whereas FLASH-RT altered only a limited set of transcripts, indicating a more restrained and physiologically contained response. Results partly similar have been reported with proton FLASH-RT: in one study at 30 Gy, skin transcriptomics 5 days post-irradiation revealed strong upregulation of apoptotic signaling and keratin pathways in CONV-RT (*11*). Interestingly, they reported enrichment of tissue repair pathways uniquely in the FLASH group, whereas we did not observe this feature, likely reflecting both differences in radiation source (protons vs. electrons) and timing of analysis, since our later endpoint (70 days post-irradiation) may have captured a stage when active repair had already subsided. Interestingly, in the present study both modalities downregulated leptin and perilipin1, indicating an effect on dermal white adipose tissue (*18*). However, the type and intensity of transcriptional changes were far greater with CONV-RT, supporting the view that FLASH-RT not only reduces the magnitude of tissue injury but also prevents the initiation of extensive inflammatory cascades and fibrosis. This is important clinically, as radiation dermatitis is a common and often dose-limiting toxicity in cancer patients (*29, 30*) and fibrosis may lead to a multitude of pathologies (*31*).

Beyond the irradiated field, we observed striking differences in systemic physiology. CONV-RT, but not FLASH-RT, triggered widespread systemic alterations, including increased circulating IL-6, reduced leptin, elevated ghrelin, and a persistently high ghrelin/leptin ratio. This profile is reminiscent of a starvation-like state (*32*), consistent with decreased glucose levels and increased food intake. Such systemic perturbations can compromise patient well-being during therapy and potentially exacerbate cancer-associated cachexia (*33*). In contrast, FLASH-treated animals maintained metabolic and inflammatory profiles similar to controls, suggesting that FLASH irradiation spares not only local tissue integrity but also systemic regulation. These results highlight a critical point: while radiotherapy is often considered a localized treatment, its systemic consequences can be profound, and radiotherapy-induced cachexia has been previously described in patients (*34*). The ability of FLASH to minimize these systemic side effects could represent a transformative advantage, improving patient quality of life and treatment tolerance. During perceived starvation and/or cachexia, the body initiates multiple compensatory mechanisms to preserve the homeostasis of vital functions. Among these, one of the most prominent is skeletal muscle catabolism (*35*), which serves to release amino acids for gluconeogenesis. Indeed, impairments in muscle structure and altered expression profile were prominent in the muscle of CONV-treated mice. Structural analyses confirmed these deficits, with CONV samples showing disrupted myofibrillar organization and muscle atrophy in line with what previously reported (*11*). Additionally, we observed mitochondrial abnormalities and widespread cytoskeletal damage. FLASH-treated muscles, by contrast, largely retained normal architecture, with only reversible tubular aggregates observed. At the molecular level, RNA sequencing again underscored the differential impact of the two modalities. CONV-RT induced widespread transcriptional remodeling, with enrichment of biological processes linked to starvation responses, lipid metabolism, unfolded protein response, and muscle remodeling. Notably, genes associated with muscle structural integrity were significantly downregulated. FLASH-RT, on the other hand, produced only minor transcriptional changes, primarily involving immune-related pathways with a substantial preservation of muscle homeostasis and prevention of maladaptive remodeling as that triggered by CONV irradiation.

Limitations of the study include the use of a single murine tumor model, a single irradiation site, and single-fraction high-dose delivery, limiting generalizability. Analyses were focused on skin and skeletal muscle, with other organs at risk not examined. Transcriptomic assessment at a single late time point may have missed early dynamic changes, and mechanistic conclusions remain inferential rather than directly tested. Differences in radiation source, fractionation, and timing may also affect outcomes and translation to humans.

In conclusion, this work not only confirms that FLASH-RT achieves tumor control comparable to CONV-RT while sparing normal tissue, systemic homeostasis, and muscle integrity, but also provides novel mechanistic insights into how this protection is achieved. Transcriptomic, histological and functional analysis collectively demonstrate that FLASH triggers a uniquely restrained biological response, sharply reducing the early wave of radiation-induced signaling cascades, systemic stress-like metabolic alterations, and muscle oxidative damage that characterize CONV-RT. This mechanistic understanding highlights the novelty of FLASH as more than a protective modality: it represents a fundamentally different biological response that could expand the therapeutic window of radiotherapy.

## MATERIALS AND METHODS

### Experimental Design

This study aims to investigate the effects of electron CONV-RT versus ultra-high dose FLASH-RT in both a syngeneic melanoma model and a naïve model, evaluating its impact on tumor progression and healthy tissues potentially affected by the therapy, as well as its systemic effects. RT was delivered by a novel linear accelerator (Linac) specifically designed to enable reproducible and finely tunable control of electron beam parameters. The energy of the delivered dose remains relatively uniform within the first 1–2 cm from the surface, enabling us to investigate the biological effects of both CONV- and FLASH-RT not only in the skin but also in the underlying muscle tissue.

C57BL/6J mice were randomly assigned to either the melanoma (N=4-8 mice per group) or the naïve model (n = 5-8 mice per group) (Fig. 1A, 2A). In the former, mice were inoculated with B16-F10 cells in the left hindlimb and received irradiation (19 Gy or 35 Gy) five days later. In the naïve model, mice received 35 Gy irradiation to the same limb 1–2 days after being housed. For consistency throughout the manuscript, “Day 0” is defined as the day of tumor inoculation in the melanoma model and the day of RT in the naïve model. Melanoma mice were monitored for up to 35 days, while naïve mice were sacrificed 70 days post-RT to assess mid- and long-term effects. Animals reaching humane endpoints, due to either tumor burden or general health deterioration, were euthanized earlier.

Survival, physical parameters, glucose regulation, and systemic levels of inflammatory markers and metabolic hormones were evaluated in both models longitudinally from day 0 to sacrifice. Ultrasound analysis was performed to evaluate tumor progression in the melanoma model. In the naïve model we investigated the impact of CONV versus FLASH-RT on skin and muscle tissues exposed to RT by high-throughput gene expression profiling, histological analysis using both optical and electron microscopy. In addition, for the skin we scored toxicity using a scale devised for this study and for muscle we evaluated motor and strength deficits. Data analysis was performed blindly.

### Animals

Male C57BL/6J mice (6–7 weeks old) were purchased from Charles River Laboratories Italia SRL. Following a 2-week acclimatization period in the animal facility, animals were housed individually (melanoma tumor model) or in groups of 3–4 per cage (naïve group). Food and water were available ad libitum. Body weight (BW) was monitored at baseline and then weekly throughout the experimental period. Food and water intake were recorded twice per week. At the time of sacrifice, skin and muscle tissues from the irradiated area were dissected and either snap-frozen in liquid nitrogen for RNA and/or protein extraction or formalin-fixed for histological analysis.

All procedures were carried out in accordance with the Helsinki guidelines for animal experimentation and approved under Authorized Protocol 1006/2023-PR, dated November 28, 2023.

### Irradiation Study Design

All the irradiations were performed at the Centro Pisano FLASH Radiotherapy (CPFR) using the ElectronFlash linear accelerator (*13*), which delivers electron beams of 7 or 9 MeV with the possibility of modifying the average dose-rate (ADR) and dose per pulse (DPP) by varying the e-beam current and the pulse repetition frequency (PRF). This allows to switch the irradiation characteristics from the conventional condition (CONV) to the “FLASH” condition, while keeping the experimental setup. All set parameters are summarized in Table 1.

**Table 1.** Radiotherapy experimental parameters: relevant parameters used in each irradiation condition.

|  | Number of pulses | DPP [Gy] | Total dose [Gy] | Pulse duration [μs] | PRF [Hz] | ADR [Gy/s] |
| --- | --- | --- | --- | --- | --- | --- |
| FLASH-RT | 4 | 4.75 | 19 | 4 | 145 | 940 |
|  | 7 | 4.75 | 33.3 | 4 | 170 | 940 |
| CONV-RT | 103 | 0.19 | 19 | 4 | 1 | 0.18 |
|  | 180 | 0.19 | 33.3 | 4 | 1 | 0.18 |

DPP, dose per pulse; PRF, pulse repetition frequency, ADR, average dose rate

All irradiations were performed on anesthetized mice (ketamine 100 mg/kg / xylazine 10 mg/kg, i.p.) using a collimator with 10 mm diameter centered on the animal’s left thigh, where the tumoral lesion was located in the case of tumor bearing mice. This choice allows the precise positioning of the electron beam on the tumoral lesion and the maximum spare of surrounding tissues, thanks to the homogeneous irradiated area (7 mm diameter). As shown in Fig. 1B, the animal was fixed by its teeth on a 3D printed support perpendicular to the applicator, so that the positioning of each animal would be reproducible.

The dosimetric characterization of our beams was carried out using the fD detector (*36*) and and measurement other methods, developed at CPFR and/ or provided by other centers (*37–45*). An extended description can be found in the Supplementary Materials.

### In Vivo Xenograft Melanoma Tumor Model

B16-F10 murine melanoma cells were cultured in standard conditions according to ATCC protocol. C57BL/6 were anesthetized using volatile isoflurane (induction: 2.5%, maintenance: 1.5%; flow 1 l/min), and the left leg was depilated at the thigh level with shaving cream. A sterile 1 mL syringe fitted with a 26G needle was used to slowly inject a cell suspension (2×10^5^ B16-F10 cells in 50 µL of phosphate-buffered saline) intradermally into the leg at an approximately 10° angle relative to the skin surface. After the procedure, animals were allowed to recover in their cages.

### 3D-Ultrasound measurement of tumor volume

Tumor volumes were measured at several time points along the study by ultrasound using a Vevo 3100 system (FUJIFILM VisualSonics Inc.) as described in the Supplementary materials. 3D – mode scans (70-150 μm intervals) were imported into the VevoLab software (FUJIFILM VisualSonics Inc.), where the melanoma tumors were identified and semi-automatically marked in the cross-sectional 2D images. Tumor volumes were calculated using the volumetric analysis tool in VevoLab, based on multiple outlined tumor perimeters.

### Blood collection and systemic analyses

Blood samples were collected from the tail vein after a six-hour fast to measure glycemia and plasma protein levels, both at baseline (Day 0) and at multiple timepoints throughout the experimental procedure. Glucose levels were measured using a OneTouch glucometer (LifeScan). For plasma collection, blood was drawn into EDTA-coated tubes and centrifuged in a refrigerated microcentrifuge (2,000×g, 10 min, 4 °C). The plasma fraction was then stored at −80 °C until analysis. Plasma leptin levels were measured using a Quantikine ELISA Kit (#MOB00B, R&D Systems). Plasma levels of TNF-α, IL-6, ghrelin, and insulin were quantified using the Milliplex Extended Hormone Panel (#MMHE-44K, Merck) Both assays were performed following the manufacturer’s protocol.

### RNA and Protein extraction and analysis from mouse tissues

Homogenization of frozen mice tissues (10-25 mg) for RNA and protein isolation was performed with a Qiagen TissueLyser II in 700 µL of TRIzol reagent (#15596-018, Thermo Fisher Scientific) or 300 µL lysis buffer (50 mM TRIS-HCl, pH 7.4, 150 mM NaCl, 10 mM EDTA, 1% v/v Triton X-100, 10% v/v glycerol) supplemented with protease and phosphatase inhibitors (1:100, #P2850, #P5726, #P8340, Sigma Aldrich), using stainless steel beads at 30 Hz for 5×30 sec.

mRNA amount of specific genes was evaluated by quantitative PCR as fold increase on TATA Binding Protein gene (TBP) expression. Protein levels of phosphorylated and total S6 ribosomal protein, AKT and STAT3 were quantified by Western Blot analysis and normalized for GAPDH expression. Detailed procedures and reagents employed are available in the Supplementary Materials.

### Transcriptomic analysis

Total RNA was assessed for quantity and quality (RIN ≥ 7.5) prior to library preparation. Ribosomal RNA was depleted, and strand-specific libraries were prepared, individually indexed, and pooled for paired-end sequencing on an Illumina NovaSeq X Plus platform. Sequencing reads were processed using the in-house RiDE pipeline (https://github.com/solida-core/ride), developed at CRS4, which includes alignment, quantification, and quality control steps. Differential expression analysis was performed using DESeq2 (*46*), followed by functional enrichment analysis with DAVID (https://davidbioinformatics.nih.gov/). Full details of the RNA-seq and bioinformatics analyses are provided in the Supplementary Materials. RNA-seq datasets are available under accession number GSEXXXX in the Gene Expression Omnibus (NCBI).

### Light microscopy

Dorsal skin samples from the irradiated area of the left hindlimb were processed for histological analysis as described by Castorina et al. (2021) (*47*). Epidermal and dermal thickness were assessed on hematoxylin and eosin (H&E)-stained sections, while macrophage infiltration was evaluated by immunohistochemistry for CD68. Tissue sections were examined under a light microscope (Zeiss Axioskop 40; Carl Zeiss GmbH), and digital images were acquired using a Zeiss Axiocam 503 color camera. Additional details are provided in Supplementary Materials.

### Semithin-sections light microscopy and Transmission electron microscopy

Murine skin and muscle samples, approximately 1 mm² in size, were fixed in 2.5% gluteraldeide and processed for subsequent microscopy analysis as extensively described in the Supplementary Materials. Semithin (800 nm) and Ultrathin sections (∼80 nm) were obtained from resin-embedded tissues. Semithin sections were stained with 1% toluidine blue and 1% sodium tetraborate and imaged using a NanoZoomer-SQ digital slide scanner (Hamamatsu Photonics) for high-resolution light microscopy analysis. Ultrathin sections were collected on copper grids and counterstained with UranyLess and lead citrate. TEM imaging was performed with a JEOL JAM-1400 Flash microscope (120 kV), and morphometric analysis of myofibril thickness at the M-line and Z-line was conducted using JEOL SightX Viewer 2.1 software.

### Statistical analysis

Graphical representation and statistical analyses were performed using GraphPad Prism. Data are presented as mean±SEM. Normality was assessed using the Shapiro–Wilk test, and homogeneity of variances using the Brown–Forsythe test. Depending on the experimental design, statistical comparisons were conducted using either one-way ANOVA (or its non-parametric equivalent, the Kruskal–Wallis test) or two-way repeated measures ANOVA with Geisser– Greenhouse correction. When assumptions for repeated measures ANOVA were violated or data were unbalanced, a mixed-effects model was applied. Where appropriate, multiple comparisons were adjusted using Tukey’s post hoc test. Survival analyses were performed using Kaplan–Meier survival curves, and group comparisons were assessed with the log-rank test for trend. Correlation analyses were carried out using Pearson’s correlation coefficient (r) to evaluate the strength and direction of linear relationships between variables, with associated p values reported for statistical significance. All statistical tests were two-tailed, and a pvalue<0.05 was considered statistically significant.

## List of Supplementary Materials

Materials and Methods

Fig. S1 to S6

Tables S1 to S9

References (*50–53*)

## Acknowledgments

we acknowledge Fondazione Pisa for funding CPFR with the grant “prog. n.134/2021”. A special thanks to Emma Buzzigoli for technical assistance and to Cecilia Ciampi, Sara Ciampi and Gianpietro Chessa for the daily management of animals.

## Funding

Piano Nazionale di Ripresa e Resilienza (PNRR), Missione 4, Componente 2, Ecosistemi dell’Innovazione–Tuscany Health Ecosystem (THE), Spoke 1 “Advanced Radiotherapies and Diagnostics in Oncology”—CUP I53C22000780001 (MM) and cascading call financed grant RADAR.

PNRR-MAD-2022-12376459, CUP B53C22009530001 (MM)

MIRO - MInibeam RadiOtherapy Program: INFN CSN5 Call 2024-2026 (MM)

## Author contributions

Conceptualization: EMDS, SC, FDM, MM

Methodology: GF, EMDS, AU, GS, CK, FF, AC, MGC, MC, FR, GA, RDP, EG, SL, TL, RC, JM, RB, SCi, FP, FDM, MM, AG, NG

Investigation: GF, EMDS, GS, FR, SL, TL, MM

Visualization: GF, EMDS, GS, CK, RDP, SL, TL, SCi, MM

Funding acquisition: SCas, SCap, FDM, MM

Project administration: FDM, MM

Supervision: CK, FF, MC, SCi, FDM, MM

Writing – original draft: GF, MM

Writing – review & editing: GF, EMDS, AU, GS, CK, AC, MGC, RDP, SL, TL, RC, JM, RB, Sci, MM

Validation: GF, GS, MC, Sci, MM

Resources: SCas, SCap, FP, FDM, MM

Formal analysis: GF, GS, CK, FF, AC, MGC, FR, RDP, EG, SL, TL, RC, JM, RB, MM

Data curation: GF, EMDS, GS, MM

## Data and materials availability

Data submission to the Gene Expression Omnibus (GEO) repository is in progress.

