## Supplementary Material, Figures and Tables for "Localized FLASH Radiotherapy Reduces Long-Term Skin and Muscle Damage While Preserving Systemic Homeostasis"

#### Supplementary Materials and Methods

**Ultrasound imaging of tumor volumes**

Ultrasound imaging was performed using a Vevo 3100 system (FUJIFILM VisualSonics Inc., Toronto, Canada) while the animals were under isoflurane anesthesia (induction: 2.5%, maintenance: 1.5%; flow 1 l/min). The left hindlimbs were shaved with depilatory cream, and the animals were secured on a heated stand. Temperature, respiration, and heart rate were monitored during the imaging process (THM-100; Indus Instruments, Houston, USA). A MS550 probe (22-55 MHz) was coupled to the mice with acoustic gel and used for all scans. B-mode and Power Doppler-mode images of the hindlimb tumor were acquired in short- and long-axis views with the animals in ventral recumbency, properly aligned to the target. Additionally, 3D-mode scans of the melanoma tumor were obtained at 70-150 µm intervals.

**RNA Isolation, cDNA synthesis and quantitative PCR**

Total RNA was extracted from 10–15 mg of frozen mouse skin and muscle tissues, collected from the hindlimb of both control and irradiated mice. After homogenization in TRIzol reagent, samples were processed for the isolation of RNA. The aqueous phase obtained after chloroform addition and centrifugation (12000 × *g* for 15 min at 4 °C) was loaded onto Direct-zol RNA Microprep columns (Zymo Research, # R2060) and RNA eluted in 15 uL RNAse-free water, as indicated by the manufacturer protocol. cDNA was synthesized using iScript Reverse Transcription Supermix for RT-qPCR (#1708840, Bio-Rad Laboratories), and quantitative real-time PCR (qPCR) was carried out using either probe-based (iTaq Universal Probes Supermix, Bio-Rad Laboratories) or SYBR Green-based (iTaq Universal SYBR Green Supermix, Bio-Rad Laboratories) systems with CFX96TM Real-Time instrument (Bio-Rad Laboratories). Probes were acquired from Applied Biosystems and exon junction-spanning primers were designed using PrimerBLAST and acquired from Eurofins. Details are displayed in Table S9 The relative amount of mRNA for CCL2, IL1b, Krt6a, Krt6b, ADAM8, BCL3, CDKN1A/p21 and PAI1/serpine1 genes was calculated as fold increase on TATA Binding Protein gene (TBP) expression (internal control). Data were obtained using ΔΔ−Ct method.

**Protein Isolation and Western Blot Analysis**

Total proteins were extracted from 22–25 mg of frozen hindlimb muscle tissue (vastus lateralis) of control and irradiated mice. After homogenization, protein lysates were cleared by centrifugation (10,000 x g, 10 min, 4°C) and concentrations were determined by Bradford Assay (#5000001, Bio-Rad Laboratories) according to the manufacturer's instructions. Equal amounts of protein (30 µg) were separated by SDS-PAGE on 4–15% Mini-PROTEAN® TGX Stain-Free™ Protein Gels (#4568083, Bio-Rad Laboratories) and transferred to nitrocellulose membranes using the Trans-Blot® Turbo™ Transfer System (#1704159, Bio-Rad Laboratories). Membranes were blocked with EveryBlot™ Blocking Buffer (#12010020, Bio-Rad Laboratories) for 30 min at room temperature, then incubated overnight at 4°C with the following primary antibodies from Cell Signaling Technology in blocking buffer: phospho-S6 (#5364, 1:1000), S6 (#2317, 1:1000), phospho-AKT (#4058, 1:1000), AKT (#9272, 1:1000), phospho-STAT3 (#9145, 1:1000), STAT3 (#9139, 1:500) and GAPDH (#2118, 1:3000). After washing with TBS-T (Tris-buffered saline containing 0.1% Tween-20), membranes were incubated with HRP-conjugated secondary antibodies from Bio-Rad Laboratories: Goat Anti-Rabbit IgG (H+L)-HRP (#1706515, 1:3000) and Goat Anti-Mouse IgG (H+L)-HRP (#1706516, 1:3000). Immunoreactive bands were detected using an enhanced chemiluminescence system (Clarity™ Western ECL Substrate, #1705060, Bio-Rad Laboratories). Densitometric analysis of the bands was performed using ImageJ software, and protein expression levels were normalized to GAPDH.

#### RNA Sequencing

*RNA evaluation* - Qualitative and quantitative RNA evaluations were performed by NanoDrop ND-1000 Spectrophotometer (ThermoFisher). To determine the integrity of RNA samples, check controls were performed by Agilent Bioanalyzer using RNA 6000 LabChip ® kit (#5067-1511) or Agilent Fragment Analyzer 5300 using RNA (15nt) kit (#5191-6572). All the samples with a RIN below 7.5 were discarded while the others were processed for libraries preparation and sequencing.

*Library preparation* - Total RNA samples (200 ng) were processed using the Illumina Stranded Total RNA Prep with Ribo-Zero Plus kit (#20040529). Ribosomal RNA (rRNA) was depleted following the manufacturer's *Sample Preparation Guide* (Document #1000000124514, version 03). Purified RNA was chemically fragmented using divalent cations under high temperature in Illumina proprietary fragmentation buffer. First strand cDNA was synthesized using random oligonucleotides and Reverse Transcriptase Enzyme and second strand cDNA synthesis was subsequently performed using DNA Polymerase I and RNase H.

After purification with Agencourt AMPure XP beads (#A63882, Beckman), DNA fragments were adenylated in their 3' ends, then Illumina–RNA Index Anchor (#20040899) adapters were ligated.

Library enrichment and indexing were carried out with Illumina RNA Unique Dual Indexes (#20091655) and PCR Mastermix in a 13 cycles amplification reaction. Finally, library products were purified with AMPure XP beads, quantified using both Tecan Infinite F200-PRO instrument and Qubit dsDNA BR reagent (#Q32853, ThermoFisher) and checked for quality on the Agilent 2100 Bioanalyzer using Agilent DNA 1000 assay (#5067-1504) or Agilent Fragment Analyzer 5300 and Double Strand DNA 910/915 (#5191-6595).

Individual indexed libraries were pooled to achieve equimolar concentrations of each sample. Specifically, 50 ng of library for each sample were collected in a vial, then the final pool was quantified using the Qubit dsDNA BR kit and the molarity was calculated considering an average library size of 340 bp.

*Sequencing on NovaSeq X Plus* - Pooled libraries were loaded and sequenced on Illumina NovaSeq X Plus using 1.5B Reagent Kit (300 Cycles) (#20104705) at 150pM final loading concentration. At the end of the run, around 140M of 151bp paired-end reads per sample were used for the bioinformatic analysis.

#### Bioinformatic analysis

*Demultiplexing* - On-board DRAGEN BCL Convert tool was used to demultiplex samples, generate individual datasets in the form of separate FASTQ files for reads 1 and 2, and to trim adapter sequences.

*Single sample pipeline* - The overall bioinformatic pipeline run for the samples is based on RiDE [https://github.com/solida-core/ride (Atzeni R, Massidda M, Cuccuru GM, Uva P)], part of the solida-core workflow collection [https://github.com/solida-core], developed at CRS4. The pipeline performs alignment, quantification, and quality controls.

*Alignment* - Reads in FASTQ format were aligned with STAR 2.5.3a [Dobin A et al, STAR: ultrafast universal RNA-seq aligner, Bioinformatics, OUP, 2013 ] using mouse reference UCSC Genome Browser mm10 *(Kent et al., 2002)*, [mm10 mouse reference: https://genome.ucsc.edu/cgi-bin/hgGateway?db=mm10]. As supporting gene track, the UCSC GenomeBrowser knownGene in GTF format was downloaded and adapted to the project aim.

*Quantification* - Transcript abundance was estimated with Kallisto 0.43.0 *(Bray et al., 2016)*, which identifies the transcript sequences using a FASTA file representing known cDNA sequences. The cDNA sequences used have been obtained from Ensembl Genome Browser, with HGNC gene symbols.

*Quality Control* - Quality control metrics were obtained for each sample with the software RSeQC *(Wang et al., 2012)*, version 5.0.3.

*Differential Expression analysis*- Differentially expressed (DE) genes were identified using the R package DeSeq2 *(Love et al., 2014)* using a FDR corrected p-value<0.05. In house scripts have been developed to evaluate the appropriate fold-change (FC) threshold and balance between p-value and FC.

*Enrichment analysis* - Functional enrichment analysis of differentially expressed genes was performed using the DAVID Bioinformatics Resources (Database for Annotation, Visualization and Integrated Discovery; https://davidbioinformatics.nih.gov/ ) *(Huang et al., 2009)*. Separate lists of upregulated and downregulated genes, annotated with official gene symbols, were uploaded with *Mus musculus* selected as the background species. Enrichment analysis was conducted specifically for Gene Ontology Biological Process (BP - direct) terms and KEGG pathways. Annotation terms containing at least two genes and with a p-value<0.05 were considered significant.

**Light microscopy**

*Sample preparation* - tissue samples were fixed in 4% paraformaldehyde in 0.1 M phosphate buffer at pH 7.4 for 24 h, progressively dehydrated in increasing concentrations of alcohol, clarified in xylene and finally embedded in paraffin as described by Castorina et al. *(Castorina et al., 2021)*. From each sample, sections 3 µm thick were cut on a sliding microtome (Leica RM 2135, Leica Microsystem) and stained with haematoxylin and eosin (H&E) for routine use.

*Immunoistochemistry* - For immunohistochemical analysis, paraffin-embedded tissue sections were processed using the BenchMark ULTRA system (Roche Tissue Diagnostics), following the manufacturer's instructions. The automated procedure included antigen retrieval with Ventana Ultra CC1 buffer (Roche Tissue Diagnostics), followed by incubation with the CONFIRM anti-CD68 (KP-1) primary antibody (Roche Tissue Diagnostics). Detection was performed using the ultraView Universal DAB Detection Kit (Roche Tissue Diagnostics). Finally, hematoxylin counterstaining and mounting were performed.

*Histological Imaging and Quantitative Analysis* - Tissue sections were examined using a light microscope (Zeiss Axioskop 40; Carl Zeiss GmbH), and digital images were captured with a Zeiss Axiocam 503 color camera for subsequent analysis of epidermal and dermal thickness, as well as CD68 protein expression. To assess tissue thickness, at least four fields of well-preserved tissue were randomly selected from each sample and imaged at ×2.5 magnification. Epidermal thickness was measured from the surface epithelium to the top of the dermal papillae, while dermal thickness was measured from the base of the dermal papillae to the underlying dermal white adipose tissue. For CD68 expression analysis, at least four fields of well-preserved tissue were randomly selected from each sample and imaged at ×20 magnification. Semi-automated image analysis was performed using ImageJ software. Images were converted to grayscale, and red-stained areas (indicating CD68 positivity) were selected using the threshold tool. CD68 expression was quantified as the percentage of CD68-positive area relative to the total selected tissue area.

**Semithin and Transmission Electron Microscopy**

Murine skin and muscle samples, approximately 1 mm² in size, were fixed in 2.5% glutaraldehyde (Electron Microscopy Sciences) in 0.1 M sodium cacodylate buffer (pH 7.4) at 4°C for a minimum of 2 hours. Samples were then washed with 0.1 M sodium cacodylate buffer (pH 7.4) to remove glutaraldehyde and post-fixed in 1% osmium tetroxide (Electron Microscopy Sciences) for 1 hour at room temperature. After rinsing with 0.1 M sodium cacodylate buffer (pH 7.4), samples were dehydrated through a graded acetone series (40%, 60%, and 100%) and infiltrated with resin/acetone mixtures at ratios of 1:1 and 2:1, followed by embedding in pure epoxy resin (Electron Microscopy Sciences) and polymerization at 60 °C for 24 hours.

Semithin sections (800 nm) were obtained using a Powertome X ultramicrotome (RCM Boeckeler) equipped with a 2.5 mm diamond knife (Diatome) and stained on glass slides with 1% toluidine blue and 1% sodium tetraborate (Electron Microscopy Sciences). The stained sections were scanned using a NanoZoomer-SQ (Hamamatsu Photonics, Hamamatsu, Japan) and analyzed with NDP.view2 software (Hamamatsu Photonics).

Ultrathin sections (~80 nm) were obtained using an 8 mm diamond knife (Diatome) and collected on 200-mesh copper grids (Electron Microscopy Sciences). Sections were counterstained with UranyLess (Electron Microscopy Sciences), followed by lead citrate (Electron Microscopy Sciences). Transmission electron microscopy (TEM) analysis was performed using a JEOL JAM-1400 Flash microscope operating at 120 kV and equipped with a high-sensitivity sCMOS camera (JEOL, Tokyo, Japan). Morphometric analysis of murine muscle myofibril thickness at both M-line and Z-line level was carried out using JEOL SightX Viewer 2.1 software (JEOL).

#### Supplementary Figures

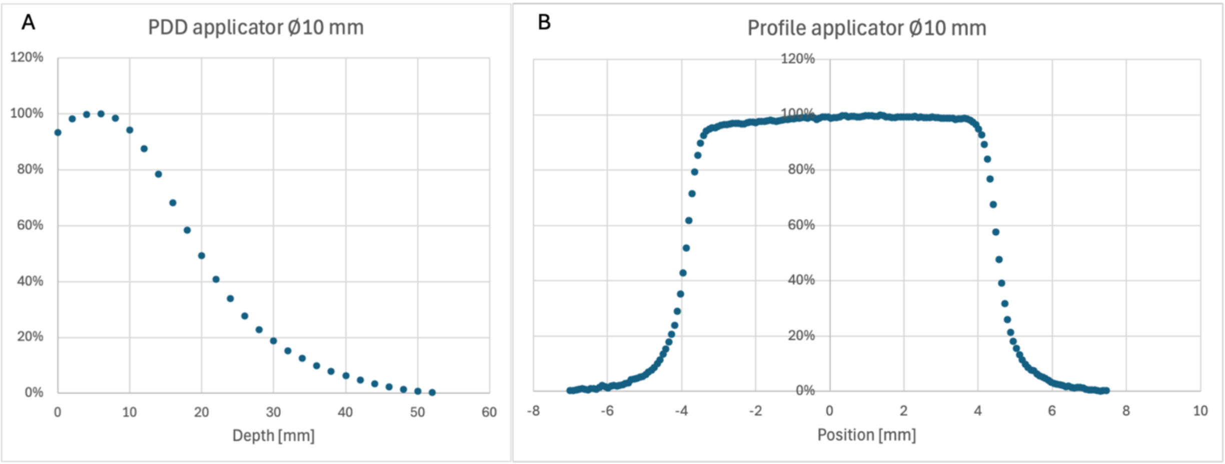


**Fig. S1. Applicator characterization.** A) Percentage-depth-dose (PDD) and B) lateral profile of Æ10mm PMMA applicator used in all irradiations (FLASH and CONV).

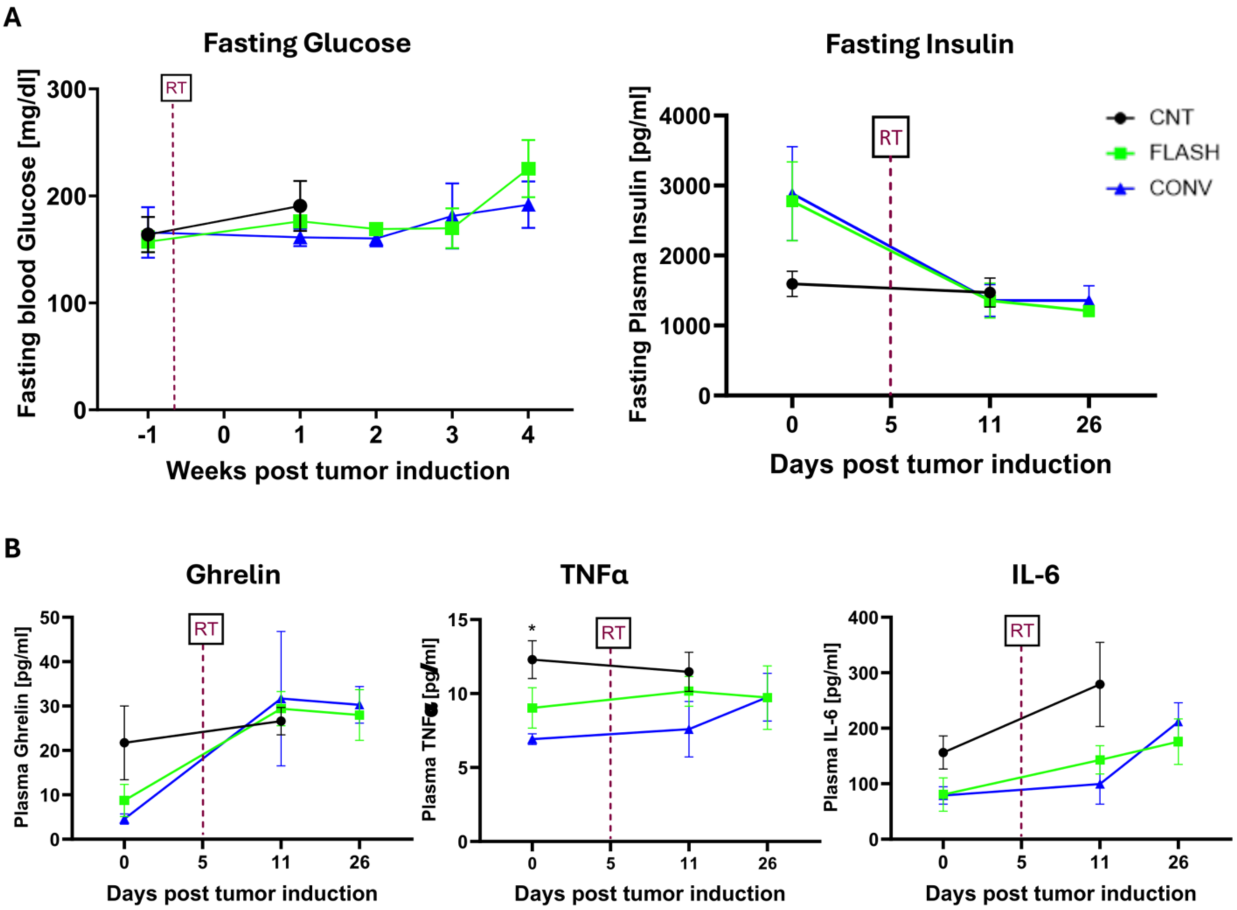


**Fig. S2. Systemic effects of conventional (CONV) and FLASH radiotherapy in the melanoma mouse model.** Mice were injected with melanoma cells in the left hindlimb and received either 35 Gy conventional (CONV) or FLASH- RT five days post-injection as described in the Materials and Methods section. A matched group of non-irradiated tumor-bearing mice served CNT. Blood samples were collected from the tail vein at baseline (pre-irradiation) and at multiple time points following irradiation to assess systemic effects. (A) Longitudinal measurements of fasting blood glucose determined using a standard glucometer (mg/dL) and plasma insulin quantified using the Milliplex Metabolic Hormone Panel assay (ng/mL, time fixed effect = 0.0049). (B) Circulating levels of ghrelin (pg/mL; time fixed effect = 0.0071), TNF-α (pg/mL; RT fixed effect = 0.0419)), and interleukin-6 (pg/mL, fixed effect: time = 0.01, RT = 0.04) measured over time using the same Milliplex assay. All data are expressed as mean ± SEM (N=3-4 per group). Data are presented as mean±SEM. Statistical analysis was performed using a matched mixed-effect model, followed by appropriate post-hoc tests. Symbols between groups: *= CNT vs CONV; *p<0.05.

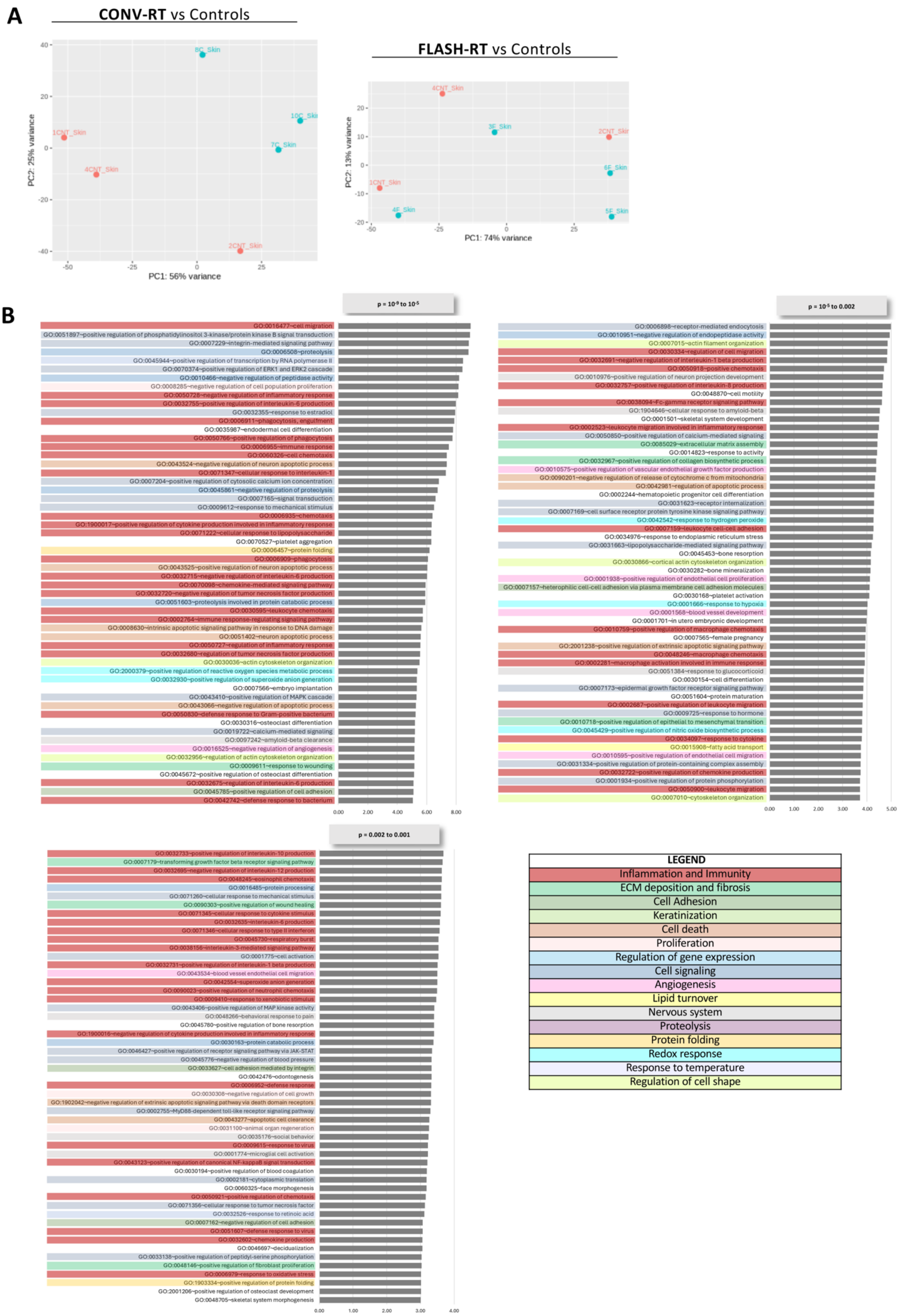
A
CONV-RT vs Controls
FLASH-RT vs Controls
PC1: 56% variance
PC2: 25% variance
PC1: 74% variance
PC2: 13% variance
B
LEGEND
Inflammation and Immunity
ECM deposition and fibrosis
Cell Adhesion
Keratinization
Cell death
Proliferation
Regulation of gene expression
Cell signaling
Angiogenesis
Lipid turnover
Nervous system
Proteolysis
Protein folding
Redox response
Response to temperature
Regulation of cell shape

**Fig. S3. Transcriptomic analysis of left hindlimb skin from mice subjected to conventional and FLASH RT, supplementary analyses.** RNA sequencing (RNA-seq) was performed on skin samples collected from the irradiated region of the left hindlimb 70 days after mice received 35 Gy electron irradiation with either conventional radiotherapy (CONV-RT, n=3) or ultra-high dose rate FLASH radiotherapy (FLASH-RT, n=4), as well as from matched non-irradiated controls (CNT, n=3), as detailed in the Material and Methods section**.** Differential gene expression analysis among irradiated and control groups was conducted using DESeq2. Genes with an absolute fold change > 1.5 and a false discovery rate (FDR)-adjusted p-value < 0.05 were considered significantly differentially expressed. The main data are shown in Figure 3. (A) Principal Component Analysis (PCA) of gene expression. PCA was performed to assess global transcriptional differences in skin samples from mice treated with either CONV or FLASH radiotherapy, compared to non-irradiated controls (CNT). The analysis reveals distinct clustering of samples, particularly between CONV-RT and CNT groups, indicating pronounced gene expression changes induced by CONV-RT. (B) Functional enrichment analysis of Gene Ontology (GO) biological process (BP) terms among upregulated genes in the CONV-RT group. Only the terms with $10^{-9} < p < 0.001$ are displayed in this picture; the top enriched terms (p < 10-9) are shown in Figure 3C. The various annotation terms collected in the figure are color-coded according to their broad categories, summarized in the legend.

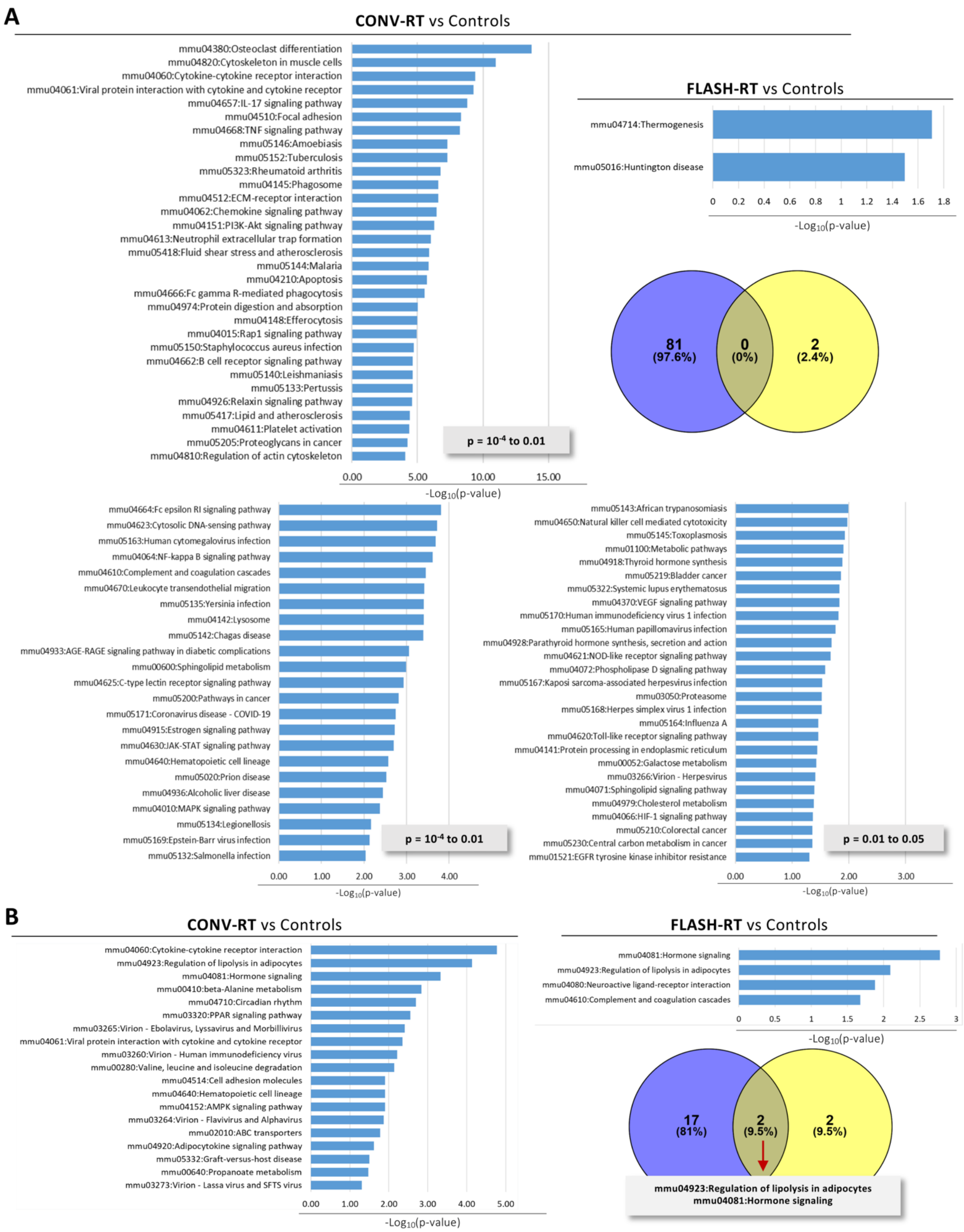


**Fig. S4. KEGG pathways enrichment analysis on the irradiation-induced differentially expressed genes the left hindlimb skin.** RNA sequencing (RNA-seq) was performed on skin samples collected from the irradiated region of the left hindlimb 70 days after mice received 35 Gy electron irradiation with either conventional radiotherapy (CONV-RT, n=3) or ultra-high dose rate FLASH radiotherapy (FLASH-RT, n=4), as well as from matched non-irradiated controls (CNT, n=3), as detailed in the Materials and Methods section . Differential gene expression analysis among irradiated and control groups was conducted using DESeq2. Genes with an absolute fold change > 1.5 and a false discovery rate (FDR)-adjusted p-value < 0.05 were considered significantly differentially expressed. The main data are shown in Figure 3. Functional enrichment analysis of KEGG pathways terms among (A) upregulated and (B) downregulated genes was performed for both irradiated groups as described in the methods; a threshold of $p<0.05$ was considered significant. Bars represent enriched KEGG terms ranked by decreasing significance (-log10 p-value). Venn diagrams summarize the number of enriched KEGG pathways identified in each analysis and their overlap.

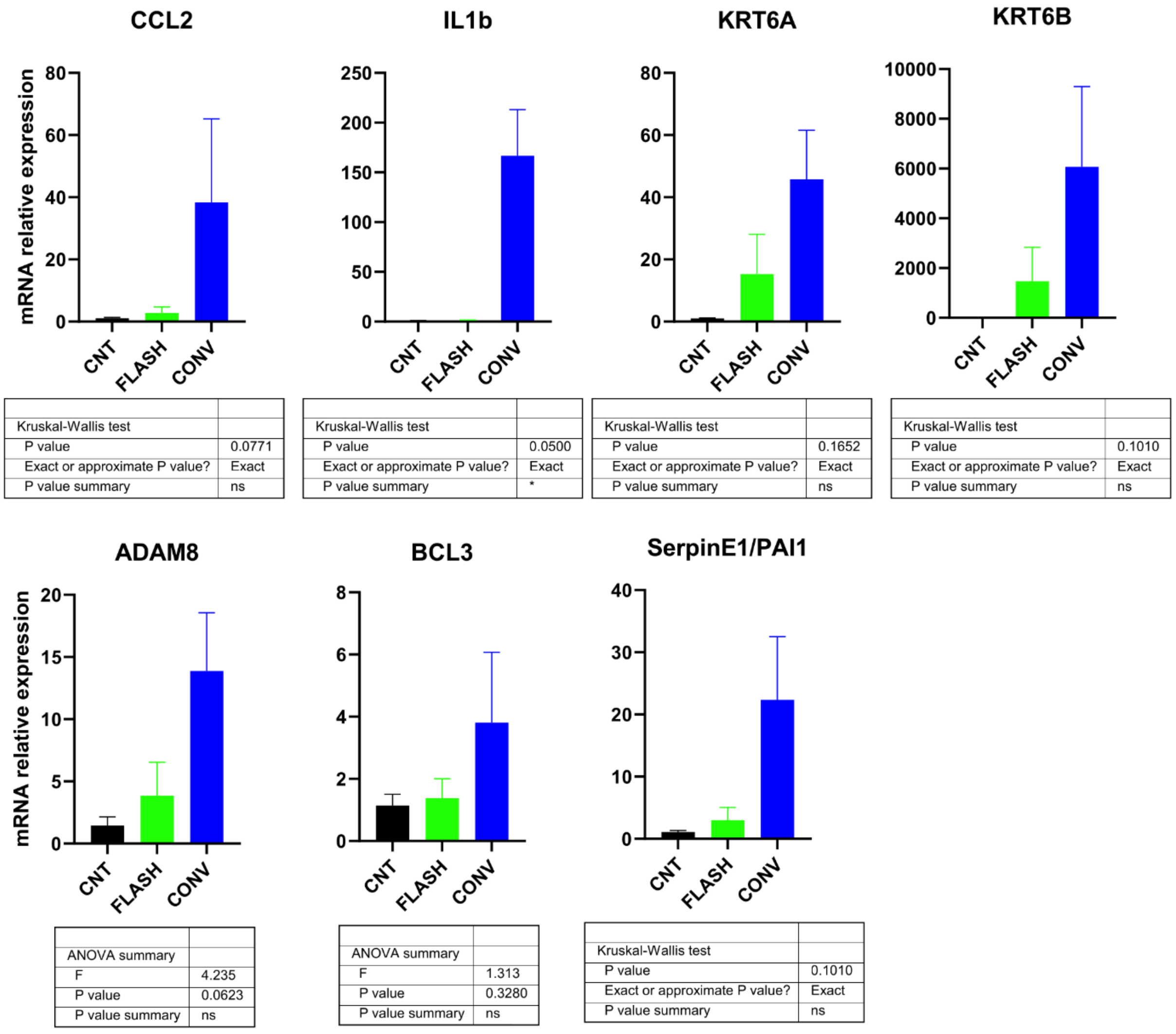


**Fig. S5. Quantitative PCR validation of gene expression changes as identified by RNA-seq in the skin of mice exposed to conventional or FLASH radiotherapy.** RNA was extracted from skin samples collected from the irradiated region of the left hindlimb 70 days after mice received 35 Gy electron irradiation via either conventional radiotherapy (CONV, n = 3) or FLASH radiotherapy (n = 4). Matched non-irradiated mice served as controls (CNT, n = 3). Following reverse transcription, qPCR was performed for the indicated targets using either probe-based or SYBR Green-based systems, with the appropriate master mixes. Relative mRNA expression levels were quantified using the ΔΔCt method, normalized to the housekeeping gene TBP, and expressed relative to non-irradiated controls (CNT). Data are presented as mean ± SEM and the tables below each graph show One-way ANOVA or Kruskal Wallis results, performed as appropriated depending on data distribution.

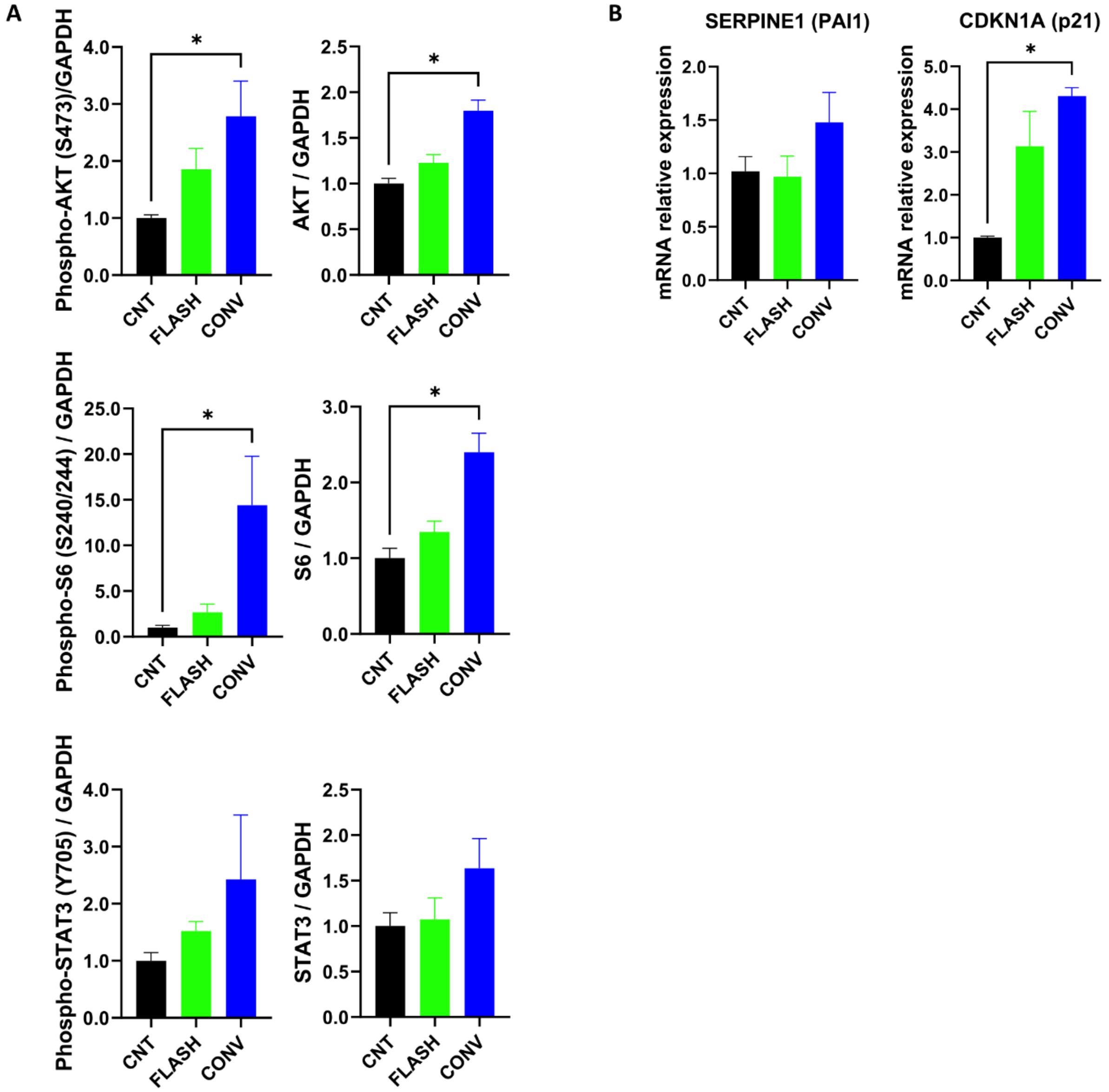


**Fig. S6. Analysis of signaling pathways regulating muscle homeostasis in response to CONV-RT and FLASH-RT.** To investigate the molecular mechanisms underlying the differential structural impact of CONV-RT and FLASH-RT on skeletal muscle, key components of anabolic, catabolic, and stress response pathways were analyzed. Total proteins and RNA were extracted from the *vastus lateralis* muscle of the irradiated left hindlimb, 70 days after exposure to 35 Gy electron irradiation delivered via CONV-RT (n = 3) or FLASH RT (n = 4). Age-matched non-irradiated mice served as controls (n = 3). (A) Western blot analysis was performed to assess phosphorylated (indicated as pAKT, pS6, pSTAT3) and total levels of AKT, S6 ribosomal protein and STAT3 (indicated as AKT, S6, STAT3 on the Y axis) using total protein lysates (30 µg per sample). Representative blots are shown in Figure 7D. Quantification is presented as bar graphs, with protein levels normalized to GAPDH and expressed relative to the control group (CNT). (B) RT-qPCR analysis of PAI-1 and p21 was performed using a SYBR Green-based approach, with the appropriate master mix. Relative mRNA expression levels were quantified using the ΔΔCt method, normalized to the housekeeping gene TBP, and expressed relative to non-irradiated controls (CNT). Data are presented as mean ± SEM. Statistical analysis was performed by Kruskal Wallis analysis, followed by appropriate post-hoc test. *$p<0.05$.

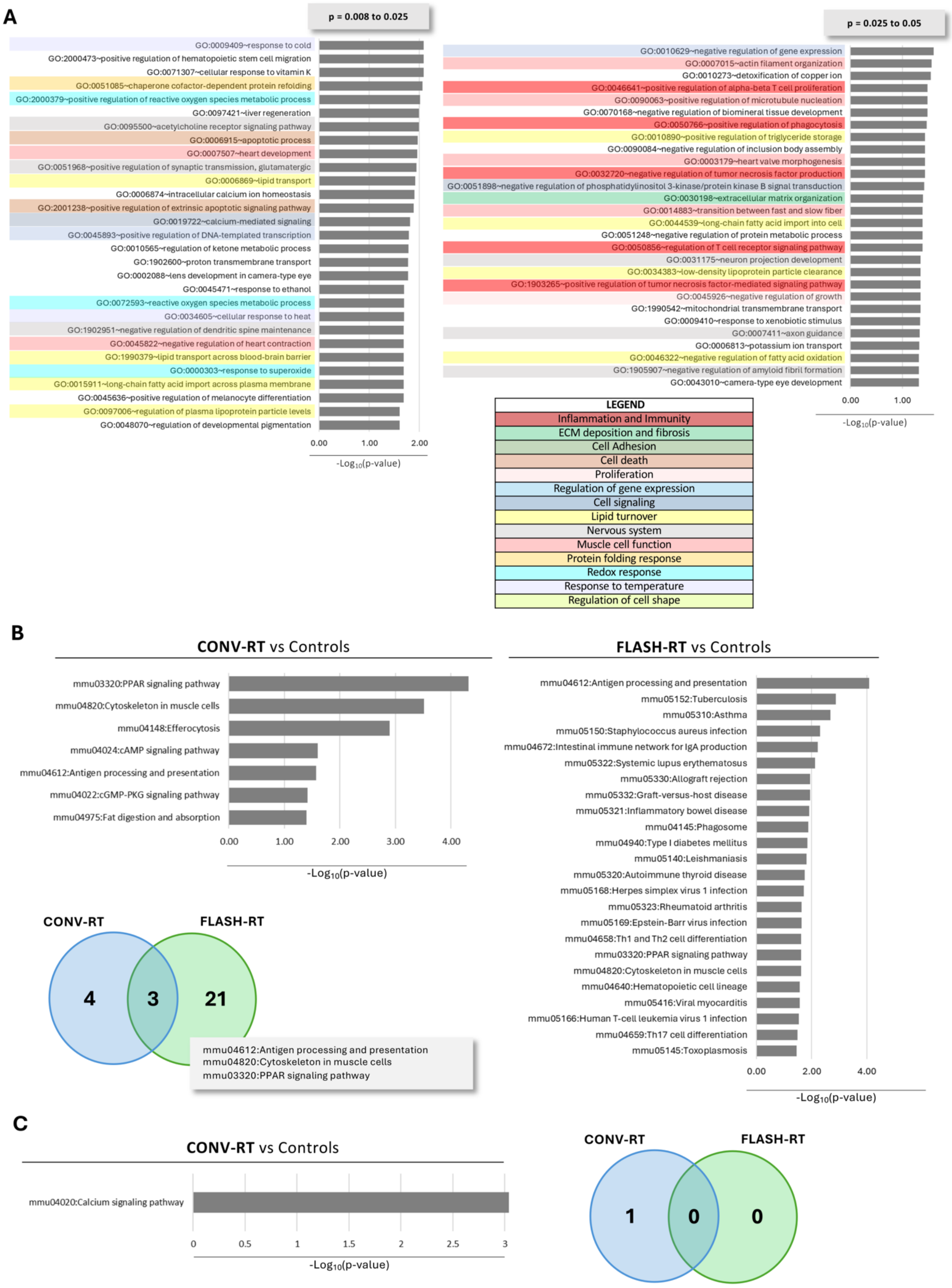


**Fig. S7. Transcriptomic analysis of left hindlimb muscle from mice subjected to conventional and FLASH RT, supplementary analyses.** RNA sequencing (RNA-seq) was performed on muscle (*vastus lateralis*) samples collected from the irradiated region of the left hindlimb 70 days after mice received 35 Gy electron irradiation with either CONV-RT, (n=3) or FLASH-RT (n=4), as well as from matched CNT (n=3), as detailed in the Methods section. Differential gene expression analysis among irradiated and control groups was conducted using DESeq2. Genes with an absolute fold change > 1.5 and a false discovery rate (FDR)-adjusted p-value < 0.05 were considered significantly differentially expressed. The main data are shown in Figure 7. (A) Functional enrichment analysis of Gene Ontology (GO) biological process (BP) terms among upregulated genes in the CONV-RT group. Only the terms with 0.008 < p < 0.05 are displayed in this picture; the top enriched terms (p <0.008) are shown in figure 7C. The various annotation terms collected in the figure are color-coded according to their broad categories, summarized in the legend. (B-C) Functional enrichment analysis of KEGG pathways terms among upregulated (B) downregulated (C) genes in both CONV-RT and FLASH-RT groups (p<0.05). Bars represent enriched KEGG pathways ranked by decreasing significance (-log10 p-value). Venn diagrams summarize the number of enriched terms identified in each analysis and their overlap.

#### Supplementary Tables

##### Table S1. Upregulated genes in CONV-RT mice skin compared to CNT

| Gene ID | Name | log2(FC) | padj |
|---|---|---|---|
| Krt6b | keratin 6B [Source:MGI Symbol;Acc:MGI:1333768] | 13.89 | 1.69E-69 |
| Cstdc5 | cystatin domain containing 5 [Source:MGI Symbol;Acc:MGI:3696883] | 9.84 | 8.71E-55 |
| Stfa3 | stefin A3 [Source:MGI Symbol;Acc:MGI:106196] | 7.74 | 1.88E-36 |
| Sh2d5 | SH2 domain containing 5 [Source:MGI Symbol;Acc:MGI:2446215] | 4.70 | 9.12E-35 |
| Krt16 | keratin 16 [Source:MGI Symbol;Acc:MGI:96690] | 8.65 | 3.00E-34 |
| S100a8 | S100 calcium binding protein A8 (calgranulin A) [Source:MGI Symbol;Acc:MGI:88244] | 8.83 | 1.61E-33 |
| Slpi | secretory leukocyte peptidase inhibitor [Source:MGI Symbol;Acc:MGI:109297] | 9.35 | 2.40E-33 |
| S100a9 | S100 calcium binding protein A9 (calgranulin B) [Source:MGI Symbol;Acc:MGI:1338947] | 10.11 | 1.59E-32 |
| Krt6a | keratin 6A [Source:MGI Symbol;Acc:MGI:1100845] | 5.56 | 1.51E-26 |
| Stfa1 | stefin A1 [Source:MGI Symbol;Acc:MGI:106198] | 9.07 | 2.54E-26 |
| Cdhr1 | cadherin-related family member 1 [Source:MGI Symbol;Acc:MGI:2157782] | 7.50 | 2.56E-24 |
| Saa3 | serum amyloid A 3 [Source:MGI Symbol;Acc:MGI:98223] | 8.83 | 4.20E-24 |
| Clcf1 | cardiotrophin-like cytokine factor 1 [Source:MGI Symbol;Acc:MGI:1930088] | 3.30 | 4.57E-23 |
| Gpr15lg | G protein coupled receptor 15 ligand [Source:MGI Symbol;Acc:MGI:1917295] | 6.64 | 1.29E-22 |
| Trem1 | triggering receptor expressed on myeloid cells 1 [Source:MGI Symbol;Acc:MGI:1930005] | 5.95 | 5.48E-21 |
| Klk6 | kallikrein related-peptidase 6 [Source:MGI Symbol;Acc:MGI:1343166] | 5.67 | 1.09E-20 |
| Il1b | interleukin 1 beta [Source:MGI Symbol;Acc:MGI:96543] | 5.71 | 2.02E-19 |
| Clec4e | C-type lectin domain family 4, member e [Source:MGI Symbol;Acc:MGI:1861232] | 5.46 | 2.02E-19 |
| Spp1 | secreted phosphoprotein 1 [Source:MGI Symbol;Acc:MGI:98389] | 6.60 | 1.35E-16 |
| Pla2g4e | phospholipase A2, group IVE [Source:MGI Symbol;Acc:MGI:1919144] | 3.18 | 2.12E-16 |
| Cxcl2 | C-X-C motif chemokine ligand 2 [Source:MGI Symbol;Acc:MGI:1340094] | 11.53 | 3.90E-16 |
| Hdc | histidine decarboxylase [Source:MGI Symbol;Acc:MGI:96062] | 5.05 | 9.03E-16 |
| Timp1 | tissue inhibitor of metalloproteinase 1 [Source:MGI Symbol;Acc:MGI:98752] | 7.43 | 9.31E-16 |
| Sprr2d | small proline-rich protein 2D [Source:MGI Symbol;Acc:MGI:1330347] | 12.37 | 1.49E-15 |
| Acod1 | aconitate decarboxylase 1 [Source:MGI Symbol;Acc:MGI:103206] | 12.08 | 2.21E-15 |
| Cthrc1 | collagen triple helix repeat containing 1 [Source:MGI Symbol;Acc:MGI:1915838] | 3.33 | 2.24E-15 |
| Osm | oncostatin M [Source:MGI Symbol;Acc:MGI:104749] | 8.00 | 1.64E-14 |
| Clec4d | C-type lectin domain family 4, member d [Source:MGI Symbol;Acc:MGI:1298389] | 7.16 | 1.31E-13 |
| Fosl1 | fos-like antigen 1 [Source:MGI Symbol;Acc:MGI:107179] | 4.13 | 1.52E-13 |
| Slc10a6 | solute carrier family 10 (sodium/bile acid cotransporter family), member 6 [Source:MGI Symbol;Acc:MGI:1923000] | 1.97 | 3.92E-13 |
| Serpinb3a | serine (or cysteine) peptidase inhibitor, clade B (ovalbumin), member 3A [Source:MGI Symbol;Acc:MGI:3573933] | 7.88 | 7.29E-13 |
| Plaur | plasminogen activator, urokinase receptor [Source:MGI Symbol;Acc:MGI:97612] | 4.02 | 8.98E-13 |
| C1qtnf5 | C1q and tumor necrosis factor related protein 5 [Source:MGI Symbol;Acc:MGI:2385958] | 4.50 | 1.14E-12 |
| Aqp3 | aquaporin 3 [Source:MGI Symbol;Acc:MGI:1333777] | 3.15 | 1.14E-12 |
| Fam83a | family with sequence similarity 83, member A [Source:MGI Symbol;Acc:MGI:2447773] | 2.93 | 1.14E-12 |
| Klk13 | kallikrein related-peptidase 13 [Source:MGI Symbol;Acc:MGI:3615275] | 4.42 | 1.75E-12 |
| Slfn4 | schlafen 4 [Source:MGI Symbol;Acc:MGI:1329010] | 4.43 | 7.31E-12 |
| Il24 | interleukin 24 [Source:MGI Symbol;Acc:MGI:2135548] | 10.83 | 7.83E-12 |
| Rptn | repetin [Source:MGI Symbol;Acc:MGI:1099055] | 6.08 | 8.11E-12 |
| Ccr1 | C-C motif chemokine receptor 1 [Source:MGI Symbol;Acc:MGI:104618] | 4.45 | 1.09E-11 |
| Lce3c | late cornified envelope 3C [Source:MGI Symbol;Acc:MGI:2135932] | 6.55 | 1.24E-11 |

| | | | |
|---|---|---|---|
| Glrx | glutaredoxin [Source:MGI Symbol;Acc:MGI:2135625] | 3.03 | 1.37E-11 |
| Csta3 | cystatin A family member 3 [Source:MGI Symbol;Acc:MGI:3644688] | 9.75 | 1.39E-11 |
| Mmp10 | matrix metallopeptidase 10 [Source:MGI Symbol;Acc:MGI:97007] | 10.85 | 1.42E-11 |
| Cxcr4 | C-X-C motif chemokine receptor 4 [Source:MGI Symbol;Acc:MGI:109563] | 2.93 | 1.50E-11 |
| Lcn2 | lipocalin 2 [Source:MGI Symbol;Acc:MGI:96757] | 6.89 | 1.53E-11 |
| Or5m3b | olfactory receptor family 5 subfamily M member 3B [Source:MGI Symbol;Acc:MGI:3030867] | 3.60 | 1.53E-11 |
| Cbln1 | cerebellin 1 precursor protein [Source:MGI Symbol;Acc:MGI:88281] | 10.97 | 1.99E-11 |
| Sprr2i | small proline-rich protein 2I [Source:MGI Symbol;Acc:MGI:1330309] | 10.08 | 1.99E-11 |
| Cxcl3 | C-X-C motif chemokine ligand 3 [Source:MGI Symbol;Acc:MGI:3037818] | 6.56 | 2.44E-11 |
| Serpinb3b | serine (or cysteine) peptidase inhibitor, clade B (ovalbumin), member 3B [Source:MGI Symbol;Acc:MGI:2683293] | 1.72 | 2.44E-11 |
| Csf3r | colony stimulating factor 3 receptor [Source:MGI Symbol;Acc:MGI:1339755] | 5.30 | 2.46E-11 |
| Retnlg | resistin like gamma [Source:MGI Symbol;Acc:MGI:2667763] | 10.33 | 2.53E-11 |
| Crabp1 | cellular retinoic acid binding protein I [Source:MGI Symbol;Acc:MGI:88490] | 2.38 | 4.47E-11 |
| Pla2g4d | phospholipase A2, group IVD [Source:MGI Symbol;Acc:MGI:1925640] | 8.00 | 4.77E-11 |
| Spink12 | serine peptidase inhibitor, Kazal type 12 [Source:MGI Symbol;Acc:MGI:1925492] | 5.64 | 5.40E-11 |
| Gsdmc | gasdermin C [Source:MGI Symbol;Acc:MGI:1933176] | 5.02 | 6.74E-11 |
| Rab31 | RAB31, member RAS oncogene family [Source:MGI Symbol;Acc:MGI:1914603] | 2.34 | 8.69E-11 |
| Gsta4 | glutathione S-transferase, alpha 4 [Source:MGI Symbol;Acc:MGI:1309515] | 4.65 | 1.29E-10 |
| Ifitm1 | interferon induced transmembrane protein 1 [Source:MGI Symbol;Acc:MGI:1915963] | 4.58 | 2.12E-10 |
| Gm15280 | predicted gene 15280 [Source:MGI Symbol;Acc:MGI:3826554] | 3.64 | 2.19E-10 |
| Zxda | zinc finger, X-linked, duplicated A [Source:MGI Symbol;Acc:MGI:1921689] | 9.51 | 2.72E-10 |
| Csf3 | colony stimulating factor 3 (granulocyte) [Source:MGI Symbol;Acc:MGI:1339751] | 5.42 | 3.01E-10 |
| Cd101 | CD101 antigen [Source:MGI Symbol;Acc:MGI:2685862] | 2.95 | 3.52E-10 |
| Cd14 | CD14 antigen [Source:MGI Symbol;Acc:MGI:88318] | 4.82 | 4.55E-10 |
| Sprr2g | small proline-rich protein 2G [Source:MGI Symbol;Acc:MGI:1330348] | 9.95 | 4.98E-10 |
| Degs2 | delta 4-desaturase, sphingolipid 2 [Source:MGI Symbol;Acc:MGI:1917309] | 4.52 | 5.80E-10 |
| Srgn | serglycin [Source:MGI Symbol;Acc:MGI:97756] | 3.41 | 6.40E-10 |
| Klk10 | kallikrein related-peptidase 10 [Source:MGI Symbol;Acc:MGI:1916790] | 1.83 | 9.10E-10 |
| Ccdc88b | coiled-coil domain containing 88B [Source:MGI Symbol;Acc:MGI:1925567] | 3.03 | 1.52E-09 |
| Ptgs2 | prostaglandin-endoperoxide synthase 2 [Source:MGI Symbol;Acc:MGI:97798] | 4.06 | 1.55E-09 |
| Cyp2g1 | cytochrome P450, family 2, subfamily g, polypeptide 1 [Source:MGI Symbol;Acc:MGI:109612] | 4.87 | 1.64E-09 |
| Nlrp3 | NLR family, pyrin domain containing 3 [Source:MGI Symbol;Acc:MGI:2653833] | 4.76 | 1.70E-09 |
| Sprr2e | small proline-rich protein 2E [Source:MGI Symbol;Acc:MGI:1330346] | 9.69 | 1.72E-09 |
| Mmp8 | matrix metallopeptidase 8 [Source:MGI Symbol;Acc:MGI:1202395] | 4.76 | 1.82E-09 |
| Srxn1 | sulfiredoxin 1 homolog (S. cerevisiae) [Source:MGI Symbol;Acc:MGI:104971] | 1.73 | 1.85E-09 |
| Chi3l1 | chitinase 3 like 1 [Source:MGI Symbol;Acc:MGI:1340899] | 4.69 | 1.88E-09 |
| Klk14 | kallikrein related-peptidase 14 [Source:MGI Symbol;Acc:MGI:2447564] | 4.09 | 1.88E-09 |
| Sprr2a2 | small proline-rich protein 2A2 [Source:MGI Symbol;Acc:MGI:3845026] | 5.71 | 2.14E-09 |
| Prss27 | serine protease 27 [Source:MGI Symbol;Acc:MGI:2450123] | 3.41 | 4.51E-09 |
| Sprr2k | small proline-rich protein 2K [Source:MGI Symbol;Acc:MGI:1330344] | 8.46 | 4.73E-09 |
| Mapk6 | mitogen-activated protein kinase 6 [Source:MGI Symbol;Acc:MGI:1354946] | 1.68 | 4.81E-09 |
| Stfa2l1 | stefin A2 like 1 [Source:MGI Symbol;Acc:MGI:3524944] | 9.42 | 5.52E-09 |
| Adam8 | a disintegrin and metallopeptidase domain 8 [Source:MGI Symbol;Acc:MGI:107825] | 3.12 | 5.52E-09 |
| Cstdc6 | cystatin domain containing 6 [Source:MGI Symbol;Acc:MGI:3696881] | 8.59 | 8.99E-09 |
| Mrgpra2b | MAS-related GPR, member A2B [Source:MGI Symbol;Acc:MGI:3033098] | 9.21 | 1.23E-08 |
| Stac2 | SH3 and cysteine rich domain 2 [Source:MGI Symbol;Acc:MGI:2144518] | 2.37 | 1.31E-08 |
| Fpr1 | formyl peptide receptor 1 [Source:MGI Symbol;Acc:MGI:107443] | 6.95 | 1.46E-08 |
| Akr1b8 | aldo-keto reductase family 1, member B8 [Source:MGI Symbol;Acc:MGI:107673] | 3.17 | 1.46E-08 |
| Cxcl1 | C-X-C motif chemokine ligand 1 [Source:MGI Symbol;Acc:MGI:108068] | 6.35 | 1.53E-08 |
| Gm49339 | predicted gene, 49339 [Source:MGI Symbol;Acc:MGI:6121530] | 4.30 | 1.68E-08 |
| Il1rl1 | interleukin 1 receptor-like 1 [Source:MGI Symbol;Acc:MGI:98427] | 4.43 | 1.69E-08 |
| Cd300lf | CD300 molecule like family member F [Source:MGI Symbol;Acc:MGI:2442359] | 3.60 | 1.89E-08 |
| Artn | artemin [Source:MGI Symbol;Acc:MGI:1333791] | 3.70 | 1.90E-08 |
| Brinp3 | bone morphogenetic protein/retinoic acid inducible neural specific 3 [Source:MGI Symbol;Acc:MGI:2443035] | 5.65 | 2.11E-08 |
| Tinagl1 | tubulointerstitial nephritis antigen-like 1 [Source:MGI Symbol;Acc:MGI:2137617] | 2.17 | 2.20E-08 |
| Adamts4 | ADAM metallopeptidase with thrombospondin type 1 motif 4 [Source:MGI Symbol;Acc:MGI:1339949] | 4.02 | 2.36E-08 |
| Phlda1 | pleckstrin homology like domain, family A, member 1 [Source:MGI Symbol;Acc:MGI:1096880] | 1.87 | 2.46E-08 |
| Qsox1 | quiescin Q6 sulfhydryl oxidase 1 [Source:MGI Symbol;Acc:MGI:1330818] | 1.37 | 2.78E-08 |
| Sprr2j-ps | small proline-rich protein 2J, pseudogene [Source:MGI Symbol;Acc:MGI:1330345] | 8.51 | 2.89E-08 |
| Inhba | inhibin beta-A [Source:MGI Symbol;Acc:MGI:96570] | 3.94 | 3.36E-08 |
| Mmp11 | matrix metallopeptidase 11 [Source:MGI Symbol;Acc:MGI:97008] | 2.87 | 3.75E-08 |
| Fscn1 | fascin actin-bundling protein 1 [Source:MGI Symbol;Acc:MGI:1352745] | 3.21 | 5.03E-08 |
| Col12a1 | collagen, type XII, alpha 1 [Source:MGI Symbol;Acc:MGI:88448] | 4.77 | 5.26E-08 |
| C5ar1 | complement component 5a receptor 1 [Source:MGI Symbol;Acc:MGI:88232] | 3.19 | 5.26E-08 |
| Mmp1b | matrix metallopeptidase 1b (interstitial collagenase) [Source:MGI Symbol;Acc:MGI:1933847] | 8.65 | 5.38E-08 |
| Dnmt3l | DNA methyltransferase 3-like [Source:MGI Symbol;Acc:MGI:1859287] | 3.07 | 6.92E-08 |
| Sgpp2 | sphingosine-1-phosphate phosphatase 2 [Source:MGI Symbol;Acc:MGI:3589109] | 1.56 | 7.00E-08 |
| Elf3 | E74-like factor 3 [Source:MGI Symbol;Acc:MGI:1101781] | 4.15 | 8.03E-08 |
| Il4ra | interleukin 4 receptor, alpha [Source:MGI Symbol;Acc:MGI:105367] | 2.78 | 8.15E-08 |
| Rbp2 | retinol binding protein 2, cellular [Source:MGI Symbol;Acc:MGI:97877] | 3.35 | 9.74E-08 |
| Cemip | cell migration inducing protein, hyaluronan binding [Source:MGI Symbol;Acc:MGI:2443629] | 4.16 | 1.07E-07 |

| | | | |
|---|---|---|---|
| Homer3 | homer scaffolding protein 3 [Source:MGI Symbol;Acc:MGI:1347359] | 1.55 | 1.49E-07 |
| Ccl3 | C-C motif chemokine ligand 3 [Source:MGI Symbol;Acc:MGI:98260] | 6.81 | 1.64E-07 |
| Ms4a6b | membrane-spanning 4-domains, subfamily A, member 6B [Source:MGI Symbol;Acc:MGI:1917024] | 3.22 | 1.94E-07 |
| Cyp4f18 | cytochrome P450, family 4, subfamily f, polypeptide 18 [Source:MGI Symbol;Acc:MGI:1919304] | 3.53 | 2.39E-07 |
| Gm18679 | predicted gene, 18679 [Source:MGI Symbol;Acc:MGI:5010864] | 8.81 | 2.45E-07 |
| Smim3 | small integral membrane protein 3 [Source:MGI Symbol;Acc:MGI:1917088] | 2.89 | 2.61E-07 |
| Cc2d2b | coiled-coil and C2 domain containing 2B [Source:MGI Symbol;Acc:MGI:3645359] | 5.30 | 2.64E-07 |
| Tnfsf11 | tumor necrosis factor (ligand) superfamily, member 11 [Source:MGI Symbol;Acc:MGI:1100089] | 5.19 | 2.77E-07 |
| Bcl3 | B cell leukemia/lymphoma 3 [Source:MGI Symbol;Acc:MGI:88140] | 2.75 | 2.77E-07 |
| Mcemp1 | mast cell expressed membrane protein 1 [Source:MGI Symbol;Acc:MGI:1916439] | 3.65 | 3.12E-07 |
| Ccdc71l | coiled-coil domain containing 71 like [Source:MGI Symbol;Acc:MGI:1919373] | 1.36 | 3.14E-07 |
| Kcnn4 | potassium intermediate/small conductance calcium-activated channel, subfamily N, member 4 [Source:MGI Symbol;Acc:MGI:1277957] | 2.46 | 3.50E-07 |
| Sptbn4 | spectrin beta, non-erythrocytic 4 [Source:MGI Symbol;Acc:MGI:1890574] | 3.68 | 3.78E-07 |
| Car2 | carbonic anhydrase 2 [Source:MGI Symbol;Acc:MGI:88269] | 3.45 | 3.83E-07 |
| Tlr13 | toll-like receptor 13 [Source:MGI Symbol;Acc:MGI:3045213] | 2.94 | 4.05E-07 |
| Slc13a3 | solute carrier family 13 (sodium-dependent dicarboxylate transporter), member 3 [Source:MGI Symbol;Acc:MGI:2149635] | 4.90 | 4.29E-07 |
| Areg | amphiregulin [Source:MGI Symbol;Acc:MGI:88068] | 3.74 | 4.29E-07 |
| Gm20784 | predicted gene, 20784 [Source:MGI Symbol;Acc:MGI:5434140] | 8.28 | 4.53E-07 |
| Hbegf | heparin-binding EGF-like growth factor [Source:MGI Symbol;Acc:MGI:96070] | 3.06 | 4.53E-07 |
| Fetub | fetuin beta [Source:MGI Symbol;Acc:MGI:1890221] | 4.66 | 4.54E-07 |
| Hck | hemopoietic cell kinase [Source:MGI Symbol;Acc:MGI:96052] | 3.40 | 4.54E-07 |
| Urah | urate (5-hydroxyiso-) hydrolase [Source:MGI Symbol;Acc:MGI:1916142] | 2.38 | 5.87E-07 |
| Slfn1 | schlafen 1 [Source:MGI Symbol;Acc:MGI:1313259] | 3.34 | 8.03E-07 |
| Ms4a7 | membrane-spanning 4-domains, subfamily A, member 7 [Source:MGI Symbol;Acc:MGI:1918846] | 2.67 | 8.03E-07 |
| Lce3a | late cornified envelope 3A [Source:MGI Symbol;Acc:MGI:3645650] | 9.10 | 8.28E-07 |
| Fbln2 | fibulin 2 [Source:MGI Symbol;Acc:MGI:95488] | 2.83 | 9.06E-07 |
| Furin | furin, paired basic amino acid cleaving enzyme [Source:MGI Symbol;Acc:MGI:97513] | 1.77 | 9.51E-07 |
| Slc15a3 | solute carrier family 15, member 3 [Source:MGI Symbol;Acc:MGI:1929691] | 3.31 | 1.09E-06 |
| Fibin | fin bud initiation factor homolog (zebrafish) [Source:MGI Symbol;Acc:MGI:1914856] | 5.06 | 1.17E-06 |
| Fpr2 | formyl peptide receptor 2 [Source:MGI Symbol;Acc:MGI:1278319] | 6.90 | 1.21E-06 |
| Acp5 | acid phosphatase 5, tartrate resistant [Source:MGI Symbol;Acc:MGI:87883] | 1.07 | 1.22E-06 |
| Sell | selectin, lymphocyte [Source:MGI Symbol;Acc:MGI:98279] | 3.48 | 1.31E-06 |
| Lilrb4b | leukocyte immunoglobulin-like receptor, subfamily B, member 4B [Source:MGI Symbol;Acc:MGI:102702] | 3.29 | 1.54E-06 |
| Trem3 | triggering receptor expressed on myeloid cells 3 [Source:MGI Symbol;Acc:MGI:1930003] | 7.87 | 1.61E-06 |
| Tnc | tenascin C [Source:MGI Symbol;Acc:MGI:101922] | 7.18 | 1.64E-06 |
| Serpine1 | serine (or cysteine) peptidase inhibitor, clade E, member 1 [Source:MGI Symbol;Acc:MGI:97608] | 3.82 | 1.73E-06 |
| Adrm1b | adhesion regulating molecule 1B [Source:MGI Symbol;Acc:MGI:3642386] | 7.06 | 1.77E-06 |
| Hhipl1 | hedgehog interacting protein-like 1 [Source:MGI Symbol;Acc:MGI:1919265] | 3.30 | 1.97E-06 |
| Car12 | carbonic anhydrase 12 [Source:MGI Symbol;Acc:MGI:1923709] | 1.43 | 2.20E-06 |
| Sirpb1b | signal-regulatory protein beta 1B [Source:MGI Symbol;Acc:MGI:3779828] | 2.98 | 2.22E-06 |
| Prl2c2 | prolactin family 2, subfamily c, member 2 [Source:MGI Symbol;Acc:MGI:97618] | 7.30 | 2.44E-06 |
| Stab1 | stabilin 1 [Source:MGI Symbol;Acc:MGI:2178742] | 1.56 | 2.58E-06 |
| Oaf | out at first homolog [Source:MGI Symbol;Acc:MGI:94852] | 2.25 | 2.92E-06 |
| Gm5150 | predicted gene 5150 [Source:MGI Symbol;Acc:MGI:3779469] | 5.44 | 2.95E-06 |
| Thbs4 | thrombospondin 4 [Source:MGI Symbol;Acc:MGI:1101779] | 2.94 | 3.03E-06 |
| Slc26a4 | solute carrier family 26, member 4 [Source:MGI Symbol;Acc:MGI:1346029] | 6.42 | 3.08E-06 |
| Col6a3 | collagen, type VI, alpha 3 [Source:MGI Symbol;Acc:MGI:88461] | 2.94 | 3.51E-06 |
| Ppard | peroxisome proliferator activator receptor delta [Source:MGI Symbol;Acc:MGI:101884] | 1.91 | 3.53E-06 |
| Ier5l | immediate early response 5-like [Source:MGI Symbol;Acc:MGI:1919750] | 2.25 | 3.56E-06 |
| Fabp5 | fatty acid binding protein 5, epidermal [Source:MGI Symbol;Acc:MGI:101790] | 3.32 | 3.75E-06 |
| Adam12 | ADAM metallopeptidase domain 12 [Source:MGI Symbol;Acc:MGI:105378] | 2.82 | 3.75E-06 |
| 2410039M03Rik | RIKEN cDNA 2410039M03 gene [Source:MGI Symbol;Acc:MGI:1917269] | 2.77 | 3.91E-06 |
| Cotl1 | coactosin like F-actin binding protein 1 [Source:MGI Symbol;Acc:MGI:1919292] | 2.12 | 3.98E-06 |
| Lce3f | late cornified envelope 3F [Source:MGI Symbol;Acc:MGI:1916770] | 11.32 | 4.01E-06 |
| Olah | oleoyl-ACP hydrolase [Source:MGI Symbol;Acc:MGI:2139018] | 4.78 | 4.22E-06 |
| Jdp2 | Jun dimerization protein 2 [Source:MGI Symbol;Acc:MGI:1932093] | 2.22 | 4.45E-06 |
| Ier3 | immediate early response 3 [Source:MGI Symbol;Acc:MGI:104814] | 2.34 | 4.52E-06 |
| Ccn5 | cellular communication network factor 5 [Source:MGI Symbol;Acc:MGI:1328326] | 2.05 | 5.34E-06 |
| Fcgr4 | Fc receptor, IgG, low affinity IV [Source:MGI Symbol;Acc:MGI:2179523] | 4.21 | 5.47E-06 |
| Tyrobp | TYRO protein tyrosine kinase binding protein [Source:MGI Symbol;Acc:MGI:1277211] | 2.60 | 5.47E-06 |
| Rbp1 | retinol binding protein 1, cellular [Source:MGI Symbol;Acc:MGI:97876] | 2.14 | 5.47E-06 |
| Bdkrb2 | bradykinin receptor, beta 2 [Source:MGI Symbol;Acc:MGI:102845] | 2.32 | 5.51E-06 |
| Gm15844 | predicted gene 15844 [Source:MGI Symbol;Acc:MGI:3802079] | 2.03 | 5.51E-06 |
| Tnfsf14 | tumor necrosis factor (ligand) superfamily, member 14 [Source:MGI Symbol;Acc:MGI:1355317] | 3.26 | 5.69E-06 |
| Tcf23 | transcription factor 23 [Source:MGI Symbol;Acc:MGI:1934960] | 4.04 | 6.09E-06 |
| Dchs1 | dachsous cadherin related 1 [Source:MGI Symbol;Acc:MGI:2685011] | 2.40 | 6.25E-06 |
| Treml4 | triggering receptor expressed on myeloid cells-like 4 [Source:MGI Symbol;Acc:MGI:1923239] | 7.12 | 7.61E-06 |
| Col18a1 | collagen, type XVIII, alpha 1 [Source:MGI Symbol;Acc:MGI:88451] | 1.79 | 7.61E-06 |
| Cers4 | ceramide synthase 4 [Source:MGI Symbol;Acc:MGI:1914510] | 0.93 | 7.61E-06 |
| Il6 | interleukin 6 [Source:MGI Symbol;Acc:MGI:96559] | 8.03 | 8.00E-06 |

| | | | |
|---|---|---|---|
| Cpxm1 | carboxypeptidase X, M14 family member 1 [Source:MGI Symbol;Acc:MGI:1934569] | 2.85 | 8.16E-06 |
| Slc7a11 | solute carrier family 7 (cationic amino acid transporter, y+ system), member 11 [Source:MGI Symbol;Acc:MGI:1347355] | 1.92 | 8.29E-06 |
| Plek | pleckstrin [Source:MGI Symbol;Acc:MGI:1860485] | 2.64 | 8.30E-06 |
| Col24a1 | collagen, type XXIV, alpha 1 [Source:MGI Symbol;Acc:MGI:1918605] | 4.40 | 8.46E-06 |
| Ccdc85b | coiled-coil domain containing 85B [Source:MGI Symbol;Acc:MGI:2147607] | 1.17 | 8.48E-06 |
| Adam30 | a disintegrin and metallopeptidase domain 30 [Source:MGI Symbol;Acc:MGI:1918328] | 4.45 | 8.51E-06 |
| Tcim | transcriptional and immune response regulator [Source:MGI Symbol;Acc:MGI:1916318] | 1.31 | 8.55E-06 |
| Chil3 | chitinase-like 3 [Source:MGI Symbol;Acc:MGI:1330860] | 6.47 | 8.76E-06 |
| Crispld2 | cysteine-rich secretory protein LCCL domain containing 2 [Source:MGI Symbol;Acc:MGI:1926142] | 3.18 | 1.00E-05 |
| Clec2e | C-type lectin domain family 2, member e [Source:MGI Symbol;Acc:MGI:3028921] | 2.54 | 1.00E-05 |
| Trex2 | three prime repair exonuclease 2 [Source:MGI Symbol;Acc:MGI:1346343] | 2.19 | 1.03E-05 |
| Prss22 | serine protease 22 [Source:MGI Symbol;Acc:MGI:1918085] | 2.72 | 1.05E-05 |
| Defb14 | defensin beta 14 [Source:MGI Symbol;Acc:MGI:2675345] | 4.78 | 1.05E-05 |
| Bdkrb1 | bradykinin receptor, beta 1 [Source:MGI Symbol;Acc:MGI:88144] | 4.08 | 1.06E-05 |
| Nr4a2 | nuclear receptor subfamily 4, group A, member 2 [Source:MGI Symbol;Acc:MGI:1352456] | 2.13 | 1.10E-05 |
| Sirpb1c | signal-regulatory protein beta 1C [Source:MGI Symbol;Acc:MGI:3807521] | 3.33 | 1.15E-05 |
| Trex1 | three prime repair exonuclease 1 [Source:MGI Symbol;Acc:MGI:1328317] | 1.74 | 1.17E-05 |
| Ehf | ets homologous factor [Source:MGI Symbol;Acc:MGI:1270840] | 1.57 | 1.19E-05 |
| Lgmn | legumain [Source:MGI Symbol;Acc:MGI:1330838] | 2.01 | 1.21E-05 |
| Hspa5 | heat shock protein 5 [Source:MGI Symbol;Acc:MGI:95835] | 1.25 | 1.24E-05 |
| Tagln2 | transgelin 2 [Source:MGI Symbol;Acc:MGI:1312985] | 1.56 | 1.25E-05 |
| Gm49037 | predicted gene, 49037 [Source:MGI Symbol;Acc:MGI:6118410] | 1.99 | 1.28E-05 |
| Pctp | phosphatidylcholine transfer protein [Source:MGI Symbol;Acc:MGI:107375] | 1.52 | 1.28E-05 |
| Apln | apelin [Source:MGI Symbol;Acc:MGI:1353624] | 2.56 | 1.35E-05 |
| Selp | selectin, platelet [Source:MGI Symbol;Acc:MGI:98280] | 4.11 | 1.35E-05 |
| Cdh11 | cadherin 11 [Source:MGI Symbol;Acc:MGI:99217] | 2.68 | 1.37E-05 |
| Ddah1 | dimethylarginine dimethylaminohydrolase 1 [Source:MGI Symbol;Acc:MGI:1916469] | 3.61 | 1.42E-05 |
| Piezo1 | piezo-type mechanosensitive ion channel component 1 [Source:MGI Symbol;Acc:MGI:3603204] | 1.16 | 1.57E-05 |
| Hilpda | hypoxia inducible lipid droplet associated [Source:MGI Symbol;Acc:MGI:1916823] | 2.36 | 1.57E-05 |
| Cd177 | CD177 antigen [Source:MGI Symbol;Acc:MGI:1916141] | 3.84 | 1.61E-05 |
| Ptges | prostaglandin E synthase [Source:MGI Symbol;Acc:MGI:1927593] | 2.11 | 1.61E-05 |
| Ms4a4a | membrane-spanning 4-domains, subfamily A, member 4A [Source:MGI Symbol;Acc:MGI:3643932] | 2.77 | 1.71E-05 |
| Trem2 | triggering receptor expressed on myeloid cells 2 [Source:MGI Symbol;Acc:MGI:1913150] | 2.74 | 1.78E-05 |
| Clec4n | C-type lectin domain family 4, member n [Source:MGI Symbol;Acc:MGI:1861231] | 3.05 | 1.78E-05 |
| Cd300c2 | CD300C molecule 2 [Source:MGI Symbol;Acc:MGI:2153249] | 3.15 | 1.84E-05 |
| Vmp1 | vacuole membrane protein 1 [Source:MGI Symbol;Acc:MGI:1923159] | 1.87 | 1.98E-05 |
| Arc | activity regulated cytoskeletal-associated protein [Source:MGI Symbol;Acc:MGI:88067] | 3.43 | 2.09E-05 |
| Sema3g | sema domain, immunoglobulin domain (Ig), short basic domain, secreted, (semaphorin) 3G [Source:MGI Symbol;Acc:MGI:3041242] | 2.04 | 2.11E-05 |
| Duox1 | dual oxidase 1 [Source:MGI Symbol;Acc:MGI:2139422] | 1.28 | 2.11E-05 |
| Arsj | arylsulfatase J [Source:MGI Symbol;Acc:MGI:2443513] | 2.69 | 2.13E-05 |
| Igfbp7 | insulin-like growth factor binding protein 7 [Source:MGI Symbol;Acc:MGI:1352480] | 2.20 | 2.15E-05 |
| C1qc | complement component 1, q subcomponent, C chain [Source:MGI Symbol;Acc:MGI:88225] | 2.01 | 2.25E-05 |
| Ccn4 | cellular communication network factor 4 [Source:MGI Symbol;Acc:MGI:1197008] | 3.65 | 2.45E-05 |
| Lce1g | late cornified envelope 1G [Source:MGI Symbol;Acc:MGI:1913445] | 4.33 | 2.47E-05 |
| Foxd4 | forkhead box D4 [Source:MGI Symbol;Acc:MGI:1347467] | 7.34 | 2.48E-05 |
| Fkbp10 | FK506 binding protein 10 [Source:MGI Symbol;Acc:MGI:104769] | 2.39 | 2.51E-05 |
| Nppb | natriuretic peptide type B [Source:MGI Symbol;Acc:MGI:97368] | 5.80 | 2.53E-05 |
| Art2a | ADP-ribosyltransferase 2a [Source:MGI Symbol;Acc:MGI:107546] | 2.82 | 2.67E-05 |
| Gpr176 | G protein-coupled receptor 176 [Source:MGI Symbol;Acc:MGI:2685858] | 3.16 | 2.78E-05 |
| Dusp5 | dual specificity phosphatase 5 [Source:MGI Symbol;Acc:MGI:2685183] | 1.95 | 2.86E-05 |
| Sprr2b | small proline-rich protein 2B [Source:MGI Symbol;Acc:MGI:1330352] | 8.22 | 3.00E-05 |
| Ttc21a | tetratricopeptide repeat domain 21A [Source:MGI Symbol;Acc:MGI:1921302] | 3.14 | 3.00E-05 |
| Pfkfb4 | 6-phosphofructo-2-kinase/fructose-2,6-biphosphatase 4 [Source:MGI Symbol;Acc:MGI:2687284] | 1.29 | 3.11E-05 |
| Msr1 | macrophage scavenger receptor 1 [Source:MGI Symbol;Acc:MGI:98257] | 2.91 | 3.12E-05 |
| Eno1 | enolase 1, alpha non-neuron [Source:MGI Symbol;Acc:MGI:95393] | 1.67 | 3.19E-05 |
| Inhbb | inhibin beta-B [Source:MGI Symbol;Acc:MGI:96571] | 1.52 | 3.33E-05 |
| Draxin | dorsal inhibitory axon guidance protein [Source:MGI Symbol;Acc:MGI:1917683] | 3.63 | 3.58E-05 |
| Crip1 | cysteine-rich protein 1 [Source:MGI Symbol;Acc:MGI:88501] | 1.77 | 3.73E-05 |
| Apol8 | apolipoprotein L 8 [Source:MGI Symbol;Acc:MGI:2444921] | 2.54 | 3.86E-05 |
| Mmp13 | matrix metallopeptidase 13 [Source:MGI Symbol;Acc:MGI:1340026] | 8.03 | 3.93E-05 |
| Ctsb | cathepsin B [Source:MGI Symbol;Acc:MGI:88561] | 1.30 | 3.95E-05 |
| Ripk3 | receptor-interacting serine-threonine kinase 3 [Source:MGI Symbol;Acc:MGI:2154952] | 1.87 | 3.95E-05 |
| Mmp1a | matrix metallopeptidase 1a (interstitial collagenase) [Source:MGI Symbol;Acc:MGI:1933846] | 6.99 | 3.97E-05 |
| AW551984 | expressed sequence AW551984 [Source:MGI Symbol;Acc:MGI:2143322] | 2.94 | 3.97E-05 |
| Krt14 | keratin 14 [Source:MGI Symbol;Acc:MGI:96688] | 3.25 | 4.04E-05 |
| Prss12 | serine protease 12 neurotrypsin (motopsin) [Source:MGI Symbol;Acc:MGI:1100881] | 2.10 | 4.05E-05 |
| Sprr2a3 | small proline-rich protein 2A3 [Source:MGI Symbol;Acc:MGI:3845028] | 4.38 | 4.14E-05 |
| Lrrc32 | leucine rich repeat containing 32 [Source:MGI Symbol;Acc:MGI:93882] | 2.53 | 4.17E-05 |
| Ppa1 | pyrophosphatase (inorganic) 1 [Source:MGI Symbol;Acc:MGI:97831] | 0.96 | 4.24E-05 |
| Wfdc12 | WAP four-disulfide core domain 12 [Source:MGI Symbol;Acc:MGI:2183434] | 2.93 | 4.30E-05 |
| Mmp19 | matrix metallopeptidase 19 [Source:MGI Symbol;Acc:MGI:1927899] | 2.94 | 4.34E-05 |

| | | | |
|---|---|---|---|
| Twist2 | twist basic helix-loop-helix transcription factor 2 [Source:MGI Symbol;Acc:MGI:104685] | 2.78 | 4.46E-05 |
| Gm7329 | predicted gene 7329 [Source:MGI Symbol;Acc:MGI:3646271] | 7.25 | 5.04E-05 |
| Slc11a1 | solute carrier family 11 (proton-coupled divalent metal ion transporters), member 1 [Source:MGI Symbol;Acc:MGI:1345275] | 2.79 | 5.04E-05 |
| Epgn | epithelial mitogen [Source:MGI Symbol;Acc:MGI:1919170] | 2.93 | 5.20E-05 |
| Snai1 | snail family zinc finger 1 [Source:MGI Symbol;Acc:MGI:98330] | 3.33 | 5.31E-05 |
| Fam43a | family with sequence similarity 43, member A [Source:MGI Symbol;Acc:MGI:2676309] | 1.81 | 5.59E-05 |
| Serpinb6c | serine (or cysteine) peptidase inhibitor, clade B, member 6c [Source:MGI Symbol;Acc:MGI:2145481] | 2.16 | 5.69E-05 |
| Gla | galactosidase, alpha [Source:MGI Symbol;Acc:MGI:1347344] | 1.51 | 5.69E-05 |
| Atf4 | activating transcription factor 4 [Source:MGI Symbol;Acc:MGI:88096] | 1.12 | 5.71E-05 |
| Gm14681 | predicted gene 14681 [Source:MGI Symbol;Acc:MGI:3705734] | 8.42 | 5.94E-05 |
| Lce3e | late cornified envelope 3E [Source:MGI Symbol;Acc:MGI:1916764] | 11.73 | 6.03E-05 |
| Tuba1c | tubulin, alpha 1C [Source:MGI Symbol;Acc:MGI:1095409] | 1.68 | 6.08E-05 |
| Thbs1 | thrombospondin 1 [Source:MGI Symbol;Acc:MGI:98737] | 2.59 | 6.38E-05 |
| Slc16a1 | solute carrier family 16 (monocarboxylic acid transporters), member 1 [Source:MGI Symbol;Acc:MGI:106013] | 1.73 | 6.43E-05 |
| Mxd1 | MAX dimerization protein 1 [Source:MGI Symbol;Acc:MGI:96908] | 1.92 | 6.48E-05 |
| Gm20547 | predicted gene 20547 [Source:MGI Symbol;Acc:MGI:5142012] | 7.00 | 7.02E-05 |
| Dynap | dynactin associated protein [Source:MGI Symbol;Acc:MGI:1922827] | 6.87 | 7.06E-05 |
| Ninj1 | ninjurin 1 [Source:MGI Symbol;Acc:MGI:1196617] | 1.65 | 7.37E-05 |
| Or2y12 | olfactory receptor family 2 subfamily Y member 12 [Source:MGI Symbol;Acc:MGI:3031216] | 2.43 | 7.47E-05 |
| Lilrb4a | leukocyte immunoglobulin-like receptor, subfamily B, member 4A [Source:MGI Symbol;Acc:MGI:102701] | 2.80 | 7.87E-05 |
| Ceacam16 | CEA cell adhesion molecule 16 [Source:MGI Symbol;Acc:MGI:2685615] | 4.40 | 7.94E-05 |
| Ugt1a7c | UDP glucuronosyltransferase 1 family, polypeptide A7C [Source:MGI Symbol;Acc:MGI:3032636] | 1.67 | 8.00E-05 |
| Asns | asparagine synthetase [Source:MGI Symbol;Acc:MGI:1350929] | 1.10 | 8.22E-05 |
| Ccl2 | C-C motif chemokine ligand 2 [Source:MGI Symbol;Acc:MGI:98259] | 4.18 | 8.29E-05 |
| Ly6g | lymphocyte antigen 6 family member G [Source:MGI Symbol;Acc:MGI:109440] | 7.62 | 8.76E-05 |
| Gm5414 | predicted gene 5414 [Source:MGI Symbol;Acc:MGI:3646939] | 7.55 | 8.76E-05 |
| Cyp7b1 | cytochrome P450, family 7, subfamily b, polypeptide 1 [Source:MGI Symbol;Acc:MGI:104978] | 2.74 | 8.84E-05 |
| Tpsab1 | tryptase alpha/beta 1 [Source:MGI Symbol;Acc:MGI:96943] | 4.21 | 8.86E-05 |
| Itpk1 | inositol 1,3,4-triphosphate 5/6 kinase [Source:MGI Symbol;Acc:MGI:2446159] | 1.34 | 8.99E-05 |
| C1qa | complement component 1, q subcomponent, alpha polypeptide [Source:MGI Symbol;Acc:MGI:88223] | 1.99 | 9.50E-05 |
| Serpinb12 | serine (or cysteine) peptidase inhibitor, clade B (ovalbumin), member 12 [Source:MGI Symbol;Acc:MGI:1919119] | 3.85 | 9.52E-05 |
| Txn1 | thioredoxin 1 [Source:MGI Symbol;Acc:MGI:98874] | 1.92 | 9.54E-05 |
| Ckap4 | cytoskeleton-associated protein 4 [Source:MGI Symbol;Acc:MGI:2444926] | 1.74 | 9.65E-05 |
| Cdk5r1 | cyclin dependent kinase 5, regulatory subunit 1 [Source:MGI Symbol;Acc:MGI:101764] | 1.79 | 1.02E-04 |
| Col5a1 | collagen, type V, alpha 1 [Source:MGI Symbol;Acc:MGI:88457] | 2.34 | 1.05E-04 |
| Slc9a5 | solute carrier family 9 (sodium/hydrogen exchanger), member 5 [Source:MGI Symbol;Acc:MGI:2685542] | 2.38 | 1.12E-04 |
| Shf | Src homology 2 domain containing F [Source:MGI Symbol;Acc:MGI:3613669] | 1.92 | 1.14E-04 |
| Slc20a1 | solute carrier family 20, member 1 [Source:MGI Symbol;Acc:MGI:108392] | 1.33 | 1.14E-04 |
| Sh2d3c | SH2 domain containing 3C [Source:MGI Symbol;Acc:MGI:1351631] | 2.30 | 1.15E-04 |
| Impdh1 | inosine monophosphate dehydrogenase 1 [Source:MGI Symbol;Acc:MGI:96567] | 1.71 | 1.17E-04 |
| Matn4 | matrilin 4 [Source:MGI Symbol;Acc:MGI:1328314] | 2.13 | 1.19E-04 |
| Tmem86a | transmembrane protein 86A [Source:MGI Symbol;Acc:MGI:1915143] | 1.38 | 1.22E-04 |
| Fhl2 | four and a half LIM domains 2 [Source:MGI Symbol;Acc:MGI:1338762] | 3.23 | 1.26E-04 |
| Sprr2f | small proline-rich protein 2F [Source:MGI Symbol;Acc:MGI:1330349] | 10.11 | 1.29E-04 |
| Siglecl1 | Siglec family like 1 [Source:MGI Symbol;Acc:MGI:3780678] | 7.12 | 1.33E-04 |
| Socs3 | suppressor of cytokine signaling 3 [Source:MGI Symbol;Acc:MGI:1201791] | 2.42 | 1.34E-04 |
| Ctss | cathepsin S [Source:MGI Symbol;Acc:MGI:107341] | 2.18 | 1.37E-04 |
| Selenom | selenoprotein M [Source:MGI Symbol;Acc:MGI:2149786] | 1.80 | 1.37E-04 |
| Nip7 | NIP7, nucleolar pre-rRNA processing protein [Source:MGI Symbol;Acc:MGI:1913414] | 0.89 | 1.47E-04 |
| Kcnj9 | potassium inwardly-rectifying channel, subfamily J, member 9 [Source:MGI Symbol;Acc:MGI:108007] | 1.65 | 1.51E-04 |
| Mafb | MAF bZIP transcription factor B [Source:MGI Symbol;Acc:MGI:104555] | 2.16 | 1.54E-04 |
| Hyal2 | hyaluronoglucosaminidase 2 [Source:MGI Symbol;Acc:MGI:1196334] | 1.22 | 1.57E-04 |
| Gm20056 | predicted gene, 20056 [Source:MGI Symbol;Acc:MGI:5012241] | 6.71 | 1.58E-04 |
| Trpm2 | transient receptor potential cation channel, subfamily M, member 2 [Source:MGI Symbol;Acc:MGI:1351901] | 2.68 | 1.64E-04 |
| Tspan11 | tetraspanin 11 [Source:MGI Symbol;Acc:MGI:1915748] | 2.26 | 1.64E-04 |
| Endod1 | endonuclease domain containing 1 [Source:MGI Symbol;Acc:MGI:1919196] | 0.99 | 1.72E-04 |
| Gm7463 | predicted gene 7463 [Source:MGI Symbol;Acc:MGI:3645336] | 8.23 | 1.97E-04 |
| Fgr | FGR proto-oncogene, Src family tyrosine kinase [Source:MGI Symbol;Acc:MGI:95527] | 2.65 | 1.98E-04 |
| Tnfrsf1b | tumor necrosis factor receptor superfamily, member 1b [Source:MGI Symbol;Acc:MGI:1314883] | 2.17 | 1.98E-04 |
| Nrg1 | neuregulin 1 [Source:MGI Symbol;Acc:MGI:96083] | 3.09 | 2.01E-04 |
| Sox7 | SRY (sex determining region Y)-box 7 [Source:MGI Symbol;Acc:MGI:98369] | 1.54 | 2.06E-04 |
| Prr7 | proline rich 7 (synaptic) [Source:MGI Symbol;Acc:MGI:3487246] | 2.34 | 2.11E-04 |
| Snph | syntaphilin [Source:MGI Symbol;Acc:MGI:2139270] | 2.27 | 2.18E-04 |
| Lst1 | leukocyte specific transcript 1 [Source:MGI Symbol;Acc:MGI:1096324] | 3.13 | 2.19E-04 |
| Ssbp4 | single stranded DNA binding protein 4 [Source:MGI Symbol;Acc:MGI:1924150] | 1.19 | 2.19E-04 |
| Sox18 | SRY (sex determining region Y)-box 18 [Source:MGI Symbol;Acc:MGI:103559] | 2.02 | 2.23E-04 |
| Efemp2 | epidermal growth factor-containing fibulin-like extracellular matrix protein 2 [Source:MGI Symbol;Acc:MGI:1891209] | 2.24 | 2.26E-04 |
| Tmem132a | transmembrane protein 132A [Source:MGI Symbol;Acc:MGI:2147810] | 2.12 | 2.27E-04 |
| Acot5 | acyl-CoA thioesterase 5 [Source:MGI Symbol;Acc:MGI:2384969] | 2.50 | 2.34E-04 |

| | | | |
|---|---|---|---|
| Tnn | tenascin N [Source:MGI Symbol;Acc:MGI:2665790] | 3.73 | 2.43E-04 |
| Col5a2 | collagen, type V, alpha 2 [Source:MGI Symbol;Acc:MGI:88458] | 2.14 | 2.45E-04 |
| Loxl3 | lysyl oxidase-like 3 [Source:MGI Symbol;Acc:MGI:1337004] | 1.91 | 2.45E-04 |
| Sele | selectin, endothelial cell [Source:MGI Symbol;Acc:MGI:98278] | 3.67 | 2.50E-04 |
| Fbln1 | fibulin 1 [Source:MGI Symbol;Acc:MGI:95487] | 1.27 | 2.53E-04 |
| Galnt6 | polypeptide N-acetylgalactosaminyltransferase 6 [Source:MGI Symbol;Acc:MGI:1891640] | 1.92 | 2.54E-04 |
| Aprt | adenine phosphoribosyl transferase [Source:MGI Symbol;Acc:MGI:88061] | 1.93 | 2.56E-04 |
| Slco1a7 | solute carrier organic anion transporter family, member 1a7 [Source:MGI Symbol;Acc:MGI:3643685] | 7.74 | 2.58E-04 |
| Abcc3 | ATP-binding cassette, sub-family C member 3 [Source:MGI Symbol;Acc:MGI:1923658] | 1.17 | 2.58E-04 |
| Cxcr1 | C-X-C motif chemokine receptor 1 [Source:MGI Symbol;Acc:MGI:2448715] | 7.13 | 2.62E-04 |
| Kif26b | kinesin family member 26B [Source:MGI Symbol;Acc:MGI:2447076] | 2.00 | 2.74E-04 |
| Prg4 | proteoglycan 4 (megakaryocyte stimulating factor, articular superficial zone protein) [Source:MGI Symbol;Acc:MGI:1891344] | 2.95 | 2.80E-04 |
| Sema7a | sema domain, immunoglobulin domain (Ig), and GPI membrane anchor, (semaphorin) 7A [Source:MGI Symbol;Acc:MGI:1306826] | 1.67 | 2.89E-04 |
| Nfasc | neurofascin [Source:MGI Symbol;Acc:MGI:104753] | 2.33 | 2.94E-04 |
| Gsto1 | glutathione S-transferase omega 1 [Source:MGI Symbol;Acc:MGI:1342273] | 1.48 | 2.94E-04 |
| Gabarap | gamma-aminobutyric acid receptor associated protein [Source:MGI Symbol;Acc:MGI:1861742] | 1.10 | 2.95E-04 |
| Tnfaip2 | tumor necrosis factor, alpha-induced protein 2 [Source:MGI Symbol;Acc:MGI:104960] | 2.08 | 3.01E-04 |
| Mrc2 | mannose receptor, C type 2 [Source:MGI Symbol;Acc:MGI:107818] | 1.81 | 3.01E-04 |
| Tmem51 | transmembrane protein 51 [Source:MGI Symbol;Acc:MGI:2384874] | 1.97 | 3.08E-04 |
| Masp1 | MBL associated serine protease 1 [Source:MGI Symbol;Acc:MGI:88492] | 2.12 | 3.13E-04 |
| Krt17 | keratin 17 [Source:MGI Symbol;Acc:MGI:96691] | 1.90 | 3.26E-04 |
| Mmp2 | matrix metallopeptidase 2 [Source:MGI Symbol;Acc:MGI:97009] | 2.22 | 3.26E-04 |
| Epop | elongin BC and polycomb repressive complex 2 associated protein [Source:MGI Symbol;Acc:MGI:2143991] | 3.11 | 3.27E-04 |
| Sec61b | SEC61 translocon subunit beta [Source:MGI Symbol;Acc:MGI:1913462] | 1.55 | 3.32E-04 |
| Flg | filaggrin [Source:MGI Symbol;Acc:MGI:95553] | 4.17 | 3.34E-04 |
| Sprr2a1 | small proline-rich protein 2A1 [Source:MGI Symbol;Acc:MGI:1330350] | 3.27 | 3.47E-04 |
| Ankk1 | ankyrin repeat and kinase domain containing 1 [Source:MGI Symbol;Acc:MGI:3045301] | 6.29 | 3.52E-04 |
| Btbd19 | BTB domain containing 19 [Source:MGI Symbol;Acc:MGI:1925861] | 1.57 | 3.59E-04 |
| Lrfn3 | leucine rich repeat and fibronectin type III domain containing 3 [Source:MGI Symbol;Acc:MGI:2442512] | 1.83 | 3.61E-04 |
| Ctsd | cathepsin D [Source:MGI Symbol;Acc:MGI:88562] | 1.22 | 3.74E-04 |
| Il19 | interleukin 19 [Source:MGI Symbol;Acc:MGI:1890472] | 6.38 | 3.75E-04 |
| Tnfrsf9 | tumor necrosis factor receptor superfamily, member 9 [Source:MGI Symbol;Acc:MGI:1101059] | 2.68 | 3.75E-04 |
| Fam167b | family with sequence similarity 167, member B [Source:MGI Symbol;Acc:MGI:2668032] | 3.15 | 3.82E-04 |
| Lypd3 | Ly6/Plaur domain containing 3 [Source:MGI Symbol;Acc:MGI:1919684] | 1.76 | 3.85E-04 |
| Bcat1 | branched chain aminotransferase 1, cytosolic [Source:MGI Symbol;Acc:MGI:104861] | 2.80 | 3.93E-04 |
| Cebpb | CCAAT/enhancer binding protein beta [Source:MGI Symbol;Acc:MGI:88373] | 2.21 | 3.93E-04 |
| Tpcn2 | two pore segment channel 2 [Source:MGI Symbol;Acc:MGI:2385297] | 1.32 | 3.95E-04 |
| Teddm3 | transmembrane epididymal family member 3 [Source:MGI Symbol;Acc:MGI:1913811] | 2.33 | 3.98E-04 |
| Ccl8 | C-C motif chemokine ligand 8 [Source:MGI Symbol;Acc:MGI:101878] | 1.89 | 3.98E-04 |
| Adra2a | adrenergic receptor, alpha 2a [Source:MGI Symbol;Acc:MGI:87934] | 2.39 | 4.00E-04 |
| Fcrlb | Fc receptor-like B [Source:MGI Symbol;Acc:MGI:3576487] | 4.42 | 4.01E-04 |
| Atp1a1 | ATPase, Na+/K+ transporting, alpha 1 polypeptide [Source:MGI Symbol;Acc:MGI:88105] | 1.60 | 4.01E-04 |
| Tmem119 | transmembrane protein 119 [Source:MGI Symbol;Acc:MGI:2385228] | 2.60 | 4.02E-04 |
| Tfe3 | transcription factor E3 [Source:MGI Symbol;Acc:MGI:98511] | 0.81 | 4.02E-04 |
| Pla2g7 | phospholipase A2, group VII (platelet-activating factor acetylhydrolase, plasma) [Source:MGI Symbol;Acc:MGI:1351327] | 2.20 | 4.09E-04 |
| Twf1 | twinfilin actin binding protein 1 [Source:MGI Symbol;Acc:MGI:1100520] | 1.16 | 4.09E-04 |
| Pax9 | paired box 9 [Source:MGI Symbol;Acc:MGI:97493] | 2.74 | 4.19E-04 |
| Nfkbid | nuclear factor of kappa light polypeptide gene enhancer in B cells inhibitor, delta [Source:MGI Symbol;Acc:MGI:3041243] | 2.80 | 4.22E-04 |
| Loxl1 | lysyl oxidase-like 1 [Source:MGI Symbol;Acc:MGI:106096] | 2.26 | 4.22E-04 |
| Tie1 | tyrosine kinase with immunoglobulin-like and EGF-like domains 1 [Source:MGI Symbol;Acc:MGI:99906] | 1.75 | 4.22E-04 |
| Fcmr | Fc fragment of IgM receptor [Source:MGI Symbol;Acc:MGI:1916419] | 4.02 | 4.24E-04 |
| Crabp2 | cellular retinoic acid binding protein II [Source:MGI Symbol;Acc:MGI:88491] | 1.88 | 4.57E-04 |
| Pgam1 | phosphoglycerate mutase 1 [Source:MGI Symbol;Acc:MGI:97552] | 1.63 | 4.69E-04 |
| Adamts15 | ADAM metallopeptidase with thrombospondin type 1 motif 15 [Source:MGI Symbol;Acc:MGI:2449569] | 2.02 | 4.75E-04 |
| Chpf | chondroitin polymerizing factor [Source:MGI Symbol;Acc:MGI:106576] | 1.82 | 4.75E-04 |
| Celsr3 | cadherin, EGF LAG seven-pass G-type receptor 3 [Source:MGI Symbol;Acc:MGI:1858236] | 3.09 | 4.81E-04 |
| Stom | stomatin [Source:MGI Symbol;Acc:MGI:95403] | 1.44 | 4.81E-04 |
| Tpm4 | tropomyosin 4 [Source:MGI Symbol;Acc:MGI:2449202] | 0.94 | 4.81E-04 |
| Dio2 | deiodinase, iodothyronine, type II [Source:MGI Symbol;Acc:MGI:1338833] | 2.07 | 5.00E-04 |
| Col27a1 | collagen, type XXVII, alpha 1 [Source:MGI Symbol;Acc:MGI:2672118] | 1.96 | 5.04E-04 |
| Spon1 | spondin 1, (f-spondin) extracellular matrix protein [Source:MGI Symbol;Acc:MGI:2385287] | 2.93 | 5.08E-04 |
| Aqp5 | aquaporin 5 [Source:MGI Symbol;Acc:MGI:106215] | 3.29 | 5.10E-04 |
| Fam177a | family with sequence similarity 177, member A [Source:MGI Symbol;Acc:MGI:1920635] | 1.42 | 5.14E-04 |
| Pdpn | podoplanin [Source:MGI Symbol;Acc:MGI:103098] | 2.68 | 5.15E-04 |
| Themis2 | thymocyte selection associated family member 2 [Source:MGI Symbol;Acc:MGI:2446213] | 1.85 | 5.16E-04 |
| Vash1 | vasohibin 1 [Source:MGI Symbol;Acc:MGI:2442543] | 1.92 | 5.18E-04 |
| Ccl4 | C-C motif chemokine ligand 4 [Source:MGI Symbol;Acc:MGI:98261] | 7.78 | 5.26E-04 |
| Glis3 | GLIS family zinc finger 3 [Source:MGI Symbol;Acc:MGI:2444289] | 2.67 | 5.30E-04 |
| Spatc1 | spermatogenesis and centriole associated 1 [Source:MGI Symbol;Acc:MGI:1921531] | 3.18 | 5.36E-04 |

| | | | |
|---|---|---|---|
| Dlgap2 | DLG associated protein 2 [Source:MGI Symbol;Acc:MGI:2443181] | 3.37 | 5.46E-04 |
| Creb3l3 | cAMP responsive element binding protein 3-like 3 [Source:MGI Symbol;Acc:MGI:2384786] | 1.73 | 5.56E-04 |
| Golm1 | golgi membrane protein 1 [Source:MGI Symbol;Acc:MGI:1917329] | 1.62 | 5.59E-04 |
| Ccl7 | C-C motif chemokine ligand 7 [Source:MGI Symbol;Acc:MGI:99512] | 2.82 | 5.61E-04 |
| B3gnt2 | UDP-GlcNAc:betaGal beta-1,3-N-acetylglucosaminyltransferase 2 [Source:MGI Symbol;Acc:MGI:1889505] | 0.91 | 5.70E-04 |
| Sfn | stratifin [Source:MGI Symbol;Acc:MGI:1891831] | 2.15 | 5.84E-04 |
| Ubald2 | UBA-like domain containing 2 [Source:MGI Symbol;Acc:MGI:1914635] | 0.91 | 6.08E-04 |
| Rnase2b | ribonuclease, RNase A family, 2B (liver, eosinophil-derived neurotoxin) [Source:MGI Symbol;Acc:MGI:1858598] | 1.63 | 6.12E-04 |
| Slfn10-ps | schlafen 10, pseudogene [Source:MGI Symbol;Acc:MGI:3512288] | 1.80 | 6.18E-04 |
| Adamts7 | ADAM metallopeptidase with thrombospondin type 1 motif 7 [Source:MGI Symbol;Acc:MGI:1347346] | 2.24 | 6.21E-04 |
| Ctsz | cathepsin Z [Source:MGI Symbol;Acc:MGI:1891190] | 1.58 | 6.24E-04 |
| Esd | esterase D/formylglutathione hydrolase [Source:MGI Symbol;Acc:MGI:95421] | 1.13 | 6.30E-04 |
| Bbln | bublin coiled coil protein [Source:MGI Symbol;Acc:MGI:1920987] | 1.61 | 6.31E-04 |
| Bgn | biglycan [Source:MGI Symbol;Acc:MGI:88158] | 2.40 | 6.35E-04 |
| Serpinb1a | serine (or cysteine) peptidase inhibitor, clade B, member 1a [Source:MGI Symbol;Acc:MGI:1913472] | 1.78 | 6.44E-04 |
| Pdia6 | protein disulfide isomerase associated 6 [Source:MGI Symbol;Acc:MGI:1919103] | 0.92 | 6.56E-04 |
| Pcsk5 | proprotein convertase subtilisin/kexin type 5 [Source:MGI Symbol;Acc:MGI:97515] | 3.05 | 6.61E-04 |
| Gm11597 | predicted gene 11597 [Source:MGI Symbol;Acc:MGI:3652313] | 4.89 | 6.62E-04 |
| Emilin1 | elastin microfibril interfacer 1 [Source:MGI Symbol;Acc:MGI:1926189] | 2.31 | 6.66E-04 |
| Col4a2 | collagen, type IV, alpha 2 [Source:MGI Symbol;Acc:MGI:88455] | 1.94 | 6.66E-04 |
| Tarm1 | T cell-interacting, activating receptor on myeloid cells 1 [Source:MGI Symbol;Acc:MGI:2442280] | 2.67 | 6.66E-04 |
| Cfl1 | cofilin 1, non-muscle [Source:MGI Symbol;Acc:MGI:101757] | 1.19 | 6.67E-04 |
| Pira12 | paired-Ig-like receptor A12 [Source:MGI Symbol;Acc:MGI:3709645] | 3.17 | 6.69E-04 |
| Il3ra | interleukin 3 receptor, alpha chain [Source:MGI Symbol;Acc:MGI:96553] | 1.58 | 6.77E-04 |
| Klk8 | kallikrein related-peptidase 8 [Source:MGI Symbol;Acc:MGI:1343327] | 2.49 | 6.88E-04 |
| Sat1 | spermidine/spermine N1-acetyl transferase 1 [Source:MGI Symbol;Acc:MGI:98233] | 1.27 | 6.95E-04 |
| Thbd | thrombomodulin [Source:MGI Symbol;Acc:MGI:98736] | 1.32 | 7.16E-04 |
| Spire2 | spire type actin nucleation factor 2 [Source:MGI Symbol;Acc:MGI:2446256] | 2.19 | 7.23E-04 |
| C1qb | complement component 1, q subcomponent, beta polypeptide [Source:MGI Symbol;Acc:MGI:88224] | 1.68 | 7.24E-04 |
| Kcp | kielin/chordin-like protein [Source:MGI Symbol;Acc:MGI:2141640] | 1.48 | 7.24E-04 |
| Lat2 | linker for activation of T cells family, member 2 [Source:MGI Symbol;Acc:MGI:1926479] | 1.80 | 7.34E-04 |
| Enah | ENAH actin regulator [Source:MGI Symbol;Acc:MGI:108360] | 2.06 | 7.45E-04 |
| Igsf3 | immunoglobulin superfamily, member 3 [Source:MGI Symbol;Acc:MGI:1926158] | 1.84 | 7.54E-04 |
| Col6a2 | collagen, type VI, alpha 2 [Source:MGI Symbol;Acc:MGI:88460] | 2.18 | 7.89E-04 |
| Pstpip1 | proline-serine-threonine phosphatase-interacting protein 1 [Source:MGI Symbol;Acc:MGI:1321396] | 1.56 | 7.89E-04 |
| Fcer1a | Fc receptor, IgE, high affinity I, alpha polypeptide [Source:MGI Symbol;Acc:MGI:95494] | 3.85 | 8.02E-04 |
| Actg1 | actin, gamma, cytoplasmic 1 [Source:MGI Symbol;Acc:MGI:87906] | 0.91 | 8.21E-04 |
| Sstr3 | somatostatin receptor 3 [Source:MGI Symbol;Acc:MGI:98329] | 2.19 | 8.21E-04 |
| Gm8080 | predicted gene 8080 [Source:MGI Symbol;Acc:MGI:3646133] | 3.77 | 8.24E-04 |
| Slc43a2 | solute carrier family 43, member 2 [Source:MGI Symbol;Acc:MGI:2442746] | 0.84 | 8.30E-04 |
| Aldh1a2 | aldehyde dehydrogenase family 1, subfamily A2 [Source:MGI Symbol;Acc:MGI:107928] | 3.15 | 8.36E-04 |
| Tmem95 | transmembrane protein 95 [Source:MGI Symbol;Acc:MGI:3779488] | 2.32 | 8.36E-04 |
| Mmp9 | matrix metallopeptidase 9 [Source:MGI Symbol;Acc:MGI:97011] | 2.03 | 8.36E-04 |
| Adm2 | adrenomedullin 2 [Source:MGI Symbol;Acc:MGI:2675256] | 3.75 | 8.38E-04 |
| Garem2 | GRB2 associated regulator of MAPK1 subtype 2 [Source:MGI Symbol;Acc:MGI:2685290] | 3.05 | 8.39E-04 |
| Crlf2 | cytokine receptor-like factor 2 [Source:MGI Symbol;Acc:MGI:1889506] | 2.15 | 8.39E-04 |
| Gpc1 | glypican 1 [Source:MGI Symbol;Acc:MGI:1194891] | 1.22 | 8.39E-04 |
| Hgf | hepatocyte growth factor [Source:MGI Symbol;Acc:MGI:96079] | 3.16 | 8.52E-04 |
| Slc39a2 | solute carrier family 39 (zinc transporter), member 2 [Source:MGI Symbol;Acc:MGI:2684326] | 1.33 | 8.79E-04 |
| Itgam | integrin alpha M [Source:MGI Symbol;Acc:MGI:96607] | 2.32 | 8.84E-04 |
| Tnfrsf12a | tumor necrosis factor receptor superfamily, member 12a [Source:MGI Symbol;Acc:MGI:1351484] | 1.83 | 8.84E-04 |
| Mefv | Mediterranean fever [Source:MGI Symbol;Acc:MGI:1859396] | 2.73 | 8.90E-04 |
| Tlnrd1 | talin rod domain containing 1 [Source:MGI Symbol;Acc:MGI:1891420] | 1.45 | 8.91E-04 |
| C1qtnf6 | C1q and tumor necrosis factor related protein 6 [Source:MGI Symbol;Acc:MGI:1919959] | 2.42 | 9.35E-04 |
| Slfn2 | schlafen 2 [Source:MGI Symbol;Acc:MGI:1313258] | 1.88 | 9.37E-04 |
| Col1a1 | collagen, type I, alpha 1 [Source:MGI Symbol;Acc:MGI:88467] | 1.77 | 9.47E-04 |
| Defb3 | defensin beta 3 [Source:MGI Symbol;Acc:MGI:1351612] | 7.06 | 9.52E-04 |
| Mdk | midkine [Source:MGI Symbol;Acc:MGI:96949] | 1.74 | 9.52E-04 |
| Sbno2 | strawberry notch 2 [Source:MGI Symbol;Acc:MGI:2448490] | 1.37 | 9.55E-04 |
| Tlr2 | toll-like receptor 2 [Source:MGI Symbol;Acc:MGI:1346060] | 2.57 | 9.71E-04 |
| Tmem156 | transmembrane protein 156 [Source:MGI Symbol;Acc:MGI:2685292] | 2.46 | 9.71E-04 |
| Tlr6 | toll-like receptor 6 [Source:MGI Symbol;Acc:MGI:1341296] | 2.05 | 9.75E-04 |
| Myd88 | myeloid differentiation primary response gene 88 [Source:MGI Symbol;Acc:MGI:108005] | 1.25 | 9.83E-04 |
| Gpr65 | G-protein coupled receptor 65 [Source:MGI Symbol;Acc:MGI:108031] | 3.00 | 9.91E-04 |
| Hk3 | hexokinase 3 [Source:MGI Symbol;Acc:MGI:2670962] | 2.32 | 9.98E-04 |
| Zfp469 | zinc finger protein 469 [Source:MGI Symbol;Acc:MGI:2684868] | 2.07 | 9.98E-04 |
| Megf6 | multiple EGF-like-domains 6 [Source:MGI Symbol;Acc:MGI:1919351] | 1.81 | 1.02E-03 |
| Slc37a2 | solute carrier family 37 (glycerol-3-phosphate transporter), member 2 [Source:MGI Symbol;Acc:MGI:1929693] | 1.81 | 1.03E-03 |
| Cmtm1 | CKLF-like MARVEL transmembrane domain containing 1 [Source:MGI Symbol;Acc:MGI:2447159] | 6.72 | 1.03E-03 |
| Cdk2ap2 | cyclin dependent kinase 2 associated protein 2 [Source:MGI Symbol;Acc:MGI:1098779] | 1.39 | 1.04E-03 |
| Grwd1 | glutamate-rich WD repeat containing 1 [Source:MGI Symbol;Acc:MGI:2141989] | 1.50 | 1.05E-03 |

| | | | |
|---|---|---|---|
| Nav3 | neuron navigator 3 [Source:MGI Symbol;Acc:MGI:2183703] | 1.13 | 1.08E-03 |
| Igfn1 | immunoglobulin-like and fibronectin type III domain containing 1 [Source:MGI Symbol;Acc:MGI:3045352] | 5.23 | 1.10E-03 |
| Gm12481 | predicted gene 12481 [Source:MGI Symbol;Acc:MGI:3650743] | 1.71 | 1.11E-03 |
| Tcirg1 | T cell, immune regulator 1, ATPase, H+ transporting, lysosomal V0 protein A3 [Source:MGI Symbol;Acc:MGI:1350931] | 1.47 | 1.11E-03 |
| Endou | endonuclease, polyU-specific [Source:MGI Symbol;Acc:MGI:97746] | 2.79 | 1.11E-03 |
| Ncf4 | neutrophil cytosolic factor 4 [Source:MGI Symbol;Acc:MGI:109186] | 2.31 | 1.11E-03 |
| Gad1 | glutamate decarboxylase 1 [Source:MGI Symbol;Acc:MGI:95632] | 3.90 | 1.12E-03 |
| Atp6v0c | ATPase, H+ transporting, lysosomal V0 subunit C [Source:MGI Symbol;Acc:MGI:88116] | 1.39 | 1.12E-03 |
| Gm9816 | predicted pseudogene 9816 [Source:MGI Symbol;Acc:MGI:3704465] | 7.43 | 1.13E-03 |
| Cstdc4 | cystatin domain containing 4 [Source:MGI Symbol;Acc:MGI:3645124] | 2.65 | 1.13E-03 |
| Tmprss11b | transmembrane protease, serine 11B [Source:MGI Symbol;Acc:MGI:2442893] | 2.31 | 1.13E-03 |
| Ppif | peptidylprolyl isomerase F (cyclophilin F) [Source:MGI Symbol;Acc:MGI:2145814] | 1.57 | 1.13E-03 |
| Cd300lb | CD300 molecule like family member B [Source:MGI Symbol;Acc:MGI:2685099] | 2.36 | 1.14E-03 |
| Il36g | interleukin 36G [Source:MGI Symbol;Acc:MGI:2449929] | 1.60 | 1.14E-03 |
| F5 | coagulation factor V [Source:MGI Symbol;Acc:MGI:88382] | 2.42 | 1.17E-03 |
| Or6c69b | olfactory receptor family 6 subfamily C member 69B [Source:MGI Symbol;Acc:MGI:3030644] | 3.07 | 1.18E-03 |
| Timm13 | translocase of inner mitochondrial membrane 13 [Source:MGI Symbol;Acc:MGI:1353432] | 1.62 | 1.20E-03 |
| Glipr2 | GLI pathogenesis-related 2 [Source:MGI Symbol;Acc:MGI:1917770] | 2.04 | 1.21E-03 |
| C3ar1 | complement component 3a receptor 1 [Source:MGI Symbol;Acc:MGI:1097680] | 1.90 | 1.25E-03 |
| Itgb2 | integrin beta 2 [Source:MGI Symbol;Acc:MGI:96611] | 2.04 | 1.26E-03 |
| Vcam1 | vascular cell adhesion molecule 1 [Source:MGI Symbol;Acc:MGI:98926] | 1.49 | 1.27E-03 |
| Rprm | reprimo, TP53 dependent G2 arrest mediator candidate [Source:MGI Symbol;Acc:MGI:1915124] | 6.11 | 1.29E-03 |
| Gm13398 | predicted gene 13398 [Source:MGI Symbol;Acc:MGI:3652315] | 1.72 | 1.30E-03 |
| Dusp9 | dual specificity phosphatase 9 [Source:MGI Symbol;Acc:MGI:2387107] | 1.83 | 1.31E-03 |
| Col13a1 | collagen, type XIII, alpha 1 [Source:MGI Symbol;Acc:MGI:1277201] | 1.56 | 1.31E-03 |
| Pdia3 | protein disulfide isomerase associated 3 [Source:MGI Symbol;Acc:MGI:95834] | 0.74 | 1.35E-03 |
| Tfec | transcription factor EC [Source:MGI Symbol;Acc:MGI:1333760] | 2.63 | 1.36E-03 |
| Ada | adenosine deaminase [Source:MGI Symbol;Acc:MGI:87916] | 1.41 | 1.41E-03 |
| Il10ra | interleukin 10 receptor, alpha [Source:MGI Symbol;Acc:MGI:96538] | 2.03 | 1.42E-03 |
| Rcn3 | reticulocalbin 3, EF-hand calcium binding domain [Source:MGI Symbol;Acc:MGI:1277122] | 1.66 | 1.42E-03 |
| Gjb1 | gap junction protein, beta 1 [Source:MGI Symbol;Acc:MGI:95719] | 3.47 | 1.44E-03 |
| Col5a3 | collagen, type V, alpha 3 [Source:MGI Symbol;Acc:MGI:1858212] | 2.45 | 1.44E-03 |
| Porcn | porcupine O-acyltransferase [Source:MGI Symbol;Acc:MGI:1890212] | 1.85 | 1.44E-03 |
| Rtn4rl2 | reticulon 4 receptor-like 2 [Source:MGI Symbol;Acc:MGI:2669796] | 2.61 | 1.44E-03 |
| Tmem200b | transmembrane protein 200B [Source:MGI Symbol;Acc:MGI:3646343] | 2.01 | 1.46E-03 |
| Cxcl5 | C-X-C motif chemokine ligand 5 [Source:MGI Symbol;Acc:MGI:1096868] | 3.24 | 1.47E-03 |
| Krt42 | keratin 42 [Source:MGI Symbol;Acc:MGI:1915489] | 4.69 | 1.48E-03 |
| Timm8a1 | translocase of inner mitochondrial membrane 8A1 [Source:MGI Symbol;Acc:MGI:1353433] | 0.93 | 1.49E-03 |
| Csrnp1 | cysteine-serine-rich nuclear protein 1 [Source:MGI Symbol;Acc:MGI:2387989] | 1.73 | 1.50E-03 |
| Pilrb1 | paired immunoglobin-like type 2 receptor beta 1 [Source:MGI Symbol;Acc:MGI:2450532] | 3.25 | 1.54E-03 |
| Gngt2 | guanine nucleotide binding protein (G protein), gamma transducing activity polypeptide 2 [Source:MGI Symbol;Acc:MGI:893584] | 2.10 | 1.54E-03 |
| Hmga2 | high mobility group AT-hook 2 [Source:MGI Symbol;Acc:MGI:101761] | 1.60 | 1.54E-03 |
| Ccin | calicin [Source:MGI Symbol;Acc:MGI:3045316] | 6.00 | 1.54E-03 |
| Tmsb10 | thymosin beta 10 [Source:MGI Symbol;Acc:MGI:109146] | 1.60 | 1.57E-03 |
| Gadd45a | growth arrest and DNA-damage-inducible 45 alpha [Source:MGI Symbol;Acc:MGI:107799] | 1.50 | 1.57E-03 |
| Ddx39a | DEAD box helicase 39a [Source:MGI Symbol;Acc:MGI:1915528] | 1.26 | 1.57E-03 |
| Tnfaip6 | tumor necrosis factor alpha induced protein 6 [Source:MGI Symbol;Acc:MGI:1195266] | 2.78 | 1.59E-03 |
| Ran | RAN, member RAS oncogene family [Source:MGI Symbol;Acc:MGI:1333112] | 1.62 | 1.60E-03 |
| Tomm5 | translocase of outer mitochondrial membrane 5 [Source:MGI Symbol;Acc:MGI:1915762] | 1.06 | 1.60E-03 |
| Sertad1 | SERTA domain containing 1 [Source:MGI Symbol;Acc:MGI:1913438] | 1.56 | 1.61E-03 |
| Ms4a6d | membrane-spanning 4-domains, subfamily A, member 6D [Source:MGI Symbol;Acc:MGI:1916024] | 2.47 | 1.61E-03 |
| Pcdhgc3 | protocadherin gamma subfamily C, 3 [Source:MGI Symbol;Acc:MGI:1935201] | 1.25 | 1.61E-03 |
| Gng11 | guanine nucleotide binding protein (G protein), gamma 11 [Source:MGI Symbol;Acc:MGI:1913316] | 1.87 | 1.64E-03 |
| Calr | calreticulin [Source:MGI Symbol;Acc:MGI:88252] | 0.67 | 1.64E-03 |
| Cebpd | CCAAT/enhancer binding protein delta [Source:MGI Symbol;Acc:MGI:103573] | 2.01 | 1.65E-03 |
| Lpin2 | lipin 2 [Source:MGI Symbol;Acc:MGI:1891341] | 1.01 | 1.66E-03 |
| Fam83e | family with sequence similarity 83, member E [Source:MGI Symbol;Acc:MGI:1921063] | 1.47 | 1.68E-03 |
| Dnajb11 | DnaJ heat shock protein family (Hsp40) member B11 [Source:MGI Symbol;Acc:MGI:1915088] | 0.78 | 1.68E-03 |
| Inpp5d | inositol polyphosphate-5-phosphatase D [Source:MGI Symbol;Acc:MGI:107357] | 1.70 | 1.70E-03 |
| Slc5a1 | solute carrier family 5 (sodium/glucose cotransporter), member 1 [Source:MGI Symbol;Acc:MGI:107678] | 1.60 | 1.70E-03 |
| Tm4sf19 | transmembrane 4 L six family member 19 [Source:MGI Symbol;Acc:MGI:3645933] | 2.75 | 1.71E-03 |
| Slc25a39 | solute carrier family 25, member 39 [Source:MGI Symbol;Acc:MGI:1196386] | 1.26 | 1.72E-03 |
| Ctsa | cathepsin A [Source:MGI Symbol;Acc:MGI:97748] | 1.22 | 1.72E-03 |
| Has3 | hyaluronan synthase 3 [Source:MGI Symbol;Acc:MGI:109599] | 1.56 | 1.72E-03 |
| Plvap | plasmalemma vesicle associated protein [Source:MGI Symbol;Acc:MGI:1890497] | 2.05 | 1.73E-03 |
| Col6a1 | collagen, type VI, alpha 1 [Source:MGI Symbol;Acc:MGI:88459] | 2.10 | 1.74E-03 |
| Pmepa1 | prostate transmembrane protein, androgen induced 1 [Source:MGI Symbol;Acc:MGI:1929600] | 1.98 | 1.74E-03 |
| Vav1 | vav 1 oncogene [Source:MGI Symbol;Acc:MGI:98923] | 1.73 | 1.74E-03 |
| Mrgpra2a | MAS-related GPR, member A2A [Source:MGI Symbol;Acc:MGI:3821888] | 6.04 | 1.76E-03 |
| Ugcg | UDP-glucose ceramide glucosyltransferase [Source:MGI Symbol;Acc:MGI:1332243] | 0.99 | 1.80E-03 |
| Tdh | L-threonine dehydrogenase [Source:MGI Symbol;Acc:MGI:1926231] | 3.80 | 1.83E-03 |

| | | | |
|---|---|---|---|
| Spred3 | sprouty-related EVH1 domain containing 3 [Source:MGI Symbol;Acc:MGI:2142186] | 1.03 | 1.88E-03 |
| Ppp1r14b | protein phosphatase 1, regulatory inhibitor subunit 14B [Source:MGI Symbol;Acc:MGI:107682] | 0.96 | 1.88E-03 |
| Ltf | lactotransferrin [Source:MGI Symbol;Acc:MGI:96837] | 5.64 | 1.90E-03 |
| Slco1a5 | solute carrier organic anion transporter family, member 1a5 [Source:MGI Symbol;Acc:MGI:1351865] | 3.19 | 1.90E-03 |
| Serpina9 | serine (or cysteine) peptidase inhibitor, clade A (alpha-1 antiproteinase, antitrypsin), member 9 [Source:MGI Symbol;Acc:MGI:1919157] | 2.62 | 1.90E-03 |
| Pirb | paired Ig-like receptor B [Source:MGI Symbol;Acc:MGI:894311] | 1.98 | 1.90E-03 |
| Col3a1 | collagen, type III, alpha 1 [Source:MGI Symbol;Acc:MGI:88453] | 1.76 | 1.90E-03 |
| Gpr153 | G protein-coupled receptor 153 [Source:MGI Symbol;Acc:MGI:1916157] | 1.87 | 1.92E-03 |
| Rplp0-ps1 | ribosomal protein lateral stalk subunit P0, pseudogene 1 [Source:MGI Symbol;Acc:MGI:3648599] | 3.72 | 1.93E-03 |
| Stfa2 | stefin A2 [Source:MGI Symbol;Acc:MGI:106197] | 7.80 | 1.93E-03 |
| Ppib | peptidylprolyl isomerase B [Source:MGI Symbol;Acc:MGI:97750] | 0.93 | 1.94E-03 |
| Tubb6 | tubulin, beta 6 class V [Source:MGI Symbol;Acc:MGI:1915201] | 2.21 | 1.94E-03 |
| Cyba | cytochrome b-245, alpha polypeptide [Source:MGI Symbol;Acc:MGI:1316658] | 1.81 | 1.95E-03 |
| Gmppb | GDP-mannose pyrophosphorylase B [Source:MGI Symbol;Acc:MGI:2660880] | 1.52 | 1.95E-03 |
| Myo1b | myosin IB [Source:MGI Symbol;Acc:MGI:107752] | 1.19 | 1.96E-03 |
| Aox4 | aldehyde oxidase 4 [Source:MGI Symbol;Acc:MGI:1919122] | 2.55 | 1.98E-03 |
| Syngr2 | synaptogyrin 2 [Source:MGI Symbol;Acc:MGI:1328324] | 0.94 | 1.99E-03 |
| Or2b11 | olfactory receptor family 2 subfamily B member 11 [Source:MGI Symbol;Acc:MGI:3030056] | 4.75 | 2.01E-03 |
| Cma1 | chymase 1, mast cell [Source:MGI Symbol;Acc:MGI:96941] | 1.39 | 2.02E-03 |
| Syk | spleen tyrosine kinase [Source:MGI Symbol;Acc:MGI:99515] | 1.48 | 2.05E-03 |
| 2300002M23Rik | RIKEN cDNA 2300002M23 gene [Source:MGI Symbol;Acc:MGI:1916792] | 7.15 | 2.06E-03 |
| Gm6211 | predicted gene 6211 [Source:MGI Symbol;Acc:MGI:3646534] | 4.19 | 2.06E-03 |
| Capn2 | calpain 2 [Source:MGI Symbol;Acc:MGI:88264] | 1.20 | 2.06E-03 |
| Col4a1 | collagen, type IV, alpha 1 [Source:MGI Symbol;Acc:MGI:88454] | 1.67 | 2.09E-03 |
| Siah2 | siah E3 ubiquitin protein ligase 2 [Source:MGI Symbol;Acc:MGI:108062] | 0.97 | 2.10E-03 |
| Pofut2 | protein O-fucosyltransferase 2 [Source:MGI Symbol;Acc:MGI:1916863] | 1.56 | 2.15E-03 |
| Vcan | versican [Source:MGI Symbol;Acc:MGI:102889] | 1.92 | 2.22E-03 |
| Ces2e | carboxylesterase 2E [Source:MGI Symbol;Acc:MGI:2443170] | 2.69 | 2.25E-03 |
| Abcb11 | ATP-binding cassette, sub-family B member 11 [Source:MGI Symbol;Acc:MGI:1351619] | 3.28 | 2.26E-03 |
| Map2k3 | mitogen-activated protein kinase kinase 3 [Source:MGI Symbol;Acc:MGI:1346868] | 1.02 | 2.29E-03 |
| Col1a2 | collagen, type I, alpha 2 [Source:MGI Symbol;Acc:MGI:88468] | 1.59 | 2.30E-03 |
| Lrrc8a | leucine rich repeat containing 8A VRAC subunit A [Source:MGI Symbol;Acc:MGI:2652847] | 0.92 | 2.30E-03 |
| Upp1 | uridine phosphorylase 1 [Source:MGI Symbol;Acc:MGI:1097668] | 3.21 | 2.36E-03 |
| Baalc | brain and acute leukemia, cytoplasmic [Source:MGI Symbol;Acc:MGI:1928704] | 2.11 | 2.38E-03 |
| Pfn1 | profilin 1 [Source:MGI Symbol;Acc:MGI:97549] | 0.97 | 2.38E-03 |
| Vash2 | vasohibin 2 [Source:MGI Symbol;Acc:MGI:2444826] | 1.95 | 2.41E-03 |
| Tpd52 | tumor protein D52 [Source:MGI Symbol;Acc:MGI:107749] | 0.83 | 2.41E-03 |
| Kcnk1 | potassium channel, subfamily K, member 1 [Source:MGI Symbol;Acc:MGI:109322] | 2.36 | 2.42E-03 |
| Slc2a1 | solute carrier family 2 (facilitated glucose transporter), member 1 [Source:MGI Symbol;Acc:MGI:95755] | 2.07 | 2.44E-03 |
| Vps37b | vacuolar protein sorting 37B [Source:MGI Symbol;Acc:MGI:1916724] | 1.33 | 2.45E-03 |
| Armcx4 | armadillo repeat containing, X-linked 4 [Source:MGI Symbol;Acc:MGI:2147887] | 2.26 | 2.46E-03 |
| Reg3g | regenerating islet-derived 3 gamma [Source:MGI Symbol;Acc:MGI:109406] | 10.36 | 2.50E-03 |
| Clec5a | C-type lectin domain family 5, member a [Source:MGI Symbol;Acc:MGI:1345151] | 2.83 | 2.50E-03 |
| Plac8 | placenta-specific 8 [Source:MGI Symbol;Acc:MGI:2445289] | 2.29 | 2.50E-03 |
| Capg | capping actin protein, gelsolin like [Source:MGI Symbol;Acc:MGI:1098259] | 1.26 | 2.50E-03 |
| Ccdc107 | coiled-coil domain containing 107 [Source:MGI Symbol;Acc:MGI:1913423] | 1.54 | 2.52E-03 |
| Carmil2 | capping protein regulator and myosin 1 linker 2 [Source:MGI Symbol;Acc:MGI:2685431] | 1.49 | 2.52E-03 |
| Cd300ld | CD300 molecule like family member d [Source:MGI Symbol;Acc:MGI:2442358] | 1.64 | 2.53E-03 |
| Nme2 | NME/NM23 nucleoside diphosphate kinase 2 [Source:MGI Symbol;Acc:MGI:97356] | 0.99 | 2.54E-03 |
| Pdgfra | platelet derived growth factor receptor, alpha polypeptide [Source:MGI Symbol;Acc:MGI:97530] | 2.32 | 2.56E-03 |
| Lyz2 | lysozyme 2 [Source:MGI Symbol;Acc:MGI:96897] | 1.45 | 2.56E-03 |
| Dok3 | docking protein 3 [Source:MGI Symbol;Acc:MGI:1351490] | 1.73 | 2.57E-03 |
| Ptprn | protein tyrosine phosphatase receptor type N [Source:MGI Symbol;Acc:MGI:102765] | 2.19 | 2.58E-03 |
| Prl2c3 | prolactin family 2, subfamily c, member 3 [Source:MGI Symbol;Acc:MGI:1341833] | 1.51 | 2.60E-03 |
| Arhgef25 | Rho guanine nucleotide exchange factor 25 [Source:MGI Symbol;Acc:MGI:1277173] | 1.16 | 2.61E-03 |
| Pgap6 | post-glycosylphosphatidylinositol attachment to proteins 6 [Source:MGI Symbol;Acc:MGI:1926283] | 1.97 | 2.62E-03 |
| Klf18 | Kruppel-like transcription factor 18 [Source:MGI Symbol;Acc:MGI:3651666] | 6.46 | 2.62E-03 |
| Tnf | tumor necrosis factor [Source:MGI Symbol;Acc:MGI:104798] | 2.54 | 2.66E-03 |
| Ccdc102a | coiled-coil domain containing 102A [Source:MGI Symbol;Acc:MGI:2686927] | 1.64 | 2.68E-03 |
| Pitx1 | paired-like homeodomain transcription factor 1 [Source:MGI Symbol;Acc:MGI:107374] | 4.23 | 2.69E-03 |
| Arpc1b | actin related protein 2/3 complex, subunit 1B [Source:MGI Symbol;Acc:MGI:1343142] | 1.36 | 2.69E-03 |
| Snx20 | sorting nexin 20 [Source:MGI Symbol;Acc:MGI:1918857] | 1.70 | 2.73E-03 |
| Gm15583 | predicted gene 15583 [Source:MGI Symbol;Acc:MGI:3783031] | 4.80 | 2.74E-03 |
| Avpi1 | arginine vasopressin-induced 1 [Source:MGI Symbol;Acc:MGI:1916784] | 1.93 | 2.74E-03 |
| Ifi202b | interferon activated gene 202B [Source:MGI Symbol;Acc:MGI:1347083] | 1.31 | 2.74E-03 |
| Ell2 | elongation factor for RNA polymerase II 2 [Source:MGI Symbol;Acc:MGI:2183438] | 0.80 | 2.79E-03 |
| Dnase2b | deoxyribonuclease II beta [Source:MGI Symbol;Acc:MGI:1913283] | 3.52 | 2.79E-03 |
| Ebp | EBP cholestenol delta-isomerase [Source:MGI Symbol;Acc:MGI:107822] | 1.01 | 2.81E-03 |
| Tspan3 | tetraspanin 3 [Source:MGI Symbol;Acc:MGI:1928098] | 0.98 | 2.86E-03 |
| Fyb1 | FYN binding protein 1 [Source:MGI Symbol;Acc:MGI:1346327] | 1.57 | 2.87E-03 |
| Cstb | cystatin B [Source:MGI Symbol;Acc:MGI:109514] | 1.55 | 2.87E-03 |

| | | | |
|---|---|---|---|
| Lce3d | late cornified envelope 3D [Source:MGI Symbol;Acc:MGI:3642919] | 6.60 | 2.89E-03 |
| Eva1b | eva-1 homolog B [Source:MGI Symbol;Acc:MGI:1922063] | 1.92 | 2.91E-03 |
| Syt8 | synaptotagmin VIII [Source:MGI Symbol;Acc:MGI:1859867] | 1.91 | 2.91E-03 |
| Clec7a | C-type lectin domain family 7, member a [Source:MGI Symbol;Acc:MGI:1861431] | 2.56 | 2.94E-03 |
| Xdh | xanthine dehydrogenase [Source:MGI Symbol;Acc:MGI:98973] | 1.10 | 2.96E-03 |
| Scgb1b27 | secretoglobin, family 1B, member 27 [Source:MGI Symbol;Acc:MGI:87862] | 6.15 | 2.97E-03 |
| Prl2c5 | prolactin family 2, subfamily c, member 5 [Source:MGI Symbol;Acc:MGI:1858413] | 4.57 | 2.99E-03 |
| Dynlt4 | dynein light chain Tctex-type 4 [Source:MGI Symbol;Acc:MGI:3045358] | 2.57 | 3.06E-03 |
| Cnfn | cornifelin [Source:MGI Symbol;Acc:MGI:1919633] | 2.50 | 3.06E-03 |
| Coro1a | coronin, actin binding protein 1A [Source:MGI Symbol;Acc:MGI:1345961] | 1.96 | 3.10E-03 |
| Cdk17 | cyclin dependent kinase 17 [Source:MGI Symbol;Acc:MGI:97517] | 1.18 | 3.20E-03 |
| Sema6b | sema domain, transmembrane domain (TM), and cytoplasmic domain, (semaphorin) 6B [Source:MGI Symbol;Acc:MGI:1202889] | 1.95 | 3.20E-03 |
| Nos3 | nitric oxide synthase 3, endothelial cell [Source:MGI Symbol;Acc:MGI:97362] | 1.84 | 3.20E-03 |
| Igsf8 | immunoglobulin superfamily, member 8 [Source:MGI Symbol;Acc:MGI:2154090] | 1.47 | 3.20E-03 |
| Pilra | paired immunoglobin-like type 2 receptor alpha [Source:MGI Symbol;Acc:MGI:2450529] | 2.56 | 3.24E-03 |
| Cox6a1 | cytochrome c oxidase subunit 6A1 [Source:MGI Symbol;Acc:MGI:103099] | 1.61 | 3.28E-03 |
| 5033423K11Rik | RIKEN cDNA 5033423K11 gene [Source:MGI Symbol;Acc:MGI:1923233] | 1.99 | 3.28E-03 |
| Ereg | epiregulin [Source:MGI Symbol;Acc:MGI:107508] | 1.07 | 3.30E-03 |
| Cyp17a1 | cytochrome P450, family 17, subfamily a, polypeptide 1 [Source:MGI Symbol;Acc:MGI:88586] | 1.36 | 3.35E-03 |
| Myo7a | myosin VIIA [Source:MGI Symbol;Acc:MGI:104510] | 2.00 | 3.36E-03 |
| Csf2rb | colony stimulating factor 2 receptor, beta, low-affinity (granulocyte-macrophage) [Source:MGI Symbol;Acc:MGI:1339759] | 2.25 | 3.44E-03 |
| Pus7l | pseudouridylate synthase 7-like [Source:MGI Symbol;Acc:MGI:1926145] | 1.01 | 3.48E-03 |
| Hnf1b | HNF1 homeobox B [Source:MGI Symbol;Acc:MGI:98505] | 6.57 | 3.50E-03 |
| Mfsd10 | major facilitator superfamily domain containing 10 [Source:MGI Symbol;Acc:MGI:1915544] | 0.99 | 3.53E-03 |
| Ccl6 | C-C motif chemokine ligand 6 [Source:MGI Symbol;Acc:MGI:98263] | 1.83 | 3.58E-03 |
| Pcsk9 | proprotein convertase subtilisin/kexin type 9 [Source:MGI Symbol;Acc:MGI:2140260] | 2.11 | 3.75E-03 |
| Scarf2 | scavenger receptor class F, member 2 [Source:MGI Symbol;Acc:MGI:1858430] | 1.88 | 3.75E-03 |
| Sdk1 | sidekick cell adhesion molecule 1 [Source:MGI Symbol;Acc:MGI:2444413] | 2.24 | 3.75E-03 |
| Psmb5 | proteasome (prosome, macropain) subunit, beta type 5 [Source:MGI Symbol;Acc:MGI:1194513] | 1.42 | 3.77E-03 |
| Ctsl | cathepsin L [Source:MGI Symbol;Acc:MGI:88564] | 1.09 | 3.77E-03 |
| Rnf144a | ring finger protein 144A [Source:MGI Symbol;Acc:MGI:1344401] | 2.08 | 3.78E-03 |
| Cgref1 | cell growth regulator with EF hand domain 1 [Source:MGI Symbol;Acc:MGI:1915817] | 1.41 | 3.85E-03 |
| Nek6 | NIMA (never in mitosis gene a)-related expressed kinase 6 [Source:MGI Symbol;Acc:MGI:1891638] | 2.06 | 3.95E-03 |
| Elovl3 | ELOVL fatty acid elongase 3 [Source:MGI Symbol;Acc:MGI:1195976] | 1.61 | 3.95E-03 |
| Ralgds | ral guanine nucleotide dissociation stimulator [Source:MGI Symbol;Acc:MGI:107485] | 1.22 | 4.02E-03 |
| Pitpnm3 | PITPNM family member 3 [Source:MGI Symbol;Acc:MGI:2685726] | 1.38 | 4.03E-03 |
| P3h3 | prolyl 3-hydroxylase 3 [Source:MGI Symbol;Acc:MGI:1315208] | 1.45 | 4.06E-03 |
| Akr1d1 | aldo-keto reductase family 1, member D1 [Source:MGI Symbol;Acc:MGI:2384785] | 2.88 | 4.13E-03 |
| Pgf | placental growth factor [Source:MGI Symbol;Acc:MGI:105095] | 2.20 | 4.18E-03 |
| Gm10163 | predicted pseudogene 10163 [Source:MGI Symbol;Acc:MGI:3704341] | 1.14 | 4.18E-03 |
| Plekho1 | pleckstrin homology domain containing, family O member 1 [Source:MGI Symbol;Acc:MGI:1914470] | 1.55 | 4.26E-03 |
| Kctd11 | potassium channel tetramerisation domain containing 11 [Source:MGI Symbol;Acc:MGI:2448712] | 1.45 | 4.29E-03 |
| Tomm20 | translocase of outer mitochondrial membrane 20 [Source:MGI Symbol;Acc:MGI:1915202] | 0.96 | 4.29E-03 |
| Gm49325 | predicted gene, 49325 [Source:MGI Symbol;Acc:MGI:6121509] | 1.44 | 4.30E-03 |
| Nt5dc2 | 5'-nucleotidase domain containing 2 [Source:MGI Symbol;Acc:MGI:1917271] | 2.09 | 4.34E-03 |
| Sp9 | trans-acting transcription factor 9 [Source:MGI Symbol;Acc:MGI:3574660] | 2.24 | 4.36E-03 |
| Rpl41 | ribosomal protein L41 [Source:MGI Symbol;Acc:MGI:1915195] | 1.78 | 4.36E-03 |
| Srpx2 | sushi-repeat-containing protein, X-linked 2 [Source:MGI Symbol;Acc:MGI:1916042] | 2.70 | 4.36E-03 |
| Atp6v1b2 | ATPase, H+ transporting, lysosomal V1 subunit B2 [Source:MGI Symbol;Acc:MGI:109618] | 0.90 | 4.39E-03 |
| Sult2b1 | sulfotransferase family, cytosolic, 2B, member 1 [Source:MGI Symbol;Acc:MGI:1926342] | 1.13 | 4.45E-03 |
| Tfpi2 | tissue factor pathway inhibitor 2 [Source:MGI Symbol;Acc:MGI:108543] | 3.04 | 4.55E-03 |
| Rpl36al | ribosomal protein L36A-like [Source:MGI Symbol;Acc:MGI:1913733] | 0.73 | 4.55E-03 |
| Ccnb1ip1 | cyclin B1 interacting protein 1 [Source:MGI Symbol;Acc:MGI:2685134] | 1.86 | 4.64E-03 |
| Rasip1 | Ras interacting protein 1 [Source:MGI Symbol;Acc:MGI:1917153] | 1.43 | 4.70E-03 |
| Hmox1 | heme oxygenase 1 [Source:MGI Symbol;Acc:MGI:96163] | 1.02 | 4.73E-03 |
| Tspo | translocator protein [Source:MGI Symbol;Acc:MGI:88222] | 1.83 | 4.79E-03 |
| Adgrg3 | adhesion G protein-coupled receptor G3 [Source:MGI Symbol;Acc:MGI:1859670] | 1.21 | 4.84E-03 |
| Cfap157 | cilia and flagella associated protein 157 [Source:MGI Symbol;Acc:MGI:2447809] | 0.92 | 4.84E-03 |
| Timm22 | translocase of inner mitochondrial membrane 22 [Source:MGI Symbol;Acc:MGI:1929742] | 0.76 | 4.84E-03 |
| En1 | engrailed 1 [Source:MGI Symbol;Acc:MGI:95389] | 2.73 | 4.92E-03 |
| Neu2 | neuraminidase 2 [Source:MGI Symbol;Acc:MGI:1344417] | 1.59 | 4.92E-03 |
| Ms4a14 | membrane-spanning 4-domains, subfamily A, member 14 [Source:MGI Symbol;Acc:MGI:2686122] | 2.41 | 4.92E-03 |
| Lyn | LYN proto-oncogene, Src family tyrosine kinase [Source:MGI Symbol;Acc:MGI:96892] | 1.67 | 4.92E-03 |
| Gm38119 | predicted gene, 38119 [Source:MGI Symbol;Acc:MGI:5611347] | 6.15 | 4.93E-03 |
| Smox | spermine oxidase [Source:MGI Symbol;Acc:MGI:2445356] | 1.79 | 5.00E-03 |
| Kif1a | kinesin family member 1A [Source:MGI Symbol;Acc:MGI:108391] | 1.96 | 5.02E-03 |
| Socs5 | suppressor of cytokine signaling 5 [Source:MGI Symbol;Acc:MGI:2385459] | 0.86 | 5.03E-03 |
| Chrm4 | cholinergic receptor, muscarinic 4 [Source:MGI Symbol;Acc:MGI:88399] | 2.41 | 5.08E-03 |
| Gk | glycerol kinase [Source:MGI Symbol;Acc:MGI:106594] | 1.50 | 5.30E-03 |
| B3gnt9 | UDP-GlcNAc:betaGal beta-1,3-N-acetylglucosaminyltransferase 9 [Source:MGI Symbol;Acc:MGI:2142841] | 1.57 | 5.31E-03 |

| | | | |
|---|---|---|---|
| Emc6 | ER membrane protein complex subunit 6 [Source:MGI Symbol;Acc:MGI:1913298] | 0.76 | 5.43E-03 |
| Saa1 | serum amyloid A 1 [Source:MGI Symbol;Acc:MGI:98221] | 9.05 | 5.46E-03 |
| Uqcr11 | ubiquinol-cytochrome c reductase, complex III subunit XI [Source:MGI Symbol;Acc:MGI:1913844] | 1.71 | 5.46E-03 |
| Galk1 | galactokinase 1 [Source:MGI Symbol;Acc:MGI:95730] | 1.45 | 5.47E-03 |
| Pitpnm1 | phosphatidylinositol transfer protein, membrane-associated 1 [Source:MGI Symbol;Acc:MGI:1197524] | 1.51 | 5.60E-03 |
| Gpsm3 | G-protein signalling modulator 3 (AGS3-like, C. elegans) [Source:MGI Symbol;Acc:MGI:2146785] | 1.70 | 5.62E-03 |
| Thbs2 | thrombospondin 2 [Source:MGI Symbol;Acc:MGI:98738] | 2.07 | 5.65E-03 |
| Cd63 | CD63 antigen [Source:MGI Symbol;Acc:MGI:99529] | 1.60 | 5.66E-03 |
| Soat1 | sterol O-acyltransferase 1 [Source:MGI Symbol;Acc:MGI:104665] | 0.98 | 5.69E-03 |
| Gm8288 | predicted gene 8288 [Source:MGI Symbol;Acc:MGI:3645459] | 5.75 | 5.70E-03 |
| Map7d1 | MAP7 domain containing 1 [Source:MGI Symbol;Acc:MGI:2384297] | 1.46 | 5.70E-03 |
| Kcnd1 | potassium voltage-gated channel, Shal-related family, member 1 [Source:MGI Symbol;Acc:MGI:96671] | 1.31 | 5.70E-03 |
| Nlrp12 | NLR family, pyrin domain containing 12 [Source:MGI Symbol;Acc:MGI:2676630] | 2.00 | 5.73E-03 |
| Comtd1 | catechol-O-methyltransferase domain containing 1 [Source:MGI Symbol;Acc:MGI:1916406] | 1.46 | 5.81E-03 |
| Kcng2 | potassium voltage-gated channel, subfamily G, member 2 [Source:MGI Symbol;Acc:MGI:3694646] | 1.33 | 5.84E-03 |
| AA467197 | expressed sequence AA467197 [Source:MGI Symbol;Acc:MGI:3034182] | 2.04 | 5.85E-03 |
| Spsb1 | splA/ryanodine receptor domain and SOCS box containing 1 [Source:MGI Symbol;Acc:MGI:1921896] | 2.03 | 5.85E-03 |
| Pdia5 | protein disulfide isomerase associated 5 [Source:MGI Symbol;Acc:MGI:1919849] | 1.14 | 5.85E-03 |
| Tmed1 | transmembrane p24 trafficking protein 1 [Source:MGI Symbol;Acc:MGI:106201] | 0.89 | 5.85E-03 |
| Apol9b | apolipoprotein L 9b [Source:MGI Symbol;Acc:MGI:1919148] | 2.12 | 5.88E-03 |
| Nos2 | nitric oxide synthase 2, inducible [Source:MGI Symbol;Acc:MGI:97361] | 3.16 | 5.93E-03 |
| Dhx58 | DExH-box helicase 58 [Source:MGI Symbol;Acc:MGI:1931560] | 1.40 | 5.95E-03 |
| Nherf1 | NHERF family PDZ scaffold protein 1 [Source:MGI Symbol;Acc:MGI:1349482] | 1.63 | 5.95E-03 |
| Map2k1 | mitogen-activated protein kinase kinase 1 [Source:MGI Symbol;Acc:MGI:1346866] | 1.00 | 6.03E-03 |
| Muc19 | mucin 19 [Source:MGI Symbol;Acc:MGI:2676278] | 2.57 | 6.19E-03 |
| Fmnl3 | formin-like 3 [Source:MGI Symbol;Acc:MGI:109569] | 1.56 | 6.19E-03 |
| Junb | jun B proto-oncogene [Source:MGI Symbol;Acc:MGI:96647] | 1.55 | 6.19E-03 |
| Hspg2 | perlecan (heparan sulfate proteoglycan 2) [Source:MGI Symbol;Acc:MGI:96257] | 1.43 | 6.19E-03 |
| Myh9 | myosin, heavy polypeptide 9, non-muscle [Source:MGI Symbol;Acc:MGI:107717] | 0.98 | 6.36E-03 |
| Nadk | NAD kinase [Source:MGI Symbol;Acc:MGI:2183149] | 0.77 | 6.41E-03 |
| Arhgef2 | Rho/Rac guanine nucleotide exchange factor 2 [Source:MGI Symbol;Acc:MGI:103264] | 1.09 | 6.42E-03 |
| Plscr1 | phospholipid scramblase 1 [Source:MGI Symbol;Acc:MGI:893575] | 1.97 | 6.44E-03 |
| Lrp1 | low density lipoprotein receptor-related protein 1 [Source:MGI Symbol;Acc:MGI:96828] | 1.39 | 6.44E-03 |
| Fbn2 | fibrillin 2 [Source:MGI Symbol;Acc:MGI:95490] | 1.18 | 6.44E-03 |
| Rhoc | ras homolog family member C [Source:MGI Symbol;Acc:MGI:106028] | 1.68 | 6.48E-03 |
| Ctsc | cathepsin C [Source:MGI Symbol;Acc:MGI:109553] | 1.22 | 6.48E-03 |
| Tbx2 | T-box 2 [Source:MGI Symbol;Acc:MGI:98494] | 1.34 | 6.49E-03 |
| Apobec1 | apolipoprotein B mRNA editing enzyme, catalytic polypeptide 1 [Source:MGI Symbol;Acc:MGI:103298] | 1.03 | 6.58E-03 |
| Selenon | selenoprotein N [Source:MGI Symbol;Acc:MGI:2151208] | 1.60 | 6.65E-03 |
| Pdgfrb | platelet derived growth factor receptor, beta polypeptide [Source:MGI Symbol;Acc:MGI:97531] | 1.97 | 6.69E-03 |
| Pik3ap1 | phosphoinositide-3-kinase adaptor protein 1 [Source:MGI Symbol;Acc:MGI:1933177] | 1.64 | 6.69E-03 |
| Gm6815 | predicted gene 6815 [Source:MGI Symbol;Acc:MGI:3648348] | 5.83 | 6.72E-03 |
| Mgam | maltase-glucoamylase [Source:MGI Symbol;Acc:MGI:1203495] | 3.08 | 6.73E-03 |
| Scand1 | SCAN domain-containing 1 [Source:MGI Symbol;Acc:MGI:1343132] | 1.84 | 6.79E-03 |
| Slc2a10 | solute carrier family 2 (facilitated glucose transporter), member 10 [Source:MGI Symbol;Acc:MGI:2156687] | 1.36 | 6.79E-03 |
| Mmp14 | matrix metallopeptidase 14 (membrane-inserted) [Source:MGI Symbol;Acc:MGI:101900] | 1.57 | 6.84E-03 |
| Wfdc17 | WAP four-disulfide core domain 17 [Source:MGI Symbol;Acc:MGI:3649773] | 2.27 | 6.89E-03 |
| Ier5 | immediate early response 5 [Source:MGI Symbol;Acc:MGI:1337072] | 1.88 | 6.89E-03 |
| Il10 | interleukin 10 [Source:MGI Symbol;Acc:MGI:96537] | 2.31 | 6.89E-03 |
| Igf2r | insulin-like growth factor 2 receptor [Source:MGI Symbol;Acc:MGI:96435] | 0.84 | 6.97E-03 |
| Raet1d | retinoic acid early transcript delta [Source:MGI Symbol;Acc:MGI:1861032] | 5.71 | 7.12E-03 |
| Pdia4 | protein disulfide isomerase associated 4 [Source:MGI Symbol;Acc:MGI:104864] | 0.79 | 7.16E-03 |
| Ak5 | adenylate kinase 5 [Source:MGI Symbol;Acc:MGI:2677491] | 1.48 | 7.17E-03 |
| Pcdhgc5 | protocadherin gamma subfamily C, 5 [Source:MGI Symbol;Acc:MGI:1935205] | 1.28 | 7.20E-03 |
| Bmp1 | bone morphogenetic protein 1 [Source:MGI Symbol;Acc:MGI:88176] | 1.69 | 7.22E-03 |
| Ptgir | prostaglandin I receptor (IP) [Source:MGI Symbol;Acc:MGI:99535] | 1.96 | 7.24E-03 |
| Il27ra | interleukin 27 receptor, alpha [Source:MGI Symbol;Acc:MGI:1355318] | 1.66 | 7.24E-03 |
| Serpinb3d | serine (or cysteine) peptidase inhibitor, clade B (ovalbumin), member 3D [Source:MGI Symbol;Acc:MGI:2683295] | 5.50 | 7.25E-03 |
| Ccdc124 | coiled-coil domain containing 124 [Source:MGI Symbol;Acc:MGI:1916403] | 1.16 | 7.25E-03 |
| Zdhhc14 | zinc finger, DHHC domain containing 14 [Source:MGI Symbol;Acc:MGI:2653229] | 1.32 | 7.26E-03 |
| Socs1 | suppressor of cytokine signaling 1 [Source:MGI Symbol;Acc:MGI:1354910] | 1.88 | 7.29E-03 |
| Nmt1 | N-myristoyltransferase 1 [Source:MGI Symbol;Acc:MGI:102579] | 0.64 | 7.32E-03 |
| Flnb | filamin, beta [Source:MGI Symbol;Acc:MGI:2446089] | 1.14 | 7.37E-03 |
| Elk3 | ELK3, member of ETS oncogene family [Source:MGI Symbol;Acc:MGI:101762] | 1.67 | 7.43E-03 |
| Mfge8 | milk fat globule EGF and factor V/VIII domain containing [Source:MGI Symbol;Acc:MGI:102768] | 1.01 | 7.43E-03 |
| Cct3 | chaperonin containing TCP1 subunit 3 [Source:MGI Symbol;Acc:MGI:104708] | 1.10 | 7.45E-03 |
| Gm13889 | predicted gene 13889 [Source:MGI Symbol;Acc:MGI:3652053] | 1.63 | 7.47E-03 |
| Pdcd1 | programmed cell death 1 [Source:MGI Symbol;Acc:MGI:104879] | 5.06 | 7.54E-03 |
| Gm11956 | predicted gene 11956 [Source:MGI Symbol;Acc:MGI:3649806] | 5.83 | 7.58E-03 |
| Fcgr1 | Fc receptor, IgG, high affinity I [Source:MGI Symbol;Acc:MGI:95498] | 2.02 | 7.60E-03 |
| Tgfa | transforming growth factor alpha [Source:MGI Symbol;Acc:MGI:98724] | 1.20 | 7.67E-03 |
| Tpst2 | protein-tyrosine sulfotransferase 2 [Source:MGI Symbol;Acc:MGI:1309516] | 0.82 | 7.70E-03 |

| | | | |
|---|---|---|---|
| Atox1 | antioxidant 1 copper chaperone [Source:MGI Symbol;Acc:MGI:1333855] | 1.20 | 7.72E-03 |
| Cfp | complement factor properdin [Source:MGI Symbol;Acc:MGI:97545] | 1.44 | 7.74E-03 |
| Gm18979 | predicted gene, 18979 [Source:MGI Symbol;Acc:MGI:5011164] | 6.18 | 7.78E-03 |
| Gm14567 | predicted gene 14567 [Source:MGI Symbol;Acc:MGI:3705732] | 3.47 | 7.78E-03 |
| Gpr84 | G protein-coupled receptor 84 [Source:MGI Symbol;Acc:MGI:1934129] | 2.81 | 7.78E-03 |
| Slc39a1 | solute carrier family 39 (zinc transporter), member 1 [Source:MGI Symbol;Acc:MGI:1353474] | 1.19 | 7.78E-03 |
| Adamts1 | ADAM metallopeptidase with thrombospondin type 1 motif 1 [Source:MGI Symbol;Acc:MGI:109249] | 1.42 | 7.80E-03 |
| Entpd7 | ectonucleoside triphosphate diphosphohydrolase 7 [Source:MGI Symbol;Acc:MGI:2135885] | 1.07 | 7.95E-03 |
| Pmaip1 | phorbol-12-myristate-13-acetate-induced protein 1 [Source:MGI Symbol;Acc:MGI:1930146] | 1.60 | 8.00E-03 |
| Bfsp1 | beaded filament structural protein 1, in lens-CP94 [Source:MGI Symbol;Acc:MGI:101770] | 1.47 | 8.01E-03 |
| Fxyd5 | FXYD domain-containing ion transport regulator 5 [Source:MGI Symbol;Acc:MGI:1201785] | 2.25 | 8.02E-03 |
| Fes | feline sarcoma oncogene [Source:MGI Symbol;Acc:MGI:95514] | 1.64 | 8.02E-03 |
| Rpl38 | ribosomal protein L38 [Source:MGI Symbol;Acc:MGI:1914921] | 1.22 | 8.05E-03 |
| Tesmin | testis expressed metallothionein like [Source:MGI Symbol;Acc:MGI:1340029] | 2.00 | 8.17E-03 |
| F7 | coagulation factor VII [Source:MGI Symbol;Acc:MGI:109325] | 3.26 | 8.20E-03 |
| Creld2 | cysteine-rich with EGF-like domains 2 [Source:MGI Symbol;Acc:MGI:1923987] | 1.10 | 8.20E-03 |
| Pxdc1 | PX domain containing 1 [Source:MGI Symbol;Acc:MGI:1914145] | 1.44 | 8.23E-03 |
| Aplnr | apelin receptor [Source:MGI Symbol;Acc:MGI:1346086] | 2.20 | 8.26E-03 |
| Pde10a | phosphodiesterase 10A [Source:MGI Symbol;Acc:MGI:1345143] | 1.50 | 8.27E-03 |
| Gmfg | glia maturation factor, gamma [Source:MGI Symbol;Acc:MGI:1927135] | 2.03 | 8.30E-03 |
| Actb | actin, beta [Source:MGI Symbol;Acc:MGI:87904] | 0.85 | 8.33E-03 |
| Gas1 | growth arrest specific 1 [Source:MGI Symbol;Acc:MGI:95655] | 1.24 | 8.47E-03 |
| Gls2 | glutaminase 2 (liver, mitochondrial) [Source:MGI Symbol;Acc:MGI:2143539] | 1.39 | 8.49E-03 |
| Postn | periostin, osteoblast specific factor [Source:MGI Symbol;Acc:MGI:1926321] | 2.03 | 8.56E-03 |
| Oaz1-ps | ornithine decarboxylase antizyme 1, pseudogene [Source:MGI Symbol;Acc:MGI:108188] | 2.28 | 8.62E-03 |
| Slc41a2 | solute carrier family 41, member 2 [Source:MGI Symbol;Acc:MGI:2442940] | 1.83 | 8.62E-03 |
| Nes | nestin [Source:MGI Symbol;Acc:MGI:101784] | 1.06 | 8.75E-03 |
| Nhp2 | NHP2 ribonucleoprotein [Source:MGI Symbol;Acc:MGI:1098547] | 1.35 | 8.76E-03 |
| Ftl1 | ferritin light polypeptide 1 [Source:MGI Symbol;Acc:MGI:95589] | 1.20 | 8.77E-03 |
| Rdh10 | retinol dehydrogenase 10 (all-trans) [Source:MGI Symbol;Acc:MGI:1924238] | 1.25 | 8.78E-03 |
| P4hb | prolyl 4-hydroxylase, beta polypeptide [Source:MGI Symbol;Acc:MGI:97464] | 1.05 | 8.79E-03 |
| Gm18445 | predicted gene, 18445 [Source:MGI Symbol;Acc:MGI:5010630] | 2.91 | 8.79E-03 |
| Cald1 | caldesmon 1 [Source:MGI Symbol;Acc:MGI:88250] | 1.47 | 8.89E-03 |
| Eno2 | enolase 2, gamma neuronal [Source:MGI Symbol;Acc:MGI:95394] | 1.15 | 8.92E-03 |
| Gng8 | guanine nucleotide binding protein (G protein), gamma 8 [Source:MGI Symbol;Acc:MGI:109163] | 1.73 | 8.92E-03 |
| Sorcs2 | sortilin-related VPS10 domain containing receptor 2 [Source:MGI Symbol;Acc:MGI:1932289] | 1.81 | 8.98E-03 |
| Rps6ka4 | ribosomal protein S6 kinase, polypeptide 4 [Source:MGI Symbol;Acc:MGI:1930076] | 1.41 | 9.08E-03 |
| Prelid1 | PRELI domain containing 1 [Source:MGI Symbol;Acc:MGI:1913744] | 1.61 | 9.13E-03 |
| Zc3h12a | zinc finger CCCH type containing 12A [Source:MGI Symbol;Acc:MGI:2385891] | 1.31 | 9.17E-03 |
| Olfml2a | olfactomedin-like 2A [Source:MGI Symbol;Acc:MGI:2444741] | 1.33 | 9.17E-03 |
| Mafg | v-maf musculoaponeurotic fibrosarcoma oncogene family, protein G (avian) [Source:MGI Symbol;Acc:MGI:96911] | 0.93 | 9.17E-03 |
| Gja5 | gap junction protein, alpha 5 [Source:MGI Symbol;Acc:MGI:95716] | 1.84 | 9.18E-03 |
| Slc66a2 | solute carrier family 66 member 2 [Source:MGI Symbol;Acc:MGI:1914193] | 1.54 | 9.18E-03 |
| Uck2 | uridine-cytidine kinase 2 [Source:MGI Symbol;Acc:MGI:1931744] | 1.49 | 9.19E-03 |
| Glmp | glycosylated lysosomal membrane protein [Source:MGI Symbol;Acc:MGI:1913318] | 0.89 | 9.19E-03 |
| Mkx | mohawk homeobox [Source:MGI Symbol;Acc:MGI:2687286] | 1.28 | 9.20E-03 |
| Sdc3 | syndecan 3 [Source:MGI Symbol;Acc:MGI:1349163] | 1.27 | 9.20E-03 |
| Klf10 | Kruppel-like transcription factor 10 [Source:MGI Symbol;Acc:MGI:1101353] | 0.81 | 9.24E-03 |
| Crybb3 | crystallin, beta B3 [Source:MGI Symbol;Acc:MGI:102717] | 1.23 | 9.29E-03 |
| Cd53 | CD53 antigen [Source:MGI Symbol;Acc:MGI:88341] | 1.79 | 9.31E-03 |
| B4galt5 | UDP-Gal:betaGlcNAc beta 1,4-galactosyltransferase, polypeptide 5 [Source:MGI Symbol;Acc:MGI:1927169] | 1.51 | 9.35E-03 |
| Itgb3 | integrin beta 3 [Source:MGI Symbol;Acc:MGI:96612] | 1.88 | 9.41E-03 |
| Irx3 | Iroquois related homeobox 3 [Source:MGI Symbol;Acc:MGI:1197522] | 1.89 | 9.46E-03 |
| Itm2c | integral membrane protein 2C [Source:MGI Symbol;Acc:MGI:1927594] | 1.11 | 9.52E-03 |
| Map1a | microtubule-associated protein 1 A [Source:MGI Symbol;Acc:MGI:1306776] | 1.17 | 9.57E-03 |
| Samsn1 | SAM domain, SH3 domain and nuclear localization signals, 1 [Source:MGI Symbol;Acc:MGI:1914992] | 1.39 | 9.62E-03 |
| C920009B18Rik | RIKEN cDNA C920009B18 gene [Source:MGI Symbol;Acc:MGI:3583961] | 3.24 | 9.62E-03 |
| Rhof | ras homolog family member F (in filopodia) [Source:MGI Symbol;Acc:MGI:1345629] | 1.27 | 9.64E-03 |
| Fxyd4 | FXYD domain-containing ion transport regulator 4 [Source:MGI Symbol;Acc:MGI:1889005] | 1.81 | 9.65E-03 |
| Cxcr2 | C-X-C motif chemokine receptor 2 [Source:MGI Symbol;Acc:MGI:105303] | 1.37 | 9.74E-03 |
| Chst11 | carbohydrate sulfotransferase 11 [Source:MGI Symbol;Acc:MGI:1927166] | 2.01 | 9.75E-03 |
| Skil | SKI-like [Source:MGI Symbol;Acc:MGI:106203] | 1.04 | 9.76E-03 |
| Ostc | oligosaccharyltransferase complex subunit (non-catalytic) [Source:MGI Symbol;Acc:MGI:1913607] | 0.98 | 9.81E-03 |
| Tsku | tsukushi, small leucine rich proteoglycan [Source:MGI Symbol;Acc:MGI:2443855] | 1.49 | 9.98E-03 |
| Sirpd | signal regulatory protein delta [Source:MGI Symbol;Acc:MGI:3780136] | 4.66 | 1.00E-02 |
| Sgms2 | sphingomyelin synthase 2 [Source:MGI Symbol;Acc:MGI:1921692] | 1.34 | 1.01E-02 |
| Cyb561d2 | cytochrome b-561 domain containing 2 [Source:MGI Symbol;Acc:MGI:1929280] | 0.88 | 1.01E-02 |
| Arpc3 | actin related protein 2/3 complex, subunit 3 [Source:MGI Symbol;Acc:MGI:1928375] | 1.11 | 1.02E-02 |
| Rundc3a | RUN domain containing 3A [Source:MGI Symbol;Acc:MGI:1858752] | 1.65 | 1.02E-02 |
| Aplp1 | amyloid beta precursor like protein 1 [Source:MGI Symbol;Acc:MGI:88046] | 2.17 | 1.03E-02 |

| | | | |
|---|---|---|---|
| Slc16a3 | solute carrier family 16 (monocarboxylic acid transporters), member 3 [Source:MGI Symbol;Acc:MGI:1933438] | 2.17 | 1.04E-02 |
| Syce2 | synaptonemal complex central element protein 2 [Source:MGI Symbol;Acc:MGI:1919096] | 1.20 | 1.04E-02 |
| Dpagt1 | dolichyl-phosphate N-acetylglucosaminephosphotransferase 1 [Source:MGI Symbol;Acc:MGI:1196396] | 0.74 | 1.04E-02 |
| Rpl12 | ribosomal protein L12 [Source:MGI Symbol;Acc:MGI:98002] | 1.21 | 1.04E-02 |
| Zmiz1 | zinc finger, MIZ-type containing 1 [Source:MGI Symbol;Acc:MGI:3040693] | 0.59 | 1.04E-02 |
| Adcy7 | adenylate cyclase 7 [Source:MGI Symbol;Acc:MGI:102891] | 0.89 | 1.05E-02 |
| Pelo | pelota mRNA surveillance and ribosome rescue factor [Source:MGI Symbol;Acc:MGI:2145154] | 1.08 | 1.06E-02 |
| Cd44 | CD44 antigen [Source:MGI Symbol;Acc:MGI:88338] | 1.36 | 1.06E-02 |
| Zfp92 | zinc finger protein 92 [Source:MGI Symbol;Acc:MGI:108094] | 0.70 | 1.07E-02 |
| Nsmce1 | NSE1 homolog, SMC5-SMC6 complex component [Source:MGI Symbol;Acc:MGI:1914961] | 0.93 | 1.07E-02 |
| Adgra2 | adhesion G protein-coupled receptor A2 [Source:MGI Symbol;Acc:MGI:1925810] | 1.53 | 1.07E-02 |
| Scarf1 | scavenger receptor class F, member 1 [Source:MGI Symbol;Acc:MGI:2449455] | 1.53 | 1.07E-02 |
| Hc | hemolytic complement [Source:MGI Symbol;Acc:MGI:96031] | 2.99 | 1.08E-02 |
| Gml | glycosylphosphatidylinositol anchored molecule like [Source:MGI Symbol;Acc:MGI:3644767] | 5.77 | 1.08E-02 |
| Serpinh1 | serine (or cysteine) peptidase inhibitor, clade H, member 1 [Source:MGI Symbol;Acc:MGI:88283] | 1.51 | 1.08E-02 |
| Sh3bgrl3 | SH3 domain binding glutamic acid-rich protein-like 3 [Source:MGI Symbol;Acc:MGI:1920973] | 0.89 | 1.09E-02 |
| Ddost | dolichyl-di-phosphooligosaccharide-protein glycotransferase [Source:MGI Symbol;Acc:MGI:1194508] | 0.71 | 1.09E-02 |
| Gsta13 | glutathione S-transferase alpha 13 [Source:MGI Symbol;Acc:MGI:3826440] | 3.65 | 1.09E-02 |
| Sting1 | stimulator of interferon response cGAMP interactor 1 [Source:MGI Symbol;Acc:MGI:1919762] | 2.07 | 1.09E-02 |
| St6galnac4 | ST6 (alpha-N-acetyl-neuraminyl-2,3-beta-galactosyl-1,3)-N-acetylgalactosaminide alpha-2,6-sialyltransferase 4 [Source:MGI Symbol;Acc:MGI:1341894] | 1.51 | 1.09E-02 |
| Txndc5 | thioredoxin domain containing 5 [Source:MGI Symbol;Acc:MGI:2145316] | 1.23 | 1.09E-02 |
| Dnaaf5 | dynein, axonemal assembly factor 5 [Source:MGI Symbol;Acc:MGI:3616079] | 0.79 | 1.10E-02 |
| Mup16 | major urinary protein 16 [Source:MGI Symbol;Acc:MGI:3780250] | 5.83 | 1.10E-02 |
| 9130401M01Rik | RIKEN cDNA 9130401M01 gene [Source:MGI Symbol;Acc:MGI:1923008] | 0.97 | 1.10E-02 |
| Mif | macrophage migration inhibitory factor (glycosylation-inhibiting factor) [Source:MGI Symbol;Acc:MGI:96982] | 0.82 | 1.10E-02 |
| Pgd | phosphogluconate dehydrogenase [Source:MGI Symbol;Acc:MGI:97553] | 0.82 | 1.10E-02 |
| Fabp12 | fatty acid binding protein 12 [Source:MGI Symbol;Acc:MGI:1922747] | 2.44 | 1.11E-02 |
| Eif5a | eukaryotic translation initiation factor 5A [Source:MGI Symbol;Acc:MGI:106248] | 0.98 | 1.11E-02 |
| Tpsb2 | tryptase beta 2 [Source:MGI Symbol;Acc:MGI:96942] | 1.66 | 1.12E-02 |
| Rcn1 | reticulocalbin 1 [Source:MGI Symbol;Acc:MGI:104559] | 0.98 | 1.12E-02 |
| Rbm47 | RNA binding motif protein 47 [Source:MGI Symbol;Acc:MGI:2384294] | 0.83 | 1.12E-02 |
| Peds1 | plasmanylethanolamine desaturase 1 [Source:MGI Symbol;Acc:MGI:2142624] | 0.90 | 1.12E-02 |
| Begain | brain-enriched guanylate kinase-associated [Source:MGI Symbol;Acc:MGI:3044626] | 1.61 | 1.13E-02 |
| Anxa2r2 | annexin A2 receptor 2 [Source:MGI Symbol;Acc:MGI:5595238] | 3.40 | 1.14E-02 |
| Rgs1 | regulator of G-protein signaling 1 [Source:MGI Symbol;Acc:MGI:1354694] | 1.46 | 1.15E-02 |
| Lrrc25 | leucine rich repeat containing 25 [Source:MGI Symbol;Acc:MGI:2445284] | 1.79 | 1.15E-02 |
| Ctla2a | cytotoxic T lymphocyte-associated protein 2 alpha [Source:MGI Symbol;Acc:MGI:88554] | 1.56 | 1.15E-02 |
| Pla1a | phospholipase A1 member A [Source:MGI Symbol;Acc:MGI:1934677] | 1.92 | 1.15E-02 |
| Oas1a | 2'-5' oligoadenylate synthetase 1A [Source:MGI Symbol;Acc:MGI:2180860] | 1.85 | 1.15E-02 |
| Tnfrsf23 | tumor necrosis factor receptor superfamily, member 23 [Source:MGI Symbol;Acc:MGI:1930269] | 1.41 | 1.16E-02 |
| Surf4 | surfeit gene 4 [Source:MGI Symbol;Acc:MGI:98445] | 0.97 | 1.16E-02 |
| Tmed9 | transmembrane p24 trafficking protein 9 [Source:MGI Symbol;Acc:MGI:1914761] | 0.70 | 1.16E-02 |
| Gadl1 | glutamate decarboxylase-like 1 [Source:MGI Symbol;Acc:MGI:1920998] | 2.99 | 1.17E-02 |
| Adamts9 | ADAM metallopeptidase with thrombospondin type 1 motif 9 [Source:MGI Symbol;Acc:MGI:1916320] | 1.34 | 1.18E-02 |
| Or2aa1 | olfactory receptor family 2 subfamily AA member 1 [Source:MGI Symbol;Acc:MGI:3030057] | 6.15 | 1.19E-02 |
| Prkcsh | protein kinase C substrate 80K-H [Source:MGI Symbol;Acc:MGI:107877] | 0.92 | 1.19E-02 |
| Hyou1 | hypoxia up-regulated 1 [Source:MGI Symbol;Acc:MGI:108030] | 0.85 | 1.19E-02 |
| Nat8 | N-acetyltransferase 8 (GCN5-related) [Source:MGI Symbol;Acc:MGI:1915646] | 2.70 | 1.20E-02 |
| Renbp | renin binding protein [Source:MGI Symbol;Acc:MGI:105940] | 1.03 | 1.20E-02 |
| Fcgr2b | Fc receptor, IgG, low affinity IIb [Source:MGI Symbol;Acc:MGI:95499] | 1.77 | 1.21E-02 |
| Ap2s1 | adaptor-related protein complex 2, sigma 1 subunit [Source:MGI Symbol;Acc:MGI:2141861] | 1.18 | 1.23E-02 |
| Nr4a3 | nuclear receptor subfamily 4, group A, member 3 [Source:MGI Symbol;Acc:MGI:1352457] | 1.25 | 1.23E-02 |
| Midn | midnolin [Source:MGI Symbol;Acc:MGI:1890222] | 1.18 | 1.24E-02 |
| Fa2h | fatty acid 2-hydroxylase [Source:MGI Symbol;Acc:MGI:2443327] | 0.83 | 1.24E-02 |
| Defb4 | defensin beta 4 [Source:MGI Symbol;Acc:MGI:1927667] | 3.91 | 1.24E-02 |
| Elob | elongin B [Source:MGI Symbol;Acc:MGI:1914923] | 0.89 | 1.24E-02 |
| Il36a | interleukin 36A [Source:MGI Symbol;Acc:MGI:1859324] | 2.08 | 1.25E-02 |
| Degs1 | delta 4-desaturase, sphingolipid 1 [Source:MGI Symbol;Acc:MGI:1097711] | 0.89 | 1.25E-02 |
| Vcl | vinculin [Source:MGI Symbol;Acc:MGI:98927] | 0.76 | 1.26E-02 |
| Mrpl20 | mitochondrial ribosomal protein L20 [Source:MGI Symbol;Acc:MGI:2137221] | 1.06 | 1.27E-02 |
| Pld3 | phospholipase D family member 3 [Source:MGI Symbol;Acc:MGI:1333782] | 0.83 | 1.27E-02 |
| Mxra8 | matrix-remodelling associated 8 [Source:MGI Symbol;Acc:MGI:1922011] | 1.77 | 1.28E-02 |
| Sys1 | SYS1 Golgi-localized integral membrane protein homolog (S. cerevisiae) [Source:MGI Symbol;Acc:MGI:1913710] | 0.77 | 1.28E-02 |
| Ptgfrn | prostaglandin F2 receptor negative regulator [Source:MGI Symbol;Acc:MGI:1277114] | 0.89 | 1.28E-02 |
| Lrat | lecithin-retinol acyltransferase (phosphatidylcholine-retinol-O-acyltransferase) [Source:MGI Symbol;Acc:MGI:1891259] | 2.27 | 1.28E-02 |
| Slc39a14 | solute carrier family 39 (zinc transporter), member 14 [Source:MGI Symbol;Acc:MGI:2384851] | 2.13 | 1.28E-02 |
| Chsy1 | chondroitin sulfate synthase 1 [Source:MGI Symbol;Acc:MGI:2681120] | 1.79 | 1.28E-02 |

| | | | |
|---|---|---|---|
| Ppic | peptidylprolyl isomerase C [Source:MGI Symbol;Acc:MGI:97751] | 1.65 | 1.28E-02 |
| AB124611 | cDNA sequence AB124611 [Source:MGI Symbol;Acc:MGI:3043001] | 1.72 | 1.29E-02 |
| Tssc4 | tumor-suppressing subchromosomal transferable fragment 4 [Source:MGI Symbol;Acc:MGI:1861712] | 1.39 | 1.30E-02 |
| Znhit2 | zinc finger, HIT domain containing 2 [Source:MGI Symbol;Acc:MGI:1352481] | 1.56 | 1.30E-02 |
| Nfatc4 | nuclear factor of activated T cells, cytoplasmic, calcineurin dependent 4 [Source:MGI Symbol;Acc:MGI:1920431] | 1.04 | 1.30E-02 |
| Amn | amnionless [Source:MGI Symbol;Acc:MGI:1934943] | 2.61 | 1.30E-02 |
| Atp5mj | ATP synthase membrane subunit j [Source:MGI Symbol;Acc:MGI:1917507] | 0.90 | 1.30E-02 |
| Clstn2 | calsyntenin 2 [Source:MGI Symbol;Acc:MGI:1929897] | 2.62 | 1.31E-02 |
| Rhob | ras homolog family member B [Source:MGI Symbol;Acc:MGI:107949] | 1.07 | 1.31E-02 |
| Nipa2 | non imprinted in Prader-Willi/Angelman syndrome 2 homolog (human) [Source:MGI Symbol;Acc:MGI:1913918] | 0.72 | 1.32E-02 |
| Rpl7a | ribosomal protein L7A [Source:MGI Symbol;Acc:MGI:1353472] | 0.97 | 1.32E-02 |
| Rab20 | RAB20, member RAS oncogene family [Source:MGI Symbol;Acc:MGI:102789] | 1.59 | 1.32E-02 |
| Lrfn4 | leucine rich repeat and fibronectin type III domain containing 4 [Source:MGI Symbol;Acc:MGI:2385612] | 1.24 | 1.32E-02 |
| Gtf2f1 | general transcription factor IIF, polypeptide 1 [Source:MGI Symbol;Acc:MGI:1923848] | 0.73 | 1.33E-02 |
| Olfml3 | olfactomedin-like 3 [Source:MGI Symbol;Acc:MGI:1914877] | 1.75 | 1.34E-02 |
| Msn | moesin [Source:MGI Symbol;Acc:MGI:97167] | 1.34 | 1.34E-02 |
| Stc1 | stanniocalcin 1 [Source:MGI Symbol;Acc:MGI:109131] | 2.43 | 1.34E-02 |
| Bzw1 | basic leucine zipper and W2 domains 1 [Source:MGI Symbol;Acc:MGI:1914132] | 0.77 | 1.34E-02 |
| Meox1 | mesenchyme homeobox 1 [Source:MGI Symbol;Acc:MGI:103220] | 2.24 | 1.34E-02 |
| Tmem121 | transmembrane protein 121 [Source:MGI Symbol;Acc:MGI:1916445] | 2.48 | 1.36E-02 |
| Lce1e | late cornified envelope 1E [Source:MGI Symbol;Acc:MGI:1915944] | 2.79 | 1.36E-02 |
| Tmem132e | transmembrane protein 132E [Source:MGI Symbol;Acc:MGI:2685490] | 2.20 | 1.37E-02 |
| Grap | GRB2-related adaptor protein [Source:MGI Symbol;Acc:MGI:1918770] | 1.29 | 1.37E-02 |
| Cad | carbamoyl-phosphate synthetase 2, aspartate transcarbamylase, and dihydroorotase [Source:MGI Symbol;Acc:MGI:1916969] | 1.43 | 1.38E-02 |
| Maff | v-maf musculoaponeurotic fibrosarcoma oncogene family, protein F (avian) [Source:MGI Symbol;Acc:MGI:96910] | 1.19 | 1.38E-02 |
| Sfrp2 | secreted frizzled-related protein 2 [Source:MGI Symbol;Acc:MGI:108078] | 2.22 | 1.38E-02 |
| Mrpl36 | mitochondrial ribosomal protein L36 [Source:MGI Symbol;Acc:MGI:2137228] | 0.75 | 1.38E-02 |
| Ptchd1 | patched domain containing 1 [Source:MGI Symbol;Acc:MGI:2685233] | 2.79 | 1.38E-02 |
| Gpx2 | glutathione peroxidase 2 [Source:MGI Symbol;Acc:MGI:106609] | 1.74 | 1.38E-02 |
| B230207O21Rik | RIKEN cDNA B230207O21 gene [Source:MGI Symbol;Acc:MGI:3704213] | 5.70 | 1.39E-02 |
| Arl11 | ADP-ribosylation factor-like 11 [Source:MGI Symbol;Acc:MGI:2444054] | 1.37 | 1.39E-02 |
| Col14a1 | collagen, type XIV, alpha 1 [Source:MGI Symbol;Acc:MGI:1341272] | 1.28 | 1.39E-02 |
| Sgsm1 | small G protein signaling modulator 1 [Source:MGI Symbol;Acc:MGI:107320] | 0.92 | 1.39E-02 |
| Gad1-ps | glutamate decarboxylase 1, pseudogene [Source:MGI Symbol;Acc:MGI:95633] | 2.58 | 1.40E-02 |
| Ube2s | ubiquitin-conjugating enzyme E2S [Source:MGI Symbol;Acc:MGI:1925141] | 1.45 | 1.42E-02 |
| Pgs1 | phosphatidylglycerophosphate synthase 1 [Source:MGI Symbol;Acc:MGI:1921701] | 1.00 | 1.42E-02 |
| Tmem125 | transmembrane protein 125 [Source:MGI Symbol;Acc:MGI:1923409] | 1.32 | 1.42E-02 |
| Fbxl15 | F-box and leucine-rich repeat protein 15 [Source:MGI Symbol;Acc:MGI:1915681] | 2.16 | 1.43E-02 |
| D8Ertd738e | DNA segment, Chr 8, ERATO Doi 738, expressed [Source:MGI Symbol;Acc:MGI:1289231] | 1.41 | 1.43E-02 |
| Ccdc12 | coiled-coil domain containing 12 [Source:MGI Symbol;Acc:MGI:1919904] | 1.02 | 1.43E-02 |
| Bcl2l11 | BCL2 like 11 [Source:MGI Symbol;Acc:MGI:1197519] | 1.15 | 1.43E-02 |
| Tlcd2 | TLC domain containing 2 [Source:MGI Symbol;Acc:MGI:1917141] | 1.05 | 1.44E-02 |
| Atosb | atos homolog B [Source:MGI Symbol;Acc:MGI:2441854] | 0.83 | 1.48E-02 |
| Akt1 | thymoma viral proto-oncogene 1 [Source:MGI Symbol;Acc:MGI:87986] | 0.90 | 1.48E-02 |
| Pira2 | paired-Ig-like receptor A2 [Source:MGI Symbol;Acc:MGI:1195970] | 2.24 | 1.48E-02 |
| Tubb2a | tubulin, beta 2A class IIA [Source:MGI Symbol;Acc:MGI:107861] | 1.64 | 1.48E-02 |
| Nubp2 | nucleotide binding protein 2 [Source:MGI Symbol;Acc:MGI:1347072] | 0.98 | 1.48E-02 |
| Tmem165 | transmembrane protein 165 [Source:MGI Symbol;Acc:MGI:894407] | 0.97 | 1.48E-02 |
| Scpep1 | serine carboxypeptidase 1 [Source:MGI Symbol;Acc:MGI:1921867] | 0.76 | 1.48E-02 |
| Dpysl4 | dihydropyrimidinase-like 4 [Source:MGI Symbol;Acc:MGI:1349764] | 1.45 | 1.48E-02 |
| Gm18787 | predicted gene, 18787 [Source:MGI Symbol;Acc:MGI:5010972] | 2.76 | 1.49E-02 |
| Zfp593 | zinc finger protein 593 [Source:MGI Symbol;Acc:MGI:1915290] | 1.46 | 1.49E-02 |
| Nfil3 | nuclear factor, interleukin 3, regulated [Source:MGI Symbol;Acc:MGI:109495] | 1.56 | 1.52E-02 |
| Akr1c18 | aldo-keto reductase family 1, member C18 [Source:MGI Symbol;Acc:MGI:2145420] | 2.02 | 1.53E-02 |
| Ptges3 | prostaglandin E synthase 3 [Source:MGI Symbol;Acc:MGI:1929282] | 0.67 | 1.53E-02 |
| B4galnt1 | beta-1,4-N-acetyl-galactosaminyl transferase 1 [Source:MGI Symbol;Acc:MGI:1342057] | 1.39 | 1.54E-02 |
| Csf2rb2 | colony stimulating factor 2 receptor, beta 2, low-affinity (granulocyte-macrophage) [Source:MGI Symbol;Acc:MGI:1339760] | 1.47 | 1.54E-02 |
| Entpd1 | ectonucleoside triphosphate diphosphohydrolase 1 [Source:MGI Symbol;Acc:MGI:102805] | 1.45 | 1.57E-02 |
| Pygo2 | pygopus 2 [Source:MGI Symbol;Acc:MGI:1916161] | 1.35 | 1.57E-02 |
| Pf4 | platelet factor 4 [Source:MGI Symbol;Acc:MGI:1888711] | 1.48 | 1.57E-02 |
| Grem2 | gremlin 2, DAN family BMP antagonist [Source:MGI Symbol;Acc:MGI:1344367] | 2.12 | 1.58E-02 |
| Lamc2 | laminin, gamma 2 [Source:MGI Symbol;Acc:MGI:99913] | 1.18 | 1.58E-02 |
| Tlx1 | T cell leukemia, homeobox 1 [Source:MGI Symbol;Acc:MGI:98769] | 5.62 | 1.58E-02 |
| Rps24 | ribosomal protein S24 [Source:MGI Symbol;Acc:MGI:98147] | 0.67 | 1.58E-02 |
| Ggn | gametogenetin [Source:MGI Symbol;Acc:MGI:2181461] | 1.03 | 1.58E-02 |
| Atp6ap2 | ATPase, H+ transporting, lysosomal accessory protein 2 [Source:MGI Symbol;Acc:MGI:1917745] | 0.65 | 1.58E-02 |
| Barhl1 | BarH like homeobox 1 [Source:MGI Symbol;Acc:MGI:1859288] | 1.53 | 1.58E-02 |

| | | | |
|---|---|---|---|
| Kdm6b | KDM1 lysine (K)-specific demethylase 6B [Source:MGI Symbol;Acc:MGI:2448492] | 0.95 | 1.58E-02 |
| Tfdp1 | transcription factor Dp 1 [Source:MGI Symbol;Acc:MGI:101934] | 1.00 | 1.60E-02 |
| Ncf2 | neutrophil cytosolic factor 2 [Source:MGI Symbol;Acc:MGI:97284] | 1.51 | 1.60E-02 |
| Yy2 | Yy2 transcription factor [Source:MGI Symbol;Acc:MGI:3837947] | 4.00 | 1.61E-02 |
| Ehd3 | EH-domain containing 3 [Source:MGI Symbol;Acc:MGI:1928900] | 1.85 | 1.62E-02 |
| Cmtm3 | CKLF-like MARVEL transmembrane domain containing 3 [Source:MGI Symbol;Acc:MGI:2447162] | 1.46 | 1.62E-02 |
| C1qtnf2 | C1q and tumor necrosis factor related protein 2 [Source:MGI Symbol;Acc:MGI:1916433] | 1.74 | 1.64E-02 |
| Pcdhb9 | protocadherin beta 9 [Source:MGI Symbol;Acc:MGI:2136744] | 2.11 | 1.64E-02 |
| Itga2b | integrin alpha 2b [Source:MGI Symbol;Acc:MGI:96601] | 0.93 | 1.64E-02 |
| Gm11425 | predicted gene 11425 [Source:MGI Symbol;Acc:MGI:3650957] | 1.94 | 1.65E-02 |
| Pik3r5 | phosphoinositide-3-kinase regulatory subunit 5 [Source:MGI Symbol;Acc:MGI:2443588] | 1.48 | 1.66E-02 |
| Agap2 | ArfGAP with GTPase domain, ankyrin repeat and PH domain 2 [Source:MGI Symbol;Acc:MGI:3580016] | 1.14 | 1.66E-02 |
| Gm12844 | predicted gene 12844 [Source:MGI Symbol;Acc:MGI:3651053] | 5.83 | 1.67E-02 |
| Anapc15 | anaphase promoting complex C subunit 15 [Source:MGI Symbol;Acc:MGI:1922680] | 0.90 | 1.67E-02 |
| Ngp | neutrophilic granule protein [Source:MGI Symbol;Acc:MGI:105983] | 3.04 | 1.68E-02 |
| Prss35 | serine protease 35 [Source:MGI Symbol;Acc:MGI:2444800] | 1.92 | 1.68E-02 |
| Hic1 | hypermethylated in cancer 1 [Source:MGI Symbol;Acc:MGI:1338010] | 1.42 | 1.69E-02 |
| Plcg2 | phospholipase C, gamma 2 [Source:MGI Symbol;Acc:MGI:97616] | 0.87 | 1.71E-02 |
| Efhd2 | EF hand domain containing 2 [Source:MGI Symbol;Acc:MGI:106504] | 1.14 | 1.71E-02 |
| Grina | glutamate receptor, ionotropic, N-methyl D-aspartate-associated protein 1 (glutamate binding) [Source:MGI Symbol;Acc:MGI:1913418] | 0.90 | 1.71E-02 |
| Atp1a3 | ATPase, Na+/K+ transporting, alpha 3 polypeptide [Source:MGI Symbol;Acc:MGI:88107] | 1.42 | 1.72E-02 |
| Pop7 | processing of precursor 7, ribonuclease P family, (S. cerevisiae) [Source:MGI Symbol;Acc:MGI:1921347] | 1.19 | 1.72E-02 |
| Maged1 | MAGE family member D1 [Source:MGI Symbol;Acc:MGI:1930187] | 1.20 | 1.72E-02 |
| Creb3l1 | cAMP responsive element binding protein 3-like 1 [Source:MGI Symbol;Acc:MGI:1347062] | 1.34 | 1.73E-02 |
| Nradd | neurotrophin receptor associated death domain [Source:MGI Symbol;Acc:MGI:1914419] | 1.11 | 1.74E-02 |
| Nme1 | NME/NM23 nucleoside diphosphate kinase 1 [Source:MGI Symbol;Acc:MGI:97355] | 1.42 | 1.74E-02 |
| Trim30a | tripartite motif-containing 30A [Source:MGI Symbol;Acc:MGI:98178] | 1.43 | 1.75E-02 |
| Tbc1d9 | TBC1 domain family, member 9 [Source:MGI Symbol;Acc:MGI:1918560] | 0.88 | 1.75E-02 |
| Map3k6 | mitogen-activated protein kinase kinase kinase 6 [Source:MGI Symbol;Acc:MGI:1855691] | 1.51 | 1.75E-02 |
| Psmb4 | proteasome (prosome, macropain) subunit, beta type 4 [Source:MGI Symbol;Acc:MGI:1098257] | 0.68 | 1.75E-02 |
| Gm21863 | predicted gene, 21863 [Source:MGI Symbol;Acc:MGI:5434027] | 2.65 | 1.77E-02 |
| Sprr1b | small proline-rich protein 1B [Source:MGI Symbol;Acc:MGI:106659] | 7.62 | 1.78E-02 |
| Gm11839 | predicted gene 11839 [Source:MGI Symbol;Acc:MGI:3651162] | 1.21 | 1.78E-02 |
| Sub1 | SUB1 homolog, transcriptional regulator [Source:MGI Symbol;Acc:MGI:104811] | 0.65 | 1.78E-02 |
| Cd5 | CD5 antigen [Source:MGI Symbol;Acc:MGI:88340] | 0.80 | 1.79E-02 |
| Pcdhgc4 | protocadherin gamma subfamily C, 4 [Source:MGI Symbol;Acc:MGI:1935203] | 0.92 | 1.79E-02 |
| Nol12 | nucleolar protein 12 [Source:MGI Symbol;Acc:MGI:2146285] | 0.85 | 1.79E-02 |
| Unc93b1 | unc-93 homolog B1, TLR signaling regulator [Source:MGI Symbol;Acc:MGI:1859307] | 1.33 | 1.80E-02 |
| Scamp4 | secretory carrier membrane protein 4 [Source:MGI Symbol;Acc:MGI:1928947] | 1.15 | 1.80E-02 |
| Myof | myoferlin [Source:MGI Symbol;Acc:MGI:1919192] | 0.75 | 1.80E-02 |
| Runx1 | runt related transcription factor 1 [Source:MGI Symbol;Acc:MGI:99852] | 1.66 | 1.81E-02 |
| Ccnd3 | cyclin D3 [Source:MGI Symbol;Acc:MGI:88315] | 1.05 | 1.81E-02 |
| Nop56 | NOP56 ribonucleoprotein [Source:MGI Symbol;Acc:MGI:1914384] | 0.97 | 1.81E-02 |
| Ap1s1 | adaptor protein complex AP-1, sigma 1 [Source:MGI Symbol;Acc:MGI:1098244] | 0.90 | 1.81E-02 |
| Tspan14 | tetraspanin 14 [Source:MGI Symbol;Acc:MGI:1196325] | 0.71 | 1.81E-02 |
| Scamp5 | secretory carrier membrane protein 5 [Source:MGI Symbol;Acc:MGI:1928948] | 1.38 | 1.82E-02 |
| Atp6v0b | ATPase, H+ transporting, lysosomal V0 subunit B [Source:MGI Symbol;Acc:MGI:1890510] | 1.01 | 1.82E-02 |
| Impg2 | interphotoreceptor matrix proteoglycan 2 [Source:MGI Symbol;Acc:MGI:3044955] | 0.77 | 1.83E-02 |
| Traf1 | TNF receptor-associated factor 1 [Source:MGI Symbol;Acc:MGI:101836] | 1.26 | 1.83E-02 |
| Unc93a | unc-93 homolog A [Source:MGI Symbol;Acc:MGI:1933250] | 1.23 | 1.83E-02 |
| Bysl | bystin-like [Source:MGI Symbol;Acc:MGI:1858419] | 0.93 | 1.83E-02 |
| Col11a2 | collagen, type XI, alpha 2 [Source:MGI Symbol;Acc:MGI:88447] | 1.80 | 1.83E-02 |
| Prex1 | phosphatidylinositol-3,4,5-trisphosphate-dependent Rac exchange factor 1 [Source:MGI Symbol;Acc:MGI:3040696] | 1.04 | 1.83E-02 |
| Cdhr17 | cadherin related family member 17 [Source:MGI Symbol;Acc:MGI:5579416] | 4.94 | 1.83E-02 |
| Itgb8 | integrin beta 8 [Source:MGI Symbol;Acc:MGI:1338035] | 0.59 | 1.84E-02 |
| Phlda2 | pleckstrin homology like domain, family A, member 2 [Source:MGI Symbol;Acc:MGI:1202307] | 2.04 | 1.84E-02 |
| Slc39a7 | solute carrier family 39 (zinc transporter), member 7 [Source:MGI Symbol;Acc:MGI:95909] | 0.93 | 1.84E-02 |
| Lce1f | late cornified envelope 1F [Source:MGI Symbol;Acc:MGI:1915078] | 2.83 | 1.85E-02 |
| Ifi30 | interferon gamma inducible protein 30 [Source:MGI Symbol;Acc:MGI:2137648] | 1.02 | 1.88E-02 |
| Ndufc2 | NADH:ubiquinone oxidoreductase subunit C2 [Source:MGI Symbol;Acc:MGI:1344370] | 0.83 | 1.88E-02 |
| Ncln | nicalin [Source:MGI Symbol;Acc:MGI:1926081] | 0.75 | 1.88E-02 |
| Hint1 | histidine triad nucleotide binding protein 1 [Source:MGI Symbol;Acc:MGI:1321133] | 1.04 | 1.89E-02 |
| Grn | granulin [Source:MGI Symbol;Acc:MGI:95832] | 1.04 | 1.89E-02 |
| Tgfb1 | transforming growth factor, beta 1 [Source:MGI Symbol;Acc:MGI:98725] | 1.35 | 1.91E-02 |
| 1700017B05Rik | RIKEN cDNA 1700017B05 gene [Source:MGI Symbol;Acc:MGI:1921461] | 1.31 | 1.91E-02 |
| Pisd | phosphatidylserine decarboxylase [Source:MGI Symbol;Acc:MGI:2445114] | 0.70 | 1.91E-02 |
| Rab5c | RAB5C, member RAS oncogene family [Source:MGI Symbol;Acc:MGI:105306] | 0.82 | 1.91E-02 |
| Lrrc59 | leucine rich repeat containing 59 [Source:MGI Symbol;Acc:MGI:2138133] | 1.08 | 1.91E-02 |
| Lmna | lamin A [Source:MGI Symbol;Acc:MGI:96794] | 0.94 | 1.91E-02 |
| Il17a | interleukin 17A [Source:MGI Symbol;Acc:MGI:107364] | 6.11 | 1.91E-02 |

| | | | |
|---|---|---|---|
| Relt | RELT tumor necrosis factor receptor [Source:MGI Symbol;Acc:MGI:2443373] | 1.39 | 1.92E-02 |
| Pabpc1l | poly(A) binding protein, cytoplasmic 1-like [Source:MGI Symbol;Acc:MGI:1922908] | 1.24 | 1.92E-02 |
| Polr2f | polymerase (RNA) II (DNA directed) polypeptide F [Source:MGI Symbol;Acc:MGI:1349393] | 1.13 | 1.92E-02 |
| Aadacl3 | arylacetamide deacetylase like 3 [Source:MGI Symbol;Acc:MGI:2685281] | 0.77 | 1.92E-02 |
| Bola2 | bolA family member 2 [Source:MGI Symbol;Acc:MGI:1913412] | 1.16 | 1.92E-02 |
| Slc27a4 | solute carrier family 27 (fatty acid transporter), member 4 [Source:MGI Symbol;Acc:MGI:1347347] | 0.95 | 1.92E-02 |
| Tgm2 | transglutaminase 2, C polypeptide [Source:MGI Symbol;Acc:MGI:98731] | 1.60 | 1.92E-02 |
| Gm6377 | predicted gene 6377 [Source:MGI Symbol;Acc:MGI:3647255] | 1.67 | 1.93E-02 |
| H2-M9 | histocompatibility 2, M region locus 9 [Source:MGI Symbol;Acc:MGI:1276570] | 1.89 | 1.93E-02 |
| Saa2 | serum amyloid A 2 [Source:MGI Symbol;Acc:MGI:98222] | 5.14 | 1.93E-02 |
| Cilp | cartilage intermediate layer protein, nucleotide pyrophosphohydrolase [Source:MGI Symbol;Acc:MGI:2444507] | 1.52 | 1.94E-02 |
| Cacna1g | calcium channel, voltage-dependent, T type, alpha 1G subunit [Source:MGI Symbol;Acc:MGI:1201678] | 1.39 | 1.95E-02 |
| Csta1 | cystatin A1 [Source:MGI Symbol;Acc:MGI:3524930] | 2.39 | 1.95E-02 |
| Rpl36 | ribosomal protein L36 [Source:MGI Symbol;Acc:MGI:1860603] | 0.96 | 1.95E-02 |
| Gm7666 | predicted pseudogene 7666 [Source:MGI Symbol;Acc:MGI:3648907] | 4.21 | 1.95E-02 |
| Cpn1 | carboxypeptidase N, polypeptide 1 [Source:MGI Symbol;Acc:MGI:2135874] | 3.36 | 1.95E-02 |
| Tbx6 | T-box 6 [Source:MGI Symbol;Acc:MGI:102539] | 1.03 | 1.95E-02 |
| Trpv2 | transient receptor potential cation channel, subfamily V, member 2 [Source:MGI Symbol;Acc:MGI:1341836] | 1.37 | 1.95E-02 |
| Col28a1 | collagen, type XXVIII, alpha 1 [Source:MGI Symbol;Acc:MGI:2685312] | 1.80 | 1.96E-02 |
| Tmem176a | transmembrane protein 176A [Source:MGI Symbol;Acc:MGI:1913308] | 1.23 | 1.96E-02 |
| Adam19 | ADAM metallopeptidase domain 19 [Source:MGI Symbol;Acc:MGI:105377] | 1.28 | 1.97E-02 |
| Rab5if | RAB5 interacting factor [Source:MGI Symbol;Acc:MGI:1914638] | 0.73 | 1.97E-02 |
| Fcer1g | Fc receptor, IgE, high affinity I, gamma polypeptide [Source:MGI Symbol;Acc:MGI:95496] | 1.67 | 2.01E-02 |
| Azin2 | antizyme inhibitor 2 [Source:MGI Symbol;Acc:MGI:2442093] | 1.24 | 2.01E-02 |
| Ndufb7 | NADH:ubiquinone oxidoreductase subunit B7 [Source:MGI Symbol;Acc:MGI:1914166] | 1.07 | 2.01E-02 |
| Vasp | vasodilator-stimulated phosphoprotein [Source:MGI Symbol;Acc:MGI:109268] | 1.03 | 2.01E-02 |
| Gch1 | GTP cyclohydrolase 1 [Source:MGI Symbol;Acc:MGI:95675] | 1.22 | 2.02E-02 |
| Sp140l1 | Sp140 nuclear body protein like 1 [Source:MGI Symbol;Acc:MGI:3037746] | 2.48 | 2.02E-02 |
| Ubd | ubiquitin D [Source:MGI Symbol;Acc:MGI:1344410] | 3.76 | 2.02E-02 |
| Gpr35 | G protein-coupled receptor 35 [Source:MGI Symbol;Acc:MGI:1929509] | 1.09 | 2.02E-02 |
| H2ac8 | H2A clustered histone 8 [Source:MGI Symbol;Acc:MGI:2448290] | 1.56 | 2.03E-02 |
| Spata20 | spermatogenesis associated 20 [Source:MGI Symbol;Acc:MGI:2183449] | 3.03 | 2.04E-02 |
| Sirpa | signal-regulatory protein alpha [Source:MGI Symbol;Acc:MGI:108563] | 1.23 | 2.04E-02 |
| Rbm19 | RNA binding motif protein 19 [Source:MGI Symbol;Acc:MGI:1921361] | 0.82 | 2.04E-02 |
| Samhd1 | SAM domain and HD domain, 1 [Source:MGI Symbol;Acc:MGI:1927468] | 0.92 | 2.05E-02 |
| Calm4 | calmodulin 4 [Source:MGI Symbol;Acc:MGI:1931464] | 2.69 | 2.05E-02 |
| Mcm3 | minichromosome maintenance complex component 3 [Source:MGI Symbol;Acc:MGI:101845] | 1.26 | 2.06E-02 |
| Ankrd13a | ankyrin repeat domain 13a [Source:MGI Symbol;Acc:MGI:1915670] | 0.78 | 2.06E-02 |
| Tymp | thymidine phosphorylase [Source:MGI Symbol;Acc:MGI:1920212] | 1.12 | 2.06E-02 |
| Dsc2 | desmocollin 2 [Source:MGI Symbol;Acc:MGI:103221] | 2.11 | 2.06E-02 |
| 1110065P20Rik | RIKEN cDNA 1110065P20 gene [Source:MGI Symbol;Acc:MGI:1916170] | 1.98 | 2.06E-02 |
| Mapkapk2 | MAP kinase-activated protein kinase 2 [Source:MGI Symbol;Acc:MGI:109298] | 1.16 | 2.06E-02 |
| Evl | Ena-vasodilator stimulated phosphoprotein [Source:MGI Symbol;Acc:MGI:1194884] | 1.51 | 2.06E-02 |
| Coq10b | coenzyme Q10B [Source:MGI Symbol;Acc:MGI:1915126] | 0.92 | 2.07E-02 |
| Prok2 | prokineticin 2 [Source:MGI Symbol;Acc:MGI:1354178] | 3.28 | 2.07E-02 |
| Irak4 | interleukin-1 receptor-associated kinase 4 [Source:MGI Symbol;Acc:MGI:2182474] | 0.77 | 2.07E-02 |
| Gm10157 | predicted gene 10157 [Source:MGI Symbol;Acc:MGI:3642264] | 1.23 | 2.08E-02 |
| Csf1r | colony stimulating factor 1 receptor [Source:MGI Symbol;Acc:MGI:1339758] | 1.37 | 2.08E-02 |
| Sec61a1 | SEC61 translocon subunit alpha 1 [Source:MGI Symbol;Acc:MGI:1858417] | 0.89 | 2.08E-02 |
| Tle6 | transducin-like enhancer of split 6 [Source:MGI Symbol;Acc:MGI:2149593] | 1.24 | 2.09E-02 |
| Pglyrp3 | peptidoglycan recognition protein 3 [Source:MGI Symbol;Acc:MGI:2685266] | 0.97 | 2.09E-02 |
| Gm47072 | predicted gene, 47072 [Source:MGI Symbol;Acc:MGI:6095791] | 2.05 | 2.11E-02 |
| Cd33 | CD33 molecule [Source:MGI Symbol;Acc:MGI:99440] | 1.83 | 2.11E-02 |
| Nts | neurotensin [Source:MGI Symbol;Acc:MGI:1328351] | 3.07 | 2.13E-02 |
| Catip | ciliogenesis associated TTC17 interacting protein [Source:MGI Symbol;Acc:MGI:2685062] | 1.51 | 2.13E-02 |
| Kcnk12 | potassium channel, subfamily K, member 12 [Source:MGI Symbol;Acc:MGI:2684043] | 5.62 | 2.13E-02 |
| Hcls1 | hematopoietic cell specific Lyn substrate 1 [Source:MGI Symbol;Acc:MGI:104568] | 1.71 | 2.13E-02 |
| Ggt5 | gamma-glutamyltransferase 5 [Source:MGI Symbol;Acc:MGI:1346063] | 1.13 | 2.13E-02 |
| Glis1 | GLIS family zinc finger 1 [Source:MGI Symbol;Acc:MGI:2386723] | 2.27 | 2.13E-02 |
| Tmem14c | transmembrane protein 14C [Source:MGI Symbol;Acc:MGI:1913404] | 0.66 | 2.13E-02 |
| Cd68 | CD68 antigen [Source:MGI Symbol;Acc:MGI:88342] | 1.57 | 2.13E-02 |
| Il33 | interleukin 33 [Source:MGI Symbol;Acc:MGI:1924375] | 1.10 | 2.14E-02 |
| Psmd4 | proteasome (prosome, macropain) 26S subunit, non-ATPase, 4 [Source:MGI Symbol;Acc:MGI:1201670] | 0.84 | 2.14E-02 |
| Casp3 | caspase 3 [Source:MGI Symbol;Acc:MGI:107739] | 0.99 | 2.15E-02 |
| Tomm40 | translocase of outer mitochondrial membrane 40 [Source:MGI Symbol;Acc:MGI:1858259] | 1.02 | 2.16E-02 |
| Phgdh-ps1 | 3-phosphoglycerate dehydrogenase, pseudogene 1 [Source:MGI Symbol;Acc:MGI:1933203] | 1.80 | 2.16E-02 |
| Plekho2 | pleckstrin homology domain containing, family O member 2 [Source:MGI Symbol;Acc:MGI:2143132] | 1.28 | 2.17E-02 |
| Sdf2l1 | stromal cell-derived factor 2-like 1 [Source:MGI Symbol;Acc:MGI:2149842] | 1.01 | 2.17E-02 |
| Zswim4 | zinc finger SWIM-type containing 4 [Source:MGI Symbol;Acc:MGI:2443726] | 0.90 | 2.17E-02 |
| Ddx54 | DEAD box helicase 54 [Source:MGI Symbol;Acc:MGI:1919240] | 0.93 | 2.18E-02 |
| Rrbp1 | ribosome binding protein 1 [Source:MGI Symbol;Acc:MGI:1932395] | 0.92 | 2.18E-02 |

| | | | |
|---|---|---|---|
| Mpeg1 | macrophage expressed gene 1 [Source:MGI Symbol;Acc:MGI:1333743] | 1.42 | 2.18E-02 |
| Shank3 | SH3 and multiple ankyrin repeat domains 3 [Source:MGI Symbol;Acc:MGI:1930016] | 1.45 | 2.19E-02 |
| Osbpl10 | oxysterol binding protein-like 10 [Source:MGI Symbol;Acc:MGI:1921736] | 1.03 | 2.19E-02 |
| Ppp1r15a | protein phosphatase 1, regulatory subunit 15A [Source:MGI Symbol;Acc:MGI:1927072] | 1.06 | 2.19E-02 |
| Gm49947 | predicted gene, 49947 [Source:MGI Symbol;Acc:MGI:6270665] | 5.44 | 2.22E-02 |
| Fgd3 | FYVE, RhoGEF and PH domain containing 3 [Source:MGI Symbol;Acc:MGI:1353657] | 1.52 | 2.22E-02 |
| H4c4 | H4 clustered histone 4 [Source:MGI Symbol;Acc:MGI:2448423] | 1.36 | 2.22E-02 |
| Csrp2 | cysteine and glycine-rich protein 2 [Source:MGI Symbol;Acc:MGI:1202907] | 1.28 | 2.24E-02 |
| S100a4 | S100 calcium binding protein A4 [Source:MGI Symbol;Acc:MGI:1330282] | 1.45 | 2.25E-02 |
| Dennd3 | DENN domain containing 3 [Source:MGI Symbol;Acc:MGI:2146009] | 2.06 | 2.25E-02 |
| S100a6 | S100 calcium binding protein A6 (calcyclin) [Source:MGI Symbol;Acc:MGI:1339467] | 0.76 | 2.25E-02 |
| Peak1 | pseudopodium-enriched atypical kinase 1 [Source:MGI Symbol;Acc:MGI:2442366] | 0.88 | 2.25E-02 |
| Serpinb6d | serine (or cysteine) peptidase inhibitor, clade B, member 6d [Source:MGI Symbol;Acc:MGI:2667783] | 3.15 | 2.27E-02 |
| Ddx28 | DEAD box helicase 28 [Source:MGI Symbol;Acc:MGI:1919236] | 1.40 | 2.28E-02 |
| Eif1a | eukaryotic translation initiation factor 1A [Source:MGI Symbol;Acc:MGI:95298] | 0.91 | 2.28E-02 |
| Mgat3 | mannoside acetylglucosaminyltransferase 3 [Source:MGI Symbol;Acc:MGI:104532] | 1.36 | 2.28E-02 |
| Ctsk | cathepsin K [Source:MGI Symbol;Acc:MGI:107823] | 1.31 | 2.29E-02 |
| Cdipt | CDP-diacylglycerol--inositol 3-phosphatidyltransferase [Source:MGI Symbol;Acc:MGI:105491] | 0.72 | 2.29E-02 |
| Ssr2 | signal sequence receptor, beta [Source:MGI Symbol;Acc:MGI:1913506] | 0.91 | 2.29E-02 |
| Psma7 | proteasome subunit alpha 7 [Source:MGI Symbol;Acc:MGI:1347070] | 1.02 | 2.29E-02 |
| Irgc | immunity related GTPase cinema [Source:MGI Symbol;Acc:MGI:2685948] | 2.93 | 2.29E-02 |
| Galns | galactosamine (N-acetyl)-6-sulfatase [Source:MGI Symbol;Acc:MGI:1355303] | 1.42 | 2.29E-02 |
| Nop2 | NOP2 nucleolar protein [Source:MGI Symbol;Acc:MGI:107891] | 0.93 | 2.29E-02 |
| Atxn7l3 | ataxin 7-like 3 [Source:MGI Symbol;Acc:MGI:3036270] | 1.36 | 2.32E-02 |
| Apobec3 | apolipoprotein B mRNA editing enzyme, catalytic polypeptide 3 [Source:MGI Symbol;Acc:MGI:1933111] | 0.91 | 2.32E-02 |
| Gata2 | GATA binding protein 2 [Source:MGI Symbol;Acc:MGI:95662] | 1.19 | 2.32E-02 |
| Cops9 | COP9 signalosome subunit 9 [Source:MGI Symbol;Acc:MGI:1914165] | 1.35 | 2.32E-02 |
| Rcc1 | regulator of chromosome condensation 1 [Source:MGI Symbol;Acc:MGI:1913989] | 1.06 | 2.34E-02 |
| Gpx1 | glutathione peroxidase 1 [Source:MGI Symbol;Acc:MGI:104887] | 1.01 | 2.34E-02 |
| Gm14767 | predicted gene 14767 [Source:MGI Symbol;Acc:MGI:3705402] | 2.59 | 2.35E-02 |
| Bak1 | BCL2-antagonist/killer 1 [Source:MGI Symbol;Acc:MGI:1097161] | 1.00 | 2.35E-02 |
| Ndrg4 | N-myc downstream regulated gene 4 [Source:MGI Symbol;Acc:MGI:2384590] | 1.18 | 2.35E-02 |
| Sptlc2 | serine palmitoyltransferase, long chain base subunit 2 [Source:MGI Symbol;Acc:MGI:108074] | 0.68 | 2.35E-02 |
| Rap1gap | Rap1 GTPase-activating protein [Source:MGI Symbol;Acc:MGI:109338] | 1.00 | 2.36E-02 |
| Gm15740 | predicted gene 15740 [Source:MGI Symbol;Acc:MGI:3783182] | 2.73 | 2.36E-02 |
| Pfdn1 | prefoldin 1 [Source:MGI Symbol;Acc:MGI:1914449] | 0.86 | 2.38E-02 |
| Mdfi | MyoD family inhibitor [Source:MGI Symbol;Acc:MGI:107687] | 1.18 | 2.39E-02 |
| Dkk3 | dickkopf WNT signaling pathway inhibitor 3 [Source:MGI Symbol;Acc:MGI:1354952] | 1.77 | 2.39E-02 |
| Src | Rous sarcoma oncogene [Source:MGI Symbol;Acc:MGI:98397] | 0.86 | 2.39E-02 |
| Rab44 | RAB44, member RAS oncogene family [Source:MGI Symbol;Acc:MGI:3045302] | 1.20 | 2.39E-02 |
| Maged2 | MAGE family member D2 [Source:MGI Symbol;Acc:MGI:1933391] | 1.39 | 2.40E-02 |
| Arhgap30 | Rho GTPase activating protein 30 [Source:MGI Symbol;Acc:MGI:2684948] | 1.09 | 2.40E-02 |
| Tnfsfm13 | tumor necrosis factor (ligand) superfamily, membrane-bound member 13 [Source:MGI Symbol;Acc:MGI:3845075] | 1.34 | 2.40E-02 |
| Ccno | cyclin O [Source:MGI Symbol;Acc:MGI:2145534] | 2.79 | 2.41E-02 |
| Trem6l | triggering receptor expressed on myeloid cells-like 6 [Source:MGI Symbol;Acc:MGI:2443478] | 1.42 | 2.42E-02 |
| Sdc1 | syndecan 1 [Source:MGI Symbol;Acc:MGI:1349162] | 0.94 | 2.42E-02 |
| Fasl | Fas ligand [Source:MGI Symbol;Acc:MGI:99255] | 2.63 | 2.42E-02 |
| Cd81 | CD81 antigen [Source:MGI Symbol;Acc:MGI:1096398] | 0.81 | 2.43E-02 |
| H4c9 | H4 clustered histone 9 [Source:MGI Symbol;Acc:MGI:2448432] | 1.59 | 2.44E-02 |
| Rrad | Ras-related associated with diabetes [Source:MGI Symbol;Acc:MGI:1930943] | 2.37 | 2.44E-02 |
| Akt1s1 | AKT1 substrate 1 [Source:MGI Symbol;Acc:MGI:1914855] | 1.45 | 2.44E-02 |
| Nxt1 | NTF2-related export protein 1 [Source:MGI Symbol;Acc:MGI:1929619] | 0.80 | 2.44E-02 |
| Atp12a | ATPase, H+/K+ transporting, nongastric, alpha polypeptide [Source:MGI Symbol;Acc:MGI:1926943] | 1.79 | 2.44E-02 |
| Col6a5 | collagen, type VI, alpha 5 [Source:MGI Symbol;Acc:MGI:3648134] | 1.74 | 2.44E-02 |
| Psg18 | pregnancy specific beta-1-glycoprotein 18 [Source:MGI Symbol;Acc:MGI:1347251] | 5.52 | 2.44E-02 |
| Pear1 | platelet endothelial aggregation receptor 1 [Source:MGI Symbol;Acc:MGI:1920432] | 1.29 | 2.46E-02 |
| Lilra6 | leukocyte immunoglobulin-like receptor, subfamily A (with TM domain), member 6 [Source:MGI Symbol;Acc:MGI:1195969] | 2.32 | 2.46E-02 |
| Notch4 | notch 4 [Source:MGI Symbol;Acc:MGI:107471] | 1.38 | 2.47E-02 |
| Lif | leukemia inhibitory factor [Source:MGI Symbol;Acc:MGI:96787] | 1.52 | 2.50E-02 |
| Abhd17a | abhydrolase domain containing 17A [Source:MGI Symbol;Acc:MGI:106388] | 1.53 | 2.51E-02 |
| Tmem269 | transmembrane protein 269 [Source:MGI Symbol;Acc:MGI:1922430] | 2.34 | 2.51E-02 |
| Ldhc | lactate dehydrogenase C [Source:MGI Symbol;Acc:MGI:96764] | 4.10 | 2.53E-02 |
| Klf7 | Kruppel-like transcription factor 7 (ubiquitous) [Source:MGI Symbol;Acc:MGI:1935151] | 0.85 | 2.54E-02 |
| Etv6 | ets variant 6 [Source:MGI Symbol;Acc:MGI:109336] | 0.77 | 2.55E-02 |
| Fgf7 | fibroblast growth factor 7 [Source:MGI Symbol;Acc:MGI:95521] | 2.41 | 2.56E-02 |
| Lgals3bp | lectin, galactoside-binding, soluble, 3 binding protein [Source:MGI Symbol;Acc:MGI:99554] | 1.44 | 2.56E-02 |
| Osgin1 | oxidative stress induced growth inhibitor 1 [Source:MGI Symbol;Acc:MGI:1919089] | 1.18 | 2.58E-02 |
| Il2rg | interleukin 2 receptor, gamma chain [Source:MGI Symbol;Acc:MGI:96551] | 1.69 | 2.58E-02 |
| Sparc | secreted acidic cysteine rich glycoprotein [Source:MGI Symbol;Acc:MGI:98373] | 1.26 | 2.58E-02 |
| Gpr39 | G protein-coupled receptor 39 [Source:MGI Symbol;Acc:MGI:1918361] | 3.68 | 2.59E-02 |
| P3h1 | prolyl 3-hydroxylase 1 [Source:MGI Symbol;Acc:MGI:1888921] | 1.63 | 2.59E-02 |

| | | | |
|---|---|---|---|
| Flna | filamin, alpha [Source:MGI Symbol;Acc:MGI:95556] | 1.01 | 2.62E-02 |
| Fblim1 | filamin binding LIM protein 1 [Source:MGI Symbol;Acc:MGI:1921452] | 0.96 | 2.62E-02 |
| Adrm1 | adhesion regulating molecule 1 26S proteasome ubiquitin receptor [Source:MGI Symbol;Acc:MGI:1929289] | 0.91 | 2.62E-02 |
| Ngf | nerve growth factor [Source:MGI Symbol;Acc:MGI:97321] | 1.75 | 2.62E-02 |
| Uox | urate oxidase [Source:MGI Symbol;Acc:MGI:98907] | 1.58 | 2.63E-02 |
| Rpp25l | ribonuclease P/MRP 25 subunit-like [Source:MGI Symbol;Acc:MGI:1917211] | 1.55 | 2.63E-02 |
| Selplg | selectin, platelet (p-selectin) ligand [Source:MGI Symbol;Acc:MGI:106689] | 1.47 | 2.63E-02 |
| Hsd17b2 | hydroxysteroid (17-beta) dehydrogenase 2 [Source:MGI Symbol;Acc:MGI:1096386] | 1.39 | 2.63E-02 |
| Spaca4 | sperm acrosome associated 4 [Source:MGI Symbol;Acc:MGI:1916613] | 2.91 | 2.63E-02 |
| Gpm6b | glycoprotein m6b [Source:MGI Symbol;Acc:MGI:107672] | 1.88 | 2.64E-02 |
| Cd52 | CD52 antigen [Source:MGI Symbol;Acc:MGI:1346088] | 1.22 | 2.64E-02 |
| Rnf26 | ring finger protein 26 [Source:MGI Symbol;Acc:MGI:2388131] | 0.86 | 2.65E-02 |
| Psmc4 | proteasome (prosome, macropain) 26S subunit, ATPase, 4 [Source:MGI Symbol;Acc:MGI:1346093] | 0.71 | 2.66E-02 |
| Timm50 | translocase of inner mitochondrial membrane 50 [Source:MGI Symbol;Acc:MGI:1913775] | 0.82 | 2.66E-02 |
| Rac2 | Rac family small GTPase 2 [Source:MGI Symbol;Acc:MGI:97846] | 1.38 | 2.67E-02 |
| Mybph | myosin binding protein H [Source:MGI Symbol;Acc:MGI:1858196] | 2.86 | 2.67E-02 |
| Sh3tc1 | SH3 domain and tetratricopeptide repeats 1 [Source:MGI Symbol;Acc:MGI:2678949] | 1.09 | 2.69E-02 |
| Gpr31b | G protein-coupled receptor 31, D17Leh66b region [Source:MGI Symbol;Acc:MGI:1354372] | 1.85 | 2.70E-02 |
| Exoc3l4 | exocyst complex component 3-like 4 [Source:MGI Symbol;Acc:MGI:1921363] | 1.68 | 2.70E-02 |
| Actn1 | actinin, alpha 1 [Source:MGI Symbol;Acc:MGI:2137706] | 1.01 | 2.70E-02 |
| Gm21451 | predicted gene, 21451 [Source:MGI Symbol;Acc:MGI:5434806] | 1.70 | 2.72E-02 |
| Fcgr3 | Fc receptor, IgG, low affinity III [Source:MGI Symbol;Acc:MGI:95500] | 1.69 | 2.73E-02 |
| Dbn1 | drebrin 1 [Source:MGI Symbol;Acc:MGI:1931838] | 1.41 | 2.73E-02 |
| Ldlrap1 | low density lipoprotein receptor adaptor protein 1 [Source:MGI Symbol;Acc:MGI:2140175] | 0.84 | 2.73E-02 |
| Lair1 | leukocyte-associated Ig-like receptor 1 [Source:MGI Symbol;Acc:MGI:105492] | 1.19 | 2.74E-02 |
| Gpaa1 | GPI anchor attachment protein 1 [Source:MGI Symbol;Acc:MGI:1202392] | 1.25 | 2.75E-02 |
| Mydgf | myeloid derived growth factor [Source:MGI Symbol;Acc:MGI:2156020] | 0.71 | 2.75E-02 |
| Kcnc3 | potassium voltage gated channel, Shaw-related subfamily, member 3 [Source:MGI Symbol;Acc:MGI:96669] | 1.31 | 2.75E-02 |
| Gm6030 | predicted gene 6030 [Source:MGI Symbol;Acc:MGI:3645112] | 2.15 | 2.75E-02 |
| Slc24a1 | solute carrier family 24 (sodium/potassium/calcium exchanger), member 1 [Source:MGI Symbol;Acc:MGI:2384871] | 1.49 | 2.75E-02 |
| Lrrc75a | leucine rich repeat containing 75A [Source:MGI Symbol;Acc:MGI:2682293] | 0.93 | 2.75E-02 |
| Slc35f6 | solute carrier family 35, member F6 [Source:MGI Symbol;Acc:MGI:1922169] | 0.74 | 2.75E-02 |
| Gm7599 | predicted gene 7599 [Source:MGI Symbol;Acc:MGI:3644309] | 1.38 | 2.76E-02 |
| Clstn3 | calsyntenin 3 [Source:MGI Symbol;Acc:MGI:2178323] | 1.08 | 2.76E-02 |
| H4c8 | H4 clustered histone 8 [Source:MGI Symbol;Acc:MGI:2448427] | 1.47 | 2.78E-02 |
| Doc2a | double C2, alpha [Source:MGI Symbol;Acc:MGI:109446] | 3.00 | 2.78E-02 |
| Hp | haptoglobin [Source:MGI Symbol;Acc:MGI:96211] | 2.02 | 2.79E-02 |
| Cdc42ep4 | CDC42 effector protein 4 [Source:MGI Symbol;Acc:MGI:1929760] | 0.95 | 2.79E-02 |
| Manf | mesencephalic astrocyte-derived neurotrophic factor [Source:MGI Symbol;Acc:MGI:1922090] | 0.95 | 2.79E-02 |
| Trem4 | triggering receptor expressed on myeloid cells 4 [Source:MGI Symbol;Acc:MGI:2157854] | 1.09 | 2.79E-02 |
| Gm3126 | predicted gene 3126 [Source:MGI Symbol;Acc:MGI:3781302] | 5.62 | 2.80E-02 |
| Eif4a1 | eukaryotic translation initiation factor 4A1 [Source:MGI Symbol;Acc:MGI:95303] | 0.77 | 2.80E-02 |
| Sh3gl1 | SH3-domain GRB2-like 1 [Source:MGI Symbol;Acc:MGI:700010] | 0.80 | 2.83E-02 |
| Ecscr | endothelial cell surface expressed chemotaxis and apoptosis regulator [Source:MGI Symbol;Acc:MGI:1915795] | 1.78 | 2.83E-02 |
| Sephs2 | selenophosphate synthetase 2 [Source:MGI Symbol;Acc:MGI:108388] | 1.28 | 2.83E-02 |
| Morf4l1-ps1 | mortality factor 4 like 1, pseudogene 1 [Source:MGI Symbol;Acc:MGI:3612158] | 1.04 | 2.87E-02 |
| H3c7 | H3 clustered histone 7 [Source:MGI Symbol;Acc:MGI:2448329] | 1.63 | 2.88E-02 |
| 4930503B20Rik | RIKEN cDNA 4930503B20 gene [Source:MGI Symbol;Acc:MGI:1922264] | 3.12 | 2.89E-02 |
| Pld1 | phospholipase D1 [Source:MGI Symbol;Acc:MGI:109585] | 0.77 | 2.90E-02 |
| Pde1b | phosphodiesterase 1B, Ca2+-calmodulin dependent [Source:MGI Symbol;Acc:MGI:97523] | 1.53 | 2.90E-02 |
| Pgls | 6-phosphogluconolactonase [Source:MGI Symbol;Acc:MGI:1913421] | 1.24 | 2.91E-02 |
| P4ha3 | procollagen-proline, 2-oxoglutarate 4-dioxygenase (proline 4-hydroxylase), alpha polypeptide III [Source:MGI Symbol;Acc:MGI:2444049] | 1.16 | 2.92E-02 |
| Gm867 | predicted gene 867 [Source:MGI Symbol;Acc:MGI:2685713] | 2.46 | 2.95E-02 |
| Cd6 | CD6 antigen [Source:MGI Symbol;Acc:MGI:103566] | 1.42 | 2.95E-02 |
| Yif1b | Yip1 interacting factor homolog B (S. cerevisiae) [Source:MGI Symbol;Acc:MGI:1924504] | 0.87 | 2.95E-02 |
| Sprr1a | small proline-rich protein 1A [Source:MGI Symbol;Acc:MGI:106660] | 2.11 | 2.95E-02 |
| Lrp10 | low-density lipoprotein receptor-related protein 10 [Source:MGI Symbol;Acc:MGI:1929480] | 0.91 | 2.96E-02 |
| Pcyt1b | phosphate cytidylyltransferase 1, choline, beta isoform [Source:MGI Symbol;Acc:MGI:2147987] | 1.26 | 2.98E-02 |
| Gpr162 | G protein-coupled receptor 162 [Source:MGI Symbol;Acc:MGI:1315214] | 1.01 | 2.99E-02 |
| Lamb1 | laminin B1 [Source:MGI Symbol;Acc:MGI:96743] | 1.51 | 2.99E-02 |
| Pycr2 | pyrroline-5-carboxylate reductase family, member 2 [Source:MGI Symbol;Acc:MGI:1277956] | 0.99 | 2.99E-02 |
| Ifit1bl1 | interferon induced protein with tetratricpeptide repeats 1B like 1 [Source:MGI Symbol;Acc:MGI:3650685] | 1.98 | 3.00E-02 |
| Bok | BCL2-related ovarian killer [Source:MGI Symbol;Acc:MGI:1858494] | 1.54 | 3.01E-02 |
| Sirpb1a | signal-regulatory protein beta 1A [Source:MGI Symbol;Acc:MGI:2444824] | 2.52 | 3.01E-02 |
| Stard3 | StAR related lipid transfer domain containing 3 [Source:MGI Symbol;Acc:MGI:1929618] | 0.94 | 3.03E-02 |
| Wdr1 | WD repeat domain 1 [Source:MGI Symbol;Acc:MGI:1337100] | 0.64 | 3.05E-02 |
| Gm6330 | predicted gene 6330 [Source:MGI Symbol;Acc:MGI:3643757] | 1.69 | 3.05E-02 |
| Tmem88 | transmembrane protein 88 [Source:MGI Symbol;Acc:MGI:1914270] | 1.72 | 3.06E-02 |
| Mt2 | metallothionein 2 [Source:MGI Symbol;Acc:MGI:97172] | 1.36 | 3.07E-02 |

| | | | |
|---|---|---|---|
| Ppm1g | protein phosphatase 1G (formerly 2C), magnesium-dependent, gamma isoform [Source:MGI Symbol;Acc:MGI:106065] | 0.78 | 3.07E-02 |
| H2az1 | H2A.Z variant histone 1 [Source:MGI Symbol;Acc:MGI:1888388] | 1.10 | 3.09E-02 |
| Trmt61a | tRNA methyltransferase 61A [Source:MGI Symbol;Acc:MGI:2443487] | 1.14 | 3.10E-02 |
| Numbl | numb-like [Source:MGI Symbol;Acc:MGI:894702] | 0.85 | 3.10E-02 |
| Tiparp | TCDD-inducible poly(ADP-ribose) polymerase [Source:MGI Symbol;Acc:MGI:2159210] | 0.80 | 3.11E-02 |
| B3gnt3 | UDP-GlcNAc:betaGal beta-1,3-N-acetylglucosaminyltransferase 3 [Source:MGI Symbol;Acc:MGI:2152535] | 1.27 | 3.11E-02 |
| Gm4240 | predicted gene 4240 [Source:MGI Symbol;Acc:MGI:3782417] | 2.84 | 3.12E-02 |
| Nlgn2 | neuroligin 2 [Source:MGI Symbol;Acc:MGI:2681835] | 1.27 | 3.12E-02 |
| Gm45140 | predicted gene 45140 [Source:MGI Symbol;Acc:MGI:5753716] | 1.13 | 3.12E-02 |
| Nifk | nucleolar protein interacting with the FHA domain of MKI67 [Source:MGI Symbol;Acc:MGI:1915199] | 0.75 | 3.13E-02 |
| Gm10705 | predicted gene 10705 [Source:MGI Symbol;Acc:MGI:3708678] | 1.77 | 3.13E-02 |
| Map1s | microtubule-associated protein 1S [Source:MGI Symbol;Acc:MGI:2443304] | 1.32 | 3.16E-02 |
| Gm20532 | predicted gene 20532 [Source:MGI Symbol;Acc:MGI:5141997] | 1.58 | 3.17E-02 |
| Yrdc | yrdC domain containing (E.coli) [Source:MGI Symbol;Acc:MGI:2387201] | 0.79 | 3.19E-02 |
| Kcnv1 | potassium channel, subfamily V, member 1 [Source:MGI Symbol;Acc:MGI:1914748] | 3.95 | 3.19E-02 |
| Loxl2 | lysyl oxidase-like 2 [Source:MGI Symbol;Acc:MGI:2137913] | 1.76 | 3.19E-02 |
| Cdt1 | chromatin licensing and DNA replication factor 1 [Source:MGI Symbol;Acc:MGI:1914427] | 1.26 | 3.21E-02 |
| Tmem217rt | transmembrane 217, retrotransposed [Source:MGI Symbol;Acc:MGI:5662654] | 2.02 | 3.21E-02 |
| Spire1 | spire type actin nucleation factor 1 [Source:MGI Symbol;Acc:MGI:1915416] | 1.10 | 3.21E-02 |
| Gm4786 | predicted gene 4786 [Source:MGI Symbol;Acc:MGI:3644879] | 3.51 | 3.23E-02 |
| Siglec1 | sialic acid binding Ig-like lectin 1, sialoadhesin [Source:MGI Symbol;Acc:MGI:99668] | 1.49 | 3.24E-02 |
| Pth1r | parathyroid hormone 1 receptor [Source:MGI Symbol;Acc:MGI:97801] | 1.07 | 3.24E-02 |
| Polr2l | polymerase (RNA) II (DNA directed) polypeptide L [Source:MGI Symbol;Acc:MGI:1913741] | 1.04 | 3.27E-02 |
| Fam162a | family with sequence similarity 162, member A [Source:MGI Symbol;Acc:MGI:1917436] | 1.84 | 3.27E-02 |
| Acox2 | acyl-Coenzyme A oxidase 2, branched chain [Source:MGI Symbol;Acc:MGI:1934852] | 0.96 | 3.28E-02 |
| Fam171a1 | family with sequence similarity 171, member A1 [Source:MGI Symbol;Acc:MGI:2442917] | 1.14 | 3.29E-02 |
| Apobr | apolipoprotein B receptor [Source:MGI Symbol;Acc:MGI:2176230] | 1.33 | 3.30E-02 |
| Akap17a | A-kinase anchoring protein 17A [Source:MGI Symbol;Acc:MGI:6723883] | 1.68 | 3.30E-02 |
| Bcl2a1b | B cell leukemia/lymphoma 2 related protein A1b [Source:MGI Symbol;Acc:MGI:1278326] | 2.57 | 3.31E-02 |
| Rgs16 | regulator of G-protein signaling 16 [Source:MGI Symbol;Acc:MGI:108407] | 1.96 | 3.31E-02 |
| Acap1 | ArfGAP with coiled-coil, ankyrin repeat and PH domains 1 [Source:MGI Symbol;Acc:MGI:2388270] | 1.12 | 3.31E-02 |
| Agpat1 | 1-acylglycerol-3-phosphate O-acyltransferase 1 [Source:MGI Symbol;Acc:MGI:1932075] | 0.69 | 3.31E-02 |
| Rps14 | ribosomal protein S14 [Source:MGI Symbol;Acc:MGI:98107] | 1.37 | 3.32E-02 |
| Zbp1 | Z-DNA binding protein 1 [Source:MGI Symbol;Acc:MGI:1927449] | 1.38 | 3.32E-02 |
| Flt4 | FMS-like tyrosine kinase 4 [Source:MGI Symbol;Acc:MGI:95561] | 1.35 | 3.33E-02 |
| Cfap69 | cilia and flagella associated protein 69 [Source:MGI Symbol;Acc:MGI:2443778] | 2.03 | 3.33E-02 |
| Snx8 | sorting nexin 8 [Source:MGI Symbol;Acc:MGI:2443816] | 1.02 | 3.33E-02 |
| Mcm2 | minichromosome maintenance complex component 2 [Source:MGI Symbol;Acc:MGI:105380] | 0.94 | 3.35E-02 |
| Bst1 | bone marrow stromal cell antigen 1 [Source:MGI Symbol;Acc:MGI:105370] | 1.22 | 3.36E-02 |
| Vim | vimentin [Source:MGI Symbol;Acc:MGI:98932] | 1.18 | 3.38E-02 |
| Tradd | TNFRSF1A-associated via death domain [Source:MGI Symbol;Acc:MGI:109200] | 1.10 | 3.39E-02 |
| Myo9b | myosin IXb [Source:MGI Symbol;Acc:MGI:106624] | 0.79 | 3.39E-02 |
| Jph3 | junctophilin 3 [Source:MGI Symbol;Acc:MGI:1891497] | 1.01 | 3.42E-02 |
| Pef1 | penta-EF hand domain containing 1 [Source:MGI Symbol;Acc:MGI:1915148] | 0.65 | 3.42E-02 |
| Psmc3 | proteasome (prosome, macropain) 26S subunit, ATPase 3 [Source:MGI Symbol;Acc:MGI:1098754] | 0.74 | 3.43E-02 |
| Ranbp1 | RAN binding protein 1 [Source:MGI Symbol;Acc:MGI:96269] | 0.98 | 3.43E-02 |
| H2ac18 | H2A clustered histone 18 [Source:MGI Symbol;Acc:MGI:96097] | 1.56 | 3.44E-02 |
| Snrpe | small nuclear ribonucleoprotein E [Source:MGI Symbol;Acc:MGI:98346] | 0.92 | 3.44E-02 |
| Kif17 | kinesin family member 17 [Source:MGI Symbol;Acc:MGI:1098229] | 1.35 | 3.45E-02 |
| Gm48583 | predicted gene, 48583 [Source:MGI Symbol;Acc:MGI:6098152] | 1.28 | 3.47E-02 |
| Ncf1 | neutrophil cytosolic factor 1 [Source:MGI Symbol;Acc:MGI:97283] | 1.18 | 3.49E-02 |
| Gm49395 | predicted gene, 49395 [Source:MGI Symbol;Acc:MGI:6121627] | 1.16 | 3.49E-02 |
| Map3k10 | mitogen-activated protein kinase kinase kinase 10 [Source:MGI Symbol;Acc:MGI:1346879] | 1.36 | 3.50E-02 |
| Gm9790 | predicted gene 9790 [Source:MGI Symbol;Acc:MGI:3704221] | 5.36 | 3.51E-02 |
| Brk1 | BRICK1, SCAR/WAVE actin-nucleating complex subunit [Source:MGI Symbol;Acc:MGI:1915406] | 0.70 | 3.51E-02 |
| Gabrb1 | gamma-aminobutyric acid type A receptor subunit beta 1 [Source:MGI Symbol;Acc:MGI:95619] | 0.77 | 3.52E-02 |
| B4galt2 | UDP-Gal:betaGlcNAc beta 1,4- galactosyltransferase, polypeptide 2 [Source:MGI Symbol;Acc:MGI:1858493] | 1.32 | 3.52E-02 |
| Rps8 | ribosomal protein S8 [Source:MGI Symbol;Acc:MGI:98166] | 1.45 | 3.53E-02 |
| Cbr3 | carbonyl reductase 3 [Source:MGI Symbol;Acc:MGI:1309992] | 1.75 | 3.54E-02 |
| H1f2 | H1.2 linker histone, cluster member [Source:MGI Symbol;Acc:MGI:1931526] | 1.63 | 3.54E-02 |
| Tbx3 | T-box 3 [Source:MGI Symbol;Acc:MGI:98495] | 1.34 | 3.54E-02 |
| Hps1 | HPS1, biogenesis of lysosomal organelles complex 3 subunit 1 [Source:MGI Symbol;Acc:MGI:2177763] | 0.93 | 3.54E-02 |
| Arap3 | ArfGAP with RhoGAP domain, ankyrin repeat and PH domain 3 [Source:MGI Symbol;Acc:MGI:2147274] | 1.18 | 3.54E-02 |
| Ankrd66 | ankyrin repeat domain 66 [Source:MGI Symbol;Acc:MGI:1925106] | 3.04 | 3.54E-02 |
| Slc2a6 | solute carrier family 2 (facilitated glucose transporter), member 6 [Source:MGI Symbol;Acc:MGI:2443286] | 1.50 | 3.55E-02 |
| Alox5ap | arachidonate 5-lipoxygenase activating protein [Source:MGI Symbol;Acc:MGI:107505] | 1.46 | 3.55E-02 |
| Mrpl12 | mitochondrial ribosomal protein L12 [Source:MGI Symbol;Acc:MGI:1926273] | 0.86 | 3.55E-02 |
| Snx17 | sorting nexin 17 [Source:MGI Symbol;Acc:MGI:2387801] | 0.65 | 3.55E-02 |
| Mtcl2 | microtubule crosslinking factor 2 [Source:MGI Symbol;Acc:MGI:2444575] | 1.17 | 3.56E-02 |
| Nfkbie | nuclear factor of kappa light polypeptide gene enhancer in B cells inhibitor, epsilon [Source:MGI Symbol;Acc:MGI:1194908] | 1.40 | 3.57E-02 |

| | | | |
|---|---|---|---|
| Plxnd1 | plexin D1 [Source:MGI Symbol;Acc:MGI:2154244] | 1.05 | 3.58E-02 |
| B4galt1 | UDP-Gal:betaGlcNAc beta 1,4- galactosyltransferase, polypeptide 1 [Source:MGI Symbol;Acc:MGI:95705] | 0.87 | 3.58E-02 |
| Tmem208 | transmembrane protein 208 [Source:MGI Symbol;Acc:MGI:1913570] | 0.82 | 3.58E-02 |
| Lpcat1 | lysophosphatidylcholine acyltransferase 1 [Source:MGI Symbol;Acc:MGI:2384812] | 0.61 | 3.58E-02 |
| Cit | citron [Source:MGI Symbol;Acc:MGI:105313] | 0.97 | 3.59E-02 |
| Mmp23 | matrix metallopeptidase 23 [Source:MGI Symbol;Acc:MGI:1347361] | 1.29 | 3.60E-02 |
| Clip3 | CAP-GLY domain containing linker protein 3 [Source:MGI Symbol;Acc:MGI:1923936] | 1.22 | 3.60E-02 |
| Lum | lumican [Source:MGI Symbol;Acc:MGI:109347] | 1.94 | 3.61E-02 |
| Slc25a25 | solute carrier family 25 (mitochondrial carrier, phosphate carrier), member 25 [Source:MGI Symbol;Acc:MGI:1915913] | 0.64 | 3.61E-02 |
| Psmb6 | proteasome (prosome, macropain) subunit, beta type 6 [Source:MGI Symbol;Acc:MGI:104880] | 0.76 | 3.62E-02 |
| Gal | galanin and GMAP prepropeptide [Source:MGI Symbol;Acc:MGI:95637] | 2.65 | 3.62E-02 |
| Itga11 | integrin alpha 11 [Source:MGI Symbol;Acc:MGI:2442114] | 1.76 | 3.62E-02 |
| Sipa1 | signal-induced proliferation associated gene 1 [Source:MGI Symbol;Acc:MGI:107576] | 1.15 | 3.62E-02 |
| Naa38 | N(alpha)-acetyltransferase 38, NatC auxiliary subunit [Source:MGI Symbol;Acc:MGI:1925554] | 0.89 | 3.63E-02 |
| Myo10 | myosin X [Source:MGI Symbol;Acc:MGI:107716] | 0.67 | 3.63E-02 |
| Bdh1 | 3-hydroxybutyrate dehydrogenase, type 1 [Source:MGI Symbol;Acc:MGI:1919161] | 1.41 | 3.64E-02 |
| Spx | spexin hormone [Source:MGI Symbol;Acc:MGI:2442262] | 1.15 | 3.64E-02 |
| Cox8a | cytochrome c oxidase subunit 8A [Source:MGI Symbol;Acc:MGI:105959] | 0.82 | 3.64E-02 |
| Mpi | mannose phosphate isomerase [Source:MGI Symbol;Acc:MGI:97075] | 0.64 | 3.64E-02 |
| H2ax | H2A.X variant histone [Source:MGI Symbol;Acc:MGI:102688] | 1.86 | 3.64E-02 |
| Galnt3 | polypeptide N-acetylgalactosaminyltransferase 3 [Source:MGI Symbol;Acc:MGI:894695] | 0.82 | 3.64E-02 |
| Gfra4 | glial cell line derived neurotrophic factor family receptor alpha 4 [Source:MGI Symbol;Acc:MGI:1341873] | 1.58 | 3.66E-02 |
| Tamalin | trafficking regulator and scaffold protein tamalin [Source:MGI Symbol;Acc:MGI:1860303] | 0.95 | 3.68E-02 |
| Pigs | phosphatidylinositol glycan anchor biosynthesis, class S [Source:MGI Symbol;Acc:MGI:2687325] | 0.92 | 3.68E-02 |
| Cxcr5 | C-X-C motif chemokine receptor 5 [Source:MGI Symbol;Acc:MGI:103567] | 1.68 | 3.68E-02 |
| Chst14 | carbohydrate sulfotransferase 14 [Source:MGI Symbol;Acc:MGI:1919386] | 0.98 | 3.68E-02 |
| H2ac11 | H2A clustered histone 11 [Source:MGI Symbol;Acc:MGI:2448293] | 1.86 | 3.69E-02 |
| Gm43064 | predicted gene 43064 [Source:MGI Symbol;Acc:MGI:5663201] | 6.22 | 3.69E-02 |
| Fastk | Fas-activated serine/threonine kinase [Source:MGI Symbol;Acc:MGI:1913837] | 0.63 | 3.69E-02 |
| Tmem121b | transmembrane protein 121B [Source:MGI Symbol;Acc:MGI:2136977] | 2.58 | 3.69E-02 |
| Cntn2 | contactin 2 [Source:MGI Symbol;Acc:MGI:104518] | 1.81 | 3.70E-02 |
| Best1 | bestrophin 1 [Source:MGI Symbol;Acc:MGI:1346332] | 1.51 | 3.70E-02 |
| Pcdhb21 | protocadherin beta 21 [Source:MGI Symbol;Acc:MGI:2136759] | 0.96 | 3.71E-02 |
| Foxd3 | forkhead box D3 [Source:MGI Symbol;Acc:MGI:1347473] | 1.62 | 3.71E-02 |
| Hsp90aa1 | heat shock protein 90, alpha (cytosolic), class A member 1 [Source:MGI Symbol;Acc:MGI:96250] | 0.60 | 3.71E-02 |
| Srm | spermidine synthase [Source:MGI Symbol;Acc:MGI:102690] | 1.24 | 3.72E-02 |
| Zfp998 | zinc finger protein 998 [Source:MGI Symbol;Acc:MGI:1924053] | 1.74 | 3.72E-02 |
| Klk9 | kallikrein related-peptidase 9 [Source:MGI Symbol;Acc:MGI:1921082] | 2.33 | 3.77E-02 |
| Snx25 | sorting nexin 25 [Source:MGI Symbol;Acc:MGI:2142610] | 0.74 | 3.78E-02 |
| Dda1 | DET1 and DDB1 associated 1 [Source:MGI Symbol;Acc:MGI:1913748] | 0.62 | 3.78E-02 |
| Dhrs1 | dehydrogenase/reductase 1 [Source:MGI Symbol;Acc:MGI:1196314] | 0.66 | 3.79E-02 |
| Arpc4 | actin related protein 2/3 complex, subunit 4 [Source:MGI Symbol;Acc:MGI:1915339] | 1.04 | 3.79E-02 |
| Tmprss11g | transmembrane protease, serine 11g [Source:MGI Symbol;Acc:MGI:2444058] | 1.44 | 3.80E-02 |
| Gm11249 | predicted gene 11249 [Source:MGI Symbol;Acc:MGI:3650834] | 2.46 | 3.80E-02 |
| Basp1 | brain abundant, membrane attached signal protein 1 [Source:MGI Symbol;Acc:MGI:1917600] | 1.72 | 3.81E-02 |
| Nr2f1 | nuclear receptor subfamily 2, group F, member 1 [Source:MGI Symbol;Acc:MGI:1352451] | 1.26 | 3.81E-02 |
| Cend1 | cell cycle exit and neuronal differentiation 1 [Source:MGI Symbol;Acc:MGI:1929898] | 3.23 | 3.83E-02 |
| Surf2 | surfeit gene 2 [Source:MGI Symbol;Acc:MGI:98444] | 0.74 | 3.83E-02 |
| Zic4 | zinc finger protein of the cerebellum 4 [Source:MGI Symbol;Acc:MGI:107201] | 2.65 | 3.83E-02 |
| Gm7889 | predicted gene 7889 [Source:MGI Symbol;Acc:MGI:3648247] | 5.60 | 3.83E-02 |
| Adap2 | ArfGAP with dual PH domains 2 [Source:MGI Symbol;Acc:MGI:2663075] | 1.18 | 3.83E-02 |
| Rpl22l1 | ribosomal protein L22 like 1 [Source:MGI Symbol;Acc:MGI:1915278] | 0.89 | 3.83E-02 |
| Fscn2 | fascin actin-bundling protein 2 [Source:MGI Symbol;Acc:MGI:2443337] | 1.07 | 3.83E-02 |
| Epb41l3 | erythrocyte membrane protein band 4.1 like 3 [Source:MGI Symbol;Acc:MGI:103008] | 0.79 | 3.83E-02 |
| Gm7775 | predicted gene 7775 [Source:MGI Symbol;Acc:MGI:3646054] | 2.40 | 3.83E-02 |
| Gm12551 | predicted gene 12551 [Source:MGI Symbol;Acc:MGI:3651664] | 1.86 | 3.85E-02 |
| Gm12551 | perilipin 2 pseudogene [Source:NCBI gene (formerly Entrezgene);Acc:101055843] | 1.86 | 3.85E-02 |
| Psap | prosaposin [Source:MGI Symbol;Acc:MGI:97783] | 1.01 | 3.86E-02 |
| P2ry6 | pyrimidinergic receptor P2Y, G-protein coupled, 6 [Source:MGI Symbol;Acc:MGI:2673874] | 1.42 | 3.86E-02 |
| Smg9 | SMG9 nonsense mediated mRNA decay factor [Source:MGI Symbol;Acc:MGI:1919247] | 0.70 | 3.86E-02 |
| Fermt3 | fermitin family member 3 [Source:MGI Symbol;Acc:MGI:2147790] | 1.20 | 3.86E-02 |
| Eef1akmt4 | EEF1A lysine methyltransferase 4 [Source:MGI Symbol;Acc:MGI:5903914] | 1.07 | 3.86E-02 |
| Gpn2 | GPN-loop GTPase 2 [Source:MGI Symbol;Acc:MGI:2140368] | 0.76 | 3.87E-02 |
| Pdlim4 | PDZ and LIM domain 4 [Source:MGI Symbol;Acc:MGI:1353470] | 0.96 | 3.88E-02 |
| B4galnt3 | beta-1,4-N-acetyl-galactosaminyl transferase 3 [Source:MGI Symbol;Acc:MGI:3041155] | 1.05 | 3.90E-02 |
| Trpc6 | transient receptor potential cation channel, subfamily C, member 6 [Source:MGI Symbol;Acc:MGI:109523] | 0.67 | 3.90E-02 |
| Ccr3 | C-C motif chemokine receptor 3 [Source:MGI Symbol;Acc:MGI:104616] | 1.92 | 3.90E-02 |
| Serpine2 | serine (or cysteine) peptidase inhibitor, clade E, member 2 [Source:MGI Symbol;Acc:MGI:101780] | 1.70 | 3.92E-02 |
| Plod3 | procollagen-lysine, 2-oxoglutarate 5-dioxygenase 3 [Source:MGI Symbol;Acc:MGI:1347008] | 1.07 | 3.92E-02 |
| Rcl1 | RNA terminal phosphate cyclase-like 1 [Source:MGI Symbol;Acc:MGI:1913275] | 0.70 | 3.92E-02 |
| Gstp3 | glutathione S-transferase pi 3 [Source:MGI Symbol;Acc:MGI:2385078] | 1.32 | 3.93E-02 |
| Lcp2 | lymphocyte cytosolic protein 2 [Source:MGI Symbol;Acc:MGI:1321402] | 1.72 | 3.95E-02 |

| | | | |
|---|---|---|---|
| Esm1 | endothelial cell-specific molecule 1 [Source:MGI Symbol;Acc:MGI:1918940] | 2.56 | 3.95E-02 |
| Plekhg4 | pleckstrin homology domain containing, family G (with RhoGef domain) member 4 [Source:MGI Symbol;Acc:MGI:2142544] | 2.98 | 3.96E-02 |
| Snrpb | small nuclear ribonucleoprotein B [Source:MGI Symbol;Acc:MGI:98342] | 0.78 | 3.99E-02 |
| St8sia2 | ST8 alpha-N-acetyl-neuraminide alpha-2,8-sialyltransferase 2 [Source:MGI Symbol;Acc:MGI:106020] | 1.42 | 3.99E-02 |
| Emp3 | epithelial membrane protein 3 [Source:MGI Symbol;Acc:MGI:1098729] | 1.61 | 3.99E-02 |
| Spi1 | Spi-1 proto-oncogene [Source:MGI Symbol;Acc:MGI:98282] | 1.32 | 3.99E-02 |
| Dok1 | docking protein 1 [Source:MGI Symbol;Acc:MGI:893587] | 1.28 | 4.00E-02 |
| Rpl15-ps6 | ribosomal protein L15, pseudogene 6 [Source:MGI Symbol;Acc:MGI:3642192] | 2.87 | 4.01E-02 |
| Mrpl17 | mitochondrial ribosomal protein L17 [Source:MGI Symbol;Acc:MGI:1351608] | 1.23 | 4.01E-02 |
| Bop1 | block of proliferation 1 [Source:MGI Symbol;Acc:MGI:1334460] | 1.07 | 4.01E-02 |
| Nt5c | 5',3'-nucleotidase, cytosolic [Source:MGI Symbol;Acc:MGI:1354954] | 0.86 | 4.01E-02 |
| Aars1 | alanyl-tRNA synthetase 1 [Source:MGI Symbol;Acc:MGI:2384560] | 0.62 | 4.01E-02 |
| Xxylt1 | xyloside xylosyltransferase 1 [Source:MGI Symbol;Acc:MGI:2146443] | 0.96 | 4.01E-02 |
| Dnlz | DNL-type zinc finger [Source:MGI Symbol;Acc:MGI:106559] | 0.80 | 4.01E-02 |
| Vmn2r32 | vomeronasal 2, receptor 32 [Source:MGI Symbol;Acc:MGI:1316696] | 6.31 | 4.05E-02 |
| Zdhhc12 | zinc finger, DHHC domain containing 12 [Source:MGI Symbol;Acc:MGI:1913470] | 1.16 | 4.06E-02 |
| Spic | Spi-C transcription factor (Spi-1/PU.1 related) [Source:MGI Symbol;Acc:MGI:1341168] | 0.82 | 4.07E-02 |
| Serpina3n | serine (or cysteine) peptidase inhibitor, clade A, member 3N [Source:MGI Symbol;Acc:MGI:105045] | 1.57 | 4.08E-02 |
| Mfap4 | microfibrillar-associated protein 4 [Source:MGI Symbol;Acc:MGI:1342276] | 1.16 | 4.08E-02 |
| Runx2 | runt related transcription factor 2 [Source:MGI Symbol;Acc:MGI:99829] | 1.05 | 4.09E-02 |
| Foxd2 | forkhead box D2 [Source:MGI Symbol;Acc:MGI:1347471] | 1.77 | 4.09E-02 |
| Rpl18 | ribosomal protein L18 [Source:MGI Symbol;Acc:MGI:98003] | 1.13 | 4.10E-02 |
| Sox13 | SRY (sex determining region Y)-box 13 [Source:MGI Symbol;Acc:MGI:98361] | 0.89 | 4.10E-02 |
| Gm18786 | predicted gene, 18786 [Source:MGI Symbol;Acc:MGI:5010971] | 3.95 | 4.11E-02 |
| Gm17455 | predicted gene, 17455 [Source:MGI Symbol;Acc:MGI:4937089] | 2.77 | 4.13E-02 |
| Neu1 | neuraminidase 1 [Source:MGI Symbol;Acc:MGI:97305] | 1.35 | 4.13E-02 |
| Itprip | inositol 1,4,5-triphosphate receptor interacting protein [Source:MGI Symbol;Acc:MGI:3042776] | 0.79 | 4.13E-02 |
| Serinc2 | serine incorporator 2 [Source:MGI Symbol;Acc:MGI:1919132] | 1.59 | 4.14E-02 |
| Fmnl1 | formin-like 1 [Source:MGI Symbol;Acc:MGI:1888994] | 1.39 | 4.14E-02 |
| Rpl34 | ribosomal protein L34 [Source:MGI Symbol;Acc:MGI:1915686] | 0.72 | 4.14E-02 |
| Crb2 | crumbs family member 2 [Source:MGI Symbol;Acc:MGI:2679260] | 1.52 | 4.15E-02 |
| Gm5641 | predicted gene 5641 [Source:MGI Symbol;Acc:MGI:3645731] | 1.00 | 4.15E-02 |
| Egfl7 | EGF-like domain 7 [Source:MGI Symbol;Acc:MGI:2449923] | 1.29 | 4.15E-02 |
| Pcbp1 | poly(rC) binding protein 1 [Source:MGI Symbol;Acc:MGI:1345635] | 0.75 | 4.15E-02 |
| Ifi206 | interferon activated gene 206 [Source:MGI Symbol;Acc:MGI:3646410] | 2.22 | 4.17E-02 |
| Itga1 | integrin alpha 1 [Source:MGI Symbol;Acc:MGI:96599] | 0.85 | 4.18E-02 |
| Mgat2 | mannoside acetylglucosaminyltransferase 2 [Source:MGI Symbol;Acc:MGI:2384966] | 0.73 | 4.18E-02 |
| Bmp2k | BMP2 inducible kinase [Source:MGI Symbol;Acc:MGI:2155456] | 0.65 | 4.18E-02 |
| Rpl28-ps1 | ribosomal protein L28, pseudogene 1 [Source:MGI Symbol;Acc:MGI:3705349] | 0.83 | 4.18E-02 |
| Slc38a5 | solute carrier family 38, member 5 [Source:MGI Symbol;Acc:MGI:2148066] | 2.48 | 4.20E-02 |
| Ly6d | lymphocyte antigen 6 family member D [Source:MGI Symbol;Acc:MGI:96881] | 1.19 | 4.21E-02 |
| Pxn | paxillin [Source:MGI Symbol;Acc:MGI:108295] | 0.81 | 4.22E-02 |
| Tbcb | tubulin folding cofactor B [Source:MGI Symbol;Acc:MGI:1913661] | 0.99 | 4.22E-02 |
| Slc26a9 | solute carrier family 26, member 9 [Source:MGI Symbol;Acc:MGI:2444594] | 1.23 | 4.24E-02 |
| Tldc2 | TBC/LysM associated domain containing 2 [Source:MGI Symbol;Acc:MGI:2686178] | 2.19 | 4.25E-02 |
| Fam110d | family with sequence similarity 110, member D [Source:MGI Symbol;Acc:MGI:1919940] | 1.52 | 4.25E-02 |
| Selenoh | selenoprotein H [Source:MGI Symbol;Acc:MGI:1919907] | 1.33 | 4.25E-02 |
| Rps9 | ribosomal protein S9 [Source:MGI Symbol;Acc:MGI:1924096] | 1.18 | 4.25E-02 |
| Shc1 | src homology 2 domain-containing transforming protein C1 [Source:MGI Symbol;Acc:MGI:98296] | 0.69 | 4.25E-02 |
| Gm9396 | predicted gene 9396 [Source:MGI Symbol;Acc:MGI:3645563] | 2.51 | 4.27E-02 |
| Nars1 | asparaginyl-tRNA synthetase 1 [Source:MGI Symbol;Acc:MGI:1917473] | 0.73 | 4.27E-02 |
| Ube2m | ubiquitin-conjugating enzyme E2M [Source:MGI Symbol;Acc:MGI:108278] | 0.60 | 4.27E-02 |
| Hsd11b2 | hydroxysteroid 11-beta dehydrogenase 2 [Source:MGI Symbol;Acc:MGI:104720] | 0.94 | 4.28E-02 |
| Snrpd2 | small nuclear ribonucleoprotein D2 [Source:MGI Symbol;Acc:MGI:98345] | 0.98 | 4.28E-02 |
| Riox1 | ribosomal oxygenase 1 [Source:MGI Symbol;Acc:MGI:1919202] | 1.18 | 4.29E-02 |
| Bdh2 | 3-hydroxybutyrate dehydrogenase, type 2 [Source:MGI Symbol;Acc:MGI:1917022] | 0.84 | 4.29E-02 |
| Pira1 | paired-Ig-like receptor A1 [Source:MGI Symbol;Acc:MGI:1195971] | 1.69 | 4.29E-02 |
| H2aj | H2J.A histone [Source:MGI Symbol;Acc:MGI:3606192] | 1.48 | 4.29E-02 |
| Gm17971 | predicted gene, 17971 [Source:MGI Symbol;Acc:MGI:5010156] | 1.15 | 4.29E-02 |
| Rrp12 | ribosomal RNA processing 12 homolog [Source:MGI Symbol;Acc:MGI:2147437] | 0.85 | 4.30E-02 |
| Dcakd | dephospho-CoA kinase domain containing [Source:MGI Symbol;Acc:MGI:1915337] | 0.97 | 4.31E-02 |
| Arrb2 | arrestin, beta 2 [Source:MGI Symbol;Acc:MGI:99474] | 1.08 | 4.32E-02 |
| Zyx | zyxin [Source:MGI Symbol;Acc:MGI:103072] | 1.04 | 4.32E-02 |
| Krt1 | keratin 1 [Source:MGI Symbol;Acc:MGI:96698] | 2.23 | 4.33E-02 |
| Ebi3 | Epstein-Barr virus induced gene 3 [Source:MGI Symbol;Acc:MGI:1354171] | 1.67 | 4.35E-02 |
| Lipg | lipase, endothelial [Source:MGI Symbol;Acc:MGI:1341803] | 1.14 | 4.35E-02 |
| Tor2a | torsin family 2, member A [Source:MGI Symbol;Acc:MGI:1353596] | 0.79 | 4.36E-02 |
| Kcnab2 | potassium voltage-gated channel, shaker-related subfamily, beta member 2 [Source:MGI Symbol;Acc:MGI:109239] | 1.13 | 4.39E-02 |
| Gm45855 | predicted gene 45855 [Source:MGI Symbol;Acc:MGI:5804970] | 1.83 | 4.41E-02 |
| Dact3 | dishevelled-binding antagonist of beta-catenin 3 [Source:MGI Symbol;Acc:MGI:3654828] | 1.26 | 4.41E-02 |
| Trappc1 | trafficking protein particle complex 1 [Source:MGI Symbol;Acc:MGI:1098727] | 0.64 | 4.43E-02 |

| | | | |
|---|---|---|---|
| Plk3 | polo like kinase 3 [Source:MGI Symbol;Acc:MGI:109604] | 1.43 | 4.43E-02 |
| Ercc1 | excision repair cross-complementing rodent repair deficiency, complementation group 1 [Source:MGI Symbol;Acc:MGI:95412] | 0.68 | 4.44E-02 |
| Impa2 | inositol monophosphatase 2 [Source:MGI Symbol;Acc:MGI:2149728] | 1.44 | 4.44E-02 |
| Hs3st1 | heparan sulfate (glucosamine) 3-O-sulfotransferase 1 [Source:MGI Symbol;Acc:MGI:1201606] | 1.13 | 4.44E-02 |
| Muc1 | mucin 1, transmembrane [Source:MGI Symbol;Acc:MGI:97231] | 1.38 | 4.46E-02 |
| Il10rb | interleukin 10 receptor, beta [Source:MGI Symbol;Acc:MGI:109380] | 0.92 | 4.46E-02 |
| Adcy4 | adenylate cyclase 4 [Source:MGI Symbol;Acc:MGI:99674] | 1.14 | 4.47E-02 |
| Synpo | synaptopodin [Source:MGI Symbol;Acc:MGI:1099446] | 1.32 | 4.48E-02 |
| H1f1 | H1.1 linker histone, cluster member [Source:MGI Symbol;Acc:MGI:1931523] | 2.15 | 4.49E-02 |
| Gm49909 | predicted gene, 49909 [Source:MGI Symbol;Acc:MGI:6270609] | 1.98 | 4.51E-02 |
| Plet1 | placenta expressed transcript 1 [Source:MGI Symbol;Acc:MGI:1923759] | 1.50 | 4.51E-02 |
| Plppr2 | phospholipid phosphatase related 2 [Source:MGI Symbol;Acc:MGI:2384575] | 0.96 | 4.51E-02 |
| Il17c | interleukin 17C [Source:MGI Symbol;Acc:MGI:2446486] | 2.03 | 4.51E-02 |
| Rusc2 | RUN and SH3 domain containing 2 [Source:MGI Symbol;Acc:MGI:2140371] | 0.82 | 4.52E-02 |
| Cd300ld5 | CD300 molecule like family member D5 [Source:MGI Symbol;Acc:MGI:3702661] | 1.70 | 4.52E-02 |
| Nid2 | nidogen 2 [Source:MGI Symbol;Acc:MGI:1298229] | 1.47 | 4.52E-02 |
| Atp2a2 | ATPase, Ca++ transporting, cardiac muscle, slow twitch 2 [Source:MGI Symbol;Acc:MGI:88110] | 0.71 | 4.52E-02 |
| Psmd3 | proteasome (prosome, macropain) 26S subunit, non-ATPase, 3 [Source:MGI Symbol;Acc:MGI:98858] | 0.78 | 4.53E-02 |
| Lyve1 | lymphatic vessel endothelial hyaluronan receptor 1 [Source:MGI Symbol;Acc:MGI:2136348] | 1.24 | 4.53E-02 |
| Gpsm1 | G-protein signalling modulator 1 (AGS3-like, C. elegans) [Source:MGI Symbol;Acc:MGI:1915089] | 1.11 | 4.53E-02 |
| Nkpd1 | NTPase, KAP family P-loop domain containing 1 [Source:MGI Symbol;Acc:MGI:1916797] | 1.85 | 4.54E-02 |
| Angptl4 | angiopoietin-like 4 [Source:MGI Symbol;Acc:MGI:1888999] | 1.93 | 4.55E-02 |
| Rpl39l | ribosomal protein L39-like [Source:MGI Symbol;Acc:MGI:1915422] | 1.74 | 4.58E-02 |
| F2r | coagulation factor II thrombin receptor [Source:MGI Symbol;Acc:MGI:101802] | 1.37 | 4.58E-02 |
| Stoml2 | stomatin (Epb7.2)-like 2 [Source:MGI Symbol;Acc:MGI:1913842] | 0.75 | 4.58E-02 |
| Tmem144 | transmembrane protein 144 [Source:MGI Symbol;Acc:MGI:1917902] | 0.75 | 4.59E-02 |
| Rrp7a | ribosomal RNA processing 7 homolog A [Source:MGI Symbol;Acc:MGI:1922028] | 0.64 | 4.60E-02 |
| Gm9761 | predicted gene 9761 [Source:MGI Symbol;Acc:MGI:3708640] | 3.89 | 4.61E-02 |
| Ido2 | indoleamine 2,3-dioxygenase 2 [Source:MGI Symbol;Acc:MGI:2142489] | 1.60 | 4.61E-02 |
| Poglut2 | protein O-glucosyltransferase 2 [Source:MGI Symbol;Acc:MGI:1919300] | 0.69 | 4.61E-02 |
| Tomm6 | translocase of outer mitochondrial membrane 6 [Source:MGI Symbol;Acc:MGI:1913369] | 0.82 | 4.64E-02 |
| Gm49528 | predicted gene, 49528 [Source:MGI Symbol;Acc:MGI:6155229] | 1.22 | 4.65E-02 |
| Fen1 | flap structure specific endonuclease 1 [Source:MGI Symbol;Acc:MGI:102779] | 1.22 | 4.67E-02 |
| 1700123O20Rik | RIKEN cDNA 1700123O20 gene [Source:MGI Symbol;Acc:MGI:1920893] | 0.92 | 4.67E-02 |
| Unc119b | unc-119 lipid binding chaperone B [Source:MGI Symbol;Acc:MGI:2147162] | 0.74 | 4.68E-02 |
| Hif1a | hypoxia inducible factor 1, alpha subunit [Source:MGI Symbol;Acc:MGI:106918] | 1.07 | 4.68E-02 |
| Ippk | inositol 1,3,4,5,6-pentakisphosphate 2-kinase [Source:MGI Symbol;Acc:MGI:1922928] | 0.84 | 4.70E-02 |
| Uqcrq | ubiquinol-cytochrome c reductase, complex III subunit VII [Source:MGI Symbol;Acc:MGI:107807] | 1.28 | 4.71E-02 |
| Gm11475 | predicted gene 11475 [Source:MGI Symbol;Acc:MGI:3652278] | 4.37 | 4.71E-02 |
| Grpr | gastrin releasing peptide receptor [Source:MGI Symbol;Acc:MGI:95836] | 2.38 | 4.71E-02 |
| Gpr68 | G protein-coupled receptor 68 [Source:MGI Symbol;Acc:MGI:2441763] | 0.90 | 4.71E-02 |
| Plod2 | procollagen lysine, 2-oxoglutarate 5-dioxygenase 2 [Source:MGI Symbol;Acc:MGI:1347007] | 1.47 | 4.72E-02 |
| Itgal | integrin alpha L [Source:MGI Symbol;Acc:MGI:96606] | 1.54 | 4.73E-02 |
| Siglece | sialic acid binding Ig-like lectin E [Source:MGI Symbol;Acc:MGI:1932475] | 1.32 | 4.76E-02 |
| Rnf223 | ring finger 223 [Source:MGI Symbol;Acc:MGI:3588193] | 1.36 | 4.76E-02 |
| Rap2a | RAS related protein 2a [Source:MGI Symbol;Acc:MGI:97855] | 0.75 | 4.77E-02 |
| Arhgap45 | Rho GTPase activating protein 45 [Source:MGI Symbol;Acc:MGI:1917969] | 1.27 | 4.81E-02 |
| F10 | coagulation factor X [Source:MGI Symbol;Acc:MGI:103107] | 1.19 | 4.81E-02 |
| Ntng2 | netrin G2 [Source:MGI Symbol;Acc:MGI:2159341] | 0.88 | 4.81E-02 |
| Mlkl | mixed lineage kinase domain-like [Source:MGI Symbol;Acc:MGI:1921818] | 0.90 | 4.82E-02 |
| Rps12 | ribosomal protein S12 [Source:MGI Symbol;Acc:MGI:98105] | 0.70 | 4.86E-02 |
| Ppia | peptidylprolyl isomerase A [Source:MGI Symbol;Acc:MGI:97749] | 0.80 | 4.87E-02 |
| Gm8307 | predicted gene 8307 [Source:MGI Symbol;Acc:MGI:3646941] | 2.99 | 4.87E-02 |
| Vwa1 | von Willebrand factor A domain containing 1 [Source:MGI Symbol;Acc:MGI:2179729] | 0.93 | 4.90E-02 |
| Ehd4 | EH-domain containing 4 [Source:MGI Symbol;Acc:MGI:1919619] | 0.78 | 4.90E-02 |
| Col8a1 | collagen, type VIII, alpha 1 [Source:MGI Symbol;Acc:MGI:88463] | 0.77 | 4.92E-02 |
| Rab13 | RAB13, member RAS oncogene family [Source:MGI Symbol;Acc:MGI:1927232] | 0.95 | 4.94E-02 |
| Rps27 | ribosomal protein S27 [Source:MGI Symbol;Acc:MGI:1888676] | 0.80 | 4.94E-02 |
| Rpl19 | ribosomal protein L19 [Source:MGI Symbol;Acc:MGI:98020] | 0.78 | 4.96E-02 |
| Cndp2 | CNDP dipeptidase 2 [Source:MGI Symbol;Acc:MGI:1913304] | 0.68 | 4.96E-02 |
| Adgre5 | adhesion G protein-coupled receptor E5 [Source:MGI Symbol;Acc:MGI:1347095] | 1.32 | 4.97E-02 |
| Lrrc47 | leucine rich repeat containing 47 [Source:MGI Symbol;Acc:MGI:1920196] | 1.05 | 4.98E-02 |
| Shmt2 | serine hydroxymethyltransferase 2 (mitochondrial) [Source:MGI Symbol;Acc:MGI:1277989] | 0.89 | 4.98E-02 |
| Pitx3 | paired-like homeodomain transcription factor 3 [Source:MGI Symbol;Acc:MGI:1100498] | 1.86 | 4.98E-02 |
| Ccl21a | C-C motif chemokine ligand 21 (serine) [Source:MGI Symbol;Acc:MGI:1349183] | 1.18 | 4.99E-02 |
| Gm6166 | predicted gene 6166 [Source:MGI Symbol;Acc:MGI:3645893] | 4.70 | 4.99E-02 |
| Gm15936 | predicted gene 15936 [Source:MGI Symbol;Acc:MGI:3802030] | 2.08 | 4.99E-02 |
| Tsen34 | tRNA splicing endonuclease subunit 34 [Source:MGI Symbol;Acc:MGI:1913328] | 1.24 | 4.99E-02 |
| Limd2 | LIM domain containing 2 [Source:MGI Symbol;Acc:MGI:1915053] | 0.90 | 4.99E-02 |
| Gm21742 | predicted gene, 21742 [Source:MGI Symbol;Acc:MGI:5433906] | 1.78 | 4.99E-02 |

#### Table S2. Downregulated genes in CONV-RT mice skin compared to CNT

| Gene ID | Name | log2(FC) | padj |
|---|---|---|---|
| Slc26a7 | solute carrier family 26, member 7 [Source:MGI Symbol;Acc:MGI:2384791] | -2.33 | 5.09E-19 |
| Pkib | protein kinase inhibitor beta, cAMP dependent, testis specific [Source:MGI Symbol;Acc:MGI:101937] | -2.15 | 6.26E-19 |
| Tnfaip8l3 | tumor necrosis factor, alpha-induced protein 8-like 3 [Source:MGI Symbol;Acc:MGI:2685363] | -2.59 | 1.25E-18 |
| Lep | leptin [Source:MGI Symbol;Acc:MGI:104663] | -4.00 | 7.87E-13 |
| Npr3 | natriuretic peptide receptor 3 [Source:MGI Symbol;Acc:MGI:97373] | -3.97 | 8.38E-13 |
| H2-M2 | histocompatibility 2, M region locus 2 [Source:MGI Symbol;Acc:MGI:95914] | -2.58 | 1.14E-12 |
| Abca5 | ATP-binding cassette, sub-family A member 5 [Source:MGI Symbol;Acc:MGI:2386607] | -2.35 | 2.22E-12 |
| Entpd2 | ectonucleoside triphosphate diphosphohydrolase 2 [Source:MGI Symbol;Acc:MGI:1096863] | -1.67 | 1.09E-11 |
| Gpha2 | glycoprotein hormone alpha 2 [Source:MGI Symbol;Acc:MGI:2156541] | -2.62 | 5.01E-10 |
| Il31ra | interleukin 31 receptor A [Source:MGI Symbol;Acc:MGI:2180511] | -2.41 | 5.04E-10 |
| Tgm7 | transglutaminase 7 [Source:MGI Symbol;Acc:MGI:2151164] | -4.21 | 1.88E-09 |
| Hlf | hepatic leukemia factor [Source:MGI Symbol;Acc:MGI:96108] | -2.18 | 2.90E-09 |
| Klhl14 | kelch-like 14 [Source:MGI Symbol;Acc:MGI:1921249] | -2.92 | 4.84E-09 |
| H2-Q5 | histocompatibility 2, Q region locus 5 [Source:MGI Symbol;Acc:MGI:95934] | -2.20 | 5.07E-09 |
| Il12rb2 | interleukin 12 receptor, beta 2 [Source:MGI Symbol;Acc:MGI:1270861] | -2.92 | 5.35E-09 |
| Clca3a1 | chloride channel accessory 3A1 [Source:MGI Symbol;Acc:MGI:1316732] | -1.94 | 9.03E-09 |
| Tox3 | TOX high mobility group box family member 3 [Source:MGI Symbol;Acc:MGI:3039593] | -2.81 | 1.68E-08 |
| Satb1 | special AT-rich sequence binding protein 1 [Source:MGI Symbol;Acc:MGI:105084] | -2.02 | 2.02E-08 |
| Cd209e | CD209e antigen [Source:MGI Symbol;Acc:MGI:2157948] | -2.48 | 2.37E-08 |
| Tef | thyrotroph embryonic factor [Source:MGI Symbol;Acc:MGI:98663] | -1.62 | 2.38E-08 |
| H2-Q6 | histocompatibility 2, Q region locus 6 [Source:MGI Symbol;Acc:MGI:95935] | -1.75 | 2.42E-08 |
| Ifnk | interferon kappa [Source:MGI Symbol;Acc:MGI:2683287] | -2.02 | 3.75E-08 |
| Tppp | tubulin polymerization promoting protein [Source:MGI Symbol;Acc:MGI:1920198] | -1.88 | 3.84E-08 |
| Usp18 | ubiquitin specific peptidase 18 [Source:MGI Symbol;Acc:MGI:1344364] | -1.49 | 5.64E-08 |
| Ptger3 | prostaglandin E receptor 3 (subtype EP3) [Source:MGI Symbol;Acc:MGI:97795] | -2.49 | 9.43E-08 |
| Lsamp | limbic system-associated membrane protein [Source:MGI Symbol;Acc:MGI:1261760] | -2.45 | 1.34E-07 |
| Cspg4b | chondroitin sulfate proteoglycan 4B [Source:MGI Symbol;Acc:MGI:3040697] | -1.45 | 1.45E-07 |
| Gm12294 | predicted gene 12294 [Source:MGI Symbol;Acc:MGI:3649526] | -2.80 | 1.98E-07 |
| H2-M5 | histocompatibility 2, M region locus 5 [Source:MGI Symbol;Acc:MGI:95917] | -2.57 | 2.11E-07 |
| Grik1 | glutamate receptor, ionotropic, kainate 1 [Source:MGI Symbol;Acc:MGI:95814] | -2.74 | 2.37E-07 |
| P2rx2 | purinergic receptor P2X, ligand-gated ion channel, 2 [Source:MGI Symbol;Acc:MGI:2665170] | -2.71 | 2.57E-07 |
| Sncg | synuclein, gamma [Source:MGI Symbol;Acc:MGI:1298397] | -2.64 | 2.63E-07 |
| Serpina3c | serine (or cysteine) peptidase inhibitor, clade A, member 3C [Source:MGI Symbol;Acc:MGI:102848] | -2.39 | 2.77E-07 |
| Plin1 | perilipin 1 [Source:MGI Symbol;Acc:MGI:1890505] | -3.02 | 5.25E-07 |
| Rspo3 | R-spondin 3 [Source:MGI Symbol;Acc:MGI:1920030] | -1.77 | 5.87E-07 |
| Scgb1a1 | secretoglobin, family 1A, member 1 [Source:MGI Symbol;Acc:MGI:98919] | -3.15 | 6.46E-07 |
| Slc6a2 | solute carrier family 6 (neurotransmitter transporter, noradrenalin), member 2 [Source:MGI Symbol;Acc:MGI:1270850] | -1.60 | 7.72E-07 |
| Gldc | glycine decarboxylase [Source:MGI Symbol;Acc:MGI:1341155] | -1.48 | 8.02E-07 |
| Irf4 | interferon regulatory factor 4 [Source:MGI Symbol;Acc:MGI:1096873] | -1.92 | 8.02E-07 |
| Stxbp6 | syntaxin binding protein 6 (amisyn) [Source:MGI Symbol;Acc:MGI:2384963] | -1.68 | 9.26E-07 |
| Skint7 | selection and upkeep of intraepithelial T cells 7 [Source:MGI Symbol;Acc:MGI:3041190] | -3.22 | 9.26E-07 |
| Arhgap44 | Rho GTPase activating protein 44 [Source:MGI Symbol;Acc:MGI:2144423] | -1.46 | 1.09E-06 |
| F830045P16Rik | RIKEN cDNA F830045P16 gene [Source:MGI Symbol;Acc:MGI:3045317] | -4.45 | 1.17E-06 |
| Prkag2 | protein kinase, AMP-activated, gamma 2 non-catalytic subunit [Source:MGI Symbol;Acc:MGI:1336153] | -1.24 | 1.41E-06 |
| Ell3 | elongation factor RNA polymerase II-like 3 [Source:MGI Symbol;Acc:MGI:2673679] | -2.08 | 1.46E-06 |
| Sectm1b | secreted and transmembrane 1B [Source:MGI Symbol;Acc:MGI:1929083] | -2.01 | 1.47E-06 |
| Efemp1 | epidermal growth factor-containing fibulin-like extracellular matrix protein 1 [Source:MGI Symbol;Acc:MGI:1339998] | -1.60 | 1.54E-06 |
| C7 | complement component 7 [Source:MGI Symbol;Acc:MGI:88235] | -3.18 | 1.60E-06 |
| Vipr1 | vasoactive intestinal peptide receptor 1 [Source:MGI Symbol;Acc:MGI:109272] | -1.80 | 1.87E-06 |
| Shisa7 | shisa family member 7 [Source:MGI Symbol;Acc:MGI:3605641] | -2.63 | 2.03E-06 |
| Xkr4 | X-linked Kx blood group related 4 [Source:MGI Symbol;Acc:MGI:3528744] | -2.05 | 2.26E-06 |
| A530016L24Rik | RIKEN cDNA A530016L24 gene [Source:MGI Symbol;Acc:MGI:2443020] | -2.64 | 2.50E-06 |
| Gm5127 | predicted gene 5127 [Source:MGI Symbol;Acc:MGI:3648285] | -2.26 | 3.36E-06 |
| Clxn | calaxin [Source:MGI Symbol;Acc:MGI:1914043] | -1.53 | 3.45E-06 |
| H2-Q7 | histocompatibility 2, Q region locus 7 [Source:MGI Symbol;Acc:MGI:95936] | -1.52 | 4.11E-06 |
| Pck1 | phosphoenolpyruvate carboxykinase 1, cytosolic [Source:MGI Symbol;Acc:MGI:97501] | -3.11 | 5.23E-06 |
| Ntf3 | neurotrophin 3 [Source:MGI Symbol;Acc:MGI:97380] | -1.84 | 5.28E-06 |
| Enpp5 | ectonucleotide pyrophosphatase/phosphodiesterase 5 [Source:MGI Symbol;Acc:MGI:1933830] | -1.96 | 5.43E-06 |
| Smoc1 | SPARC related modular calcium binding 1 [Source:MGI Symbol;Acc:MGI:1929878] | -2.07 | 5.51E-06 |
| Kel | Kell blood group [Source:MGI Symbol;Acc:MGI:1346053] | -7.26 | 5.51E-06 |
| Cd55 | CD55 molecule, decay accelerating factor for complement [Source:MGI Symbol;Acc:MGI:104850] | -1.78 | 6.10E-06 |
| Egfl6 | EGF-like-domain, multiple 6 [Source:MGI Symbol;Acc:MGI:1858599] | -1.23 | 6.28E-06 |
| Mill2 | MHC I like leukocyte 2 [Source:MGI Symbol;Acc:MGI:2179989] | -1.44 | 6.69E-06 |
| Cdh4 | cadherin 4 [Source:MGI Symbol;Acc:MGI:99218] | -2.00 | 6.83E-06 |

| | | | |
|---|---|---|---|
| Lrrn3 | leucine rich repeat protein 3, neuronal [Source:MGI Symbol;Acc:MGI:106036] | -1.59 | 7.50E-06 |
| Cd59a | CD59a antigen [Source:MGI Symbol;Acc:MGI:109177] | -1.97 | 7.68E-06 |
| Gm17043 | predicted gene 17043 [Source:MGI Symbol;Acc:MGI:4937870] | -4.34 | 7.70E-06 |
| Rasl10a | RAS-like, family 10, member A [Source:MGI Symbol;Acc:MGI:1922918] | -1.42 | 7.87E-06 |
| Cd3e | CD3 antigen, epsilon polypeptide [Source:MGI Symbol;Acc:MGI:88332] | -1.49 | 8.02E-06 |
| Ccdc88c | coiled-coil domain containing 88C [Source:MGI Symbol;Acc:MGI:1915589] | -1.23 | 8.29E-06 |
| Adcy1 | adenylate cyclase 1 [Source:MGI Symbol;Acc:MGI:99677] | -2.16 | 1.03E-05 |
| Gm45713 | predicted gene 45713 [Source:MGI Symbol;Acc:MGI:5804828] | -2.55 | 1.19E-05 |
| Acvr1c | activin A receptor, type IC [Source:MGI Symbol;Acc:MGI:2661081] | -2.69 | 1.35E-05 |
| Prr32 | proline rich 32 [Source:MGI Symbol;Acc:MGI:1916050] | -7.09 | 1.38E-05 |
| Igfbp3 | insulin-like growth factor binding protein 3 [Source:MGI Symbol;Acc:MGI:96438] | -1.12 | 1.46E-05 |
| Kcnj16 | potassium inwardly-rectifying channel, subfamily J, member 16 [Source:MGI Symbol;Acc:MGI:1314842] | -2.26 | 1.46E-05 |
| Klhdc1 | kelch domain containing 1 [Source:MGI Symbol;Acc:MGI:2672853] | -1.80 | 1.50E-05 |
| Cutal | cutA divalent cation tolerance homolog-like [Source:MGI Symbol;Acc:MGI:1925246] | -2.26 | 1.55E-05 |
| Nr1d1 | nuclear receptor subfamily 1, group D, member 1 [Source:MGI Symbol;Acc:MGI:2444210] | -1.77 | 1.56E-05 |
| Mgl2 | macrophage galactose N-acetyl-galactosamine specific lectin 2 [Source:MGI Symbol;Acc:MGI:2385729] | -2.23 | 1.62E-05 |
| Cdh18 | cadherin 18 [Source:MGI Symbol;Acc:MGI:1344366] | -3.74 | 1.74E-05 |
| Nrcam | neuronal cell adhesion molecule [Source:MGI Symbol;Acc:MGI:104750] | -1.39 | 1.76E-05 |
| Fry | FRY microtubule binding protein [Source:MGI Symbol;Acc:MGI:2443895] | -1.34 | 1.84E-05 |
| Cyp4b1 | cytochrome P450, family 4, subfamily b, polypeptide 1 [Source:MGI Symbol;Acc:MGI:103225] | -1.34 | 1.94E-05 |
| Nlrp1c-ps | NLR family, pyrin domain containing 1C, pseudogene [Source:NCBI gene (formerly Entrezgene);Acc:627984] | -1.61 | 2.30E-05 |
| Nlrp1c-ps | NLR family, pyrin domain containing 1C, pseudogene [Source:MGI Symbol;Acc:MGI:3582962] | -1.61 | 2.30E-05 |
| Plscr4 | phospholipid scramblase 4 [Source:MGI Symbol;Acc:MGI:2143267] | -1.60 | 2.55E-05 |
| Hsf2 | heat shock factor 2 [Source:MGI Symbol;Acc:MGI:96239] | -1.35 | 2.78E-05 |
| Serpina3h | serine (or cysteine) peptidase inhibitor, clade A, member 3H [Source:MGI Symbol;Acc:MGI:2182839] | -2.26 | 3.00E-05 |
| Serpina3h | serine (or cysteine) peptidase inhibitor, clade A, member 3H [Source:NCBI gene (formerly Entrezgene);Acc:546546] | -2.26 | 3.00E-05 |
| Ccdc3 | coiled-coil domain containing 3 [Source:MGI Symbol;Acc:MGI:1921436] | -1.86 | 3.04E-05 |
| Scg3 | secretogranin III [Source:MGI Symbol;Acc:MGI:103032] | -1.73 | 3.11E-05 |
| Rab27a | RAB27A, member RAS oncogene family [Source:MGI Symbol;Acc:MGI:1861441] | -1.05 | 3.23E-05 |
| Opcml | opioid binding protein/cell adhesion molecule-like [Source:MGI Symbol;Acc:MGI:97397] | -1.52 | 3.37E-05 |
| Cyp2b23 | cytochrome P450, family 2, subfamily b, polypeptide 23 [Source:MGI Symbol;Acc:MGI:3646735] | -2.80 | 3.45E-05 |
| Trarg1 | trafficking regulator of GLUT4 (SLC2A4) 1 [Source:MGI Symbol;Acc:MGI:3029307] | -2.15 | 4.10E-05 |
| Rbp4 | retinol binding protein 4, plasma [Source:MGI Symbol;Acc:MGI:97879] | -2.98 | 4.34E-05 |
| Cd96 | CD96 antigen [Source:MGI Symbol;Acc:MGI:1934368] | -2.43 | 4.53E-05 |
| Micu1 | mitochondrial calcium uptake 1 [Source:MGI Symbol;Acc:MGI:2384909] | -1.50 | 4.86E-05 |
| Arrdc3 | arrestin domain containing 3 [Source:MGI Symbol;Acc:MGI:2145242] | -0.84 | 5.20E-05 |
| Cachd1 | cache domain containing 1 [Source:MGI Symbol;Acc:MGI:2444177] | -1.70 | 5.76E-05 |
| AW146154 | expressed sequence AW146154 [Source:MGI Symbol;Acc:MGI:2142212] | -1.39 | 5.81E-05 |
| Bche | butyrylcholinesterase [Source:MGI Symbol;Acc:MGI:894278] | -1.89 | 6.17E-05 |
| Trim50 | tripartite motif-containing 50 [Source:MGI Symbol;Acc:MGI:2664992] | -1.55 | 6.87E-05 |
| Sh3d21 | SH3 domain containing 21 [Source:MGI Symbol;Acc:MGI:1914188] | -1.49 | 7.05E-05 |
| Ark2c | arkadia (RNF111) C-terminal like ring finger ubiquitin ligase 2C [Source:MGI Symbol;Acc:MGI:2444521] | -1.38 | 7.28E-05 |
| Nectin3 | nectin cell adhesion molecule 3 [Source:MGI Symbol;Acc:MGI:1930171] | -1.37 | 7.47E-05 |
| Mrap | melanocortin 2 receptor accessory protein [Source:MGI Symbol;Acc:MGI:1924287] | -2.67 | 7.60E-05 |
| Arhgap18 | Rho GTPase activating protein 18 [Source:MGI Symbol;Acc:MGI:1921160] | -1.12 | 7.80E-05 |
| Upk3b | uroplakin 3B [Source:MGI Symbol;Acc:MGI:2140882] | -3.40 | 8.22E-05 |
| Adgre4 | adhesion G protein-coupled receptor E4 [Source:MGI Symbol;Acc:MGI:1196464] | -1.99 | 8.25E-05 |
| Gm44289 | predicted gene, 44289 [Source:MGI Symbol;Acc:MGI:5690681] | -2.85 | 8.31E-05 |
| Il18r1 | interleukin 18 receptor 1 [Source:MGI Symbol;Acc:MGI:105383] | -1.21 | 8.99E-05 |
| Fgf18 | fibroblast growth factor 18 [Source:MGI Symbol;Acc:MGI:1277980] | -2.43 | 9.41E-05 |
| Ccl27al | C-C motif chemokine ligand 27A like [Source:MGI Symbol;Acc:MGI:3713752] | -2.12 | 9.89E-05 |
| Adgrg2 | adhesion G protein-coupled receptor G2 [Source:MGI Symbol;Acc:MGI:2446854] | -1.56 | 1.02E-04 |
| Il11ra2 | interleukin 11 receptor subunit alpha 2 [Source:MGI Symbol;Acc:MGI:109123] | -2.33 | 1.09E-04 |
| Ptpn21 | protein tyrosine phosphatase, non-receptor type 21 [Source:MGI Symbol;Acc:MGI:1344406] | -1.61 | 1.09E-04 |
| Cttnbp2 | cortactin binding protein 2 [Source:MGI Symbol;Acc:MGI:1353467] | -1.87 | 1.13E-04 |
| Cyp2j9 | cytochrome P450, family 2, subfamily j, polypeptide 9 [Source:MGI Symbol;Acc:MGI:1921769] | -1.79 | 1.17E-04 |
| Vps13a | vacuolar protein sorting 13A [Source:MGI Symbol;Acc:MGI:2444304] | -1.17 | 1.22E-04 |
| Cry2 | cryptochrome circadian regulator 2 [Source:MGI Symbol;Acc:MGI:1270859] | -1.08 | 1.23E-04 |
| Slc4a8 | solute carrier family 4 (anion exchanger), member 8 [Source:MGI Symbol;Acc:MGI:1928745] | -1.58 | 1.24E-04 |
| Rab11fip4 | RAB11 family interacting protein 4 (class II) [Source:MGI Symbol;Acc:MGI:2442920] | -1.81 | 1.25E-04 |
| Or10g6 | olfactory receptor family 10 subfamily G member 6 [Source:MGI Symbol;Acc:MGI:3030815] | -1.56 | 1.26E-04 |
| Aoc3 | amine oxidase, copper containing 3 [Source:MGI Symbol;Acc:MGI:1306797] | -2.10 | 1.28E-04 |
| Lypd1 | Ly6/Plaur domain containing 1 [Source:MGI Symbol;Acc:MGI:1919835] | -1.47 | 1.37E-04 |
| Gm15519 | predicted gene 15519 [Source:MGI Symbol;Acc:MGI:3782965] | -3.64 | 1.37E-04 |
| Pdzd7 | PDZ domain containing 7 [Source:MGI Symbol;Acc:MGI:3608325] | -1.08 | 1.38E-04 |
| Ifitm6 | interferon induced transmembrane protein 6 [Source:MGI Symbol;Acc:MGI:2686976] | -1.92 | 1.47E-04 |
| Zc2hc1a | zinc finger, C2HC-type containing 1A [Source:MGI Symbol;Acc:MGI:1914556] | -1.28 | 1.52E-04 |
| Wipf3 | WAS/WASL interacting protein family, member 3 [Source:MGI Symbol;Acc:MGI:3044681] | -1.63 | 1.61E-04 |
| Sult5a1 | sulfotransferase family 5A, member 1 [Source:MGI Symbol;Acc:MGI:1931463] | -2.10 | 1.64E-04 |
| Rab6b | RAB6B, member RAS oncogene family [Source:MGI Symbol;Acc:MGI:107283] | -1.12 | 1.65E-04 |
| Abat | 4-aminobutyrate aminotransferase [Source:MGI Symbol;Acc:MGI:2443582] | -1.07 | 1.68E-04 |

| | | | |
|---|---|---|---|
| Kctd12 | potassium channel tetramerisation domain containing 12 [Source:MGI Symbol;Acc:MGI:2145823] | -1.17 | 1.68E-04 |
| Pdzrn4 | PDZ domain containing RING finger 4 [Source:MGI Symbol;Acc:MGI:3056996] | -2.12 | 1.72E-04 |
| Lama1 | laminin, alpha 1 [Source:MGI Symbol;Acc:MGI:99892] | -1.85 | 1.72E-04 |
| Oxtr | oxytocin receptor [Source:MGI Symbol;Acc:MGI:109147] | -1.60 | 1.77E-04 |
| Apol7a | apolipoprotein L 7a [Source:MGI Symbol;Acc:MGI:1923011] | -1.74 | 1.77E-04 |
| Myocd | myocardin [Source:MGI Symbol;Acc:MGI:2137495] | -1.40 | 2.04E-04 |
| Cdon | cell adhesion molecule-related/down-regulated by oncogenes [Source:MGI Symbol;Acc:MGI:1926387] | -1.19 | 2.09E-04 |
| Slc6a4 | solute carrier family 6 (neurotransmitter transporter, serotonin), member 4 [Source:MGI Symbol;Acc:MGI:96285] | -1.63 | 2.09E-04 |
| Il11ra3 | interleukin 11 receptor subunit alpha 3 [Source:MGI Symbol;Acc:MGI:3801997] | -2.35 | 2.09E-04 |
| Cyp4a10 | cytochrome P450, family 4, subfamily a, polypeptide 10 [Source:MGI Symbol;Acc:MGI:88611] | -2.56 | 2.09E-04 |
| Fsip1 | fibrous sheath-interacting protein 1 [Source:MGI Symbol;Acc:MGI:1918563] | -1.77 | 2.15E-04 |
| Hyal5 | hyaluronoglucosaminidase 5 [Source:MGI Symbol;Acc:MGI:1921718] | -3.42 | 2.23E-04 |
| Trhde | TRH-degrading enzyme [Source:MGI Symbol;Acc:MGI:2384311] | -2.63 | 2.30E-04 |
| Zfp467 | zinc finger protein 467 [Source:MGI Symbol;Acc:MGI:1916160] | -1.18 | 2.48E-04 |
| Utrn | utrophin [Source:MGI Symbol;Acc:MGI:104631] | -0.83 | 2.62E-04 |
| Gm5141 | predicted gene 5141 [Source:MGI Symbol;Acc:MGI:3779466] | -1.71 | 2.67E-04 |
| Bmp3 | bone morphogenetic protein 3 [Source:MGI Symbol;Acc:MGI:88179] | -1.32 | 2.71E-04 |
| Rev3l | REV3 like, DNA directed polymerase zeta catalytic subunit [Source:MGI Symbol;Acc:MGI:1337131] | -0.85 | 2.90E-04 |
| Gm7399 | predicted gene 7399 [Source:MGI Symbol;Acc:MGI:3647495] | -1.75 | 2.90E-04 |
| Amy1 | amylase 1, salivary [Source:MGI Symbol;Acc:MGI:88019] | -2.66 | 3.01E-04 |
| Lrrc31 | leucine rich repeat containing 31 [Source:MGI Symbol;Acc:MGI:2443864] | -1.73 | 3.21E-04 |
| Tex47 | testis expressed 47 [Source:MGI Symbol;Acc:MGI:1918170] | -4.00 | 3.59E-04 |
| Art4 | ADP-ribosyltransferase 4 [Source:MGI Symbol;Acc:MGI:1202710] | -1.63 | 3.62E-04 |
| Timp4 | tissue inhibitor of metalloproteinase 4 [Source:MGI Symbol;Acc:MGI:109125] | -1.96 | 3.81E-04 |
| Ackr4 | atypical chemokine receptor 4 [Source:MGI Symbol;Acc:MGI:2181676] | -1.67 | 3.85E-04 |
| Sema3c | sema domain, immunoglobulin domain (Ig), short basic domain, secreted, (semaphorin) 3C [Source:MGI Symbol;Acc:MGI:107557] | -1.68 | 3.98E-04 |
| Adrb3 | adrenergic receptor, beta 3 [Source:MGI Symbol;Acc:MGI:87939] | -1.54 | 4.01E-04 |
| D3Ertd751e | DNA segment, Chr 3, ERATO Doi 751, expressed [Source:MGI Symbol;Acc:MGI:1289213] | -1.33 | 4.01E-04 |
| Mill1 | MHC I like leukocyte 1 [Source:MGI Symbol;Acc:MGI:2179988] | -1.62 | 4.01E-04 |
| Pak3 | p21 (RAC1) activated kinase 3 [Source:MGI Symbol;Acc:MGI:1339656] | -1.45 | 4.04E-04 |
| Tlr11 | toll-like receptor 11 [Source:MGI Symbol;Acc:MGI:3045226] | -2.04 | 4.16E-04 |
| Vmn1r214 | vomeronasal 1 receptor 214 [Source:MGI Symbol;Acc:MGI:2159663] | -1.41 | 4.18E-04 |
| Ccdc171 | coiled-coil domain containing 171 [Source:MGI Symbol;Acc:MGI:1922152] | -0.98 | 4.19E-04 |
| Slc2a12 | solute carrier family 2 (facilitated glucose transporter), member 12 [Source:MGI Symbol;Acc:MGI:3052471] | -2.27 | 4.22E-04 |
| Phactr2 | phosphatase and actin regulator 2 [Source:MGI Symbol;Acc:MGI:2446138] | -1.06 | 4.23E-04 |
| Plcxd3 | phosphatidylinositol-specific phospholipase C, X domain containing 3 [Source:MGI Symbol;Acc:MGI:2442605] | -2.79 | 4.24E-04 |
| Prrg3 | proline rich Gla (G-carboxyglutamic acid) 3 (transmembrane) [Source:MGI Symbol;Acc:MGI:2685214] | -1.38 | 4.36E-04 |
| Slc36a2 | solute carrier family 36 (proton/amino acid symporter), member 2 [Source:MGI Symbol;Acc:MGI:1891430] | -1.76 | 4.52E-04 |
| Hmgcll1 | 3-hydroxymethyl-3-methylglutaryl-Coenzyme A lyase-like 1 [Source:MGI Symbol;Acc:MGI:2446108] | -1.56 | 4.69E-04 |
| Slc24a3 | solute carrier family 24 (sodium/potassium/calcium exchanger), member 3 [Source:MGI Symbol;Acc:MGI:2137513] | -2.03 | 4.71E-04 |
| Hsd3b2 | hydroxy-delta-5-steroid dehydrogenase, 3 beta- and steroid delta-isomerase 2 [Source:MGI Symbol;Acc:MGI:96234] | -1.21 | 4.72E-04 |
| Pon1 | paraoxonase 1 [Source:MGI Symbol;Acc:MGI:103295] | -3.75 | 4.72E-04 |
| Smoc2 | SPARC related modular calcium binding 2 [Source:MGI Symbol;Acc:MGI:1929881] | -0.87 | 5.03E-04 |
| Cfap96 | cilia and flagella associated protein 96 [Source:MGI Symbol;Acc:MGI:1916729] | -1.35 | 5.10E-04 |
| Sema3d | sema domain, immunoglobulin domain (Ig), short basic domain, secreted, (semaphorin) 3D [Source:MGI Symbol;Acc:MGI:1860118] | -2.21 | 5.42E-04 |
| Rbbp8 | retinoblastoma binding protein 8, endonuclease [Source:MGI Symbol;Acc:MGI:2442995] | -0.97 | 5.42E-04 |
| Kif21a | kinesin family member 21A [Source:MGI Symbol;Acc:MGI:109188] | -1.70 | 5.42E-04 |
| Gm49545 | predicted gene, 49545 [Source:MGI Symbol;Acc:MGI:6155252] | -1.97 | 5.42E-04 |
| Chrnb4 | cholinergic receptor, nicotinic, beta polypeptide 4 [Source:MGI Symbol;Acc:MGI:87892] | -2.57 | 5.43E-04 |
| Sec14l4 | SEC14-like lipid binding 4 [Source:MGI Symbol;Acc:MGI:2144095] | -0.78 | 5.47E-04 |
| Ipmk | inositol polyphosphate multikinase [Source:MGI Symbol;Acc:MGI:1916968] | -0.92 | 5.53E-04 |
| Ddb2 | damage specific DNA binding protein 2 [Source:MGI Symbol;Acc:MGI:1355314] | -1.01 | 5.59E-04 |
| Defb11 | defensin beta 11 [Source:MGI Symbol;Acc:MGI:2179197] | -6.30 | 5.61E-04 |
| Aff3 | AF4/FMR2 family, member 3 [Source:MGI Symbol;Acc:MGI:106927] | -1.55 | 5.79E-04 |
| Gm17743 | predicted gene, 17743 [Source:MGI Symbol;Acc:MGI:5009819] | -2.33 | 5.87E-04 |
| Apol6 | apolipoprotein L 6 [Source:MGI Symbol;Acc:MGI:1919189] | -1.94 | 6.15E-04 |
| Foxn3 | forkhead box N3 [Source:MGI Symbol;Acc:MGI:1918625] | -1.00 | 6.38E-04 |
| Pramel57 | PRAME like 57 [Source:MGI Symbol;Acc:MGI:3781464] | -3.61 | 6.43E-04 |
| Ablim3 | actin binding LIM protein family, member 3 [Source:MGI Symbol;Acc:MGI:2442582] | -1.55 | 6.49E-04 |
| Bbox1 | gamma-butyrobetaine hydroxylase 1 [Source:MGI Symbol;Acc:MGI:1891372] | -2.34 | 6.56E-04 |
| Dach2 | dachshund family transcription factor 2 [Source:MGI Symbol;Acc:MGI:1890446] | -2.00 | 6.69E-04 |
| Bcl2 | B cell leukemia/lymphoma 2 [Source:MGI Symbol;Acc:MGI:88138] | -0.86 | 6.89E-04 |
| Clasp2 | CLIP associating protein 2 [Source:MGI Symbol;Acc:MGI:1923749] | -0.85 | 6.93E-04 |
| Fn3krp | fructosamine 3 kinase related protein [Source:MGI Symbol;Acc:MGI:2679256] | -0.90 | 7.18E-04 |
| Atp6v0e2 | ATPase, H+ transporting, lysosomal V0 subunit E2 [Source:MGI Symbol;Acc:MGI:1923502] | -1.56 | 7.19E-04 |
| Ccl27b | C-C motif chemokine ligand 27b [Source:MGI Symbol;Acc:MGI:1891389] | -2.97 | 7.19E-04 |

| | | | |
|---|---|---|---|
| Gm973 | predicted gene 973 [Source:MGI Symbol;Acc:MGI:2685819] | -1.57 | 7.31E-04 |
| Vstm2a | V-set and transmembrane domain containing 2A [Source:MGI Symbol;Acc:MGI:2384826] | -2.87 | 7.45E-04 |
| 1700025G04Rik | RIKEN cDNA 1700025G04 gene [Source:MGI Symbol;Acc:MGI:1916649] | -1.05 | 7.80E-04 |
| Gm5949 | predicted gene 5949 [Source:MGI Symbol;Acc:MGI:3645374] | -2.80 | 8.27E-04 |
| Ppara | peroxisome proliferator activated receptor alpha [Source:MGI Symbol;Acc:MGI:104740] | -1.30 | 8.52E-04 |
| Plin4 | perilipin 4 [Source:MGI Symbol;Acc:MGI:1929709] | -2.11 | 8.73E-04 |
| Gm48892 | predicted gene, 48892 [Source:MGI Symbol;Acc:MGI:6098655] | -3.85 | 9.64E-04 |
| Abca6 | ATP-binding cassette, sub-family A member 6 [Source:MGI Symbol;Acc:MGI:1923434] | -2.01 | 9.75E-04 |
| Glt8d2 | glycosyltransferase 8 domain containing 2 [Source:MGI Symbol;Acc:MGI:1922032] | -1.39 | 1.01E-03 |
| Crebrf | CREB3 regulatory factor [Source:MGI Symbol;Acc:MGI:1924378] | -1.28 | 1.02E-03 |
| Akap17b | A kinase anchor protein 17B [Source:MGI Symbol;Acc:MGI:2443758] | -0.97 | 1.02E-03 |
| Prdm9 | PR domain containing 9 [Source:MGI Symbol;Acc:MGI:2384854] | -1.04 | 1.04E-03 |
| Col4a4 | collagen, type IV, alpha 4 [Source:MGI Symbol;Acc:MGI:104687] | -1.28 | 1.08E-03 |
| Abcd2 | ATP-binding cassette, sub-family D member 2 [Source:MGI Symbol;Acc:MGI:1349467] | -2.36 | 1.09E-03 |
| Clca1 | chloride channel accessory 1 [Source:MGI Symbol;Acc:MGI:1346342] | -2.99 | 1.14E-03 |
| Slc26a8 | solute carrier family 26, member 8 [Source:MGI Symbol;Acc:MGI:2385046] | -1.27 | 1.14E-03 |
| Crim1 | cysteine rich transmembrane BMP regulator 1 [Source:MGI Symbol;Acc:MGI:1354756] | -0.84 | 1.15E-03 |
| Vmn1r28 | vomeronasal 1 receptor 28 [Source:MGI Symbol;Acc:MGI:2159461] | -1.41 | 1.15E-03 |
| Tcaf2 | TRPM8 channel-associated factor 2 [Source:MGI Symbol;Acc:MGI:2385258] | -1.08 | 1.16E-03 |
| Scin | scinderin [Source:MGI Symbol;Acc:MGI:1306794] | -1.43 | 1.23E-03 |
| Or51g1 | olfactory receptor family 51 subfamily G member 1 [Source:MGI Symbol;Acc:MGI:3030412] | -1.32 | 1.26E-03 |
| Smad9 | SMAD family member 9 [Source:MGI Symbol;Acc:MGI:1859993] | -1.30 | 1.27E-03 |
| Ighd | immunoglobulin heavy constant delta [Source:MGI Symbol;Acc:MGI:96447] | -2.89 | 1.29E-03 |
| Zfp652 | zinc finger protein 652 [Source:MGI Symbol;Acc:MGI:2442221] | -1.28 | 1.35E-03 |
| Cidec | cell death-inducing DFFA-like effector c [Source:MGI Symbol;Acc:MGI:95585] | -2.36 | 1.40E-03 |
| Col8a2 | collagen, type VIII, alpha 2 [Source:MGI Symbol;Acc:MGI:88464] | -1.23 | 1.44E-03 |
| Pecr | peroxisomal trans-2-enoyl-CoA reductase [Source:MGI Symbol;Acc:MGI:2148199] | -1.15 | 1.52E-03 |
| Krt7 | keratin 7 [Source:MGI Symbol;Acc:MGI:96704] | -1.17 | 1.53E-03 |
| Slc22a3 | solute carrier family 22 (organic cation transporter), member 3 [Source:MGI Symbol;Acc:MGI:1333817] | -1.68 | 1.53E-03 |
| Upb1 | ureidopropionase, beta [Source:MGI Symbol;Acc:MGI:2143535] | -1.16 | 1.56E-03 |
| Klhdc7a | kelch domain containing 7A [Source:MGI Symbol;Acc:MGI:2444612] | -1.46 | 1.57E-03 |
| Btc | betacellulin, epidermal growth factor family member [Source:MGI Symbol;Acc:MGI:99439] | -1.65 | 1.58E-03 |
| Inmt | indolethylamine N-methyltransferase [Source:MGI Symbol;Acc:MGI:102963] | -3.45 | 1.59E-03 |
| BC064078 | cDNA sequence BC064078 [Source:MGI Symbol;Acc:MGI:3040692] | -1.81 | 1.65E-03 |
| Sorcs3 | sortilin-related VPS10 domain containing receptor 3 [Source:MGI Symbol;Acc:MGI:1913923] | -2.26 | 1.68E-03 |
| Or14p1 | olfactory receptor family 14 subfamily P member 1 [Source:MGI Symbol;Acc:MGI:3030299] | -1.22 | 1.68E-03 |
| Elavl2 | ELAV like RNA binding protein 1 [Source:MGI Symbol;Acc:MGI:1100887] | -3.32 | 1.69E-03 |
| Ccr4 | C-C motif chemokine receptor 4 [Source:MGI Symbol;Acc:MGI:107824] | -1.49 | 1.69E-03 |
| Mef2a | myocyte enhancer factor 2A [Source:MGI Symbol;Acc:MGI:99532] | -1.05 | 1.70E-03 |
| Enpep | glutamyl aminopeptidase [Source:MGI Symbol;Acc:MGI:106645] | -2.02 | 1.74E-03 |
| Itpkb | inositol 1,4,5-trisphosphate 3-kinase B [Source:MGI Symbol;Acc:MGI:109235] | -0.68 | 1.77E-03 |
| Ccl27a | C-C motif chemokine ligand 27A [Source:MGI Symbol;Acc:MGI:1343459] | -1.92 | 1.80E-03 |
| Hspa12a | heat shock protein 12A [Source:MGI Symbol;Acc:MGI:1920692] | -1.06 | 1.85E-03 |
| Cd209a | CD209a antigen [Source:MGI Symbol;Acc:MGI:2157942] | -1.51 | 1.88E-03 |
| Pik3c2g | phosphatidylinositol-4-phosphate 3-kinase catalytic subunit type 2 gamma [Source:MGI Symbol;Acc:MGI:1203730] | -2.24 | 1.89E-03 |
| Mapkbp1 | mitogen-activated protein kinase binding protein 1 [Source:MGI Symbol;Acc:MGI:1347004] | -0.86 | 1.90E-03 |
| Ptprd | protein tyrosine phosphatase receptor type D [Source:MGI Symbol;Acc:MGI:97812] | -1.09 | 1.90E-03 |
| Dab1 | disabled 1 [Source:MGI Symbol;Acc:MGI:108554] | -2.44 | 1.90E-03 |
| Tdrp | testis development related protein [Source:MGI Symbol;Acc:MGI:1919398] | -1.45 | 1.90E-03 |
| Pnpla3 | patatin-like phospholipase domain containing 3 [Source:MGI Symbol;Acc:MGI:2151796] | -1.70 | 1.90E-03 |
| Ngfr | nerve growth factor receptor (TNFR superfamily, member 16) [Source:MGI Symbol;Acc:MGI:97323] | -0.94 | 1.92E-03 |
| Lpl | lipoprotein lipase [Source:MGI Symbol;Acc:MGI:96820] | -1.95 | 1.92E-03 |
| B3galt2 | UDP-Gal:betaGlcNAc beta 1,3-galactosyltransferase, polypeptide 2 [Source:MGI Symbol;Acc:MGI:1349461] | -0.99 | 1.93E-03 |
| Clca3a2 | chloride channel accessory 3A2 [Source:MGI Symbol;Acc:MGI:1931471] | -2.34 | 1.95E-03 |
| Mpc1-ps | mitochondrial pyruvate carrier 1, pseudogene [Source:MGI Symbol;Acc:MGI:3781628] | -6.22 | 1.95E-03 |
| Glb1l2 | galactosidase, beta 1-like 2 [Source:MGI Symbol;Acc:MGI:2388283] | -1.59 | 1.96E-03 |
| Slc5a7 | solute carrier family 5 (choline transporter), member 7 [Source:MGI Symbol;Acc:MGI:1927126] | -2.68 | 1.99E-03 |
| Abhd14b | abhydrolase domain containing 14b [Source:MGI Symbol;Acc:MGI:1923741] | -1.18 | 2.00E-03 |
| Armc2 | armadillo repeat containing 2 [Source:MGI Symbol;Acc:MGI:1916449] | -1.14 | 2.01E-03 |
| Serpinb10 | serine (or cysteine) peptidase inhibitor, clade B (ovalbumin), member 10 [Source:MGI Symbol;Acc:MGI:2138648] | -0.97 | 2.03E-03 |
| Nlrc5 | NLR family, CARD domain containing 5 [Source:MGI Symbol;Acc:MGI:3612191] | -1.04 | 2.04E-03 |
| Mcu | mitochondrial calcium uniporter [Source:MGI Symbol;Acc:MGI:3026965] | -0.78 | 2.06E-03 |
| Nova1 | NOVA alternative splicing regulator 1 [Source:MGI Symbol;Acc:MGI:104297] | -1.38 | 2.10E-03 |
| Cfap100 | cilia and flagella associated protein 100 [Source:MGI Symbol;Acc:MGI:2141635] | -1.63 | 2.12E-03 |
| Kcna1 | potassium voltage-gated channel, shaker-related subfamily, member 1 [Source:MGI Symbol;Acc:MGI:96654] | -1.35 | 2.15E-03 |
| Ar | androgen receptor [Source:MGI Symbol;Acc:MGI:88064] | -1.38 | 2.18E-03 |
| Gm17655 | predicted gene, 17655 [Source:MGI Symbol;Acc:MGI:4937289] | -1.72 | 2.20E-03 |
| Ndufs5-ps | NADH:ubiquinone oxidoreductase core subunit S5, pseudogene [Source:MGI Symbol;Acc:MGI:3612445] | -2.07 | 2.20E-03 |
| Txnip | thioredoxin interacting protein [Source:MGI Symbol;Acc:MGI:1889549] | -1.47 | 2.25E-03 |
| Gm21985 | predicted gene 21985 [Source:MGI Symbol;Acc:MGI:5439454] | -1.27 | 2.26E-03 |

| | | | |
|---|---|---|---|
| Fam81a | family with sequence similarity 81, member A [Source:MGI Symbol;Acc:MGI:1924136] | -1.04 | 2.29E-03 |
| Scd1 | stearoyl-Coenzyme A desaturase 1 [Source:MGI Symbol;Acc:MGI:98239] | -1.55 | 2.30E-03 |
| Gm19120 | predicted gene, 19120 [Source:MGI Symbol;Acc:MGI:5011305] | -1.87 | 2.34E-03 |
| Mcf2l | mcf.2 transforming sequence-like [Source:MGI Symbol;Acc:MGI:103263] | -1.15 | 2.36E-03 |
| Limd1 | LIM domains containing 1 [Source:MGI Symbol;Acc:MGI:1352502] | -0.82 | 2.37E-03 |
| Caprin2 | caprin family member 2 [Source:MGI Symbol;Acc:MGI:2448541] | -0.84 | 2.37E-03 |
| Ikzf2 | IKAROS family zinc finger 2 [Source:MGI Symbol;Acc:MGI:1342541] | -1.88 | 2.38E-03 |
| Erbb4 | erb-b2 receptor tyrosine kinase 4 [Source:MGI Symbol;Acc:MGI:104771] | -4.23 | 2.38E-03 |
| Thra | thyroid hormone receptor alpha [Source:MGI Symbol;Acc:MGI:98742] | -0.95 | 2.38E-03 |
| Net1 | neuroepithelial cell transforming gene 1 [Source:MGI Symbol;Acc:MGI:1927138] | -1.15 | 2.39E-03 |
| Wfdc3 | WAP four-disulfide core domain 3 [Source:MGI Symbol;Acc:MGI:1923897] | -1.53 | 2.49E-03 |
| Sucnr1 | succinate receptor 1 [Source:MGI Symbol;Acc:MGI:1934135] | -2.87 | 2.51E-03 |
| Gm47630 | predicted gene, 47630 [Source:MGI Symbol;Acc:MGI:6096700] | -1.06 | 2.52E-03 |
| Ppp2r3a | protein phosphatase 2, regulatory subunit B", alpha [Source:MGI Symbol;Acc:MGI:2442104] | -1.36 | 2.52E-03 |
| Usp53 | ubiquitin specific peptidase 53 [Source:MGI Symbol;Acc:MGI:2139607] | -0.90 | 2.53E-03 |
| Hoxa7 | homeobox A7 [Source:MGI Symbol;Acc:MGI:96179] | -1.30 | 2.57E-03 |
| Gm15712 | predicted gene 15712 [Source:MGI Symbol;Acc:MGI:3783154] | -1.30 | 2.61E-03 |
| Ankfn1 | ankyrin-repeat and fibronectin type III domain containing 1 [Source:MGI Symbol;Acc:MGI:2686021] | -3.43 | 2.65E-03 |
| Elfn2 | leucine rich repeat and fibronectin type III, extracellular 2 [Source:MGI Symbol;Acc:MGI:3608416] | -3.72 | 2.65E-03 |
| Adra1a | adrenergic receptor, alpha 1a [Source:MGI Symbol;Acc:MGI:104773] | -1.68 | 2.66E-03 |
| Gm5403 | predicted gene 5403 [Source:MGI Symbol;Acc:MGI:3644700] | -1.47 | 2.71E-03 |
| Lgals12 | lectin, galactose binding, soluble 12 [Source:MGI Symbol;Acc:MGI:1929094] | -1.40 | 2.72E-03 |
| Gm36378 | predicted gene, 36378 [Source:MGI Symbol;Acc:MGI:5595537] | -1.63 | 2.72E-03 |
| Plppr4 | phospholipid phosphatase related 4 [Source:MGI Symbol;Acc:MGI:106530] | -2.21 | 2.76E-03 |
| Inpp4b | inositol polyphosphate-4-phosphatase, type II [Source:MGI Symbol;Acc:MGI:2158925] | -1.40 | 2.79E-03 |
| Defb12 | defensin beta 12 [Source:MGI Symbol;Acc:MGI:1924924] | -5.42 | 2.79E-03 |
| Srgap2 | SLIT-ROBO Rho GTPase activating protein 2 [Source:MGI Symbol;Acc:MGI:109605] | -0.90 | 2.82E-03 |
| Mturn | maturin, neural progenitor differentiation regulator homolog (Xenopus) [Source:MGI Symbol;Acc:MGI:1915485] | -1.01 | 2.87E-03 |
| Hepacam2 | HEPACAM family member 2 [Source:MGI Symbol;Acc:MGI:2141520] | -1.14 | 2.87E-03 |
| Zfp773 | zinc finger protein 773 [Source:MGI Symbol;Acc:MGI:1923623] | -1.60 | 2.87E-03 |
| Skint2 | selection and upkeep of intraepithelial T cells 2 [Source:MGI Symbol;Acc:MGI:3649629] | -2.01 | 2.88E-03 |
| Zkscan7 | zinc finger with KRAB and SCAN domains 7 [Source:MGI Symbol;Acc:MGI:3040678] | -1.01 | 2.90E-03 |
| Gm13169 | predicted gene 13169 [Source:MGI Symbol;Acc:MGI:3651793] | -6.14 | 2.94E-03 |
| Klf8 | Kruppel-like transcription factor 8 [Source:MGI Symbol;Acc:MGI:2442430] | -1.09 | 2.95E-03 |
| Psg17 | pregnancy specific beta-1-glycoprotein 17 [Source:MGI Symbol;Acc:MGI:1347250] | -2.53 | 2.99E-03 |
| Myoc | myocilin [Source:MGI Symbol;Acc:MGI:1202864] | -3.57 | 3.04E-03 |
| Prkar2b | protein kinase, cAMP dependent regulatory, type II beta [Source:MGI Symbol;Acc:MGI:97760] | -1.24 | 3.07E-03 |
| Zfp932 | zinc finger protein 932 [Source:MGI Symbol;Acc:MGI:1916754] | -0.98 | 3.08E-03 |
| Ptpn14 | protein tyrosine phosphatase, non-receptor type 14 [Source:MGI Symbol;Acc:MGI:102467] | -1.39 | 3.20E-03 |
| Msh6 | mutS homolog 6 [Source:MGI Symbol;Acc:MGI:1343961] | -0.74 | 3.20E-03 |
| Cd36 | CD36 molecule [Source:MGI Symbol;Acc:MGI:107899] | -1.79 | 3.24E-03 |
| Gm52977 | predicted gene, 52977 [Source:MGI Symbol;Acc:MGI:6388859] | -1.98 | 3.26E-03 |
| Zfp788 | zinc finger protein 788 [Source:MGI Symbol;Acc:MGI:1914857] | -0.78 | 3.27E-03 |
| Tshr | thyroid stimulating hormone receptor [Source:MGI Symbol;Acc:MGI:98849] | -2.05 | 3.30E-03 |
| Fam135b | family with sequence similarity 135, member B [Source:MGI Symbol;Acc:MGI:1917613] | -1.56 | 3.33E-03 |
| Ssbp2 | single-stranded DNA binding protein 2 [Source:MGI Symbol;Acc:MGI:1914220] | -0.67 | 3.34E-03 |
| Fam149b | family with sequence similarity 149, member B [Source:MGI Symbol;Acc:MGI:2145567] | -0.77 | 3.38E-03 |
| Chat | choline acetyltransferase [Source:MGI Symbol;Acc:MGI:88392] | -2.69 | 3.44E-03 |
| Aadacl2fm3 | AADACL2 family member 3 [Source:MGI Symbol;Acc:MGI:3643798] | -3.84 | 3.58E-03 |
| Aldh6a1 | aldehyde dehydrogenase family 6, subfamily A1 [Source:MGI Symbol;Acc:MGI:1915077] | -1.49 | 3.64E-03 |
| Mpz | myelin protein zero [Source:MGI Symbol;Acc:MGI:103177] | -1.34 | 3.66E-03 |
| Disp1 | dispatched RND transporter family member 1 [Source:MGI Symbol;Acc:MGI:1916147] | -1.18 | 3.67E-03 |
| Ttc3 | tetratricopeptide repeat domain 3 [Source:MGI Symbol;Acc:MGI:1276539] | -0.66 | 3.72E-03 |
| Ppp2r2c | protein phosphatase 2, regulatory subunit B, gamma [Source:MGI Symbol;Acc:MGI:2442660] | -0.91 | 3.75E-03 |
| Rpgrip1l | Rpgrip1-like [Source:MGI Symbol;Acc:MGI:1920563] | -0.92 | 3.78E-03 |
| Abca8a | ATP-binding cassette, sub-family A member 8a [Source:MGI Symbol;Acc:MGI:2386846] | -1.97 | 3.83E-03 |
| Skint4 | selection and upkeep of intraepithelial T cells 4 [Source:MGI Symbol;Acc:MGI:2444425] | -1.57 | 3.84E-03 |
| Rgs9 | regulator of G-protein signaling 9 [Source:MGI Symbol;Acc:MGI:1338824] | -1.10 | 3.87E-03 |
| Pdcd2 | programmed cell death 2 [Source:MGI Symbol;Acc:MGI:104643] | -1.06 | 3.87E-03 |
| Or8b50 | olfactory receptor family 8 subfamily B member 50 [Source:MGI Symbol;Acc:MGI:3030748] | -0.96 | 3.88E-03 |
| Chrdl1 | chordin-like 1 [Source:MGI Symbol;Acc:MGI:1933172] | -1.97 | 3.91E-03 |
| 4930562C15Rik | RIKEN cDNA 4930562C15 gene [Source:MGI Symbol;Acc:MGI:1926059] | -1.17 | 3.98E-03 |
| Sult1e1 | sulfotransferase family 1E, member 1 [Source:MGI Symbol;Acc:MGI:98431] | -3.87 | 4.05E-03 |
| Ephx1 | epoxide hydrolase 1, microsomal [Source:MGI Symbol;Acc:MGI:95405] | -0.98 | 4.18E-03 |
| Gdf10 | growth differentiation factor 10 [Source:MGI Symbol;Acc:MGI:95684] | -1.06 | 4.18E-03 |
| Cyp2j6 | cytochrome P450, family 2, subfamily j, polypeptide 6 [Source:MGI Symbol;Acc:MGI:1270148] | -1.30 | 4.25E-03 |
| Ppp1r9a | protein phosphatase 1, regulatory subunit 9A [Source:MGI Symbol;Acc:MGI:2442401] | -1.36 | 4.27E-03 |
| Or52d1 | olfactory receptor family 52 subfamily D member 1 [Source:MGI Symbol;Acc:MGI:3030480] | -1.17 | 4.28E-03 |
| Flt3l | FMS-like tyrosine kinase 3 ligand [Source:MGI Symbol;Acc:MGI:95560] | -1.08 | 4.31E-03 |
| Kcna2 | potassium voltage-gated channel, shaker-related subfamily, member 2 [Source:MGI Symbol;Acc:MGI:96659] | -1.29 | 4.34E-03 |

| | | | |
|---|---|---|---|
| Gm15824 | predicted gene 15824 [Source:MGI Symbol;Acc:MGI:3801976] | -1.26 | 4.36E-03 |
| 4930522L14Rik | RIKEN cDNA 4930522L14 gene [Source:MGI Symbol;Acc:MGI:1925270] | -1.44 | 4.36E-03 |
| 4930522L14Rik | RIKEN cDNA 4930522L14 gene [Source:NCBI gene (formerly Entrezgene);Acc:100041734] | -1.44 | 4.36E-03 |
| Aldh1a7 | aldehyde dehydrogenase family 1, subfamily A7 [Source:MGI Symbol;Acc:MGI:1347050] | -2.05 | 4.37E-03 |
| Sult1a1 | sulfotransferase family 1A, phenol-preferring, member 1 [Source:MGI Symbol;Acc:MGI:102896] | -1.04 | 4.39E-03 |
| Zfp960 | zinc finger protein 960 [Source:MGI Symbol;Acc:MGI:3052731] | -1.10 | 4.39E-03 |
| Zfp93 | zinc finger protein 93 [Source:MGI Symbol;Acc:MGI:107611] | -1.10 | 4.39E-03 |
| Tspyl4 | TSPY-like 4 [Source:MGI Symbol;Acc:MGI:106393] | -1.02 | 4.46E-03 |
| Nxt2 | nuclear transport factor 2-like export factor 2 [Source:MGI Symbol;Acc:MGI:2147914] | -0.95 | 4.60E-03 |
| Sorbs1 | sorbin and SH3 domain containing 1 [Source:MGI Symbol;Acc:MGI:700014] | -1.15 | 4.62E-03 |
| Gm44216 | predicted gene, 44216 [Source:MGI Symbol;Acc:MGI:5690608] | -1.77 | 4.62E-03 |
| Acss3 | acyl-CoA synthetase short-chain family member 3 [Source:MGI Symbol;Acc:MGI:2685720] | -0.95 | 4.64E-03 |
| Ccl24 | C-C motif chemokine ligand 24 [Source:MGI Symbol;Acc:MGI:1928953] | -1.07 | 4.68E-03 |
| Or4c126 | olfactory receptor family 4 subfamily C member 126 [Source:MGI Symbol;Acc:MGI:3031095] | -1.16 | 4.73E-03 |
| B3galt5 | UDP-Gal:betaGlcNAc beta 1,3-galactosyltransferase, polypeptide 5 [Source:MGI Symbol;Acc:MGI:2136878] | -1.22 | 4.79E-03 |
| Rps18-ps5 | ribosomal protein S18, pseudogene 5 [Source:MGI Symbol;Acc:MGI:3649931] | -3.60 | 4.79E-03 |
| Ntng1 | netrin G1 [Source:MGI Symbol;Acc:MGI:1934028] | -1.80 | 4.88E-03 |
| Mmd | monocyte to macrophage differentiation-associated [Source:MGI Symbol;Acc:MGI:1914718] | -1.15 | 4.92E-03 |
| Cpxm2 | carboxypeptidase X, M14 family member 2 [Source:MGI Symbol;Acc:MGI:1926006] | -1.58 | 4.92E-03 |
| Tmod2 | tropomodulin 2 [Source:MGI Symbol;Acc:MGI:1355335] | -0.96 | 4.94E-03 |
| Gal3st4 | galactose-3-O-sulfotransferase 4 [Source:MGI Symbol;Acc:MGI:1916254] | -1.59 | 4.96E-03 |
| Vmn1r5 | vomeronasal 1 receptor 5 [Source:MGI Symbol;Acc:MGI:2159455] | -1.05 | 5.30E-03 |
| Marveld2 | MARVEL (membrane-associating) domain containing 2 [Source:MGI Symbol;Acc:MGI:2446166] | -1.36 | 5.30E-03 |
| Hsf3 | heat shock transcription factor 3 [Source:MGI Symbol;Acc:MGI:3045337] | -1.51 | 5.38E-03 |
| Gm18599 | predicted gene, 18599 [Source:MGI Symbol;Acc:MGI:5010784] | -2.27 | 5.44E-03 |
| Zfp2 | zinc finger protein 2 [Source:MGI Symbol;Acc:MGI:99167] | -1.18 | 5.47E-03 |
| Rad51b | RAD51 paralog B [Source:MGI Symbol;Acc:MGI:1099436] | -1.20 | 5.49E-03 |
| Or1o11 | olfactory receptor family 1 subfamily O member 11 [Source:MGI Symbol;Acc:MGI:2177491] | -5.22 | 5.70E-03 |
| Trpm3 | transient receptor potential cation channel, subfamily M, member 3 [Source:MGI Symbol;Acc:MGI:2443101] | -1.07 | 5.74E-03 |
| Zfp846 | zinc finger protein 846 [Source:MGI Symbol;Acc:MGI:1924012] | -0.92 | 5.78E-03 |
| Ppm1k | protein phosphatase 1K (PP2C domain containing) [Source:MGI Symbol;Acc:MGI:2442111] | -0.91 | 5.88E-03 |
| Col4a6 | collagen, type IV, alpha 6 [Source:MGI Symbol;Acc:MGI:2152695] | -1.55 | 5.89E-03 |
| Adam1b | a disintegrin and metallopeptidase domain 1b [Source:MGI Symbol;Acc:MGI:2429506] | -2.13 | 5.90E-03 |
| Lrrcc1 | leucine rich repeat and coiled-coil domain containing 1 [Source:MGI Symbol;Acc:MGI:1918960] | -0.94 | 5.91E-03 |
| Rnf13 | ring finger protein 13 [Source:MGI Symbol;Acc:MGI:1346341] | -1.17 | 5.91E-03 |
| Cnksr2 | connector enhancer of kinase suppressor of Ras 2 [Source:MGI Symbol;Acc:MGI:2661175] | -1.61 | 6.11E-03 |
| Zkscan8 | zinc finger with KRAB and SCAN domains 8 [Source:MGI Symbol;Acc:MGI:1913815] | -0.79 | 6.21E-03 |
| Gm29487 | predicted gene 29487 [Source:MGI Symbol;Acc:MGI:5580193] | -1.26 | 6.21E-03 |
| Gm7909 | predicted gene 7909 [Source:MGI Symbol;Acc:MGI:3645643] | -1.31 | 6.21E-03 |
| Krt24 | keratin 24 [Source:MGI Symbol;Acc:MGI:1922956] | -1.71 | 6.21E-03 |
| Cd3g | CD3 antigen, gamma polypeptide [Source:MGI Symbol;Acc:MGI:88333] | -0.94 | 6.24E-03 |
| Ahr | aryl-hydrocarbon receptor [Source:MGI Symbol;Acc:MGI:105043] | -0.89 | 6.24E-03 |
| Zfp995 | zinc finger protein 995 [Source:MGI Symbol;Acc:MGI:1917331] | -0.74 | 6.26E-03 |
| Ghr | growth hormone receptor [Source:MGI Symbol;Acc:MGI:95708] | -1.30 | 6.34E-03 |
| Gm21013 | predicted gene, 21013 [Source:MGI Symbol;Acc:MGI:5434368] | -2.17 | 6.35E-03 |
| Dlg2 | discs large MAGUK scaffold protein 2 [Source:MGI Symbol;Acc:MGI:1344351] | -1.73 | 6.38E-03 |
| Fgf12 | fibroblast growth factor 12 [Source:MGI Symbol;Acc:MGI:109183] | -1.19 | 6.40E-03 |
| Scml2 | Scm polycomb group protein like 2 [Source:MGI Symbol;Acc:MGI:1340042] | -1.58 | 6.44E-03 |
| Aff1 | AF4/FMR2 family, member 1 [Source:MGI Symbol;Acc:MGI:1100819] | -0.85 | 6.50E-03 |
| Ighm | immunoglobulin heavy constant mu [Source:MGI Symbol;Acc:MGI:96448] | -1.91 | 6.55E-03 |
| Trgc1 | T cell receptor gamma, constant 1 [Source:MGI Symbol;Acc:MGI:98625] | -1.56 | 6.58E-03 |
| Zfp1007 | zinc finger protein 1007 [Source:MGI Symbol;Acc:MGI:1924450] | -0.90 | 6.60E-03 |
| Rgs18 | regulator of G-protein signaling 18 [Source:MGI Symbol;Acc:MGI:1927498] | -1.50 | 6.66E-03 |
| Gm13312 | predicted gene 13312 [Source:MGI Symbol;Acc:MGI:3651094] | -1.52 | 6.82E-03 |
| Csrnp2 | cysteine-serine-rich nuclear protein 2 [Source:MGI Symbol;Acc:MGI:2386852] | -0.87 | 6.84E-03 |
| Retnla | resistin like alpha [Source:MGI Symbol;Acc:MGI:1888504] | -3.06 | 6.84E-03 |
| Slc7a10 | solute carrier family 7 (cationic amino acid transporter, y+ system), member 10 [Source:MGI Symbol;Acc:MGI:1858261] | -2.20 | 6.89E-03 |
| Tasor | transcription activation suppressor [Source:MGI Symbol;Acc:MGI:1921694] | -0.67 | 6.94E-03 |
| Nox1 | NADPH oxidase 1 [Source:MGI Symbol;Acc:MGI:2450016] | -1.85 | 6.97E-03 |
| Ptprr | protein tyrosine phosphatase receptor type R [Source:MGI Symbol;Acc:MGI:109559] | -0.86 | 6.98E-03 |
| Gm18259 | predicted gene, 18259 [Source:MGI Symbol;Acc:MGI:5010444] | -1.19 | 7.00E-03 |
| Akr1c21 | aldo-keto reductase family 1, member C21 [Source:MGI Symbol;Acc:MGI:1924587] | -3.16 | 7.00E-03 |
| Usp17lc | ubiquitin specific peptidase 17-like C [Source:MGI Symbol;Acc:MGI:107698] | -1.54 | 7.01E-03 |
| Gm47753 | predicted gene, 47753 [Source:MGI Symbol;Acc:MGI:6096899] | -1.72 | 7.08E-03 |
| Clec2i | C-type lectin domain family 2, member i [Source:MGI Symbol;Acc:MGI:2136650] | -1.36 | 7.24E-03 |
| Abca3 | ATP-binding cassette, sub-family A member 3 [Source:MGI Symbol;Acc:MGI:1351617] | -0.70 | 7.26E-03 |
| Tpk1 | thiamine pyrophosphokinase [Source:MGI Symbol;Acc:MGI:1352500] | -0.81 | 7.31E-03 |
| Trim34a | tripartite motif-containing 34A [Source:MGI Symbol;Acc:MGI:2137359] | -1.10 | 7.38E-03 |
| Syt3 | synaptotagmin III [Source:MGI Symbol;Acc:MGI:99665] | -1.37 | 7.38E-03 |
| Trim2 | tripartite motif-containing 2 [Source:MGI Symbol;Acc:MGI:1933163] | -1.23 | 7.39E-03 |

| | | | |
|---|---|---|---|
| Abca8b | ATP-binding cassette, sub-family A member 8b [Source:MGI Symbol;Acc:MGI:1351668] | -1.49 | 7.39E-03 |
| Vmn2r57 | vomeronasal 2, receptor 57 [Source:MGI Symbol;Acc:MGI:3703084] | -2.28 | 7.54E-03 |
| Gm12889 | predicted gene 12889 [Source:MGI Symbol;Acc:MGI:3652132] | -0.93 | 7.59E-03 |
| Rpl30-ps11 | ribosomal protein L30, pseudogene 11 [Source:MGI Symbol;Acc:MGI:3642364] | -5.63 | 7.76E-03 |
| 1700019D03Rik | RIKEN cDNA 1700019D03 gene [Source:MGI Symbol;Acc:MGI:1914330] | -1.58 | 7.78E-03 |
| Cecr2 | CECR2, histone acetyl-lysine reader [Source:MGI Symbol;Acc:MGI:1923799] | -0.99 | 7.90E-03 |
| C9 | complement component 9 [Source:MGI Symbol;Acc:MGI:1098282] | -1.78 | 7.90E-03 |
| Fbxo16 | F-box protein 16 [Source:MGI Symbol;Acc:MGI:1354706] | -1.71 | 7.99E-03 |
| Slco2b1 | solute carrier organic anion transporter family, member 2b1 [Source:MGI Symbol;Acc:MGI:1351872] | -1.29 | 8.01E-03 |
| Lctl | lactase-like [Source:MGI Symbol;Acc:MGI:2183549] | -2.33 | 8.02E-03 |
| Gga2 | golgi associated, gamma adaptin ear containing, ARF binding protein 2 [Source:MGI Symbol;Acc:MGI:1921355] | -0.75 | 8.05E-03 |
| Rgs22 | regulator of G-protein signalling 22 [Source:MGI Symbol;Acc:MGI:3613651] | -1.49 | 8.05E-03 |
| Septin3 | septin 3 [Source:MGI Symbol;Acc:MGI:1345148] | -1.17 | 8.08E-03 |
| Camta1 | calmodulin binding transcription activator 1 [Source:MGI Symbol;Acc:MGI:2140230] | -0.99 | 8.11E-03 |
| Bmp5 | bone morphogenetic protein 5 [Source:MGI Symbol;Acc:MGI:88181] | -2.04 | 8.12E-03 |
| Crp | C-reactive protein, pentraxin-related [Source:MGI Symbol;Acc:MGI:88512] | -1.03 | 8.17E-03 |
| Daglb | diacylglycerol lipase, beta [Source:MGI Symbol;Acc:MGI:2442032] | -0.67 | 8.26E-03 |
| Fam221a | family with sequence similarity 221, member A [Source:MGI Symbol;Acc:MGI:2442161] | -1.48 | 8.27E-03 |
| Zfp869 | zinc finger protein 869 [Source:MGI Symbol;Acc:MGI:1914119] | -0.78 | 8.30E-03 |
| Kcnn2 | potassium intermediate/small conductance calcium-activated channel, subfamily N, member 2 [Source:MGI Symbol;Acc:MGI:2153182] | -1.09 | 8.36E-03 |
| Adam1a | a disintegrin and metallopeptidase domain 1a [Source:MGI Symbol;Acc:MGI:2429504] | -0.99 | 8.39E-03 |
| Gm379 | predicted gene 379 [Source:MGI Symbol;Acc:MGI:2685225] | -0.91 | 8.47E-03 |
| Hyal6 | hyaluronoglucosaminidase 6 [Source:MGI Symbol;Acc:MGI:1921659] | -3.37 | 8.56E-03 |
| Wrn | Werner syndrome RecQ like helicase [Source:MGI Symbol;Acc:MGI:109635] | -0.75 | 8.59E-03 |
| Ubash3a | ubiquitin associated and SH3 domain containing, A [Source:MGI Symbol;Acc:MGI:1926074] | -0.86 | 8.61E-03 |
| Krt15 | keratin 15 [Source:MGI Symbol;Acc:MGI:96689] | -2.27 | 8.64E-03 |
| Fut8 | fucosyltransferase 8 [Source:MGI Symbol;Acc:MGI:1858901] | -0.91 | 8.65E-03 |
| Maob | monoamine oxidase B [Source:MGI Symbol;Acc:MGI:96916] | -1.64 | 8.65E-03 |
| Zbtb20 | zinc finger and BTB domain containing 20 [Source:MGI Symbol;Acc:MGI:1929213] | -1.13 | 8.79E-03 |
| Rhobtb1 | Rho-related BTB domain containing 1 [Source:MGI Symbol;Acc:MGI:1916538] | -0.84 | 8.87E-03 |
| Ccdc27 | coiled-coil domain containing 27 [Source:MGI Symbol;Acc:MGI:2685881] | -1.59 | 8.87E-03 |
| Mccc1 | methylcrotonoyl-Coenzyme A carboxylase 1 (alpha) [Source:MGI Symbol;Acc:MGI:1919289] | -0.92 | 8.92E-03 |
| Pik3ip1 | phosphoinositide-3-kinase interacting protein 1 [Source:MGI Symbol;Acc:MGI:1917016] | -1.62 | 9.04E-03 |
| Rorc | RAR-related orphan receptor gamma [Source:MGI Symbol;Acc:MGI:104856] | -1.37 | 9.08E-03 |
| Gm3604 | predicted gene 3604 [Source:MGI Symbol;Acc:MGI:3781781] | -1.58 | 9.09E-03 |
| Zfp39 | zinc finger protein 39 [Source:MGI Symbol;Acc:MGI:99183] | -1.22 | 9.12E-03 |
| Nog | noggin [Source:MGI Symbol;Acc:MGI:104327] | -1.32 | 9.13E-03 |
| Zc3h6 | zinc finger CCCH type containing 6 [Source:MGI Symbol;Acc:MGI:1926001] | -1.18 | 9.17E-03 |
| Gm48349 | predicted gene, 48349 [Source:MGI Symbol;Acc:MGI:6097813] | -1.22 | 9.20E-03 |
| Gm49500 | predicted gene, 49500 [Source:MGI Symbol;Acc:MGI:6155185] | -1.71 | 9.45E-03 |
| Akap7 | A kinase anchor protein 7 [Source:MGI Symbol;Acc:MGI:1859150] | -1.28 | 9.58E-03 |
| Fmo5 | flavin containing monooxygenase 5 [Source:MGI Symbol;Acc:MGI:1310004] | -0.93 | 9.58E-03 |
| Retn | resistin [Source:MGI Symbol;Acc:MGI:1888506] | -1.54 | 9.61E-03 |
| Nr3c2 | nuclear receptor subfamily 3, group C, member 2 [Source:MGI Symbol;Acc:MGI:99459] | -1.53 | 9.72E-03 |
| Or6c3 | olfactory receptor family 6 subfamily C member 3 [Source:MGI Symbol;Acc:MGI:3030622] | -1.43 | 9.76E-03 |
| Mmp27 | matrix metallopeptidase 27 [Source:MGI Symbol;Acc:MGI:3039232] | -0.97 | 9.77E-03 |
| Pax5 | paired box 5 [Source:MGI Symbol;Acc:MGI:97489] | -3.82 | 9.82E-03 |
| Rgsl1 | regulator of G-protein signaling like 1 [Source:MGI Symbol;Acc:MGI:2685048] | -1.14 | 9.90E-03 |
| Tnfrsf19 | tumor necrosis factor receptor superfamily, member 19 [Source:MGI Symbol;Acc:MGI:1352474] | -1.54 | 9.90E-03 |
| Fnbp1 | formin binding protein 1 [Source:MGI Symbol;Acc:MGI:109606] | -0.70 | 1.00E-02 |
| Rab30 | RAB30, member RAS oncogene family [Source:MGI Symbol;Acc:MGI:1923235] | -1.17 | 1.00E-02 |
| Micu2 | mitochondrial calcium uptake 2 [Source:MGI Symbol;Acc:MGI:1915764] | -0.72 | 1.02E-02 |
| Mob3b | MOB kinase activator 3B [Source:MGI Symbol;Acc:MGI:2664539] | -1.12 | 1.04E-02 |
| Smco3 | single-pass membrane protein with coiled-coil domains 3 [Source:MGI Symbol;Acc:MGI:2443451] | -2.28 | 1.05E-02 |
| Or2a52 | olfactory receptor family 2 subfamily A member 52 [Source:MGI Symbol;Acc:MGI:3030271] | -4.54 | 1.07E-02 |
| Ces1d | carboxylesterase 1D [Source:MGI Symbol;Acc:MGI:2148202] | -1.49 | 1.08E-02 |
| Npy1r | neuropeptide Y receptor Y1 [Source:MGI Symbol;Acc:MGI:104963] | -1.14 | 1.09E-02 |
| B4galnt2 | beta-1,4-N-acetyl-galactosaminyl transferase 2 [Source:MGI Symbol;Acc:MGI:1342058] | -1.53 | 1.10E-02 |
| Scd3 | stearoyl-coenzyme A desaturase 3 [Source:MGI Symbol;Acc:MGI:1353437] | -1.83 | 1.10E-02 |
| Vmn1r42 | vomeronasal 1 receptor 42 [Source:MGI Symbol;Acc:MGI:2148511] | -1.07 | 1.11E-02 |
| Gna14 | guanine nucleotide binding protein, alpha 14 [Source:MGI Symbol;Acc:MGI:95769] | -1.27 | 1.11E-02 |
| Zscan18 | zinc finger and SCAN domain containing 18 [Source:MGI Symbol;Acc:MGI:3643810] | -1.08 | 1.11E-02 |
| Trim66 | tripartite motif-containing 66 [Source:MGI Symbol;Acc:MGI:2152406] | -1.36 | 1.12E-02 |
| Mc2r | melanocortin 2 receptor [Source:MGI Symbol;Acc:MGI:96928] | -3.39 | 1.13E-02 |
| Serpina1c | serine (or cysteine) peptidase inhibitor, clade A, member 1C [Source:MGI Symbol;Acc:MGI:891969] | -4.18 | 1.13E-02 |
| Elac1 | elaC ribonuclease Z 1 [Source:MGI Symbol;Acc:MGI:1890495] | -0.90 | 1.14E-02 |
| Gm6291 | predicted gene 6291 [Source:MGI Symbol;Acc:MGI:3648803] | -1.43 | 1.14E-02 |
| Ccdc40 | coiled-coil domain containing 40 [Source:MGI Symbol;Acc:MGI:2443893] | -2.31 | 1.14E-02 |
| Teddm2 | transmembrane epididymal family member 2 [Source:MGI Symbol;Acc:MGI:1923273] | -1.00 | 1.17E-02 |

| | | | |
|---|---|---|---|
| Alox12e | arachidonate lipoxygenase, epidermal [Source:MGI Symbol;Acc:MGI:1274790] | -2.04 | 1.17E-02 |
| Enox2 | ecto-NOX disulfide-thiol exchanger 2 [Source:MGI Symbol;Acc:MGI:2384799] | -0.94 | 1.20E-02 |
| Kank1 | KN motif and ankyrin repeat domains 1 [Source:MGI Symbol;Acc:MGI:2147707] | -1.18 | 1.21E-02 |
| Heca | hdc homolog, cell cycle regulator [Source:MGI Symbol;Acc:MGI:2685715] | -1.00 | 1.21E-02 |
| Gdi1 | GDP dissociation inhibitor 1 [Source:MGI Symbol;Acc:MGI:99846] | -0.73 | 1.23E-02 |
| Apol10b | apolipoprotein L 10B [Source:MGI Symbol;Acc:MGI:3043522] | -1.14 | 1.24E-02 |
| Gm3188 | predicted gene 3188 [Source:MGI Symbol;Acc:MGI:3781367] | -1.17 | 1.24E-02 |
| Mtmr11 | myotubularin related protein 11 [Source:MGI Symbol;Acc:MGI:2652817] | -0.73 | 1.25E-02 |
| Cd226 | CD226 antigen [Source:MGI Symbol;Acc:MGI:3039602] | -1.23 | 1.26E-02 |
| Rasgrf1 | RAS protein-specific guanine nucleotide-releasing factor 1 [Source:MGI Symbol;Acc:MGI:99694] | -0.75 | 1.27E-02 |
| Dzip1 | DAZ interacting protein 1 [Source:MGI Symbol;Acc:MGI:1914311] | -0.83 | 1.28E-02 |
| Grid1 | glutamate receptor, ionotropic, delta 1 [Source:MGI Symbol;Acc:MGI:95812] | -1.38 | 1.30E-02 |
| Tnk1 | tyrosine kinase, non-receptor, 1 [Source:MGI Symbol;Acc:MGI:1930958] | -0.76 | 1.30E-02 |
| Trdmt1 | tRNA aspartic acid methyltransferase 1 [Source:MGI Symbol;Acc:MGI:1274787] | -0.68 | 1.30E-02 |
| Cyp39a1 | cytochrome P450, family 39, subfamily a, polypeptide 1 [Source:MGI Symbol;Acc:MGI:1927096] | -1.16 | 1.30E-02 |
| Or10w1 | olfactory receptor family 10 subfamily W member 1 [Source:MGI Symbol;Acc:MGI:3031324] | -1.18 | 1.30E-02 |
| Diras2 | DIRAS family, GTP-binding RAS-like 2 [Source:MGI Symbol;Acc:MGI:1915453] | -1.38 | 1.30E-02 |
| Or8b41 | olfactory receptor family 8 subfamily B member 41 [Source:MGI Symbol;Acc:MGI:3030724] | -1.12 | 1.30E-02 |
| Ccdc141 | coiled-coil domain containing 141 [Source:MGI Symbol;Acc:MGI:1919735] | -0.88 | 1.30E-02 |
| Gvin-ps6 | GTPase, very large interferon inducible, pseudogene 6 [Source:MGI Symbol;Acc:MGI:3647753] | -1.62 | 1.31E-02 |
| Ifih1 | interferon induced with helicase C domain 1 [Source:MGI Symbol;Acc:MGI:1918836] | -0.79 | 1.31E-02 |
| Dpyd | dihydropyrimidine dehydrogenase [Source:MGI Symbol;Acc:MGI:2139667] | -1.85 | 1.31E-02 |
| Rif1 | replication timing regulatory factor 1 [Source:MGI Symbol;Acc:MGI:1098622] | -0.71 | 1.32E-02 |
| Smim1 | small integral membrane protein 1 [Source:MGI Symbol;Acc:MGI:1916109] | -1.02 | 1.32E-02 |
| Vit | vitrin [Source:MGI Symbol;Acc:MGI:1921449] | -1.01 | 1.33E-02 |
| BC024139 | cDNA sequence BC024139 [Source:MGI Symbol;Acc:MGI:2442591] | -1.23 | 1.35E-02 |
| Nr1d2 | nuclear receptor subfamily 1, group D, member 2 [Source:MGI Symbol;Acc:MGI:2449205] | -1.02 | 1.35E-02 |
| Gm15079 | predicted gene 15079 [Source:MGI Symbol;Acc:MGI:3705703] | -2.97 | 1.35E-02 |
| 4930523C07Rik | RIKEN cDNA 4930523C07 gene [Source:MGI Symbol;Acc:MGI:1914897] | -0.82 | 1.36E-02 |
| Adcy6 | adenylate cyclase 6 [Source:MGI Symbol;Acc:MGI:87917] | -0.80 | 1.37E-02 |
| Mccc2 | methylcrotonoyl-Coenzyme A carboxylase 2 (beta) [Source:MGI Symbol;Acc:MGI:1925288] | -1.04 | 1.37E-02 |
| Map3k21 | mitogen-activated protein kinase kinase kinase 21 [Source:MGI Symbol;Acc:MGI:2385307] | -1.88 | 1.37E-02 |
| Ankrd29 | ankyrin repeat domain 29 [Source:MGI Symbol;Acc:MGI:2687055] | -1.09 | 1.37E-02 |
| Wdr93 | WD repeat domain 93 [Source:MGI Symbol;Acc:MGI:3646885] | -1.62 | 1.37E-02 |
| Cdkn1b | cyclin dependent kinase inhibitor 1B [Source:MGI Symbol;Acc:MGI:104565] | -0.86 | 1.38E-02 |
| Mrgprb2 | MAS-related GPR, member B2 [Source:MGI Symbol;Acc:MGI:2441674] | -1.73 | 1.38E-02 |
| Mcc | mutated in colorectal cancers [Source:MGI Symbol;Acc:MGI:96930] | -1.06 | 1.38E-02 |
| Cacna1e | calcium channel, voltage-dependent, R type, alpha 1E subunit [Source:MGI Symbol;Acc:MGI:106217] | -1.10 | 1.38E-02 |
| Pcmtd2 | protein-L-isoaspartate (D-aspartate) O-methyltransferase domain containing 2 [Source:MGI Symbol;Acc:MGI:1923927] | -0.84 | 1.40E-02 |
| Ccdc122 | coiled-coil domain containing 122 [Source:MGI Symbol;Acc:MGI:1918358] | -0.85 | 1.40E-02 |
| Il20ra | interleukin 20 receptor, alpha [Source:MGI Symbol;Acc:MGI:3605069] | -1.74 | 1.40E-02 |
| Dgkk | diacylglycerol kinase kappa [Source:MGI Symbol;Acc:MGI:3580254] | -1.97 | 1.40E-02 |
| Psmb9 | proteasome (prosome, macropain) subunit, beta type 9 (large multifunctional peptidase 2) [Source:MGI Symbol;Acc:MGI:1346526] | -0.92 | 1.42E-02 |
| Rpl31-ps20 | ribosomal protein L31, pseudogene 20 [Source:MGI Symbol;Acc:MGI:3644512] | -5.52 | 1.42E-02 |
| Apol7e | apolipoprotein L 7e [Source:MGI Symbol;Acc:MGI:3704456] | -0.72 | 1.42E-02 |
| Il34 | interleukin 34 [Source:MGI Symbol;Acc:MGI:1923777] | -1.24 | 1.44E-02 |
| Or2h2b-ps1 | olfactory receptor family 2 subfamily H member 2B, pseudogene 1 [Source:MGI Symbol;Acc:MGI:3030587] | -1.85 | 1.44E-02 |
| Gm4297 | predicted gene 4297 [Source:MGI Symbol;Acc:MGI:3782476] | -0.98 | 1.45E-02 |
| Otud1 | OTU domain containing 1 [Source:MGI Symbol;Acc:MGI:1918448] | -1.47 | 1.48E-02 |
| Phka2 | phosphorylase kinase alpha 2 [Source:MGI Symbol;Acc:MGI:97577] | -0.82 | 1.48E-02 |
| Cyp3a13 | cytochrome P450, family 3, subfamily a, polypeptide 13 [Source:MGI Symbol;Acc:MGI:88610] | -1.02 | 1.48E-02 |
| Ovol2 | ovo like zinc finger 2 [Source:MGI Symbol;Acc:MGI:1338039] | -1.26 | 1.48E-02 |
| Phf24 | PHD finger protein 24 [Source:MGI Symbol;Acc:MGI:2140712] | -1.74 | 1.48E-02 |
| Mucl1 | mucin-like 1 [Source:MGI Symbol;Acc:MGI:98393] | -2.31 | 1.48E-02 |
| Rgmb | repulsive guidance molecule family member B [Source:MGI Symbol;Acc:MGI:1916049] | -1.16 | 1.50E-02 |
| Or10ab4 | olfactory receptor family 10 subfamily AB member 4 [Source:MGI Symbol;Acc:MGI:3030313] | -0.90 | 1.52E-02 |
| Wnk2 | WNK lysine deficient protein kinase 2 [Source:MGI Symbol;Acc:MGI:1922857] | -1.61 | 1.52E-02 |
| Ccdc17 | coiled-coil domain containing 17 [Source:MGI Symbol;Acc:MGI:1915667] | -1.17 | 1.54E-02 |
| Tnfsf18 | tumor necrosis factor (ligand) superfamily, member 18 [Source:MGI Symbol;Acc:MGI:2673064] | -2.37 | 1.54E-02 |
| Appl2 | adaptor protein, phosphotyrosine interaction, PH domain and leucine zipper containing 2 [Source:MGI Symbol;Acc:MGI:2384914] | -0.92 | 1.54E-02 |
| Gm11870 | predicted gene 11870 [Source:MGI Symbol;Acc:MGI:3650285] | -0.80 | 1.55E-02 |
| Or10ak16 | olfactory receptor family 10 subfamily AK member 16 [Source:MGI Symbol;Acc:MGI:3031164] | -0.77 | 1.56E-02 |
| Nfib | nuclear factor I/B [Source:MGI Symbol;Acc:MGI:103188] | -0.88 | 1.56E-02 |
| Mtcp1 | mature T cell proliferation 1 [Source:MGI Symbol;Acc:MGI:102699] | -1.04 | 1.57E-02 |
| Gm36262 | predicted gene, 36262 [Source:MGI Symbol;Acc:MGI:5595421] | -1.18 | 1.58E-02 |
| Inhca | inhibitor of carbonic anhydrase [Source:MGI Symbol;Acc:MGI:1919025] | -1.17 | 1.59E-02 |

| | | | |
|---|---|---|---|
| D7Ertd443e | DNA segment, Chr 7, ERATO Doi 443, expressed [Source:MGI Symbol;Acc:MGI:1196431] | -1.44 | 1.60E-02 |
| Zfp953 | zinc finger protein 953 [Source:MGI Symbol;Acc:MGI:3612873] | -0.94 | 1.60E-02 |
| Agmo | alkylglycerol monooxygenase [Source:MGI Symbol;Acc:MGI:2442495] | -1.28 | 1.60E-02 |
| Sh3gl2 | SH3-domain GRB2-like 2 [Source:MGI Symbol;Acc:MGI:700009] | -1.10 | 1.61E-02 |
| Gm30280 | predicted gene, 30280 [Source:MGI Symbol;Acc:MGI:5589439] | -1.54 | 1.61E-02 |
| Ypel2 | yippee like 2 [Source:MGI Symbol;Acc:MGI:1925114] | -0.97 | 1.61E-02 |
| Acad12 | acyl-Coenzyme A dehydrogenase family, member 12 [Source:MGI Symbol;Acc:MGI:2443320] | -0.75 | 1.62E-02 |
| Sh2b1 | SH2B adaptor protein 1 [Source:MGI Symbol;Acc:MGI:1201407] | -0.64 | 1.64E-02 |
| Gm48891 | predicted gene, 48891 [Source:MGI Symbol;Acc:MGI:6098654] | -4.16 | 1.65E-02 |
| Slc12a6 | solute carrier family 12, member 6 [Source:MGI Symbol;Acc:MGI:2135960] | -0.78 | 1.65E-02 |
| R3hdm2 | R3H domain containing 2 [Source:MGI Symbol;Acc:MGI:1919000] | -0.88 | 1.66E-02 |
| Cckbr | cholecystokinin B receptor [Source:MGI Symbol;Acc:MGI:99479] | -1.99 | 1.66E-02 |
| Msh3 | mutS homolog 3 [Source:MGI Symbol;Acc:MGI:109519] | -0.62 | 1.67E-02 |
| Serpina3e-ps | serine (or cysteine) peptidase inhibitor, clade A, member 3E, pseudogene [Source:MGI Symbol;Acc:MGI:2182837] | -2.61 | 1.68E-02 |
| Anxa8 | annexin A8 [Source:MGI Symbol;Acc:MGI:1201374] | -0.97 | 1.69E-02 |
| Slc2a2 | solute carrier family 2 (facilitated glucose transporter), member 2 [Source:MGI Symbol;Acc:MGI:1095438] | -1.01 | 1.69E-02 |
| Kcnu1 | potassium channel, subfamily U, member 1 [Source:MGI Symbol;Acc:MGI:1202300] | -2.08 | 1.69E-02 |
| Scn3a | sodium channel, voltage-gated, type III, alpha [Source:MGI Symbol;Acc:MGI:98249] | -1.06 | 1.70E-02 |
| Cep70 | centrosomal protein 70 [Source:MGI Symbol;Acc:MGI:1915371] | -0.76 | 1.70E-02 |
| Zfp993 | zinc finger protein 993 [Source:MGI Symbol;Acc:MGI:3713585] | -1.52 | 1.72E-02 |
| Or6e1 | olfactory receptor family 6 subfamily E member 1 [Source:MGI Symbol;Acc:MGI:1333764] | -0.84 | 1.72E-02 |
| Tfap2b | transcription factor AP-2 beta [Source:MGI Symbol;Acc:MGI:104672] | -1.14 | 1.72E-02 |
| Smim8 | small integral membrane protein 8 [Source:MGI Symbol;Acc:MGI:1913541] | -1.12 | 1.72E-02 |
| Tceanc | transcription elongation factor A (SII) N-terminal and central domain containing [Source:MGI Symbol;Acc:MGI:2685236] | -0.73 | 1.73E-02 |
| Phyhip | phytanoyl-CoA hydroxylase interacting protein [Source:MGI Symbol;Acc:MGI:1860417] | -1.22 | 1.73E-02 |
| Adam22 | a disintegrin and metallopeptidase domain 22 [Source:MGI Symbol;Acc:MGI:1340046] | -1.12 | 1.73E-02 |
| Orm1 | orosomucoid 1 [Source:MGI Symbol;Acc:MGI:97443] | -1.31 | 1.74E-02 |
| Gm17910 | predicted gene, 17910 [Source:MGI Symbol;Acc:MGI:5010095] | -1.10 | 1.77E-02 |
| Zfp975 | zinc finger protein 975 [Source:MGI Symbol;Acc:MGI:3648690] | -0.92 | 1.78E-02 |
| Gm44578 | predicted gene 44578 [Source:MGI Symbol;Acc:MGI:5753154] | -0.90 | 1.79E-02 |
| L3mbtl4 | L3MBTL4 histone methyl-lysine binding protein [Source:MGI Symbol;Acc:MGI:2444889] | -1.25 | 1.79E-02 |
| Btnl9 | butyrophilin-like 9 [Source:MGI Symbol;Acc:MGI:2442439] | -1.17 | 1.81E-02 |
| Cd209c | CD209c antigen [Source:MGI Symbol;Acc:MGI:2157945] | -0.67 | 1.81E-02 |
| Gm6811 | predicted gene 6811 [Source:MGI Symbol;Acc:MGI:3648804] | -4.87 | 1.81E-02 |
| Slc43a1 | solute carrier family 43, member 1 [Source:MGI Symbol;Acc:MGI:1931352] | -0.90 | 1.83E-02 |
| Kdm4c | lysine (K)-specific demethylase 4C [Source:MGI Symbol;Acc:MGI:1924054] | -0.59 | 1.83E-02 |
| Gm35315 | predicted gene, 35315 [Source:MGI Symbol;Acc:MGI:5594474] | -1.33 | 1.85E-02 |
| Dst | dystonin [Source:MGI Symbol;Acc:MGI:104627] | -0.87 | 1.85E-02 |
| Hif3a | hypoxia inducible factor 3, alpha subunit [Source:MGI Symbol;Acc:MGI:1859778] | -0.90 | 1.88E-02 |
| Pla2r1 | phospholipase A2 receptor 1 [Source:MGI Symbol;Acc:MGI:102468] | -1.03 | 1.91E-02 |
| Kir3dl1 | killer cell immunoglobulin-like receptor, three domains, long cytoplasmic tail, 1 [Source:MGI Symbol;Acc:MGI:2652397] | -3.04 | 1.91E-02 |
| Heph | hephaestin [Source:MGI Symbol;Acc:MGI:1332240] | -1.15 | 1.92E-02 |
| Ap1s3 | adaptor-related protein complex AP-1, sigma 3 [Source:MGI Symbol;Acc:MGI:1891304] | -0.82 | 1.92E-02 |
| Cers6 | ceramide synthase 6 [Source:MGI Symbol;Acc:MGI:2442564] | -0.83 | 1.93E-02 |
| Acer3 | alkaline ceramidase 3 [Source:MGI Symbol;Acc:MGI:1913440] | -0.63 | 1.94E-02 |
| Zfp1008 | zinc finger protein 1008 [Source:MGI Symbol;Acc:MGI:2443901] | -1.02 | 1.94E-02 |
| Pyy | peptide YY [Source:MGI Symbol;Acc:MGI:99924] | -3.28 | 1.95E-02 |
| Arhgef28 | Rho guanine nucleotide exchange factor 28 [Source:MGI Symbol;Acc:MGI:1346016] | -0.96 | 1.95E-02 |
| Lyz3 | lysozyme 3 [Source:MGI Symbol;Acc:MGI:1924647] | -1.67 | 1.95E-02 |
| Sema3e | sema domain, immunoglobulin domain (Ig), short basic domain, secreted, (semaphorin) 3E [Source:MGI Symbol;Acc:MGI:1340034] | -1.07 | 1.95E-02 |
| Tatdn3 | TatD DNase domain containing 3 [Source:MGI Symbol;Acc:MGI:1916222] | -0.80 | 1.96E-02 |
| Trgv5 | T cell receptor gamma, variable 5 [Source:MGI Symbol;Acc:MGI:98635] | -3.33 | 1.96E-02 |
| Chl1 | cell adhesion molecule L1-like [Source:MGI Symbol;Acc:MGI:1098266] | -0.83 | 1.96E-02 |
| Tgfb2 | transforming growth factor, beta 2 [Source:MGI Symbol;Acc:MGI:98726] | -1.24 | 1.97E-02 |
| Tmem100 | transmembrane protein 100 [Source:MGI Symbol;Acc:MGI:1915138] | -1.30 | 1.99E-02 |
| Gm7041 | predicted gene 7041 [Source:MGI Symbol;Acc:MGI:3646096] | -3.11 | 1.99E-02 |
| Dixdc1 | DIX domain containing 1 [Source:MGI Symbol;Acc:MGI:2679721] | -0.79 | 2.01E-02 |
| Teddm1a | transmembrane epididymal protein 1A [Source:MGI Symbol;Acc:MGI:2668439] | -2.26 | 2.02E-02 |
| Svopl | SV2 related protein homolog (rat)-like [Source:MGI Symbol;Acc:MGI:2444335] | -0.74 | 2.03E-02 |
| Sebox | SEBOX homeobox [Source:MGI Symbol;Acc:MGI:108012] | -1.37 | 2.05E-02 |
| Ivns1abp | influenza virus NS1A binding protein [Source:MGI Symbol;Acc:MGI:2152389] | -0.67 | 2.06E-02 |
| AI987944 | expressed sequence AI987944 [Source:MGI Symbol;Acc:MGI:2142079] | -0.99 | 2.06E-02 |
| Trim21 | tripartite motif-containing 21 [Source:MGI Symbol;Acc:MGI:106657] | -1.23 | 2.06E-02 |
| Zfp455 | zinc finger protein 455 [Source:MGI Symbol;Acc:MGI:3040708] | -1.27 | 2.08E-02 |
| Lrba | LPS-responsive beige-like anchor [Source:MGI Symbol;Acc:MGI:1933162] | -0.85 | 2.08E-02 |
| Osr2 | odd-skipped related 2 [Source:MGI Symbol;Acc:MGI:1930813] | -1.25 | 2.09E-02 |
| Rslcan18 | regulator of sex-limitation candidate 18 [Source:MGI Symbol;Acc:MGI:5433745] | -0.88 | 2.11E-02 |
| Gm21569 | predicted gene, 21569 [Source:MGI Symbol;Acc:MGI:5434924] | -0.99 | 2.11E-02 |

| | | | |
|---|---|---|---|
| Itga8 | integrin alpha 8 [Source:MGI Symbol;Acc:MGI:109442] | -1.11 | 2.11E-02 |
| Platr25 | pluripotency associated transcript 25 [Source:MGI Symbol;Acc:MGI:3645613] | -0.86 | 2.11E-02 |
| Or5d35 | olfactory receptor family 5 subfamily D member 35 [Source:MGI Symbol;Acc:MGI:3030995] | -1.02 | 2.11E-02 |
| Vmn2r112 | vomeronasal 2, receptor 112 [Source:MGI Symbol;Acc:MGI:3644292] | -0.70 | 2.13E-02 |
| Scrn3 | secernin 3 [Source:MGI Symbol;Acc:MGI:1921866] | -0.97 | 2.13E-02 |
| Or2a20 | olfactory receptor family 2 subfamily A member 2 [Source:MGI Symbol;Acc:MGI:3030268] | -5.28 | 2.13E-02 |
| Arhgef5 | Rho guanine nucleotide exchange factor 5 [Source:MGI Symbol;Acc:MGI:1858952] | -0.93 | 2.13E-02 |
| Ankrd34c | ankyrin repeat domain 34C [Source:MGI Symbol;Acc:MGI:2685617] | -1.13 | 2.14E-02 |
| Gm4724 | predicted gene 4724 [Source:MGI Symbol;Acc:MGI:3782904] | -1.57 | 2.14E-02 |
| Or5g9 | olfactory receptor family 5 subfamily G member 9 [Source:MGI Symbol;Acc:MGI:3030843] | -1.07 | 2.16E-02 |
| Bcorl1 | BCL6 co-repressor-like 1 [Source:MGI Symbol;Acc:MGI:2443910] | -0.98 | 2.16E-02 |
| Arl4a | ADP-ribosylation factor-like 4A [Source:MGI Symbol;Acc:MGI:99437] | -1.00 | 2.16E-02 |
| Plaat3 | phospholipase A and acyltransferase 3 [Source:MGI Symbol;Acc:MGI:2179715] | -1.27 | 2.17E-02 |
| Macrod2 | mono-ADP ribosylhydrolase 2 [Source:MGI Symbol;Acc:MGI:1920149] | -0.99 | 2.17E-02 |
| Wasf3 | WASP family, member 3 [Source:MGI Symbol;Acc:MGI:2658986] | -1.20 | 2.19E-02 |
| Gm15720 | predicted gene 15720 [Source:MGI Symbol;Acc:MGI:3783164] | -1.48 | 2.20E-02 |
| Cdin1 | CDAN1 interacting nuclease 1 [Source:MGI Symbol;Acc:MGI:3026886] | -0.73 | 2.21E-02 |
| Cd209d | CD209d antigen [Source:MGI Symbol;Acc:MGI:2157947] | -1.09 | 2.22E-02 |
| Irs3 | insulin receptor substrate 3 [Source:MGI Symbol;Acc:MGI:1194882] | -1.71 | 2.24E-02 |
| Psme1 | proteasome (prosome, macropain) activator subunit 1 (PA28 alpha) [Source:MGI Symbol;Acc:MGI:1096367] | -0.67 | 2.24E-02 |
| Gm18669 | predicted gene, 18669 [Source:MGI Symbol;Acc:MGI:5010854] | -2.89 | 2.24E-02 |
| Flrt1 | fibronectin leucine rich transmembrane protein 1 [Source:MGI Symbol;Acc:MGI:3026647] | -1.10 | 2.25E-02 |
| Gpc6 | glypican 6 [Source:MGI Symbol;Acc:MGI:1346322] | -0.84 | 2.27E-02 |
| Or12j4 | olfactory receptor family 12 subfamily J member 4 [Source:MGI Symbol;Acc:MGI:3030367] | -0.81 | 2.28E-02 |
| Or52h7 | olfactory receptor family 52 subfamily H member 7 [Source:MGI Symbol;Acc:MGI:3030486] | -0.90 | 2.28E-02 |
| Gm47597 | predicted gene, 47597 [Source:MGI Symbol;Acc:MGI:6096648] | -1.13 | 2.28E-02 |
| Crebl2 | cAMP responsive element binding protein-like 2 [Source:MGI Symbol;Acc:MGI:1889385] | -0.77 | 2.28E-02 |
| Poli | polymerase (DNA directed), iota [Source:MGI Symbol;Acc:MGI:1347081] | -1.10 | 2.28E-02 |
| Prlr | prolactin receptor [Source:MGI Symbol;Acc:MGI:97763] | -1.19 | 2.28E-02 |
| Cfd | complement factor D [Source:MGI Symbol;Acc:MGI:87931] | -1.70 | 2.29E-02 |
| Zfp667 | zinc finger protein 667 [Source:MGI Symbol;Acc:MGI:2442757] | -1.02 | 2.29E-02 |
| Or5p60 | olfactory receptor family 5 subfamily P member 60 [Source:MGI Symbol;Acc:MGI:3030318] | -0.74 | 2.29E-02 |
| Slc6a14 | solute carrier family 6 (neurotransmitter transporter), member 14 [Source:MGI Symbol;Acc:MGI:1890216] | -0.92 | 2.30E-02 |
| Ldah | lipid droplet associated hydrolase [Source:MGI Symbol;Acc:MGI:1916082] | -0.70 | 2.30E-02 |
| Tmem266 | transmembrane protein 266 [Source:MGI Symbol;Acc:MGI:2142980] | -1.13 | 2.31E-02 |
| Adk | adenosine kinase [Source:MGI Symbol;Acc:MGI:87930] | -0.86 | 2.32E-02 |
| Skint5 | selection and upkeep of intraepithelial T cells 5 [Source:MGI Symbol;Acc:MGI:3650151] | -2.66 | 2.32E-02 |
| Prob1 | proline rich basic protein 1 [Source:MGI Symbol;Acc:MGI:2686460] | -1.81 | 2.32E-02 |
| Gm18899 | predicted gene, 18899 [Source:MGI Symbol;Acc:MGI:5011084] | -1.18 | 2.32E-02 |
| Cytl1 | cytokine-like 1 [Source:MGI Symbol;Acc:MGI:2684993] | -2.09 | 2.32E-02 |
| Gm21827 | predicted gene, 21827 [Source:MGI Symbol;Acc:MGI:5433991] | -3.90 | 2.33E-02 |
| Aqp4 | aquaporin 4 [Source:MGI Symbol;Acc:MGI:107387] | -2.61 | 2.33E-02 |
| Pkia | protein kinase inhibitor, alpha [Source:MGI Symbol;Acc:MGI:104747] | -2.83 | 2.33E-02 |
| Gm9574 | predicted gene 9574 [Source:MGI Symbol;Acc:MGI:3809525] | -1.85 | 2.34E-02 |
| Jakmip3 | janus kinase and microtubule interacting protein 3 [Source:MGI Symbol;Acc:MGI:1921254] | -2.44 | 2.35E-02 |
| Tdrd3 | tudor domain containing 3 [Source:MGI Symbol;Acc:MGI:2444023] | -0.82 | 2.35E-02 |
| Spart | spartin [Source:MGI Symbol;Acc:MGI:2139806] | -0.72 | 2.35E-02 |
| Pla2g2d | phospholipase A2, group IID [Source:MGI Symbol;Acc:MGI:1341796] | -0.77 | 2.36E-02 |
| Rad9b | RAD9 checkpoint clamp component B [Source:MGI Symbol;Acc:MGI:2385231] | -0.90 | 2.36E-02 |
| Bfsp2 | beaded filament structural protein 2, phakinin [Source:MGI Symbol;Acc:MGI:1333828] | -1.89 | 2.37E-02 |
| Gm16107 | predicted gene 16107 [Source:MGI Symbol;Acc:MGI:3801820] | -2.18 | 2.38E-02 |
| Axdnd1 | axonemal dynein light chain domain containing 1 [Source:MGI Symbol;Acc:MGI:1924602] | -0.91 | 2.38E-02 |
| Nlrp5 | NLR family, pyrin domain containing 5 [Source:MGI Symbol;Acc:MGI:1345193] | -1.20 | 2.38E-02 |
| Add3 | adducin 3 [Source:MGI Symbol;Acc:MGI:1351615] | -0.93 | 2.39E-02 |
| Gm28455 | predicted gene 28455 [Source:MGI Symbol;Acc:MGI:5579161] | -1.69 | 2.39E-02 |
| Fas | Fas cell surface death receptor [Source:MGI Symbol;Acc:MGI:95484] | -1.26 | 2.39E-02 |
| Fgf11 | fibroblast growth factor 11 [Source:MGI Symbol;Acc:MGI:109167] | -0.78 | 2.40E-02 |
| Zfp939 | zinc finger protein 939 [Source:MGI Symbol;Acc:MGI:3036240] | -1.08 | 2.40E-02 |
| Zfp853 | zinc finger protein 853 [Source:MGI Symbol;Acc:MGI:2685638] | -0.83 | 2.42E-02 |
| Aldh2 | aldehyde dehydrogenase 2, mitochondrial [Source:MGI Symbol;Acc:MGI:99600] | -0.78 | 2.42E-02 |
| Bmp4 | bone morphogenetic protein 4 [Source:MGI Symbol;Acc:MGI:88180] | -1.13 | 2.42E-02 |
| Ugt8a | UDP galactosyltransferase 8A [Source:MGI Symbol;Acc:MGI:109522] | -1.43 | 2.44E-02 |
| Foxred2 | FAD-dependent oxidoreductase domain containing 2 [Source:MGI Symbol;Acc:MGI:106315] | -0.98 | 2.45E-02 |
| Chst3 | carbohydrate sulfotransferase 3 [Source:MGI Symbol;Acc:MGI:1858224] | -1.01 | 2.46E-02 |
| Irs1 | insulin receptor substrate 1 [Source:MGI Symbol;Acc:MGI:99454] | -1.16 | 2.48E-02 |
| Kcnj2 | potassium inwardly-rectifying channel, subfamily J, member 2 [Source:MGI Symbol;Acc:MGI:104744] | -1.34 | 2.48E-02 |
| Dbp | D site albumin promoter binding protein [Source:MGI Symbol;Acc:MGI:94866] | -1.44 | 2.52E-02 |
| Gm20696 | predicted gene 20696 [Source:MGI Symbol;Acc:MGI:5313143] | -0.84 | 2.52E-02 |
| Col4a3 | collagen, type IV, alpha 3 [Source:MGI Symbol;Acc:MGI:104688] | -1.06 | 2.54E-02 |
| Or4c3 | olfactory receptor family 4 subfamily C member 3 [Source:MGI Symbol;Acc:MGI:3031098] | -1.30 | 2.54E-02 |
| Gm19154 | predicted gene, 19154 [Source:MGI Symbol;Acc:MGI:5011339] | -1.40 | 2.55E-02 |
| Ccdc196 | coiled-coil domain containing 196 [Source:MGI Symbol;Acc:MGI:3645257] | -3.02 | 2.57E-02 |
| Apcdd1 | adenomatosis polyposis coli down-regulated 1 [Source:MGI Symbol;Acc:MGI:3513977] | -0.86 | 2.57E-02 |

| | | | |
|---|---|---|---|
| Itih5 | inter-alpha-trypsin inhibitor, heavy chain 5 [Source:MGI Symbol;Acc:MGI:1925751] | -1.12 | 2.57E-02 |
| Zfp273 | zinc finger protein 273 [Source:MGI Symbol;Acc:MGI:3036278] | -0.82 | 2.57E-02 |
| Aldoc | aldolase C, fructose-bisphosphate [Source:MGI Symbol;Acc:MGI:101863] | -1.19 | 2.57E-02 |
| Tmem178b | transmembrane protein 178B [Source:MGI Symbol;Acc:MGI:3647581] | -0.99 | 2.58E-02 |
| Gm42878 | predicted gene 42878 [Source:MGI Symbol;Acc:MGI:5663015] | -0.68 | 2.59E-02 |
| Gbp2b | guanylate binding protein 2b [Source:MGI Symbol;Acc:MGI:95666] | -2.11 | 2.59E-02 |
| Jakmip2 | janus kinase and microtubule interacting protein 2 [Source:MGI Symbol;Acc:MGI:1923467] | -1.12 | 2.59E-02 |
| Acsm5 | acyl-CoA synthetase medium-chain family member 5 [Source:MGI Symbol;Acc:MGI:2444086] | -1.59 | 2.59E-02 |
| Arhgap12 | Rho GTPase activating protein 12 [Source:MGI Symbol;Acc:MGI:1922665] | -0.87 | 2.59E-02 |
| Mthfd1 | methylenetetrahydrofolate dehydrogenase (NADP+ dependent), methenyltetrahydrofolate cyclohydrolase, formyltetrahydrofolate synthase [Source:MGI Symbol;Acc:MGI:1342005] | -0.60 | 2.60E-02 |
| Trmt9b | tRNA methyltransferase 9B [Source:MGI Symbol;Acc:MGI:2442328] | -1.59 | 2.60E-02 |
| Cacna2d2 | calcium channel, voltage-dependent, alpha 2/delta subunit 2 [Source:MGI Symbol;Acc:MGI:1929813] | -1.07 | 2.61E-02 |
| Gm3619 | predicted gene 3619 [Source:MGI Symbol;Acc:MGI:3781795] | -1.42 | 2.62E-02 |
| Sfrp4 | secreted frizzled-related protein 4 [Source:MGI Symbol;Acc:MGI:892010] | -1.68 | 2.62E-02 |
| Skint11 | selection and upkeep of intraepithelial T cells 11 [Source:MGI Symbol;Acc:MGI:2685415] | -1.96 | 2.62E-02 |
| Gm6195 | predicted pseudogene 6195 [Source:MGI Symbol;Acc:MGI:3647661] | -1.59 | 2.63E-02 |
| Lrrtm3 | leucine rich repeat transmembrane neuronal 3 [Source:MGI Symbol;Acc:MGI:2389177] | -1.30 | 2.63E-02 |
| Gpm6a | glycoprotein m6a [Source:MGI Symbol;Acc:MGI:107671] | -1.46 | 2.64E-02 |
| Uevld | UEV and lactate/malate dehyrogenase domains [Source:MGI Symbol;Acc:MGI:1860490] | -0.71 | 2.65E-02 |
| Gm55889 | predicted gene, 55889 [Source:MGI Symbol;Acc:MGI:6848243] | -1.11 | 2.65E-02 |
| Zfp784 | zinc finger protein 784 [Source:MGI Symbol;Acc:MGI:3606042] | -1.80 | 2.65E-02 |
| Skint10 | selection and upkeep of intraepithelial T cells 10 [Source:MGI Symbol;Acc:MGI:2685416] | -2.34 | 2.66E-02 |
| Agr3 | anterior gradient 3 [Source:MGI Symbol;Acc:MGI:2685734] | -3.99 | 2.66E-02 |
| Cd34 | CD34 antigen [Source:MGI Symbol;Acc:MGI:88329] | -1.03 | 2.66E-02 |
| Ear2 | eosinophil-associated, ribonuclease A family, member 2 [Source:MGI Symbol;Acc:MGI:108020] | -2.33 | 2.67E-02 |
| Gm29394 | predicted gene 29394 [Source:MGI Symbol;Acc:MGI:5580100] | -0.98 | 2.68E-02 |
| Zfp607a | zinc finger protein 607A [Source:MGI Symbol;Acc:MGI:3584526] | -0.61 | 2.69E-02 |
| Or1f19 | olfactory receptor family 1 subfamily F member 19 [Source:MGI Symbol;Acc:MGI:3032605] | -0.73 | 2.70E-02 |
| Fam13a | family with sequence similarity 13, member A [Source:MGI Symbol;Acc:MGI:1889842] | -1.43 | 2.70E-02 |
| Ppil6 | peptidylprolyl isomerase (cyclophilin)-like 6 [Source:MGI Symbol;Acc:MGI:1920325] | -1.73 | 2.70E-02 |
| Lacc1 | laccase domain containing 1 [Source:MGI Symbol;Acc:MGI:2445077] | -0.62 | 2.71E-02 |
| Mblac2 | metallo-beta-lactamase domain containing 2 [Source:MGI Symbol;Acc:MGI:1920102] | -0.95 | 2.71E-02 |
| Itih6 | inter-alpha-trypsin inhibitor heavy chain family member 6 [Source:MGI Symbol;Acc:MGI:2685232] | -2.65 | 2.72E-02 |
| Or1e1f | olfactory receptor family 1 subfamily E member 1F [Source:MGI Symbol;Acc:MGI:3030231] | -0.97 | 2.72E-02 |
| Zfp991 | zinc finger protein 991 [Source:MGI Symbol;Acc:MGI:3701604] | -0.92 | 2.74E-02 |
| Kctd16 | potassium channel tetramerisation domain containing 16 [Source:MGI Symbol;Acc:MGI:1914659] | -1.32 | 2.74E-02 |
| Gm8222 | predicted gene 8222 [Source:MGI Symbol;Acc:MGI:3643833] | -1.75 | 2.74E-02 |
| Exph5 | exophilin 5 [Source:MGI Symbol;Acc:MGI:2443248] | -1.26 | 2.75E-02 |
| Scn7a | sodium channel, voltage-gated, type VII, alpha [Source:MGI Symbol;Acc:MGI:102965] | -1.06 | 2.77E-02 |
| Tmie | transmembrane inner ear [Source:MGI Symbol;Acc:MGI:2159400] | -1.70 | 2.77E-02 |
| Ezh1 | enhancer of zeste 1 polycomb repressive complex 2 subunit [Source:MGI Symbol;Acc:MGI:1097695] | -0.60 | 2.78E-02 |
| Or2j6 | olfactory receptor family 2 subfamily J member 6 [Source:MGI Symbol;Acc:MGI:3030365] | -0.85 | 2.79E-02 |
| Slc16a10 | solute carrier family 16 (monocarboxylic acid transporters), member 10 [Source:MGI Symbol;Acc:MGI:1919722] | -0.87 | 2.79E-02 |
| Bco1 | beta-carotene oxygenase 1 [Source:MGI Symbol;Acc:MGI:1926923] | -1.01 | 2.82E-02 |
| Frem2 | Fras1 related extracellular matrix protein 2 [Source:MGI Symbol;Acc:MGI:2444465] | -1.84 | 2.82E-02 |
| Or11a3-ps1 | olfactory receptor family 11 subfamily A member 3, pseudogene 1 [Source:MGI Symbol;Acc:MGI:3030589] | -2.62 | 2.82E-02 |
| Nr3c1 | nuclear receptor subfamily 3, group C, member 1 [Source:MGI Symbol;Acc:MGI:95824] | -0.74 | 2.83E-02 |
| Rasgrp1 | RAS guanyl releasing protein 1 [Source:MGI Symbol;Acc:MGI:1314635] | -1.43 | 2.83E-02 |
| Dusp26 | dual specificity phosphatase 26 [Source:MGI Symbol;Acc:MGI:1914209] | -2.17 | 2.83E-02 |
| Ccdc80 | coiled-coil domain containing 80 [Source:MGI Symbol;Acc:MGI:1915146] | -1.07 | 2.83E-02 |
| Gm13683 | predicted gene 13683 [Source:MGI Symbol;Acc:MGI:3650118] | -1.23 | 2.85E-02 |
| Dlc1 | deleted in liver cancer 1 [Source:MGI Symbol;Acc:MGI:1354949] | -0.79 | 2.86E-02 |
| Zfp974 | zinc finger protein 974 [Source:MGI Symbol;Acc:MGI:1920680] | -0.65 | 2.86E-02 |
| Micu3 | mitochondrial calcium uptake family, member 3 [Source:MGI Symbol;Acc:MGI:1925756] | -0.85 | 2.88E-02 |
| Hint3 | histidine triad nucleotide binding protein 3 [Source:MGI Symbol;Acc:MGI:1914097] | -0.72 | 2.88E-02 |
| Gm8437 | predicted gene 8437 [Source:MGI Symbol;Acc:MGI:3642987] | -2.62 | 2.89E-02 |
| Tmem230 | transmembrane protein 230 [Source:MGI Symbol;Acc:MGI:1917862] | -1.02 | 2.91E-02 |
| Sugct | succinyl-CoA glutarate-CoA transferase [Source:MGI Symbol;Acc:MGI:1923221] | -1.07 | 2.91E-02 |
| Gm18956 | predicted gene, 18956 [Source:MGI Symbol;Acc:MGI:5011141] | -5.04 | 2.94E-02 |
| Or7c19 | olfactory receptor family 7 subfamily C member 19 [Source:MGI Symbol;Acc:MGI:3030205] | -1.33 | 2.95E-02 |
| Cwf19l2 | CWF19 like cell cycle control factor 2 [Source:MGI Symbol;Acc:MGI:1918023] | -0.74 | 2.96E-02 |
| Gm7031 | predicted gene 7031 [Source:MGI Symbol;Acc:MGI:3646638] | -4.56 | 2.97E-02 |
| Vmn2r75 | vomeronasal 2, receptor 75 [Source:MGI Symbol;Acc:MGI:3648311] | -0.88 | 2.99E-02 |
| Gstt1 | glutathione S-transferase, theta 1 [Source:MGI Symbol;Acc:MGI:107379] | -1.10 | 2.99E-02 |
| Cyp4f37 | cytochrome P450, family 4, subfamily f, polypeptide 37 [Source:MGI Symbol;Acc:MGI:3780112] | -2.04 | 2.99E-02 |
| Prrt2 | proline-rich transmembrane protein 2 [Source:MGI Symbol;Acc:MGI:1916267] | -0.80 | 3.00E-02 |
| Or4a66 | olfactory receptor family 4 subfamily A member 66 [Source:MGI Symbol;Acc:MGI:3031030] | -0.82 | 3.03E-02 |
| Frmd3 | FERM domain containing 3 [Source:MGI Symbol;Acc:MGI:2442466] | -1.07 | 3.03E-02 |
| Hnmt | histamine N-methyltransferase [Source:MGI Symbol;Acc:MGI:2153181] | -1.32 | 3.03E-02 |

| | | | |
|---|---|---|---|
| Or10j3b | olfactory receptor family 10 subfamily J member 3B [Source:MGI Symbol;Acc:MGI:3031238] | -1.47 | 3.04E-02 |
| Snca | synuclein, alpha [Source:MGI Symbol;Acc:MGI:1277151] | -1.85 | 3.04E-02 |
| Cib2 | calcium and integrin binding family member 2 [Source:MGI Symbol;Acc:MGI:1929293] | -1.08 | 3.04E-02 |
| Trdv4 | T cell receptor delta variable 4 [Source:MGI Symbol;Acc:MGI:4887419] | -1.26 | 3.04E-02 |
| Prss3b | serine protease 3B [Source:MGI Symbol;Acc:MGI:1914623] | -0.88 | 3.06E-02 |
| Atp6v1g3 | ATPase, H+ transporting, lysosomal V1 subunit G3 [Source:MGI Symbol;Acc:MGI:2450548] | -1.48 | 3.06E-02 |
| Pygo1 | pygopus 1 [Source:MGI Symbol;Acc:MGI:1919385] | -1.02 | 3.06E-02 |
| Or10ag57 | olfactory receptor family 10 subfamily AG member 57 [Source:MGI Symbol;Acc:MGI:3030956] | -0.89 | 3.09E-02 |
| Or14s1-ps1 | olfactory receptor family 14 subfamily S member 1, pseudogene 1 [Source:MGI Symbol;Acc:MGI:3030592] | -5.13 | 3.10E-02 |
| Skint8 | selection and upkeep of intraepithelial T cells 8 [Source:MGI Symbol;Acc:MGI:3651523] | -1.32 | 3.11E-02 |
| Kcnk2 | potassium channel, subfamily K, member 2 [Source:MGI Symbol;Acc:MGI:109366] | -0.75 | 3.11E-02 |
| Aspa | aspartoacylase [Source:MGI Symbol;Acc:MGI:87914] | -1.06 | 3.11E-02 |
| Tut4 | terminal uridylyl transferase 4 [Source:MGI Symbol;Acc:MGI:2445126] | -0.97 | 3.12E-02 |
| Car3 | carbonic anhydrase 3 [Source:MGI Symbol;Acc:MGI:88270] | -2.12 | 3.12E-02 |
| Hnrnph3 | heterogeneous nuclear ribonucleoprotein H3 [Source:MGI Symbol;Acc:MGI:1926462] | -0.66 | 3.12E-02 |
| Ankrd26 | ankyrin repeat domain 26 [Source:MGI Symbol;Acc:MGI:1917887] | -0.80 | 3.13E-02 |
| Elmod1 | ELMO/CED-12 domain containing 1 [Source:MGI Symbol;Acc:MGI:3583900] | -1.35 | 3.13E-02 |
| Negr1 | neuronal growth regulator 1 [Source:MGI Symbol;Acc:MGI:2444846] | -1.08 | 3.13E-02 |
| Tmem177 | transmembrane protein 177 [Source:MGI Symbol;Acc:MGI:1913593] | -1.28 | 3.14E-02 |
| Or8h7 | olfactory receptor family 8 subfamily H member 7 [Source:MGI Symbol;Acc:MGI:3030931] | -1.41 | 3.16E-02 |
| Capsl | calcyphosine-like [Source:MGI Symbol;Acc:MGI:1922818] | -1.61 | 3.16E-02 |
| Gm8895 | predicted gene 8895 [Source:MGI Symbol;Acc:MGI:3643891] | -3.18 | 3.16E-02 |
| Cyp2j7 | cytochrome P450, family 2, subfamily j, polypeptide 7 [Source:MGI Symbol;Acc:MGI:2449816] | -4.59 | 3.16E-02 |
| Zfp874a | zinc finger protein 874a [Source:MGI Symbol;Acc:MGI:3040703] | -0.70 | 3.16E-02 |
| Tdo2 | tryptophan 2,3-dioxygenase [Source:MGI Symbol;Acc:MGI:1928486] | -1.68 | 3.16E-02 |
| Wdr31 | WD repeat domain 31 [Source:MGI Symbol;Acc:MGI:1918604] | -0.92 | 3.17E-02 |
| Zfp935 | zinc finger protein 935 [Source:MGI Symbol;Acc:MGI:1918758] | -0.74 | 3.18E-02 |
| Mfsd14b | major facilitator superfamily domain containing 14B [Source:MGI Symbol;Acc:MGI:1913881] | -0.76 | 3.18E-02 |
| Il6ra | interleukin 6 receptor, alpha [Source:MGI Symbol;Acc:MGI:105304] | -0.99 | 3.19E-02 |
| Adgrg6 | adhesion G protein-coupled receptor G6 [Source:MGI Symbol;Acc:MGI:1916151] | -1.00 | 3.19E-02 |
| Or4e5 | olfactory receptor family 4 subfamily E member 5 [Source:MGI Symbol;Acc:MGI:3031341] | -2.31 | 3.19E-02 |
| Gm49228 | predicted gene, 49228 [Source:MGI Symbol;Acc:MGI:6118689] | -0.84 | 3.20E-02 |
| Hemgn | hemogen [Source:MGI Symbol;Acc:MGI:2136910] | -1.76 | 3.20E-02 |
| Slc7a6 | solute carrier family 7 (cationic amino acid transporter, y+ system), member 6 [Source:MGI Symbol;Acc:MGI:2142598] | -0.66 | 3.23E-02 |
| Rnf144b | ring finger protein 144B [Source:MGI Symbol;Acc:MGI:2384986] | -1.03 | 3.23E-02 |
| Adipoq | adiponectin, C1Q and collagen domain containing [Source:MGI Symbol;Acc:MGI:106675] | -2.20 | 3.24E-02 |
| Osgep | O-sialoglycoprotein endopeptidase [Source:MGI Symbol;Acc:MGI:1913496] | -0.70 | 3.24E-02 |
| Gm1110 | predicted gene 1110 [Source:MGI Symbol;Acc:MGI:2685956] | -3.41 | 3.27E-02 |
| Scel | sciellin [Source:MGI Symbol;Acc:MGI:1891228] | -1.47 | 3.27E-02 |
| Klb | klotho beta [Source:MGI Symbol;Acc:MGI:1932466] | -1.67 | 3.27E-02 |
| Vmn1r4 | vomeronasal 1 receptor 4 [Source:MGI Symbol;Acc:MGI:2159457] | -0.83 | 3.30E-02 |
| Slc22a8 | solute carrier family 22 (organic anion transporter), member 8 [Source:MGI Symbol;Acc:MGI:1336187] | -1.40 | 3.30E-02 |
| Cstpp1 | centriolar satellite-associated tubulin polyglutamylase complex regulator 1 [Source:MGI Symbol;Acc:MGI:1915079] | -0.81 | 3.30E-02 |
| Svip | small VCP/p97-interacting protein [Source:MGI Symbol;Acc:MGI:1922994] | -1.11 | 3.34E-02 |
| Nr1h5 | nuclear receptor subfamily 1, group H, member 5 [Source:MGI Symbol;Acc:MGI:3026618] | -0.97 | 3.38E-02 |
| Flacc1 | flagellum associated containing coiled-coil domains 1 [Source:MGI Symbol;Acc:MGI:1918359] | -1.30 | 3.41E-02 |
| Tmem200a | transmembrane protein 200A [Source:MGI Symbol;Acc:MGI:1924470] | -1.06 | 3.42E-02 |
| Dnajc6 | DnaJ heat shock protein family (Hsp40) member C6 [Source:MGI Symbol;Acc:MGI:1919935] | -1.24 | 3.43E-02 |
| Vmn2r77 | vomeronasal 2, receptor 77 [Source:MGI Symbol;Acc:MGI:3643879] | -1.26 | 3.49E-02 |
| Gpc5 | glypican 5 [Source:MGI Symbol;Acc:MGI:1194894] | -1.39 | 3.50E-02 |
| Srrm4 | serine/arginine repetitive matrix 4 [Source:MGI Symbol;Acc:MGI:1916205] | -1.08 | 3.51E-02 |
| Limch1 | LIM and calponin homology domains 1 [Source:MGI Symbol;Acc:MGI:1924819] | -1.30 | 3.51E-02 |
| Ssxb16 | SSX member B16 [Source:MGI Symbol;Acc:MGI:3642927] | -1.21 | 3.51E-02 |
| 4933411K16Rik | RIKEN cDNA 4933411K16 gene [Source:MGI Symbol;Acc:MGI:1914015] | -1.23 | 3.52E-02 |
| Map2k6 | mitogen-activated protein kinase kinase 6 [Source:MGI Symbol;Acc:MGI:1346870] | -1.36 | 3.52E-02 |
| Serpina3d-ps | serine (or cysteine) peptidase inhibitor, clade A, member 3D, pseudogene [Source:MGI Symbol;Acc:MGI:2182836] | -2.61 | 3.53E-02 |
| Gm14391 | predicted gene 14391 [Source:MGI Symbol;Acc:MGI:3709324] | -0.61 | 3.54E-02 |
| Zfp182 | zinc finger protein 182 [Source:MGI Symbol;Acc:MGI:2442220] | -0.79 | 3.54E-02 |
| Zfp169 | zinc finger protein 169 [Source:MGI Symbol;Acc:MGI:1915161] | -0.92 | 3.55E-02 |
| Gm48113 | predicted gene, 48113 [Source:MGI Symbol;Acc:MGI:6097466] | -1.48 | 3.55E-02 |
| Gm7591 | predicted gene 7591 [Source:MGI Symbol;Acc:MGI:3644074] | -0.98 | 3.56E-02 |
| Gm45871 | predicted gene 45871 [Source:MGI Symbol;Acc:MGI:5804986] | -0.67 | 3.57E-02 |
| Gm13615 | predicted gene 13615 [Source:MGI Symbol;Acc:MGI:3650493] | -0.84 | 3.60E-02 |
| Stag3 | STAG3 cohesin complex component [Source:MGI Symbol;Acc:MGI:1355311] | -0.99 | 3.60E-02 |
| Cnih3 | cornichon family AMPA receptor auxiliary protein 3 [Source:MGI Symbol;Acc:MGI:1920228] | -1.39 | 3.61E-02 |
| Gm5578 | predicted pseudogene 5578 [Source:MGI Symbol;Acc:MGI:3645010] | -2.18 | 3.61E-02 |
| Ift52 | intraflagellar transport 52 [Source:MGI Symbol;Acc:MGI:2387217] | -0.66 | 3.62E-02 |

| | | | |
|---|---|---|---|
| Pcolce2 | procollagen C-endopeptidase enhancer 2 [Source:MGI Symbol;Acc:MGI:1923727] | -1.68 | 3.62E-02 |
| Chpt1 | choline phosphotransferase 1 [Source:MGI Symbol;Acc:MGI:2384841] | -1.36 | 3.62E-02 |
| Trp53bp2 | transformation related protein 53 binding protein 2 [Source:MGI Symbol;Acc:MGI:2138319] | -0.71 | 3.62E-02 |
| Wt1 | WT1 transcription factor [Source:MGI Symbol;Acc:MGI:98968] | -0.72 | 3.62E-02 |
| Or51a10 | olfactory receptor family 51 subfamily A member 10 [Source:MGI Symbol;Acc:MGI:3030476] | -0.65 | 3.64E-02 |
| Myo9a | myosin IXa [Source:MGI Symbol;Acc:MGI:107735] | -0.97 | 3.64E-02 |
| Ptchd3 | patched domain containing 3 [Source:MGI Symbol;Acc:MGI:1921925] | -4.36 | 3.64E-02 |
| Zfp160 | zinc finger protein 160 [Source:MGI Symbol;Acc:MGI:108187] | -0.74 | 3.64E-02 |
| Sh3d19 | SH3 domain protein D19 [Source:MGI Symbol;Acc:MGI:1350923] | -0.70 | 3.70E-02 |
| Ehbp1 | EH domain binding protein 1 [Source:MGI Symbol;Acc:MGI:2667252] | -0.63 | 3.70E-02 |
| Ric3 | RIC3 acetylcholine receptor chaperone [Source:MGI Symbol;Acc:MGI:2443887] | -1.11 | 3.71E-02 |
| Ecm2 | extracellular matrix protein 2, female organ and adipocyte specific [Source:MGI Symbol;Acc:MGI:3039578] | -0.97 | 3.71E-02 |
| Dynlt3 | dynein light chain Tctex-type 3 [Source:MGI Symbol;Acc:MGI:1914367] | -0.88 | 3.71E-02 |
| Gm3508 | predicted gene 3508 [Source:MGI Symbol;Acc:MGI:3781685] | -0.93 | 3.77E-02 |
| Slc5a3 | solute carrier family 5 (inositol transporters), member 3 [Source:MGI Symbol;Acc:MGI:1858226] | -1.07 | 3.77E-02 |
| Gm8925 | predicted gene 8925 [Source:MGI Symbol;Acc:MGI:3643158] | -1.86 | 3.77E-02 |
| Or51b4 | olfactory receptor family 51 subfamily B member 4 [Source:MGI Symbol;Acc:MGI:1341906] | -0.65 | 3.78E-02 |
| Pcmtd1 | protein-L-isoaspartate (D-aspartate) O-methyltransferase domain containing 1 [Source:MGI Symbol;Acc:MGI:2441773] | -0.86 | 3.78E-02 |
| 4930519F16Rik | RIKEN cDNA 4930519F16 gene [Source:MGI Symbol;Acc:MGI:1922356] | -3.05 | 3.78E-02 |
| Btbd8 | BTB domain containing 8 [Source:MGI Symbol;Acc:MGI:3646208] | -0.94 | 3.81E-02 |
| C2cd4c | C2 calcium-dependent domain containing 4C [Source:MGI Symbol;Acc:MGI:2685084] | -0.98 | 3.81E-02 |
| Gabrr2 | gamma-aminobutyric acid type A receptor subunit rho 2 [Source:MGI Symbol;Acc:MGI:95626] | -3.14 | 3.81E-02 |
| AW554918 | expressed sequence AW554918 [Source:MGI Symbol;Acc:MGI:2147376] | -0.67 | 3.83E-02 |
| Veph1 | ventricular zone expressed PH domain-containing 1 [Source:MGI Symbol;Acc:MGI:1920039] | -0.92 | 3.83E-02 |
| Cdkl2 | cyclin dependent kinase like 2 [Source:MGI Symbol;Acc:MGI:1858227] | -0.77 | 3.83E-02 |
| Cdo1 | cysteine dioxygenase 1, cytosolic [Source:MGI Symbol;Acc:MGI:105925] | -1.69 | 3.83E-02 |
| Rnase6 | ribonuclease, RNase A family, 6 [Source:MGI Symbol;Acc:MGI:1925666] | -1.28 | 3.85E-02 |
| Ptn | pleiotrophin [Source:MGI Symbol;Acc:MGI:97804] | -1.07 | 3.86E-02 |
| Gm13328 | predicted gene 13328 [Source:MGI Symbol;Acc:MGI:3651185] | -4.55 | 3.86E-02 |
| Ankhd1 | ankyrin repeat and KH domain containing 1 [Source:MGI Symbol;Acc:MGI:1921733] | -0.61 | 3.90E-02 |
| Mmut | methylmalonyl-Coenzyme A mutase [Source:MGI Symbol;Acc:MGI:97239] | -0.86 | 3.90E-02 |
| Pou6f1 | POU domain, class 6, transcription factor 1 [Source:MGI Symbol;Acc:MGI:102935] | -0.78 | 3.90E-02 |
| Hibch | 3-hydroxyisobutyryl-Coenzyme A hydrolase [Source:MGI Symbol;Acc:MGI:1923792] | -0.99 | 3.90E-02 |
| Cavin2 | caveolae associated 2 [Source:MGI Symbol;Acc:MGI:99513] | -1.23 | 3.90E-02 |
| Zfp950 | zinc finger protein 950 [Source:MGI Symbol;Acc:MGI:2652824] | -0.69 | 3.92E-02 |
| Trpv6 | transient receptor potential cation channel, subfamily V, member 6 [Source:MGI Symbol;Acc:MGI:1927259] | -1.20 | 3.92E-02 |
| Atrnl1 | attractin like 1 [Source:MGI Symbol;Acc:MGI:2147749] | -0.61 | 3.93E-02 |
| Pard3b | par-3 family cell polarity regulator beta [Source:MGI Symbol;Acc:MGI:1919301] | -0.83 | 3.93E-02 |
| Adamts14 | ADAM metallopeptidase with thrombospondin type 1 motif 14 [Source:MGI Symbol;Acc:MGI:2179942] | -0.84 | 3.94E-02 |
| Slitrk6 | SLIT and NTRK-like family, member 6 [Source:MGI Symbol;Acc:MGI:2443198] | -1.46 | 3.94E-02 |
| Mtm1 | X-linked myotubular myopathy gene 1 [Source:MGI Symbol;Acc:MGI:1099452] | -0.79 | 3.95E-02 |
| Eps8l3 | EPS8-like 3 [Source:MGI Symbol;Acc:MGI:2139743] | -1.24 | 3.97E-02 |
| Or51v14 | olfactory receptor family 51 subfamily V member 14 [Source:MGI Symbol;Acc:MGI:3030454] | -1.09 | 4.01E-02 |
| Or1o4 | olfactory receptor family 1 subfamily O member 4 [Source:MGI Symbol;Acc:MGI:2177482] | -1.46 | 4.07E-02 |
| Gm20541 | predicted gene 20541 [Source:MGI Symbol;Acc:MGI:5142006] | -2.34 | 4.08E-02 |
| Dnhd1 | dynein heavy chain domain 1 [Source:MGI Symbol;Acc:MGI:1924755] | -1.18 | 4.09E-02 |
| Or7g19 | olfactory receptor family 7 subfamily G member 19 [Source:MGI Symbol;Acc:MGI:3030666] | -1.29 | 4.10E-02 |
| H2-M6-ps | histocompatibility 2, M region locus 6, pseudogene [Source:MGI Symbol;Acc:MGI:95918] | -3.38 | 4.12E-02 |
| Prss48 | serine protease 48 [Source:MGI Symbol;Acc:MGI:2685865] | -1.11 | 4.13E-02 |
| Mrgprb9-ps | MAS-related GPR, member B9, pseudogene [Source:MGI Symbol;Acc:MGI:3033133] | -2.62 | 4.13E-02 |
| Gnpat | glyceronephosphate O-acyltransferase [Source:MGI Symbol;Acc:MGI:1343460] | -0.60 | 4.13E-02 |
| Akap12 | A kinase anchor protein 12 [Source:MGI Symbol;Acc:MGI:1932576] | -1.18 | 4.13E-02 |
| Gm11633 | predicted gene 11633 [Source:MGI Symbol;Acc:MGI:3650676] | -1.41 | 4.14E-02 |
| Vmn2r27 | vomeronasal 2, receptor27 [Source:MGI Symbol;Acc:MGI:3761517] | -0.95 | 4.14E-02 |
| Or2b7 | olfactory receptor family 2 subfamily B member 7 [Source:MGI Symbol;Acc:MGI:3031369] | -1.05 | 4.15E-02 |
| Synpo2 | synaptopodin 2 [Source:MGI Symbol;Acc:MGI:2153070] | -1.74 | 4.15E-02 |
| Gm57488 | predicted gene, 57488 [Source:MGI Symbol;Acc:MGI:7511940] | -1.58 | 4.16E-02 |
| Topaz1 | testis and ovary specific PAZ domain containing 1 [Source:MGI Symbol;Acc:MGI:3779933] | -4.93 | 4.18E-02 |
| Zfp418 | zinc finger protein 418 [Source:MGI Symbol;Acc:MGI:2444763] | -0.80 | 4.19E-02 |
| Kyat3 | kynurenine aminotransferase 3 [Source:MGI Symbol;Acc:MGI:2677849] | -0.79 | 4.21E-02 |
| Or5au1 | olfactory receptor family 5 subfamily AU member 1 [Source:MGI Symbol;Acc:MGI:3030055] | -1.16 | 4.22E-02 |
| BC085271 | cDNA sequence BC085271 [Source:MGI Symbol;Acc:MGI:3612444] | -1.60 | 4.22E-02 |
| Grb14 | growth factor receptor bound protein 14 [Source:MGI Symbol;Acc:MGI:1355324] | -1.33 | 4.22E-02 |
| Gm47759 | predicted gene, 47759 [Source:MGI Symbol;Acc:MGI:6096909] | -1.55 | 4.23E-02 |
| Gbe1 | 1,4-alpha-glucan branching enzyme 1 [Source:MGI Symbol;Acc:MGI:1921435] | -0.89 | 4.24E-02 |
| Mlh1 | mutL homolog 1 [Source:MGI Symbol;Acc:MGI:101938] | -0.64 | 4.25E-02 |
| Echdc3 | enoyl Coenzyme A hydratase domain containing 3 [Source:MGI Symbol;Acc:MGI:1915106] | -1.24 | 4.25E-02 |
| Hoxa4 | homeobox A4 [Source:MGI Symbol;Acc:MGI:96176] | -1.14 | 4.29E-02 |
| Syne2 | spectrin repeat containing, nuclear envelope 2 [Source:MGI Symbol;Acc:MGI:2449316] | -0.96 | 4.29E-02 |

| | | | |
|---|---|---|---|
| Ctso | cathepsin O [Source:MGI Symbol;Acc:MGI:2139628] | -0.90 | 4.31E-02 |
| Clca2 | chloride channel accessory 2 [Source:MGI Symbol;Acc:MGI:2139758] | -1.63 | 4.32E-02 |
| Vmn2r69 | vomeronasal 2, receptor 69 [Source:MGI Symbol;Acc:MGI:3761311] | -0.59 | 4.34E-02 |
| Meiob | meiosis specific with OB domains [Source:MGI Symbol;Acc:MGI:1922428] | -1.91 | 4.34E-02 |
| Zfp3 | zinc finger protein 3 [Source:MGI Symbol;Acc:MGI:99177] | -0.88 | 4.36E-02 |
| Lonp2 | lon peptidase 2, peroxisomal [Source:MGI Symbol;Acc:MGI:1914137] | -0.59 | 4.38E-02 |
| Sbk2 | SH3-binding domain kinase family, member 2 [Source:MGI Symbol;Acc:MGI:2685925] | -4.15 | 4.38E-02 |
| Or4d10c | olfactory receptor family 4 subfamily D member 10C [Source:MGI Symbol;Acc:MGI:3031260] | -2.08 | 4.39E-02 |
| Or5bb12 | olfactory receptor family 5 subfamily BB member 12 [Source:MGI Symbol;Acc:MGI:3031389] | -0.79 | 4.39E-02 |
| Per3 | period circadian clock 3 [Source:MGI Symbol;Acc:MGI:1277134] | -0.82 | 4.40E-02 |
| Gm47355 | predicted gene, 47355 [Source:MGI Symbol;Acc:MGI:6096260] | -0.60 | 4.41E-02 |
| 9330159F19Rik | RIKEN cDNA 9330159F19 gene [Source:MGI Symbol;Acc:MGI:3036239] | -1.94 | 4.41E-02 |
| Or14q1-ps1 | olfactory receptor family 14 subfamily Q member 1, pseudogene 1 [Source:MGI Symbol;Acc:MGI:3030590] | -1.90 | 4.43E-02 |
| Gpr141b | G protein-coupled receptor 141B [Source:MGI Symbol;Acc:MGI:2441809] | -0.81 | 4.44E-02 |
| Cyp21a2-ps | cytochrome P450, family 21, subfamily a, polypeptide 2 pseudogene [Source:MGI Symbol;Acc:MGI:3645529] | -1.41 | 4.46E-02 |
| Bcl6 | B cell leukemia/lymphoma 6 [Source:MGI Symbol;Acc:MGI:107187] | -0.59 | 4.47E-02 |
| Rrm2b | ribonucleotide reductase M2 B (TP53 inducible) [Source:MGI Symbol;Acc:MGI:2155865] | -0.65 | 4.47E-02 |
| Lrrfip2 | leucine rich repeat (in FLII) interacting protein 2 [Source:MGI Symbol;Acc:MGI:1918518] | -0.60 | 4.48E-02 |
| Hoxa2 | homeobox A2 [Source:MGI Symbol;Acc:MGI:96174] | -1.02 | 4.48E-02 |
| Mybl1 | myeloblastosis oncogene-like 1 [Source:MGI Symbol;Acc:MGI:99925] | -0.63 | 4.49E-02 |
| Slc7a13 | solute carrier family 7, (cationic amino acid transporter, y+ system) member 13 [Source:MGI Symbol;Acc:MGI:1921337] | -1.30 | 4.50E-02 |
| Paqr5 | progestin and adipoQ receptor family member V [Source:MGI Symbol;Acc:MGI:1921340] | -1.21 | 4.51E-02 |
| Or8j3b | olfactory receptor family 8 subfamily J member 3B [Source:MGI Symbol;Acc:MGI:3030891] | -0.96 | 4.51E-02 |
| Med14 | mediator complex subunit 14 [Source:MGI Symbol;Acc:MGI:1349442] | -0.66 | 4.51E-02 |
| Msantd2 | Myb/SANT-like DNA-binding domain containing 2 [Source:MGI Symbol;Acc:MGI:2384579] | -0.59 | 4.51E-02 |
| Or4c12 | olfactory receptor family 4 subfamily C member 12 [Source:MGI Symbol;Acc:MGI:3031093] | -1.45 | 4.53E-02 |
| Zfp654 | zinc finger protein 654 [Source:MGI Symbol;Acc:MGI:1919270] | -0.67 | 4.58E-02 |
| Cfap300 | cilia and flagella associated protein 300 [Source:MGI Symbol;Acc:MGI:3045346] | -0.63 | 4.59E-02 |
| Laptm4b | lysosomal-associated protein transmembrane 4B [Source:MGI Symbol;Acc:MGI:1890494] | -0.68 | 4.59E-02 |
| Gm12584 | predicted gene 12584 [Source:MGI Symbol;Acc:MGI:3651486] | -1.37 | 4.61E-02 |
| Rufy1 | RUN and FYVE domain containing 1 [Source:MGI Symbol;Acc:MGI:2429762] | -0.59 | 4.65E-02 |
| Prf1 | perforin 1 (pore forming protein) [Source:MGI Symbol;Acc:MGI:97551] | -1.12 | 4.67E-02 |
| Nkap | NFKB activating protein [Source:MGI Symbol;Acc:MGI:1914300] | -0.81 | 4.70E-02 |
| Lnpep | leucyl/cystinyl aminopeptidase [Source:MGI Symbol;Acc:MGI:2387123] | -0.66 | 4.71E-02 |
| Slc9a7 | solute carrier family 9 (sodium/hydrogen exchanger), member 7 [Source:MGI Symbol;Acc:MGI:2444530] | -0.74 | 4.71E-02 |
| Zfp990 | zinc finger protein 990 [Source:MGI Symbol;Acc:MGI:3652161] | -1.68 | 4.74E-02 |
| Paqr9 | progestin and adipoQ receptor family member IX [Source:MGI Symbol;Acc:MGI:1922802] | -1.81 | 4.74E-02 |
| Gm6918 | predicted gene 6918 [Source:MGI Symbol;Acc:MGI:3643700] | -5.14 | 4.75E-02 |
| Or13p4 | olfactory receptor family 13 subfamily P member 4 [Source:MGI Symbol;Acc:MGI:3031176] | -1.10 | 4.82E-02 |
| Gm18353 | predicted gene, 18353 [Source:MGI Symbol;Acc:MGI:5010538] | -2.59 | 4.86E-02 |
| Cimap1c | ciliary microtubule associated protein 1C [Source:MGI Symbol;Acc:MGI:2681875] | -2.40 | 4.87E-02 |
| Cyld | CYLD lysine 63 deubiquitinase [Source:MGI Symbol;Acc:MGI:1921506] | -0.71 | 4.89E-02 |
| Xcl1 | chemokine (C motif) ligand 1 [Source:MGI Symbol;Acc:MGI:104593] | -2.33 | 4.91E-02 |
| Cited2 | Cbp/p300-interacting transactivator, with Glu/Asp-rich carboxy-terminal domain, 2 [Source:MGI Symbol;Acc:MGI:1306784] | -0.88 | 4.95E-02 |
| Hoxa3 | homeobox A3 [Source:MGI Symbol;Acc:MGI:96175] | -1.12 | 4.95E-02 |
| Or12e14 | olfactory receptor family 12 subfamily E member 14 [Source:MGI Symbol;Acc:MGI:3030984] | -0.68 | 4.95E-02 |
| Or2g25 | olfactory receptor family 2 subfamily G member 25 [Source:MGI Symbol;Acc:MGI:2177500] | -1.10 | 4.95E-02 |
| Gm32256 | predicted gene, 32256 [Source:MGI Symbol;Acc:MGI:5591415] | -1.25 | 4.95E-02 |
| Optc | opticin [Source:MGI Symbol;Acc:MGI:2151113] | -1.31 | 4.95E-02 |
| Reps2 | RALBP1 associated Eps domain containing protein 2 [Source:MGI Symbol;Acc:MGI:2663511] | -1.15 | 4.96E-02 |
| Scara5 | scavenger receptor class A, member 5 [Source:MGI Symbol;Acc:MGI:1918395] | -1.17 | 4.97E-02 |
| Nudt12 | nudix hydrolase 12 [Source:MGI Symbol;Acc:MGI:1915243] | -0.85 | 4.97E-02 |
| Gls | glutaminase [Source:MGI Symbol;Acc:MGI:95752] | -0.67 | 4.98E-02 |
| Krt77 | keratin 77 [Source:MGI Symbol;Acc:MGI:3588209] | -2.18 | 4.98E-02 |
| Gm15078 | predicted gene 15078 [Source:MGI Symbol;Acc:MGI:3705413] | -2.63 | 4.99E-02 |
| Ttc8 | tetratricopeptide repeat domain 8 [Source:MGI Symbol;Acc:MGI:1923510] | -0.85 | 4.99E-02 |
| Gm20438 | predicted gene 20438 [Source:MGI Symbol;Acc:MGI:5141903] | -1.18 | 4.99E-02 |
| Spam1 | sperm adhesion molecule 1 [Source:MGI Symbol;Acc:MGI:109335] | -3.01 | 4.99E-02 |
| Vmn1r198 | vomeronasal 1 receptor 198 [Source:MGI Symbol;Acc:MGI:2159689] | -0.78 | 4.99E-02 |

### Table S3. Upregulated genes in FLASH-RT mice skin compared to CNT

| Gene ID | Name | log2(FC) | padj |
|---|---|---|---|
| Gm20784 | predicted gene, 20784 [Source:MGI Symbol;Acc:MGI:5434140] | 8.36 | 1.54E-07 |

| | | | |
|---|---|---|---|
| Gm12481 | predicted gene 12481 [Source:MGI Symbol;Acc:MGI:3650743] | 1.16 | 1.08E-04 |
| Krt6b | keratin 6B [Source:MGI Symbol;Acc:MGI:1333768] | 11.17 | 2.79E-04 |
| Mafg | v-maf musculoaponeurotic fibrosarcoma oncogene family, protein G (avian) [Source:MGI Symbol;Acc:MGI:96911] | 0.59 | 3.40E-04 |
| Akt1s1 | AKT1 substrate 1 [Source:MGI Symbol;Acc:MGI:1914855] | 0.88 | 1.21E-03 |
| Zxda | zinc finger, X-linked, duplicated A [Source:MGI Symbol;Acc:MGI:1921689] | 7.40 | 1.50E-03 |
| Akap17a | A-kinase anchoring protein 17A [Source:MGI Symbol;Acc:MGI:6723883] | 0.78 | 1.71E-03 |
| Stard3 | StAR related lipid transfer domain containing 3 [Source:MGI Symbol;Acc:MGI:1929618] | 0.71 | 2.38E-03 |
| Thbs4 | thrombospondin 4 [Source:MGI Symbol;Acc:MGI:1101779] | 2.20 | 2.84E-03 |
| Gm20683 | predicted gene 20683 [Source:MGI Symbol;Acc:MGI:5313130] | 1.21 | 4.55E-03 |
| Sac3d1 | SAC3 domain containing 1 [Source:MGI Symbol;Acc:MGI:1913656] | 1.06 | 4.63E-03 |
| Cstdc5 | cystatin domain containing 5 [Source:MGI Symbol;Acc:MGI:3696883] | 8.06 | 5.08E-03 |
| Gm20708 | predicted gene 20708 [Source:MGI Symbol;Acc:MGI:5313155] | 6.29 | 6.80E-03 |
| Ccl19-ps5 | C-C motif chemokine ligand 19, pseudogene 5 [Source:MGI Symbol;Acc:MGI:3693096] | 6.21 | 8.79E-03 |
| H60c | histocompatibility 60c [Source:MGI Symbol;Acc:MGI:3774845] | 0.62 | 8.79E-03 |
| Alpl | alkaline phosphatase, liver/bone/kidney [Source:MGI Symbol;Acc:MGI:87983] | 1.66 | 9.25E-03 |
| Tmem147 | transmembrane protein 147 [Source:MGI Symbol;Acc:MGI:1915011] | 0.83 | 9.25E-03 |
| Gm20547 | predicted gene 20547 [Source:MGI Symbol;Acc:MGI:5142012] | 5.89 | 1.12E-02 |
| Gpc1 | glypican 1 [Source:MGI Symbol;Acc:MGI:1194891] | 0.88 | 1.12E-02 |
| Slc2a8 | solute carrier family 2, (facilitated glucose transporter), member 8 [Source:MGI Symbol;Acc:MGI:1860103] | 1.06 | 1.46E-02 |
| Stfa1 | stefin A1 [Source:MGI Symbol;Acc:MGI:106198] | 6.91 | 1.50E-02 |
| Stfa3 | stefin A3 [Source:MGI Symbol;Acc:MGI:106196] | 6.06 | 1.50E-02 |
| Oaz1-ps | ornithine decarboxylase antizyme 1, pseudogene [Source:MGI Symbol;Acc:MGI:108188] | 1.25 | 1.50E-02 |
| Gm12895 | predicted gene 12895 [Source:MGI Symbol;Acc:MGI:3649907] | 2.82 | 1.51E-02 |
| Gm12896 | predicted gene 12896 [Source:MGI Symbol;Acc:MGI:3650312] | 2.82 | 1.51E-02 |
| Dpysl4 | dihydropyrimidinase-like 4 [Source:MGI Symbol;Acc:MGI:1349764] | 1.65 | 1.51E-02 |
| Rpp25l | ribonuclease P/MRP 25 subunit-like [Source:MGI Symbol;Acc:MGI:1917211] | 1.01 | 1.51E-02 |
| Uqcr11 | ubiquinol-cytochrome c reductase, complex III subunit XI [Source:MGI Symbol;Acc:MGI:1913844] | 1.03 | 1.52E-02 |
| Jdp2 | Jun dimerization protein 2 [Source:MGI Symbol;Acc:MGI:1932093] | 0.67 | 1.58E-02 |
| Map3k10 | mitogen-activated protein kinase kinase kinase 10 [Source:MGI Symbol;Acc:MGI:1346879] | 0.89 | 1.61E-02 |
| Tymp | thymidine phosphorylase [Source:MGI Symbol;Acc:MGI:1920212] | 1.14 | 2.30E-02 |
| Flg | filaggrin [Source:MGI Symbol;Acc:MGI:95553] | 2.62 | 2.64E-02 |
| Gsdmc2 | gasdermin C2 [Source:MGI Symbol;Acc:MGI:2146102] | 2.45 | 3.00E-02 |
| Cfap157 | cilia and flagella associated protein 157 [Source:MGI Symbol;Acc:MGI:2447809] | 0.99 | 3.89E-02 |
| Abhd17a | abhydrolase domain containing 17A [Source:MGI Symbol;Acc:MGI:106388] | 0.88 | 3.98E-02 |
| Atp5mc2 | ATP synthase membrane subunit c locus 2 [Source:MGI Symbol;Acc:MGI:1915192] | 0.64 | 3.98E-02 |
| Slpi | secretory leukocyte peptidase inhibitor [Source:MGI Symbol;Acc:MGI:109297] | 4.68 | 4.41E-02 |
| Ecel1 | endothelin converting enzyme-like 1 [Source:MGI Symbol;Acc:MGI:1343461] | 2.99 | 4.54E-02 |
| Ddx28 | DEAD box helicase 28 [Source:MGI Symbol;Acc:MGI:1919236] | 0.90 | 4.69E-02 |
| Rnf187 | ring finger protein 187 [Source:MGI Symbol;Acc:MGI:1914224] | 0.68 | 4.69E-02 |
| Pygo2 | pygopus 2 [Source:MGI Symbol;Acc:MGI:1916161] | 0.70 | 4.78E-02 |
| Mrps26 | mitochondrial ribosomal protein S26 [Source:MGI Symbol;Acc:MGI:1333830] | 0.61 | 4.94E-02 |

### Table S4. Downregulated genes in FLASH-RT mice skin compared to CNT

| Gene ID | Name | log2(FC) | padj |
|---|---|---|---|
| Gm47893 | predicted gene, 47893 [Source:MGI Symbol;Acc:MGI:6097126] | -3.70 | 2.49E-08 |
| Ptger3 | prostaglandin E receptor 3 (subtype EP3) [Source:MGI Symbol;Acc:MGI:97795] | -1.89 | 5.50E-05 |
| Slc5a7 | solute carrier family 5 (choline transporter), member 7 [Source:MGI Symbol;Acc:MGI:1927126] | -2.34 | 7.16E-05 |
| Gm17043 | predicted gene 17043 [Source:MGI Symbol;Acc:MGI:4937870] | -3.83 | 2.79E-04 |
| Pcolce2 | procollagen C-endopeptidase enhancer 2 [Source:MGI Symbol;Acc:MGI:1923727] | -1.24 | 2.79E-04 |
| Gm15519 | predicted gene 15519 [Source:MGI Symbol;Acc:MGI:3782965] | -3.05 | 5.15E-04 |
| Aifm2 | apoptosis-inducing factor, mitochondrion-associated 2 [Source:MGI Symbol;Acc:MGI:1918611] | -0.95 | 1.50E-03 |
| Gm21378 | predicted gene, 21378 [Source:MGI Symbol;Acc:MGI:5434733] | -1.28 | 2.38E-03 |
| Lep | leptin [Source:MGI Symbol;Acc:MGI:104663] | -3.00 | 2.43E-03 |
| Trarg1 | trafficking regulator of GLUT4 (SLC2A4) 1 [Source:MGI Symbol;Acc:MGI:3029307] | -2.13 | 3.29E-03 |
| Gm8895 | predicted gene 8895 [Source:MGI Symbol;Acc:MGI:3643891] | -2.97 | 4.37E-03 |
| Cdo1 | cysteine dioxygenase 1, cytosolic [Source:MGI Symbol;Acc:MGI:105925] | -2.02 | 4.48E-03 |
| Serpina3c | serine (or cysteine) peptidase inhibitor, clade A, member 3C [Source:MGI Symbol;Acc:MGI:102848] | -1.52 | 5.64E-03 |
| Rab6b | RAB6B, member RAS oncogene family [Source:MGI Symbol;Acc:MGI:107283] | -0.93 | 5.64E-03 |
| Lctl | lactase-like [Source:MGI Symbol;Acc:MGI:2183549] | -2.24 | 6.80E-03 |
| Col4a4 | collagen, type IV, alpha 4 [Source:MGI Symbol;Acc:MGI:104687] | -0.72 | 1.20E-02 |
| Potefam3e | POTE ankyrin domain family member 3E [Source:MGI Symbol;Acc:MGI:1923056] | -5.88 | 1.34E-02 |
| Acvr1c | activin A receptor, type IC [Source:MGI Symbol;Acc:MGI:2661081] | -2.00 | 1.40E-02 |
| Slc1a3 | solute carrier family 1 (glial high affinity glutamate transporter), member 3 [Source:MGI Symbol;Acc:MGI:99917] | -0.84 | 1.50E-02 |
| Serpina1a | serine (or cysteine) peptidase inhibitor, clade A, member 1A [Source:MGI Symbol;Acc:MGI:891971] | -2.39 | 1.51E-02 |
| Slc22a3 | solute carrier family 22 (organic cation transporter), member 3 [Source:MGI Symbol;Acc:MGI:1333817] | -1.40 | 1.51E-02 |
| Mpz | myelin protein zero [Source:MGI Symbol;Acc:MGI:103177] | -1.18 | 1.51E-02 |
| Oxtr | oxytocin receptor [Source:MGI Symbol;Acc:MGI:109147] | -1.11 | 1.51E-02 |

| | | | |
|---|---|---|---|
| Slc36a2 | solute carrier family 36 (proton/amino acid symporter), member 2 [Source:MGI Symbol;Acc:MGI:1891430] | -1.08 | 1.52E-02 |
| Serpina1c | serine (or cysteine) peptidase inhibitor, clade A, member 1C [Source:MGI Symbol;Acc:MGI:891969] | -4.32 | 1.61E-02 |
| Prr32 | proline rich 32 [Source:MGI Symbol;Acc:MGI:1916050] | -3.18 | 1.71E-02 |
| Efemp1 | epidermal growth factor-containing fibulin-like extracellular matrix protein 1 [Source:MGI Symbol;Acc:MGI:1339998] | -1.06 | 1.88E-02 |
| Ccdc122 | coiled-coil domain containing 122 [Source:MGI Symbol;Acc:MGI:1918358] | -0.74 | 1.99E-02 |
| Cyp2b19 | cytochrome P450, family 2, subfamily b, polypeptide 19 [Source:MGI Symbol;Acc:MGI:107303] | -0.70 | 1.99E-02 |
| Plin1 | perilipin 1 [Source:MGI Symbol;Acc:MGI:1890505] | -2.32 | 2.13E-02 |
| Tshr | thyroid stimulating hormone receptor [Source:MGI Symbol;Acc:MGI:98849] | -1.77 | 2.40E-02 |
| Gm3188 | predicted gene 3188 [Source:MGI Symbol;Acc:MGI:3781367] | -0.84 | 2.41E-02 |
| Cimip2a | ciliary microtubule inner protein 2A [Source:MGI Symbol;Acc:MGI:3605773] | -1.47 | 2.62E-02 |
| Fmr1 | fragile X messenger ribonucleoprotein 1 [Source:MGI Symbol;Acc:MGI:95564] | -0.72 | 2.62E-02 |
| Gm13255 | predicted gene 13255 [Source:MGI Symbol;Acc:MGI:3652189] | -0.81 | 2.62E-02 |
| Tent2-ps1 | terminal nucleotidyltransferase 2, pseudogene 1 [Source:MGI Symbol;Acc:MGI:3801944] | -1.24 | 2.96E-02 |
| Vmn1r28 | vomeronasal 1 receptor 28 [Source:MGI Symbol;Acc:MGI:2159461] | -0.77 | 2.98E-02 |
| Oprl1 | opioid receptor-like 1 [Source:MGI Symbol;Acc:MGI:97440] | -3.91 | 3.12E-02 |
| Mmd | monocyte to macrophage differentiation-associated [Source:MGI Symbol;Acc:MGI:1914718] | -0.98 | 3.12E-02 |
| A530016L24Rik | RIKEN cDNA A530016L24 gene [Source:MGI Symbol;Acc:MGI:2443020] | -2.19 | 3.46E-02 |
| Pax5 | paired box 5 [Source:MGI Symbol;Acc:MGI:97489] | -4.00 | 3.89E-02 |
| Mgl2 | macrophage galactose N-acetyl-galactosamine specific lectin 2 [Source:MGI Symbol;Acc:MGI:2385729] | -1.09 | 3.89E-02 |
| Gdf10 | growth differentiation factor 10 [Source:MGI Symbol;Acc:MGI:95684] | -0.73 | 3.98E-02 |
| Or10g6 | olfactory receptor family 10 subfamily G member 6 [Source:MGI Symbol;Acc:MGI:3030815] | -0.78 | 4.24E-02 |
| Crp | C-reactive protein, pentraxin-related [Source:MGI Symbol;Acc:MGI:88512] | -0.62 | 4.24E-02 |
| Nkain3 | Na+/K+ transporting ATPase interacting 3 [Source:MGI Symbol;Acc:MGI:2444830] | -0.69 | 4.28E-02 |
| Ccdc80 | coiled-coil domain containing 80 [Source:MGI Symbol;Acc:MGI:1915146] | -1.19 | 4.50E-02 |
| Sncg | synuclein, gamma [Source:MGI Symbol;Acc:MGI:1298397] | -2.05 | 4.52E-02 |
| C7 | complement component 7 [Source:MGI Symbol;Acc:MGI:88235] | -1.91 | 4.65E-02 |
| Npr3 | natriuretic peptide receptor 3 [Source:MGI Symbol;Acc:MGI:97373] | -1.95 | 4.69E-02 |
| Nat8f3 | N-acetyltransferase 8 (GCN5-related) family member 3 [Source:MGI Symbol;Acc:MGI:2136449] | -0.68 | 4.94E-02 |

### Table S5. Upregulated genes in CONV-RT mice muscle compared to CNT

| Gene ID | Name | log2(FC) | padj |
|---|---|---|---|
| Glul | glutamate-ammonia ligase [Source:MGI Symbol;Acc:MGI:95739] | 1.58 | 1.66E-22 |
| Cdkn1a | cyclin dependent kinase inhibitor 1A [Source:MGI Symbol;Acc:MGI:104556] | 2.67 | 3.98E-15 |
| Lgi1 | leucine-rich repeat LGI family, member 1 [Source:MGI Symbol;Acc:MGI:1861691] | 3.85 | 7.17E-07 |
| Cryab | crystallin, alpha B [Source:MGI Symbol;Acc:MGI:88516] | 1.57 | 1.60E-06 |
| Pdlim1 | PDZ and LIM domain 1 (elfin) [Source:MGI Symbol;Acc:MGI:1860611] | 2.40 | 1.84E-06 |
| Pdk4 | pyruvate dehydrogenase kinase, isoenzyme 4 [Source:MGI Symbol;Acc:MGI:1351481] | 1.54 | 1.84E-06 |
| Kcnma1 | potassium large conductance calcium-activated channel, subfamily M, alpha member 1 [Source:MGI Symbol;Acc:MGI:99923] | 0.91 | 3.04E-06 |
| Fat1 | FAT atypical cadherin 1 [Source:MGI Symbol;Acc:MGI:109168] | 1.31 | 4.59E-06 |
| Frem2 | Fras1 related extracellular matrix protein 2 [Source:MGI Symbol;Acc:MGI:2444465] | 3.86 | 3.12E-05 |
| Ucp2 | uncoupling protein 2 (mitochondrial, proton carrier) [Source:MGI Symbol;Acc:MGI:109354] | 1.80 | 3.80E-05 |
| Hspa1b | heat shock protein 1B [Source:MGI Symbol;Acc:MGI:99517] | 2.73 | 3.91E-05 |
| Plin4 | perilipin 4 [Source:MGI Symbol;Acc:MGI:1929709] | 1.00 | 8.21E-05 |
| Abra | actin-binding Rho activating protein [Source:MGI Symbol;Acc:MGI:2444891] | 1.48 | 1.36E-04 |
| Zfp703 | zinc finger protein 703 [Source:MGI Symbol;Acc:MGI:2662729] | 1.25 | 1.73E-04 |
| Dusp18 | dual specificity phosphatase 18 [Source:MGI Symbol;Acc:MGI:1922469] | 1.97 | 2.15E-04 |
| Pank1 | pantothenate kinase 1 [Source:MGI Symbol;Acc:MGI:1922985] | 1.47 | 2.44E-04 |
| Cblb | Casitas B-lineage lymphoma b [Source:MGI Symbol;Acc:MGI:2146430] | 0.94 | 2.94E-04 |
| Ttc9 | tetratricopeptide repeat domain 9 [Source:MGI Symbol;Acc:MGI:1916730] | 2.72 | 3.87E-04 |
| Zbtb16 | zinc finger and BTB domain containing 16 [Source:MGI Symbol;Acc:MGI:103222] | 0.85 | 3.87E-04 |
| Apoe | apolipoprotein E [Source:MGI Symbol;Acc:MGI:88057] | 1.12 | 4.17E-04 |
| Cebpb | CCAAT/enhancer binding protein beta [Source:MGI Symbol;Acc:MGI:88373] | 1.22 | 4.28E-04 |
| Gramd1b | GRAM domain containing 1B [Source:MGI Symbol;Acc:MGI:1925037] | 1.41 | 4.93E-04 |
| Sorbs1 | sorbin and SH3 domain containing 1 [Source:MGI Symbol;Acc:MGI:700014] | 1.26 | 8.68E-04 |
| Nnt | nicotinamide nucleotide transhydrogenase [Source:MGI Symbol;Acc:MGI:109279] | 1.83 | 1.14E-03 |
| Mup22 | major urinary protein 22 [Source:MGI Symbol;Acc:MGI:5434675] | 6.08 | 1.21E-03 |
| Arhgap26 | Rho GTPase activating protein 26 [Source:MGI Symbol;Acc:MGI:1918552] | 1.27 | 1.21E-03 |
| Aldh2 | aldehyde dehydrogenase 2, mitochondrial [Source:MGI Symbol;Acc:MGI:99600] | 0.83 | 1.31E-03 |
| Plin3 | perilipin 3 [Source:MGI Symbol;Acc:MGI:1914155] | 1.12 | 1.41E-03 |
| Plin5 | perilipin 5 [Source:MGI Symbol;Acc:MGI:1914218] | 1.79 | 1.79E-03 |
| Zranb3 | zinc finger, RAN-binding domain containing 3 [Source:MGI Symbol;Acc:MGI:1918362] | 1.36 | 1.92E-03 |
| Adora1 | adenosine A1 receptor [Source:MGI Symbol;Acc:MGI:99401] | 1.86 | 1.99E-03 |
| Slc27a1 | solute carrier family 27 (fatty acid transporter), member 1 [Source:MGI Symbol;Acc:MGI:1347098] | 0.97 | 2.22E-03 |

| | | | |
|---|---|---|---|
| Ciapin1 | cytokine induced apoptosis inhibitor 1 [Source:MGI Symbol;Acc:MGI:1922083] | 1.25 | 2.39E-03 |
| Lrrc52 | leucine rich repeat containing 52 [Source:MGI Symbol;Acc:MGI:1924118] | 5.33 | 3.25E-03 |
| Pkn3 | protein kinase N3 [Source:MGI Symbol;Acc:MGI:2388285] | 1.79 | 3.44E-03 |
| Gk | glycerol kinase [Source:MGI Symbol;Acc:MGI:106594] | 3.38 | 3.50E-03 |
| Ccr2 | C-C motif chemokine receptor 2 [Source:MGI Symbol;Acc:MGI:106185] | 2.21 | 3.78E-03 |
| H2-Eb1 | histocompatibility 2, class II antigen E beta [Source:MGI Symbol;Acc:MGI:95901] | 1.23 | 4.81E-03 |
| Ppara | peroxisome proliferator activated receptor alpha [Source:MGI Symbol;Acc:MGI:104740] | 1.60 | 5.00E-03 |
| Postn | periostin, osteoblast specific factor [Source:MGI Symbol;Acc:MGI:1926321] | 1.52 | 5.00E-03 |
| Ampd3 | adenosine monophosphate deaminase 3 [Source:MGI Symbol;Acc:MGI:1096344] | 1.41 | 5.00E-03 |
| Btg2 | BTG anti-proliferation factor 2 [Source:MGI Symbol;Acc:MGI:108384] | 1.24 | 5.00E-03 |
| Mt1 | metallothionein 1 [Source:MGI Symbol;Acc:MGI:97171] | 2.13 | 5.01E-03 |
| Arpp21 | cyclic AMP-regulated phosphoprotein, 21 [Source:MGI Symbol;Acc:MGI:107562] | 1.81 | 5.70E-03 |
| Parp9 | poly (ADP-ribose) polymerase family, member 9 [Source:MGI Symbol;Acc:MGI:1933117] | 1.56 | 6.02E-03 |
| Mgp | matrix Gla protein [Source:MGI Symbol;Acc:MGI:96976] | 1.22 | 6.08E-03 |
| Rhou | ras homolog family member U [Source:MGI Symbol;Acc:MGI:1916831] | 1.14 | 6.08E-03 |
| Ephx1 | epoxide hydrolase 1, microsomal [Source:MGI Symbol;Acc:MGI:95405] | 1.15 | 7.76E-03 |
| Ptprc | protein tyrosine phosphatase receptor type C [Source:MGI Symbol;Acc:MGI:97810] | 1.50 | 8.40E-03 |
| Dnaja4 | DnaJ heat shock protein family (Hsp40) member A4 [Source:MGI Symbol;Acc:MGI:1927638] | 1.11 | 8.53E-03 |
| Gas6 | growth arrest specific 6 [Source:MGI Symbol;Acc:MGI:95660] | 0.65 | 8.53E-03 |
| Fkbp5 | FK506 binding protein 5 [Source:MGI Symbol;Acc:MGI:104670] | 1.16 | 8.72E-03 |
| Zfp865 | zinc finger protein 865 [Source:MGI Symbol;Acc:MGI:2442656] | 1.51 | 8.78E-03 |
| Pdp2 | pyruvate dehydrogenase phosphatase catalytic subunit 2 [Source:MGI Symbol;Acc:MGI:1918878] | 1.45 | 9.33E-03 |
| Usp49 | ubiquitin specific peptidase 49 [Source:MGI Symbol;Acc:MGI:2685391] | 1.26 | 1.01E-02 |
| Pcnt | pericentrin (kendrin) [Source:MGI Symbol;Acc:MGI:102722] | 0.64 | 1.05E-02 |
| Pdlim5 | PDZ and LIM domain 5 [Source:MGI Symbol;Acc:MGI:1927489] | 0.62 | 1.20E-02 |
| Hspa1a | heat shock protein 1A [Source:MGI Symbol;Acc:MGI:96244] | 2.86 | 1.36E-02 |
| Armcx4 | armadillo repeat containing, X-linked 4 [Source:MGI Symbol;Acc:MGI:2147887] | 1.18 | 1.36E-02 |
| Cd36 | CD36 molecule [Source:MGI Symbol;Acc:MGI:107899] | 1.07 | 1.36E-02 |
| Cdk17 | cyclin dependent kinase 17 [Source:MGI Symbol;Acc:MGI:97517] | 1.30 | 1.43E-02 |
| Grin2b | glutamate receptor, ionotropic, NMDA2B (epsilon 2) [Source:MGI Symbol;Acc:MGI:95821] | 3.38 | 1.52E-02 |
| Cspg4b | chondroitin sulfate proteoglycan 4B [Source:MGI Symbol;Acc:MGI:3040697] | 2.26 | 1.52E-02 |
| Egln3 | egl-9 family hypoxia-inducible factor 3 [Source:MGI Symbol;Acc:MGI:1932288] | 1.58 | 1.52E-02 |
| Lyz2 | lysozyme 2 [Source:MGI Symbol;Acc:MGI:96897] | 1.22 | 1.52E-02 |
| Dgat2 | diacylglycerol O-acyltransferase 2 [Source:MGI Symbol;Acc:MGI:1915050] | 1.44 | 1.74E-02 |
| Slc25a34 | solute carrier family 25, member 34 [Source:MGI Symbol;Acc:MGI:2686215] | 2.10 | 1.76E-02 |
| Gabarapl1 | GABA type A receptor associated protein like 1 [Source:MGI Symbol;Acc:MGI:1914980] | 1.04 | 2.00E-02 |
| Tra2a | transformer 2 alpha [Source:MGI Symbol;Acc:MGI:1933972] | 0.82 | 2.00E-02 |
| Lyrm7 | LYR motif containing 7 [Source:MGI Symbol;Acc:MGI:1922780] | 1.48 | 2.22E-02 |
| Hsph1 | heat shock 105kDa/110kDa protein 1 [Source:MGI Symbol;Acc:MGI:105053] | 0.71 | 2.23E-02 |
| Atp2a2 | ATPase, Ca++ transporting, cardiac muscle, slow twitch 2 [Source:MGI Symbol;Acc:MGI:88110] | 4.08 | 2.27E-02 |
| Gm12338 | predicted gene 12338 [Source:MGI Symbol;Acc:MGI:3650622] | 1.64 | 2.27E-02 |
| Ctla2a | cytotoxic T lymphocyte-associated protein 2 alpha [Source:MGI Symbol;Acc:MGI:88554] | 1.56 | 2.27E-02 |
| Ucp3 | uncoupling protein 3 (mitochondrial, proton carrier) [Source:MGI Symbol;Acc:MGI:1099787] | 0.94 | 2.27E-02 |
| Barx2 | BarH-like homeobox 2 [Source:MGI Symbol;Acc:MGI:109617] | 2.64 | 2.64E-02 |
| Sulf2 | sulfatase 2 [Source:MGI Symbol;Acc:MGI:1919293] | 1.14 | 2.64E-02 |
| Acad10 | acyl-Coenzyme A dehydrogenase family, member 10 [Source:MGI Symbol;Acc:MGI:1919235] | 1.48 | 2.70E-02 |
| Fndc5 | fibronectin type III domain containing 5 [Source:MGI Symbol;Acc:MGI:1917614] | 1.02 | 2.76E-02 |
| Sema3b | sema domain, immunoglobulin domain (Ig), short basic domain, secreted, (semaphorin) 3B [Source:MGI Symbol;Acc:MGI:107561] | 1.14 | 2.84E-02 |
| Bdh1 | 3-hydroxybutyrate dehydrogenase, type 1 [Source:MGI Symbol;Acc:MGI:1919161] | 3.39 | 2.87E-02 |
| C1qc | complement component 1, q subcomponent, C chain [Source:MGI Symbol;Acc:MGI:88225] | 1.46 | 2.87E-02 |
| Slc4a3 | solute carrier family 4 (anion exchanger), member 3 [Source:MGI Symbol;Acc:MGI:109350] | 1.46 | 2.88E-02 |
| Cdc14a | CDC14 cell division cycle 14A [Source:MGI Symbol;Acc:MGI:2442676] | 0.80 | 3.03E-02 |
| Mpeg1 | macrophage expressed gene 1 [Source:MGI Symbol;Acc:MGI:1333743] | 1.64 | 3.03E-02 |
| Apol6 | apolipoprotein L 6 [Source:MGI Symbol;Acc:MGI:1919189] | 1.42 | 3.03E-02 |
| Cep250 | centrosomal protein 250 [Source:MGI Symbol;Acc:MGI:108084] | 0.94 | 3.03E-02 |
| Nlrc3 | NLR family, CARD domain containing 3 [Source:MGI Symbol;Acc:MGI:2444070] | 1.92 | 3.15E-02 |
| Idh2 | isocitrate dehydrogenase 2 (NADP+), mitochondrial [Source:MGI Symbol;Acc:MGI:96414] | 2.23 | 3.16E-02 |
| Laptm5 | lysosomal-associated protein transmembrane 5 [Source:MGI Symbol;Acc:MGI:108046] | 1.39 | 3.24E-02 |
| Adamts9 | ADAM metallopeptidase with thrombospondin type 1 motif 9 [Source:MGI Symbol;Acc:MGI:1916320] | 0.86 | 3.24E-02 |
| Per2 | period circadian clock 2 [Source:MGI Symbol;Acc:MGI:1195265] | 0.95 | 3.39E-02 |
| Mt2 | metallothionein 2 [Source:MGI Symbol;Acc:MGI:97172] | 2.49 | 3.45E-02 |
| Ctss | cathepsin S [Source:MGI Symbol;Acc:MGI:107341] | 1.67 | 3.46E-02 |
| Hspb1 | heat shock protein 1 [Source:MGI Symbol;Acc:MGI:96240] | 0.97 | 3.46E-02 |
| Rrad | Ras-related associated with diabetes [Source:MGI Symbol;Acc:MGI:1930943] | 2.37 | 3.50E-02 |
| G0s2 | G0/G1 switch gene 2 [Source:MGI Symbol;Acc:MGI:1316737] | 1.72 | 3.62E-02 |
| Rcan1 | regulator of calcineurin 1 [Source:MGI Symbol;Acc:MGI:1890564] | 1.13 | 3.63E-02 |
| Myom3 | myomesin family, member 3 [Source:MGI Symbol;Acc:MGI:2685280] | 3.48 | 3.64E-02 |
| Osgin1 | oxidative stress induced growth inhibitor 1 [Source:MGI Symbol;Acc:MGI:1919089] | 1.18 | 3.64E-02 |
| Sbk3 | SH3 domain binding kinase family, member 3 [Source:MGI Symbol;Acc:MGI:2685924] | 3.97 | 3.72E-02 |

| | | | |
|---|---|---|---|
| Adamts20 | ADAM metallopeptidase with thrombospondin type 1 motif 20 [Source:MGI Symbol;Acc:MGI:2660628] | 1.49 | 3.72E-02 |
| Ldhb | lactate dehydrogenase B [Source:MGI Symbol;Acc:MGI:96763] | 2.82 | 3.79E-02 |
| Plekhh1 | pleckstrin homology domain containing, family H (with MyTH4 domain) member 1 [Source:MGI Symbol;Acc:MGI:2144989] | 2.34 | 3.79E-02 |
| Pros1 | protein S (alpha) [Source:MGI Symbol;Acc:MGI:1095733] | 1.23 | 3.79E-02 |
| Arid5a | AT-rich interaction domain 5A [Source:MGI Symbol;Acc:MGI:2443039] | 1.04 | 3.79E-02 |
| Atp1a1 | ATPase, Na+/K+ transporting, alpha 1 polypeptide [Source:MGI Symbol;Acc:MGI:88105] | 0.83 | 3.79E-02 |
| Ly6e | lymphocyte antigen 6 family member E [Source:MGI Symbol;Acc:MGI:106651] | 0.80 | 3.79E-02 |
| Sox9 | SRY (sex determining region Y)-box 9 [Source:MGI Symbol;Acc:MGI:98371] | 1.86 | 3.89E-02 |
| Ms4a6b | membrane-spanning 4-domains, subfamily A, member 6B [Source:MGI Symbol;Acc:MGI:1917024] | 3.12 | 4.05E-02 |
| Gm43302 | predicted gene 43302 [Source:MGI Symbol;Acc:MGI:5663439] | 2.42 | 4.05E-02 |
| Tecpr2 | tectonin beta-propeller repeat containing 2 [Source:MGI Symbol;Acc:MGI:2144865] | 0.77 | 4.05E-02 |
| Siglec1 | sialic acid binding Ig-like lectin 1, sialoadhesin [Source:MGI Symbol;Acc:MGI:99668] | 1.60 | 4.22E-02 |
| Ednrb | endothelin receptor type B [Source:MGI Symbol;Acc:MGI:102720] | 1.14 | 4.22E-02 |
| Tnni1 | troponin I, skeletal, slow 1 [Source:MGI Symbol;Acc:MGI:105073] | 5.39 | 4.25E-02 |
| Myl3 | myosin, light polypeptide 3 [Source:MGI Symbol;Acc:MGI:97268] | 4.26 | 4.25E-02 |
| Parp4 | poly (ADP-ribose) polymerase family, member 4 [Source:MGI Symbol;Acc:MGI:2685589] | 0.77 | 4.25E-02 |
| Ly6c1 | lymphocyte antigen 6 family member C1 [Source:MGI Symbol;Acc:MGI:96882] | 0.77 | 4.27E-02 |
| Myoz2 | myozenin 2 [Source:MGI Symbol;Acc:MGI:1913063] | 3.77 | 4.30E-02 |
| Grk5 | G protein-coupled receptor kinase 5 [Source:MGI Symbol;Acc:MGI:109161] | 1.23 | 4.36E-02 |
| Klhl34 | kelch-like 34 [Source:MGI Symbol;Acc:MGI:2685234] | 1.93 | 4.36E-02 |
| St3gal5 | ST3 beta-galactoside alpha-2,3-sialyltransferase 5 [Source:MGI Symbol;Acc:MGI:1339963] | 0.93 | 4.37E-02 |
| Enah | ENAH actin regulator [Source:MGI Symbol;Acc:MGI:108360] | 1.11 | 4.43E-02 |
| Fosb | FBJ osteosarcoma oncogene B [Source:MGI Symbol;Acc:MGI:95575] | 2.56 | 4.63E-02 |
| Cyfip2 | cytoplasmic FMR1 interacting protein 2 [Source:MGI Symbol;Acc:MGI:1924134] | 3.86 | 4.82E-02 |
| Plbd1 | phospholipase B domain containing 1 [Source:MGI Symbol;Acc:MGI:1914107] | 1.30 | 4.82E-02 |
| Avil | advillin [Source:MGI Symbol;Acc:MGI:1333798] | 1.06 | 4.82E-02 |
| Thbs1 | thrombospondin 1 [Source:MGI Symbol;Acc:MGI:98737] | 1.04 | 4.82E-02 |
| Amotl2 | angiomotin-like 2 [Source:MGI Symbol;Acc:MGI:1929286] | 0.78 | 4.94E-02 |

### Table S6. Downregulated genes in CONV-RT mice muscle compared to CNT

| Gene ID | Name | log2(FC) | padj |
|---|---|---|---|
| Mtss2 | MTSS I-BAR domain containing 2 [Source:MGI Symbol;Acc:MGI:3039591] | -1.11 | 7.17E-07 |
| Nrep | neuronal regeneration related protein [Source:MGI Symbol;Acc:MGI:99444] | -1.43 | 1.04E-06 |
| Slc37a4 | solute carrier family 37 (glucose-6-phosphate transporter), member 4 [Source:MGI Symbol;Acc:MGI:1316650] | -1.09 | 2.99E-06 |
| Nr1d1 | nuclear receptor subfamily 1, group D, member 1 [Source:MGI Symbol;Acc:MGI:2444210] | -1.46 | 1.11E-05 |
| Sorbs2 | sorbin and SH3 domain containing 2 [Source:MGI Symbol;Acc:MGI:1924574] | -0.89 | 1.63E-05 |
| Vldlr | very low density lipoprotein receptor [Source:MGI Symbol;Acc:MGI:98935] | -0.84 | 3.57E-05 |
| Auts2 | autism susceptibility candidate 2 [Source:MGI Symbol;Acc:MGI:1919847] | -1.20 | 1.02E-04 |
| Sms | spermine synthase [Source:MGI Symbol;Acc:MGI:109490] | -0.91 | 1.04E-04 |
| Bhlhe41 | basic helix-loop-helix family, member e41 [Source:MGI Symbol;Acc:MGI:1930704] | -0.78 | 2.15E-04 |
| Mylk2 | myosin, light polypeptide kinase 2, skeletal muscle [Source:MGI Symbol;Acc:MGI:2139434] | -0.81 | 2.18E-04 |
| Stat5b | signal transducer and activator of transcription 5B [Source:MGI Symbol;Acc:MGI:103035] | -0.78 | 2.26E-04 |
| Kcmf1 | potassium channel modulatory factor 1 [Source:MGI Symbol;Acc:MGI:1921537] | -0.71 | 3.33E-04 |
| Slc41a3 | solute carrier family 41, member 3 [Source:MGI Symbol;Acc:MGI:1918949] | -0.98 | 4.26E-04 |
| Htra4 | HtrA serine peptidase 4 [Source:MGI Symbol;Acc:MGI:3036260] | -2.40 | 7.10E-04 |
| Igsf3 | immunoglobulin superfamily, member 3 [Source:MGI Symbol;Acc:MGI:1926158] | -1.41 | 1.67E-03 |
| Dyrk2 | dual-specificity tyrosine phosphorylation regulated kinase 2 [Source:MGI Symbol;Acc:MGI:1330301] | -0.77 | 3.25E-03 |
| Ampd1 | adenosine monophosphate deaminase 1 [Source:MGI Symbol;Acc:MGI:88015] | -0.86 | 3.78E-03 |
| Asph | aspartate-beta-hydroxylase [Source:MGI Symbol;Acc:MGI:1914186] | -0.75 | 4.43E-03 |
| Mllt11 | myeloid/lymphoid or mixed-lineage leukemia; translocated to, 11 [Source:MGI Symbol;Acc:MGI:1929671] | -0.80 | 4.81E-03 |
| Dusp10 | dual specificity phosphatase 10 [Source:MGI Symbol;Acc:MGI:1927070] | -1.12 | 5.00E-03 |
| Cacna2d4 | calcium channel, voltage-dependent, alpha 2/delta subunit 4 [Source:MGI Symbol;Acc:MGI:2442632] | -1.54 | 5.00E-03 |
| Mafa | MAF bZIP transcription factor A [Source:MGI Symbol;Acc:MGI:2673307] | -0.89 | 5.36E-03 |
| Smyd2 | SET and MYND domain containing 2 [Source:MGI Symbol;Acc:MGI:1915889] | -0.93 | 5.36E-03 |
| Fhod3 | formin homology 2 domain containing 3 [Source:MGI Symbol;Acc:MGI:1925847] | -0.71 | 5.38E-03 |
| Tnrc18 | trinucleotide repeat containing 18 [Source:MGI Symbol;Acc:MGI:3648294] | -0.65 | 5.70E-03 |
| Nos1 | nitric oxide synthase 1, neuronal [Source:MGI Symbol;Acc:MGI:97360] | -1.15 | 5.70E-03 |
| Ntmt2 | N-terminal Xaa-Pro-Lys N-methyltransferase 2 [Source:MGI Symbol;Acc:MGI:2685053] | -1.07 | 6.16E-03 |
| Pdgfa | platelet derived growth factor, alpha [Source:MGI Symbol;Acc:MGI:97527] | -0.96 | 6.21E-03 |
| Cmbl | carboxymethylenebutenolidase homolog [Source:MGI Symbol;Acc:MGI:1916824] | -1.01 | 6.23E-03 |
| Ttyh2 | tweety family member 2 [Source:MGI Symbol;Acc:MGI:2157091] | -0.99 | 6.52E-03 |
| Casq1 | calsequestrin 1 [Source:MGI Symbol;Acc:MGI:1309468] | -0.68 | 6.54E-03 |
| Reep1 | receptor accessory protein 1 [Source:MGI Symbol;Acc:MGI:1098827] | -0.78 | 8.18E-03 |
| H60b | histocompatibility 60b [Source:MGI Symbol;Acc:MGI:3649078] | -0.97 | 8.18E-03 |
| Zyg11b | zyg-ll family member B, cell cycle regulator [Source:MGI Symbol;Acc:MGI:2685277] | -0.60 | 8.40E-03 |

| | | | |
|---|---|---|---|
| Msrb3 | methionine sulfoxide reductase B3 [Source:MGI Symbol;Acc:MGI:2443538] | -0.59 | 8.53E-03 |
| Slc16a10 | solute carrier family 16 (monocarboxylic acid transporters), member 10 [Source:MGI Symbol;Acc:MGI:1919722] | -0.66 | 8.53E-03 |
| Gdap1 | ganglioside-induced differentiation-associated-protein 1 [Source:MGI Symbol;Acc:MGI:1338002] | -1.12 | 8.72E-03 |
| Smtnl2 | smoothelin-like 2 [Source:MGI Symbol;Acc:MGI:2442764] | -0.80 | 1.05E-02 |
| Dmxl1 | Dmx-like 1 [Source:MGI Symbol;Acc:MGI:2443926] | -1.01 | 1.10E-02 |
| Shisa4 | shisa family member 4 [Source:MGI Symbol;Acc:MGI:1924802] | -0.62 | 1.23E-02 |
| Gm57854 | predicted gene, 57854 [Source:MGI Symbol;Acc:MGI:7512655] | -0.82 | 1.36E-02 |
| St8sia5 | ST8 alpha-N-acetyl-neuraminide alpha-2,8-sialyltransferase 5 [Source:MGI Symbol;Acc:MGI:109243] | -0.90 | 1.40E-02 |
| Wnk3 | WNK lysine deficient protein kinase 3 [Source:MGI Symbol;Acc:MGI:2652875] | -1.67 | 1.42E-02 |
| Art3 | ADP-ribosyltransferase 3 [Source:MGI Symbol;Acc:MGI:1202729] | -0.63 | 1.54E-02 |
| Srcin1 | SRC kinase signaling inhibitor 1 [Source:MGI Symbol;Acc:MGI:1933179] | -1.74 | 1.54E-02 |
| Stx17 | syntaxin 17 [Source:MGI Symbol;Acc:MGI:1914977] | -0.78 | 1.85E-02 |
| Pcca | propionyl-Coenzyme A carboxylase, alpha polypeptide [Source:MGI Symbol;Acc:MGI:97499] | -0.77 | 1.88E-02 |
| Pld5 | phospholipase D family member 5 [Source:MGI Symbol;Acc:MGI:2442056] | -1.48 | 1.89E-02 |
| Atp13a5 | ATPase type 13A5 [Source:MGI Symbol;Acc:MGI:2444068] | -1.46 | 2.09E-02 |
| Ttll7 | tubulin tyrosine ligase-like family, member 7 [Source:MGI Symbol;Acc:MGI:1918142] | -0.67 | 2.25E-02 |
| Lbx1 | ladybird homeobox 1 [Source:MGI Symbol;Acc:MGI:104867] | -0.68 | 2.27E-02 |
| Peg3 | paternally expressed 3 [Source:MGI Symbol;Acc:MGI:104748] | -1.26 | 2.39E-02 |
| Lgalsl | galectin like [Source:MGI Symbol;Acc:MGI:1916114] | -0.63 | 2.48E-02 |
| Pkdcc | protein kinase domain containing, cytoplasmic [Source:MGI Symbol;Acc:MGI:2147077] | -0.68 | 2.64E-02 |
| Rgmb | repulsive guidance molecule family member B [Source:MGI Symbol;Acc:MGI:1916049] | -0.80 | 2.64E-02 |
| Gadd45b | growth arrest and DNA-damage-inducible 45 beta [Source:MGI Symbol;Acc:MGI:107776] | -1.06 | 2.64E-02 |
| Cdh23 | cadherin related 23 (otocadherin) [Source:MGI Symbol;Acc:MGI:1890219] | -1.36 | 2.64E-02 |
| Aldh1a1 | aldehyde dehydrogenase family 1, subfamily A1 [Source:MGI Symbol;Acc:MGI:1353450] | -0.85 | 2.88E-02 |
| Klhdc3 | kelch domain containing 3 [Source:MGI Symbol;Acc:MGI:2651568] | -0.64 | 3.09E-02 |
| Ptgfr | prostaglandin F receptor [Source:MGI Symbol;Acc:MGI:97796] | -2.13 | 3.31E-02 |
| Nudt4 | nudix hydrolase 4 [Source:MGI Symbol;Acc:MGI:1918457] | -0.62 | 3.45E-02 |
| Spred3 | sprouty-related EVH1 domain containing 3 [Source:MGI Symbol;Acc:MGI:2142186] | -1.62 | 3.48E-02 |
| Vwa7 | von Willebrand factor A domain containing 7 [Source:MGI Symbol;Acc:MGI:1306798] | -0.84 | 3.70E-02 |
| Fam53a | family with sequence similarity 53, member A [Source:MGI Symbol;Acc:MGI:1919225] | -0.63 | 3.74E-02 |
| Pvalb | parvalbumin [Source:MGI Symbol;Acc:MGI:97821] | -0.66 | 3.79E-02 |
| C7 | complement component 7 [Source:MGI Symbol;Acc:MGI:88235] | -1.42 | 3.79E-02 |
| Bmp5 | bone morphogenetic protein 5 [Source:MGI Symbol;Acc:MGI:88181] | -1.69 | 3.96E-02 |
| Mettl21c | methyltransferase 21C, AARS1 lysine [Source:MGI Symbol;Acc:MGI:3611450] | -1.01 | 4.63E-02 |
| Asb14 | ankyrin repeat and SOCS box-containing 14 [Source:MGI Symbol;Acc:MGI:2655107] | -0.71 | 4.67E-02 |
| Lrtm2 | leucine-rich repeats and transmembrane domains 2 [Source:MGI Symbol;Acc:MGI:2141485] | -2.29 | 4.94E-02 |

### Table S7. Upregulated genes in FLASH-RT mice muscle compared to CNT

| Gene ID | Name | log2(FC) | padj |
|---|---|---|---|
| Pdk4 | pyruvate dehydrogenase kinase, isoenzyme 4 [Source:MGI Symbol;Acc:MGI:1351481] | 1.68 | 3.74E-10 |
| Lmod2 | leiomodin 2 (cardiac) [Source:MGI Symbol;Acc:MGI:2135672] | 1.61 | 3.37E-08 |
| Cdkn1a | cyclin dependent kinase inhibitor 1A [Source:MGI Symbol;Acc:MGI:104556] | 2.38 | 3.60E-06 |
| Bdh1 | 3-hydroxybutyrate dehydrogenase, type 1 [Source:MGI Symbol;Acc:MGI:1919161] | 2.50 | 6.98E-06 |
| Abra | actin-binding Rho activating protein [Source:MGI Symbol;Acc:MGI:2444891] | 0.79 | 6.98E-06 |
| Ucp3 | uncoupling protein 3 (mitochondrial, proton carrier) [Source:MGI Symbol;Acc:MGI:1099787] | 1.04 | 1.59E-05 |
| Idh2 | isocitrate dehydrogenase 2 (NADP+), mitochondrial [Source:MGI Symbol;Acc:MGI:96414] | 1.29 | 9.14E-05 |
| Grk5 | G protein-coupled receptor kinase 5 [Source:MGI Symbol;Acc:MGI:109161] | 1.46 | 1.44E-04 |
| Lpl | lipoprotein lipase [Source:MGI Symbol;Acc:MGI:96820] | 1.15 | 1.73E-04 |
| Sorbs1 | sorbin and SH3 domain containing 1 [Source:MGI Symbol;Acc:MGI:700014] | 0.73 | 1.32E-03 |
| Myh7b | myosin, heavy chain 7B, cardiac muscle, beta [Source:MGI Symbol;Acc:MGI:3710243] | 2.43 | 1.80E-03 |
| Lgi1 | leucine-rich repeat LGI family, member 1 [Source:MGI Symbol;Acc:MGI:1861691] | 2.77 | 1.82E-03 |
| H2-Aa | histocompatibility 2, class II antigen A, alpha [Source:MGI Symbol;Acc:MGI:95895] | 1.39 | 2.96E-03 |
| H2-Eb1 | histocompatibility 2, class II antigen E beta [Source:MGI Symbol;Acc:MGI:95901] | 1.33 | 5.18E-03 |
| Depp1 | DEPP1 autophagy regulator [Source:MGI Symbol;Acc:MGI:1918730] | 2.17 | 5.83E-03 |
| Pank1 | pantothenate kinase 1 [Source:MGI Symbol;Acc:MGI:1922985] | 1.36 | 6.65E-03 |
| Slc25a34 | solute carrier family 25, member 34 [Source:MGI Symbol;Acc:MGI:2686215] | 1.99 | 6.99E-03 |
| Cd74 | CD74 antigen (invariant polypeptide of major histocompatibility complex, class II antigen-associated) [Source:MGI Symbol;Acc:MGI:96534] | 1.29 | 8.53E-03 |
| Mup22 | major urinary protein 22 [Source:MGI Symbol;Acc:MGI:5434675] | 6.11 | 8.68E-03 |
| Lyz2 | lysozyme 2 [Source:MGI Symbol;Acc:MGI:96897] | 1.32 | 8.68E-03 |
| Ldhb | lactate dehydrogenase B [Source:MGI Symbol;Acc:MGI:96763] | 1.96 | 9.33E-03 |
| Ccr2 | C-C motif chemokine receptor 2 [Source:MGI Symbol;Acc:MGI:106185] | 2.01 | 1.75E-02 |
| C1qc | complement component 1, q subcomponent, C chain [Source:MGI Symbol;Acc:MGI:88225] | 1.42 | 1.75E-02 |
| Actn2 | actinin alpha 2 [Source:MGI Symbol;Acc:MGI:109192] | 0.98 | 1.75E-02 |
| Gm15542 | predicted gene 15542 [Source:MGI Symbol;Acc:MGI:3782990] | 5.59 | 2.66E-02 |
| Slc27a1 | solute carrier family 27 (fatty acid transporter), member 1 [Source:MGI Symbol;Acc:MGI:1347098] | 0.91 | 2.66E-02 |
| Myadm | myeloid-associated differentiation marker [Source:MGI Symbol;Acc:MGI:1355332] | 0.81 | 2.66E-02 |

| | | | |
|---|---|---|---|
| Cblb | Casitas B-lineage lymphoma b [Source:MGI Symbol;Acc:MGI:2146430] | 0.63 | 2.66E-02 |
| Pdlim1 | PDZ and LIM domain 1 (elfin) [Source:MGI Symbol;Acc:MGI:1860611] | 1.51 | 2.66E-02 |
| H2-Ab1 | histocompatibility 2, class II antigen A, beta 1 [Source:MGI Symbol;Acc:MGI:103070] | 1.36 | 2.70E-02 |
| Gabarapl1 | GABA type A receptor associated protein like 1 [Source:MGI Symbol;Acc:MGI:1914980] | 0.75 | 2.77E-02 |
| Psd4 | pleckstrin and Sec7 domain containing 4 [Source:MGI Symbol;Acc:MGI:2674093] | 3.01 | 3.58E-02 |
| Mpeg1 | macrophage expressed gene 1 [Source:MGI Symbol;Acc:MGI:1333743] | 1.75 | 3.70E-02 |
| Zdhhc23 | zinc finger, DHHC domain containing 23 [Source:MGI Symbol;Acc:MGI:2685625] | 1.57 | 3.70E-02 |
| Ctss | cathepsin S [Source:MGI Symbol;Acc:MGI:107341] | 1.76 | 4.19E-02 |
| Khdrbs3 | KH domain containing, RNA binding, signal transduction associated 3 [Source:MGI Symbol;Acc:MGI:1313312] | 1.42 | 4.48E-02 |
| Palmd | palmdelphin [Source:MGI Symbol;Acc:MGI:2148896] | 0.91 | 4.51E-02 |
| Shroom4 | shroom family member 4 [Source:MGI Symbol;Acc:MGI:2685570] | 0.85 | 4.88E-02 |
| Ly6e | lymphocyte antigen 6 family member E [Source:MGI Symbol;Acc:MGI:106651] | 0.65 | 4.88E-02 |
| Lrrc8c | leucine rich repeat containing 8 family, member C [Source:MGI Symbol;Acc:MGI:2140839] | 0.97 | 4.98E-02 |
| Apln | apelin [Source:MGI Symbol;Acc:MGI:1353624] | 2.47 | 4.98E-02 |

#### Table S8. Downregulated genes in FLASH-RT mice muscle compared to CNT

| Gene ID | Name | log2(FC) | padj |
|---|---|---|---|
| Bhlhe41 | basic helix-loop-helix family, member e41 [Source:MGI Symbol;Acc:MGI:1930704] | -0.73 | 6.98E-06 |
| Ttyh2 | tweety family member 2 [Source:MGI Symbol;Acc:MGI:2157091] | -1.02 | 1.57E-03 |
| Mafa | MAF bZIP transcription factor A [Source:MGI Symbol;Acc:MGI:2673307] | -0.77 | 2.95E-03 |
| Ganc | glucosidase, alpha; neutral C [Source:MGI Symbol;Acc:MGI:1923301] | -0.68 | 2.96E-03 |
| Pla2g7 | phospholipase A2, group VII (platelet-activating factor acetylhydrolase, plasma) [Source:MGI Symbol;Acc:MGI:1351327] | -0.70 | 3.06E-03 |
| Zdhhc18 | zinc finger, DHHC domain containing 18 [Source:MGI Symbol;Acc:MGI:3527792] | -0.85 | 5.18E-03 |
| Ttll7 | tubulin tyrosine ligase-like family, member 7 [Source:MGI Symbol;Acc:MGI:1918142] | -0.60 | 5.83E-03 |
| Crocc | ciliary rootlet coiled-coil, rootletin [Source:MGI Symbol;Acc:MGI:3529431] | -1.60 | 1.57E-02 |
| Tmem161b | transmembrane protein 161B [Source:MGI Symbol;Acc:MGI:1919995] | -1.36 | 2.66E-02 |
| Cmbl | carboxymethylenebutenolidase homolog [Source:MGI Symbol;Acc:MGI:1916824] | -0.69 | 2.70E-02 |
| Pld5 | phospholipase D family member 5 [Source:MGI Symbol;Acc:MGI:2442056] | -1.37 | 2.77E-02 |
| Mrtfa | myocardin related transcription factor A [Source:MGI Symbol;Acc:MGI:2384495] | -0.69 | 3.51E-02 |
| Pkhd1l1 | polycystic kidney and hepatic disease 1-like 1 [Source:MGI Symbol;Acc:MGI:2183153] | -2.82 | 4.99E-02 |

#### Table S9. Details of Probes and Primers for qPCR

| Target | Detection system | Probe code or primer sequences (5'-3') |
|---|---|---|
| TBP | probe | Mm 00446973_m1 |
| CCL2 | probe | Mm 00441242_m1 |
| IL1b | probe | Mm 00434228_m1 |
| IL6 | probe | Mm 00446191_m1 |
| Krt6A | SYBR green / primers | FW = GCAGACCAGCCTGACAGATTTC<br>RV = GTGGGCACTTCAGGGTTTTC |
| Krt6B | SYBR green / primers | FW = TGTAACCGTGAAAAAGGATGTAGA<br>RV = ACTCTCAGGAAGTTGATCTCGT |
| ADAM8 | SYBR green / primers | FW = CTGTGAATCAGGACCACTCCAA<br>RV = ACATTGGTCAGGCAGCCTGTCT |
| BCL3 | SYBR green / primers | FW = CCGGAGGCCCTTTACTACCA<br>RV = GGAGTAGGGGTGAGTAGGCAG |
| CDKN1A/p21 | SYBR green / primers | FW = GCAGACCAGCCTGACAGATTTC<br>RV = GTGGGCACTTCAGGGTTTTC |
| PAI-1/SerpinE1 | SYBR green / primers | FW = CAGAGCAACAAGTTCAACTACACTGA<br>RV = CAGCGATGAACATGCTGAGG |
